# ADP-ribose derived Nuclear ATP is Required for Chromatin Remodeling and Hormonal Gene Regulation (97 charact)

**DOI:** 10.1101/006593

**Authors:** Roni H. G. Wright, Francois LeDily, Daniel Soronellas, Andy Pohl, Jaume Bonet, A. S. Nacht, Guillermo P. Vicent, Michael Wierer, Baldo Oliva, Miguel Beato

## Abstract

**Highlights:**

– Hormonal gene regulation requires synthesis of PAR and its degradation to ADP-ribose by PARG
– ADP-ribose is converted to ATP in the cell nuclei by hormone-activated NUDIX5/NUDT5
– Blocking nuclear ATP formation precludes hormone-induced chromatin remodeling, gene regulation and cell proliferation

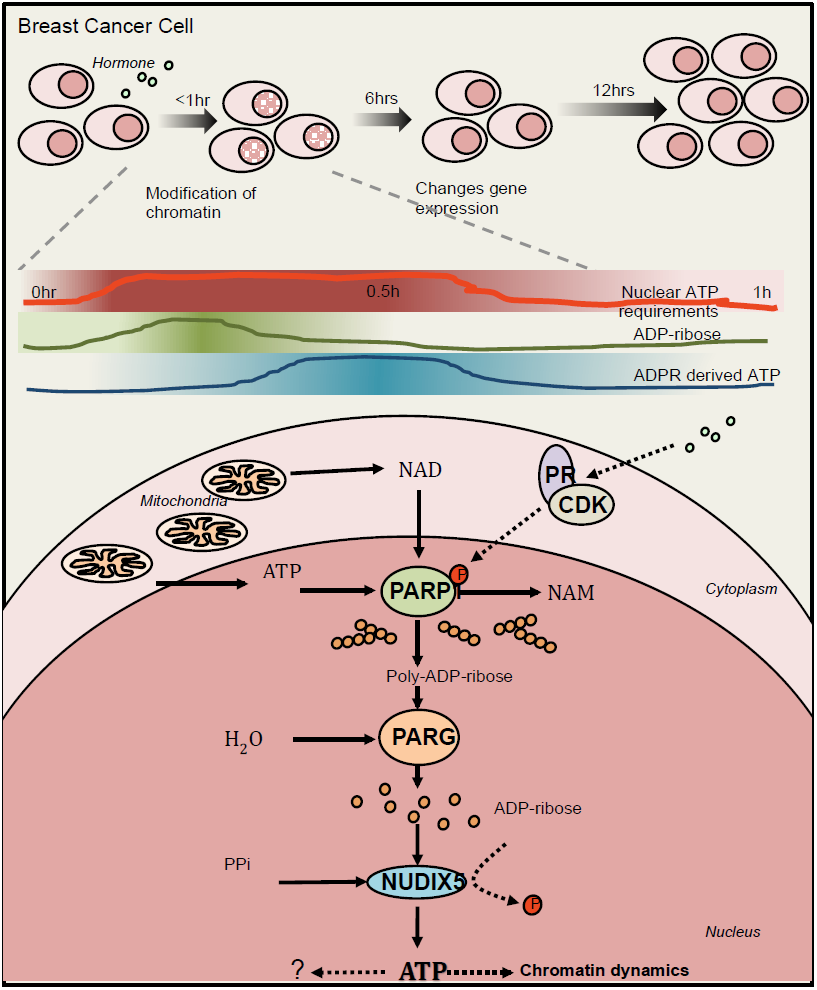

**Summary:** Key nuclear processes in eukaryotes including DNA replication or repair and gene regulation require extensive chromatin remodeling catalyzed by energy consuming enzymes. How the energetic demands of such processes are ensured in response to rapid stimuli remains unclear. We have analyzed this question in the context of the massive gene regulation changes induced by progestins in breast cancer cells and found that ATP is generated in the cell nucleus via the hydrolysis of poly-ADP-ribose to ADP-ribose. Nuclear ATP synthesis requires the combined enzymatic activities of PARP1, PARG and NUDIX5/NUDT5. Although initiated via mitochondrial derived ATP, the nuclear source of ATP is essential for hormone induced chromatin remodeling, gene regulation and cell proliferation and may also participate in DNA repair. This novel pathway reveals exciting avenues of research for drug development.

---

The nucleus of human cells requires an immense amount of energy to replicate or to repair the genome and to reprogram gene expression during differentiation or in response to external cues. Extensive changes in chromatin compaction and nucleosome organization are required to ensure gene accessibility for regulatory proteins and enzymes involved in these processes. First the chromatin fiber must be de-condensed followed by mobilization of the nucleosomes. Repositioning by sliding a single nucleosome core particle requires the breaking and reforming of hundreds of interactions between positively charged amino acids in histones and phosphates in the DNA backbone. A complete round of nucleosome remodeling *in vitro* requires the hydrolysis of ≈1000 ATP molecules by the ATPases/DNA-translocase domains that fuel the activity of dedicated ATP-dependent chromatin remodeling complexes ^1^. In the nucleus of living cells the energy cost may be even higher, due to the added complexity imposed by the folding of the chromatin fiber, linker histones and other chromosomal proteins.

The general assumption is that nuclear energetic demands are met via the diffusion of ATP from the mitochondria. Although this may be the case in steady state situations, the question remains about the source of energy for extensive or sudden changes in chromatin. Nearly 60 years ago Allfrey and Mirsky proposed that ATP could be generated in isolated nuclei ^2, 3^. It was suggested that the substrate for nuclear oxidative phosphorylation was in part generated by the ribose pathway ^4^. In 1989 an enzyme activity named ADP-ribose pyrophosphorylase was postulated in HeLa cell nuclei that catalyzed the formation of ATP and Ribose-5-phosphate from ADP-ribose (ADPR) and PPi ^5^. Moreover, ATP generation via ADPR catabolism has been shown to be involved in DNA repair and replication ^8–10^. However, whether nuclear ATP is also required for changes in gene expression is not known.

We have previously shown that in response to progestins (Pg) breast cancer cells undergo extensive changes in gene expression that require a global modification of chromatin, mediated by kinases, histone modifying enzymes and ATP-dependent chromatin remodelers ^9–12^. One key event in this response is the rapid and transient generation of poly-ADP-ribose (PAR) by PARP1 that consumes about 50% of the cellular NAD ^13^. We considered the possibility that the transient nature of PAR accumulation reflects the need for the degradation of PAR that may play a role in the energetic balance of the cell nucleus.

## Nuclear ATP levels increase in response to progesterone

To measure ATP levels in living T47D^M^ cells we utilized the ATeam constructs ^14^, a set of FRET-based ATP/ADP detectors targeted to cell nucleus, mitochondria or cytosol (Extended Figure 1A). ATP(YFP)/ADP^(CFP)^ emissions were measured at 1 minute intervals after hormone stimulation, the images were processed and the ratio of ATP/ADP determined as described ^15^ (Extended Figure 1B). The specificity and sensitivity of the ATeams were demonstrated using glucose stimulated Mito-ATeam in the presence or absence of the ATP-synthase inhibitor oligomycin. Levels of ATP increased rapidly (5–10 min) following the addition of glucose but the increase was inhibited by simultaneous treatment with oligomycin (Extended Figure 1C and D, respectively). Following addition of hormone to cells expressing the Nuclear-ATeam the nuclear ratio ATP^(YFP)^/ADP ^(CFP)^ increased rapidly to reached a low peak at 10–15 min followed by a much higher peak at 35–50 min, which was not found in solvent treated cells. Figure 1A shows snapshots at 5 min intervals of a representative cell expressing nuclear ATeam in response to hormone; Figure 1B shows quantification of 18 nuclei of cells incubated with hormone (left panel) or with solvent (right panel). Quantification of cells expressing the mitochondrial-ATeam did not show the hormone-dependent increase in ATP (Figure 1C).

**Figure 1:**
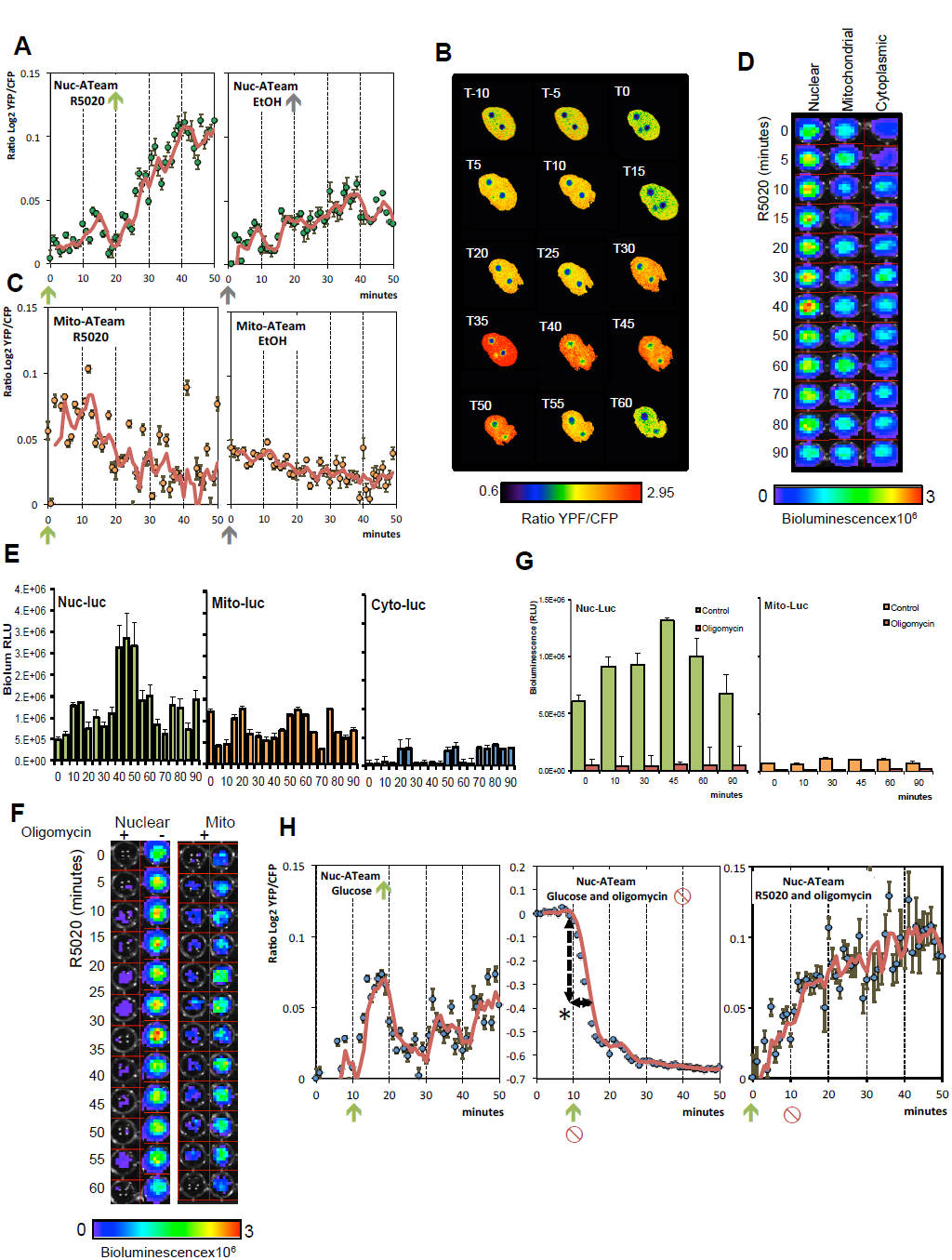
ATP levels increase in the nucleus 30–50 min after hormone following and initial rise of ATP coming from mitochondria. **A**. Representative ratio images (5 min intervals) of nuc-ATeam treated with hormone at T0. Scale bar on the bottom indicates ratio YFP/CFP **B**. ATP/ADP ratio quantification of Nuc-ATeam treated at T0 with hormone (R5020) or solvent (EtOH). Data is the average of 18 nuclear ROI’s +/−SEM, presented as log_2_^YFP/CFP^. **C**. ATP/ADP ratio quantification of Mito-ATeam treated with hormone (R5020, T0 arrow) or mock treated (EtOH alone). Data is the average of 16 nuclear ROI’s +/−SEM, presented as log_2_^YFP/CFP^. **D**. Bioluminescence image of cells transfected with Nuclear, Mitochondrial or Cytoplasmic targeted luciferase constructs and treated with hormone (R5020) for the time indicated. Scale bar represents the relative bioluminescence (RLU × 10^6^). **E**. Quantification of the data presented in (D); data is indicative of three independent experiments carried out in duplicate +/−SEM. **F**. Bioluminescence image of cells transfected with Nuclear or Mitochondrial targeted luciferase constructs. Cells were treated with hormone for the time indicated either with (+) or without (−) prior treatment with oligomycin. **G**. Quantification of the data presented in (F); data is indicative of three independent experiments carried out in duplicate +/−SEM. **H**. *Left panel*. ATP/ADP ratio quantification of Nuc-ATeam in cells treated with glucose (T10). Data is presented as log_2_^YFP/CFP^. Data is the average of 14 nuclear ROI’s +/−SEM, presented as log_2_^YFP/CFP^.*Central panel*. ATP/ADP ratio quantification of Nuc-ATeam in cells treated with glucose and oligomycin (T10). Data is presented as log_2_^YFP/CFP^. Data is the average of 8 nuclear ROI’s +/−SEM, presented as log_2_^YFP/CFP^. *Right panel*. ATP/ADP ratio quantification of Nuc-ATeam I cells treated with hormone (T0) and oligomycin (T10). Data is presented as log_2_^YFP/CFP^. Data is the average of 12 nuclear ROI’s +/−SEM, presented as log_2_^YFP/CFP^.

To confirm the FRET based results, we used nuclear, cytoplasmic and mitochondrial luciferase-based reporter constructs ^16^ to measure the levels of ATP by bioluminescence imaging (Extended Figure 1E and F). We confirmed the specific increase in nuclear ATP levels in cells treated with hormone, with a low peak at 10–15 min and a high peak at 40–50 minutes, which was not observed with the cytoplasmic or mitochondrial targeted constructs (Representative Bioluminescent image in Figure 1D and quantification in Figure 1E).

The increase in nuclear ATP was compromised by inhibiting mitochondrial ATP production with oligomycin prior to hormone addition, indicating that mitochondrial ATP is required for initiation of nuclear ATP generation (Figure 1F and G). In response to glucose the ATP levels in the nucleus increase rapidly (Figure 1H *left panel*), likely via diffusion of ATP from the mitochondria, as demonstrated by its inhibition by oligomycin (Figure 1H *middle panel*). The depletion of nuclear ATP in nuclei of cells treated with glucose and oligomycin demonstrates that the steady state nucleus contains a finite pool of ATP, which is depleted in around 10 min if it is not replenished (dashed line). However, in cells treated with hormone followed after 10 min by oligomycin, the nuclear levels of ATP continue to increase up to 40–50 min despite blocking of mitochondrial ATP production (Figure 1H *right panel*). We conclude that nuclear ATP generated after 20 minutes of hormone treatment is independent of mitochondrial ATP synthesis.

## PAR degradation to ADP-ribose is needed for nuclear ATP synthesis

Hormone induces a strong and transient increase in nuclear Poly-ADP-ribose (PAR) levels (Figure 2A *top panels*), due to the activation of PARP1 ^13^. Concomitantly with the PARP1 activation the degradation of PAR by PARG is also induced, as indicated by the opposite effect of PARP1 and PARG inhibition. While PARP1 inhibition prevents the accumulation of PAR, PARG inhibition results in persistence of elevated PAR levels (Figure 2A *bottom panels* and Figure 2B time kinetics, see also Extended Figure 2 A and B). In concert with PAR accumulation following hormone we observe a transient decrease in cellular NAD levels, which is dependent on PARP activity as demonstrated by the effect of PARP inhibition with 3AB (Figure 2C). Both PAR and NAD levels return to basal state after 60–80 min of hormone (Figure 2B and C). The recovery of NAD levels is dependent on PARG activity as shown with the selective inhibitor Tannic Acid, suggesting that ADPR is required (Figure 2C).

**Figure 2:**
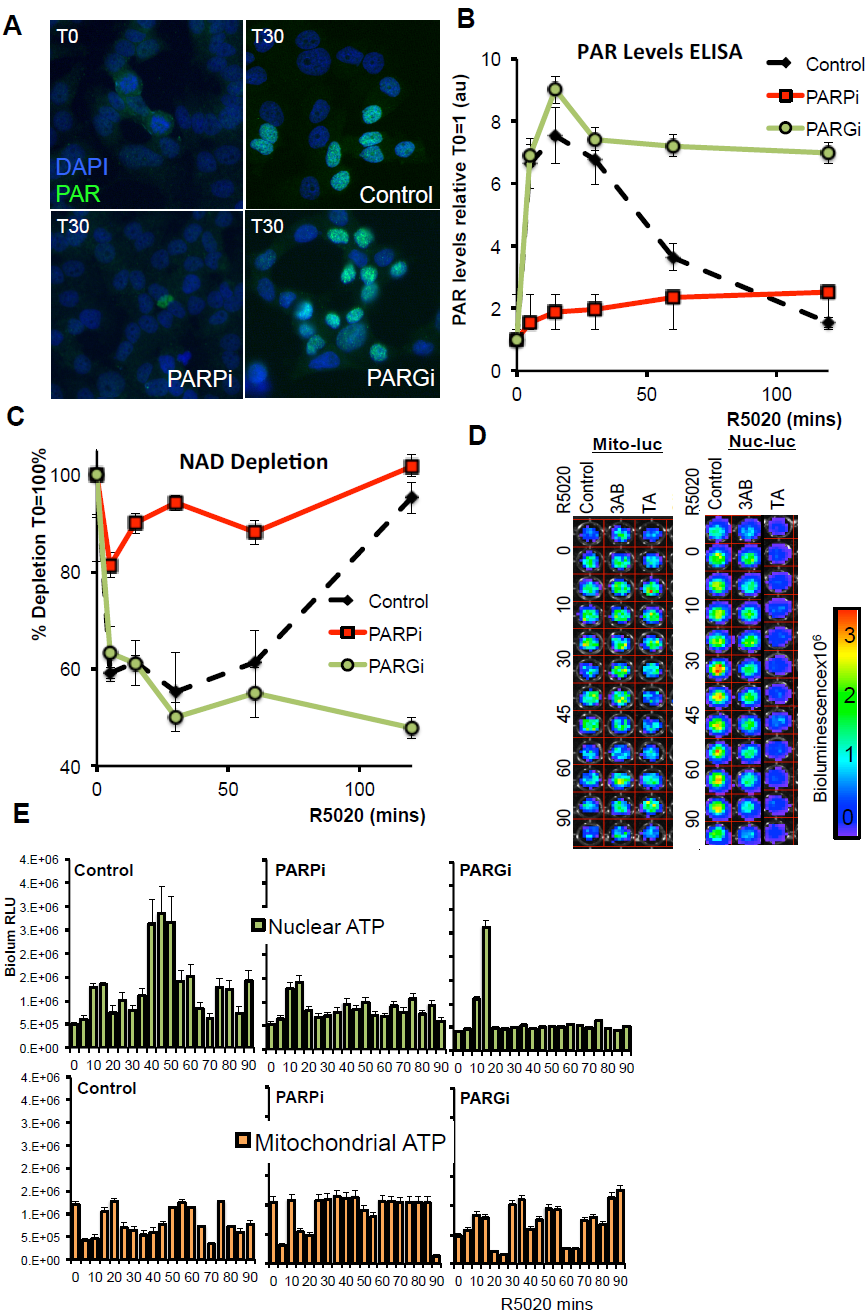
PAR degradation and identification of PARP1 target proteins. **A**. PAR levels were visualized by immunofluorescence using anti-PAR antibody in untreated cells (T0) and in cells treated with hormone for 30 minutes (T30) in the absence of inhibitors (control) or in the presence of either 3AB (PARPi) or TA (PARGi). **B**. PAR levels were determined by PAR-capture ELISA in cells treated with hormone in the absence (control) and in the presence of PARP or PARG inhibition (PARPi, PARGi) for the indicated times. **C**. NAD levels were determined and are shown as a percentage of depletion (T0=100%) in cells treated with hormone in the presence or absence of PARP or PARG inhibition (PARPi, PARGi) for the indicated times. **D**. Image of a representative luminescence assay of mitochondrial and nuclear ATP levels in T47Dwt cells treated with hormone for the indicated time periods (minutes) in the absence of inhibitors (Control) or in the presence of PARP inhibitors (3AB) or PARG inhibitor (TA). **E**. Quantitation of the data measurements as shown in D. The histograms represent the average and SEM of 3 experiments performed in duplicate.

To test whether PAR and its degradation to ADPR were needed for nuclear ATP synthesis we measured nuclear ATP in the presence of inhibitors of PAPR1 or PARG. We observed that specifically the high increase in nuclear ATP 40–50 min after hormone was dependent on PARP1 and PARG, while little changes were observed on mitochondrial or cytoplasmic ATP levels (Figure 2D and 2E, and Extended Figure 2C). These findings indicate that both the formation of PAR and its degradation to ADPR are needed for nuclear ATP generation.

## Role of NUDIX5/NUDT5 in nuclear ATP synthesis

To gain further insight into the role of PAR, we performed proteomics analysis of the PAR interactome in untreated control cells (Time 0) and in cells treated with hormone for 30 minutes (Figure 3A). Cell extracts were immunoprecipitated with an antibody against PAR and precipitated proteins were analyzed by mass spectrometry. We identified 1091 unique either PARylated or PAR binding proteins (Extended Table 1) most of them following hormone induction, including histones, kinases, phosphatases and proteins involved in both NAD and nucleotide metabolism (Extended Table 2, GO biological function analysis; Extended Tables 3 and 4), which have also been reported in other cell types, including H2A, H1, XRCC5 and PARP1 itself ^17^. Among the proteins involved in ADP and ADP-ribose metabolism we identified ARF3 (ADP-ribosylation factor 3), ADP-ribosylation factor-binding protein (GGA2) and ADP-sugar pyrophosphatase NUDT5, also known as NUDIX5. Of them only NUDT5/NUDIX5 peptides were enriched following hormone treatment. Members of the Nudix (Nucleoside Diphosphate linked to X) superfamily of enzymes are found in all classes of organisms and catalyze the hydrolysis of a wide range of substrates including nucleotide sugars (review ^18^). NUDIX5 is in the top 10% of genes overexpressed in breast cancer in correlation with PARP1 and PARG (Extended Figure 3A), a signature not shared by other members of the NUDIX family (Extended Figure 3B). NUDIX5 overexpression correlates with known PARP1 interacting proteins in invasive breast carcinoma compared to normal breast (p=7.4E-30) in addition to other cancer types (Extended Figure 3C, D).

**Figure 3:**
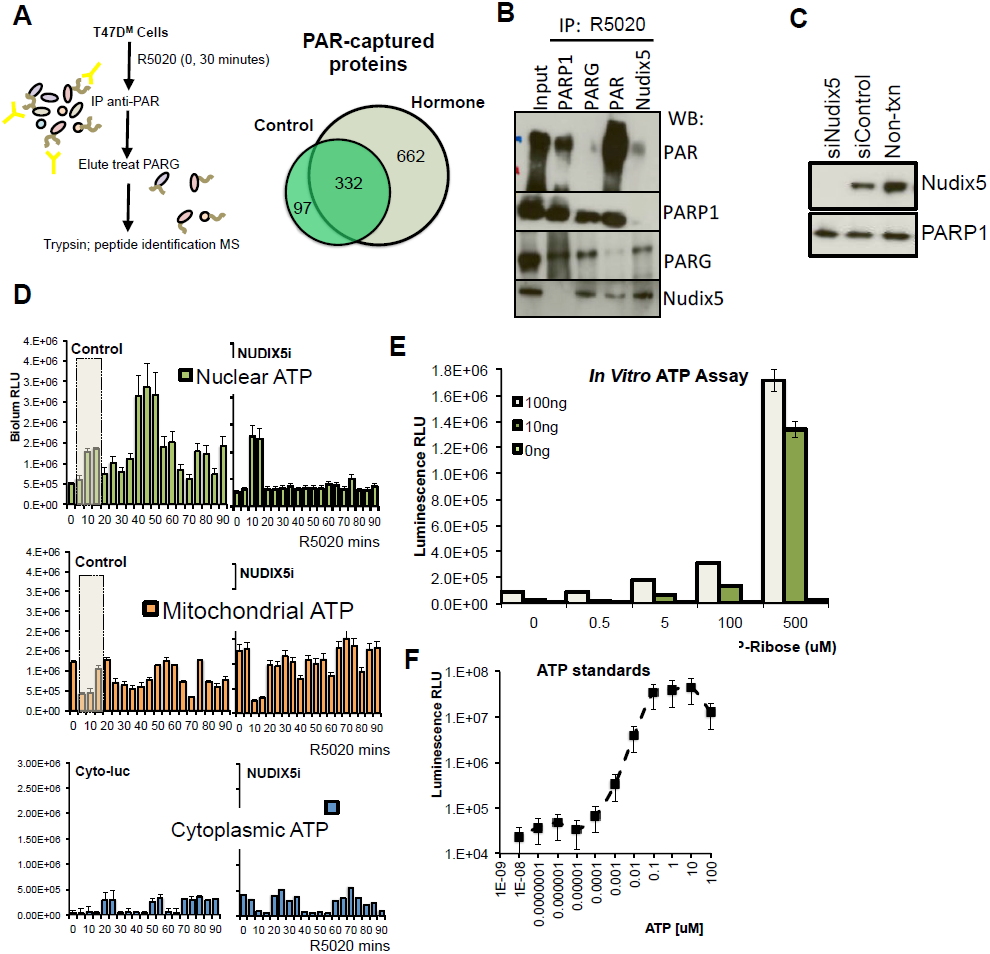
Novel PAR/PARG interactor Nudix5 plays a role in hormone induced nuclear ATP generation. **A**. Schematic representation of the method used to capture and identify PARP and PAR target protein in T47D cells. Immunoprecipitation, tryptic digest and mass spec identification was performed in duplicate and proteins were identified by at least 2 unique peptides. Venn diagram (right) indicates the number of unique proteins identified at T0 (control) and T30 (hormone) **B**. Immunoprecipitation was performed in T47D cells treated with hormone for 30 min using anti-PARP1, PARG, PAR or NUDIX5 antibodies. Protein complexes were separated by SDS-PAGE and probed using specific antibodies. **C**. T47D cells were transfected with specific siRNA against NUDIX5 and the protein levels measured by western blot using specific antibodies. **D**. Quantification of nuclear, mitochondrial and cytoplasmic targeted luciferase constructs treated with hormone for the indicated times in untreated cells (control) or in cells depleted of NUDIX5 (NUDIXi). Data represents the mean of three independent experiments carried out in duplicate +/−SEM **E**. *In vitro* ATP assay performed in the presence or absence of recombinant NUDIX5 (100ng, 10ng, 0ng) with varying concentrations of ADPR (uM). The histogram shows relative light units (RLU) and represents the mean of two independent experiments carried out in triplicate +/−SEM. **F**. *In vitro* ATP assay as described above was spiked with known concentrations of ATP (uM standards). Data represents the mean of two independent experiments carried out in triplicate +/−SEM.

This evidence led us to hypothesize a crosstalk between PARP1, PARG and NUDIX5. Indeed NUDIX5 interacts with both PAR and PARG in a hormone dependent manner (Figure 3B). NUDIX5 has been reported to use ADP-ribose as substrate and to catalyze its hydrolysis to 5-ribose-phosphate and AMP ^19^. Therefore we explored whether NUDIX5 was involved in nuclear ATP generation. When we knockdown NUDIX5 with a specific siRNA (Figure 3C), we observed no compensatory changes in gene expression of Nudix family members NUDT6, NUDT9 or NUDT12 (Extended Figure 3E), but we found a marked reduction of the nuclear ATP peak at 30–50 min following hormone with no significant effect on mitochondrial ATP levels (Figure 3D). It is important to note that the peak of nuclear ATP around 10 min following hormone (Figure 3D dash box) is independent of PAR formation or degradation, (Figure 2E) as well as of NUDIX5 (Figure 3D). We conclude that the early increase in ATP is mitochondria-dependent, while the later nuclear ATP peak at 30–50 min is mitochondria-independent but PARP, PARG and NUDIX dependent.

To directly test the enzymatic activity of NUDIX5, we synthesized recombinant human NUDIX5 *in vitro* (Extended Figure 3F) and were able to detect the production of ATP in a luciferase assay when incubated at 37^o^C in the presence of ADP-ribose, PPi and 4 mM Mg2^+^ (Figure 3E, luciferase sensitivity using ATP standards Figure 3F).

## PAR derived ATP generation is needed for cell proliferation and chromatin remodeling

To explore the molecular processes for which PAR-derived ATP is used, we analyzed the effect of inhibiting nuclear ATP synthesis on chromatin dynamics and gene expression following hormone treatment. In expression microarray experiments, inhibition of PARG or depletion of NUDIX5 compromised regulation of 50% of hormone-responsive genes, with a significant overlap between genes dependent on PARG and on NUDIX5 (Figure 4A). 70% of both NUDIX5 and PARG dependent genes required both activities and majority of these genes also depended on PARylation (Figure 4B and C). Consistent with previous reports ^13, 20^, we found that genes dependent on PARP and PARG were enriched in processes such as signal transduction, cell proliferation, and stress (GO biological process analysis; Extended Table 5 and 6, Pathway enrichment analysis Extended Table 7). Progestin induced cell proliferation is abrogated by inhibition of PAR formation ^13^, and we now found that both PARG and NUDIX5 are also required for progestin induced cell proliferation (Figure 4D). This effect was also observed with another breast cancer cell line MCF7 in response to oestrogen and the associated changes in gene expression were also dependent on PAR synthesis, degradation ADPR and NUDIX5 (Figure 4E and F, respectively; see also Extended Figure 4A). In MCF7 cells estrogens induced increases in PAR and nuclear ATP that were dependent on the activities of PARP, PARG and NUDIX5 (Extended Figure 4B, and C). Although the response to estrogens of MCF7 cells was less synchronized, we conclude that nuclear ATP synthesis is required for the proliferative response of breast cancer cells to both steroid hormones.

**Figure 4:**
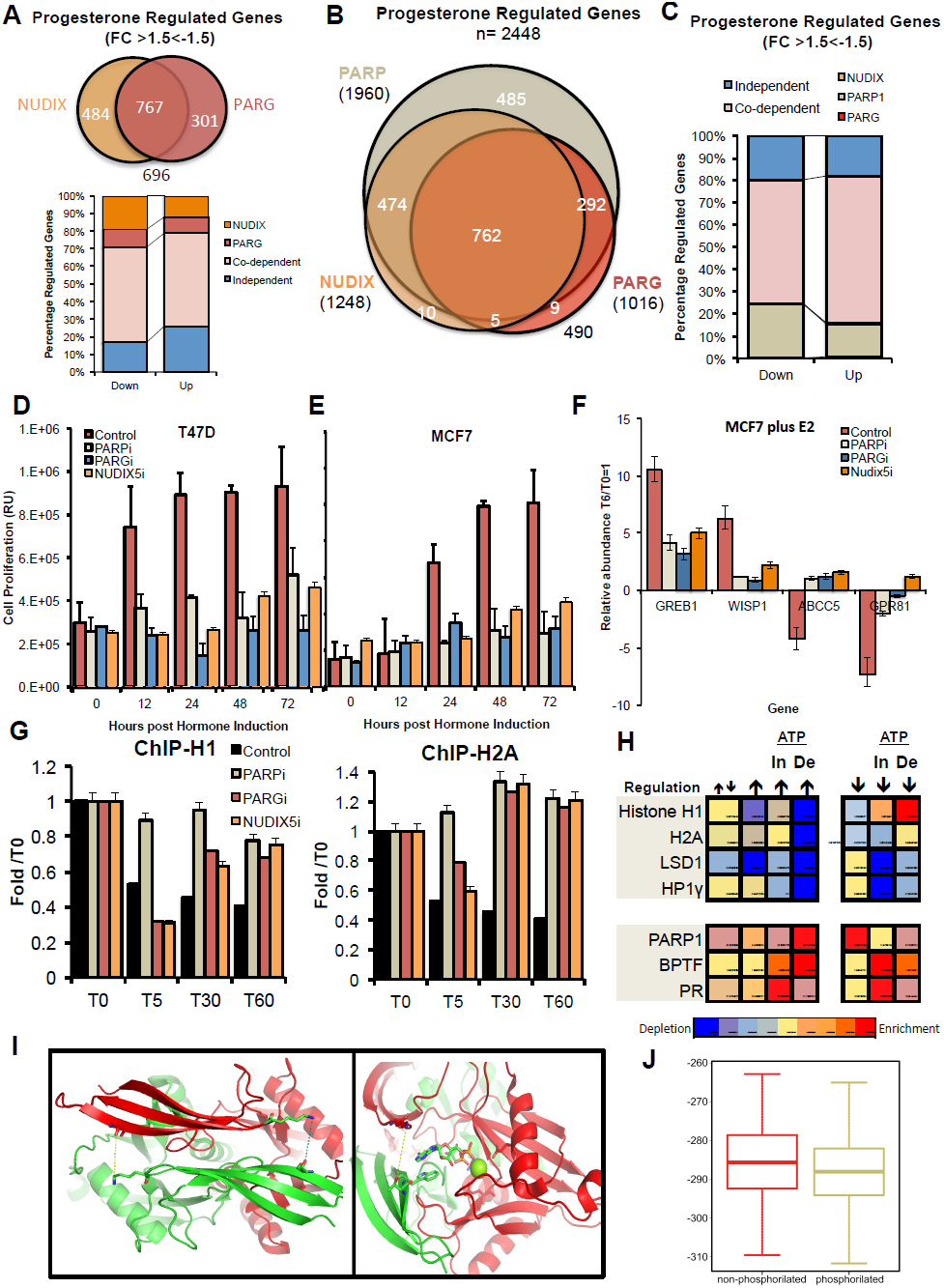
Nuclear ATP is essential for hormone regulated gene expression and chromatin remodeling. **A**. Venn diagram showing the overlap of Pg regulated genes (FC<−1.5>1.5, p<0.05) dependent on the actions of PARG and or NUDIX. Below; Breakdown of up and down progesterone regulated genes, indicating their dependence on PARG, NUDIX or both. **B**. Venn diagram showing the overlap of 2448 Pg regulated genes (FC<−1.5>1.5, p<0.05) dependent on the actions of PARP, PARG and or NUDIX. **C**. Breakdown of both up (FC>1.5, p<0.05) and down (FC<1.5, p<0.05) regulated genes and the dependence on PARP, PARG or NUDIX. **D**. Cell proliferation was measured by BrdU incorporation in T47D cells at the time points indicated following in the addition of hormone (R5020) in the presence inhibitors of PARP (3AB, PARPi) or PARG (TA, PARGi), or after knockdown of NUDIX5 (NUDIX5i). Data represents the mean of two independent experiments carried out in duplicate +/−SEM. **E**. Cell proliferation was measured by BrdU incorporation in MCF7 cells at the time points indicated following in the addition of hormone (17β-estradiol) in the presence inhibitors of PARP (3AB, PARPi) or PARG (TA, PARGi), or after knockdown of NUDIX5 (NUDIX5i). Data represents the mean of two independent experiments carried out in duplicate +/−SEM. **F**. The expression of oestrogen target genes in the presence of PARP, PARG or NUDIX5 inhibition was analysed by qRT-PCR. RNA was extracted 6 hours post hormone (17β-estradiol) addition. Data is represents the mean of three independent experiments carried out in duplicate +/−SEM. **G**. The enrichment of H1 (*left panel*) or H2A (*right panel*) was measured by ChIP using H2A or H1 specific antibodies in T47D cells treated with hormone for the indicated times in the presence or absence of PARP inhibitor (3AB, PARPi), PARG inhibitor (TA, PARGi) or siRNA targeted against NUDIX5 (NUDIX5i). Data represents the mean two independent experiments carried out in duplicate +/−SEM performed in 5 different amplified regions previously demonstrated to be depleted of both H2A and H1 following hormone treatment ^11, 13^. **H**. Heat-map showing the stronger chromatin remodeling (H1 and H2A displacement upper panel, recruitment PARP1, BPTF lower panel) over the promoter region (−2kbp) of genes up (↑) or down (↓) regulated by hormone and dependent (De) or independent (In) on nuclear ATP synthesis. **I**. Far view with the two interactions that will be fixed by the phosphorylation in (*left panel*) when the bond is broken the green monomer leaves and allows access to the ADPR (specifically to the two phosphates that are farther in this view). *Right*; View of the dimer from the active site. From the red chain we can see to GLU (93 -far- and 112 -right-) and the LYS27 that would stabilize the dimer when THR45 is phosphorylated and, thus, has more negative charge. The distance between both residues is 8 Angstroms here, a distance significantly decreased when T45 is phosphorylated (Extended Figure G and H). ADPR and the Mg ion are visualized. **J**. Comparison between the dimer stability/interface energies (using Zrank), between 500 decoys of NUDIX5 (PDB: 2DSC), non-phosphorylated (left) or phosphorylated (right). Both Wilcoxon (2.2E^−4^), and t-distributions (4.14E^−4^) support the statistical significance of this difference.

In response to progestin there are rapid and global changes in chromatin, genome topology ^11, 21^. Two consecutive chromatin remodeling events take place; an early one (1–5 min) leading to histone H1 displacement that depends on the activities of CDK2, PARP1 and NURF, and a later one (2–60 min) leading to displacement of H2A/H2B dimers catalyzed by BAF and PCAF ^11^. We confirm that the initial displacement of histone H1 depended on PARP1 activity, and found that it did not depend upon the actions of either PARG or NUDIX, while displacement at 30 and 60 min depended on all 3 activities (Figure 4G *left panel*). Similarly, initial displacement of H2A depended on PARP1, and only partially on PARG and NUDIX5, while subsequent H2A displacement required all 3 enzymatic activities (Figure 4G *right panel*). This suggests that initial nucleosome remodeling uses ATP coming form the mitochondria while the subsequent remodeling steps depend on nuclear ATP generation.

Collectively these data support the idea that ATP is required for genes which regulation involves extensive chromatin remodeling. To explore this notion we compared the quantitated chromatin remodeling (Displacement of in H1, H2A, HPIy, LSD1) −2kbp from the TSS of gene promoters, which regulation by hormone was dependent (De) or independent (In) of PARG and NUDIX5. We found that for both up-and down-regulated genes the extent of remodeling correlated well with the requirement of nuclear ATP, increased displacement or recruitment of H1 and H2A from up and down regulated genes respectively (Figure 4 H upper panel). The role of ADPR in remodeling also correlated as demonstrated by the increase in PARP1 and BPTF recruitment to up-regulated genes dependent on ATP (Figure 4H, lower panel). In contrast the levels of RNA synthesis changes in response to hormone did not correlate significantly with nuclear ATP requirement (data not shown). This is consistent with a higher requirement for ADPR derived ATP at genes which regulation required extensive chromatin remodeling.

## Discussion

Nuclear processes, including chromatin remodeling, transcription and replication require high amounts of energy in the form of ATP. Here we have described a novel mechanism of energy generation in the nuclei of breast cancer cells treated with hormone. In this model the high energetic cost of global chromatin modification is covered via the conversion of ADPR to ATP, to meet the local energy demands of the hormone response and to ensure cell survival.

Changes in nuclear ATP levels in response to hormone exhibit two distinct phases (Extended Figure 4D); an initial mitochondrial dependent phase (5–10 minutes) followed by a later increase in ATP (30–50 minutes), which is independent of the mitochondria but dependent on nuclear ADPR (Figures 1). In the initial stages histone H1 and H2A/H2B are locally displaced from the promoters of hormone responsive genes via the concerted actions of PARP1 and the chromatin remodeling factors BPTF and BAF ^11^. The energy requirements during these initial minutes are met via the pool of nuclear ATP present in the nucleus and are mitochondria dependent. It is possible that this ATP is also needed for the activity of the NMNAT-1 enzyme, which synthesizes NAD+ in the nucleus ^22^. NMNAT-1 is known to interact with PARP1 and to co-localized on hormone responsive genes, and could provide the NAD substrate for activation and PAR synthesis by PARP1 ^23^. The activity of PARP1 during these initial stages is essential for the displacement of histone H1 ^13^ and H2A/H2B, presumably via the traditional actions of PAR including direct parylation of histones ^24^. PARP1 has been shown to be capable of replacing histone H1 in chromatin and could directly participate in H1 displacement ^25^. During these initial stages of hormone activation the levels of PAR steadily increase in concert with a decrease in NAD levels.

During the second phase of hormone response and due to the direct action of PARG, PAR levels sharply decrease, accompanied by a rescue of NAD and an increase in nuclear ATP. This energy store is as a direct result of ADPR conversion to ATP via NUDIX5 and is essential for hormone dependent chromatin remodeling, gene regulation, cell proliferation and survival. In preliminary experiments we found that survival of MCF7 cells after DNA damage is dependent on PARP1, PARG and NUDIX5 (Data not shown).

Indeed we have preliminary evidence for a possible mechanism of NUDIX5 activation. Of all Nudix hydrolase family members only NUDIX5 is known to form a homodimer that hydrolyses ADPR to 5-ribose-phosphate and AMP ^26^. Phosphoproteomic analysis in T47D^M^ identified NUDIX5 as phosphorylated threonine 45 (T45) prior to hormone and dephosphorylated following hormone treatment (Extended Figure 4E). According to structural modelling T45 phosphorylation is essential for the stability of the homodimer and dephosphorylation of T45 should result in a de-stabilisation of the homodimer (Figure 4I and J, Extended Figure 4F). While hydrolysis of ADP-ribose to AMP would be energetically favorable with the homodimer, the reaction energetically and structurally feasible by the monomer in the presence of pyrophosphate would be the hydrolysis of ADP-ribose to 5-ribose phosphate and ATP. To test this hypothesis we generated NUDIX5 T45 phospho-null and phospho-mimetic mutants (T45A and T45D) and found that the T45D mutant behaves as dominant negative for Pg induced cell proliferation and gene regulation (Figure 4G and H respectively). Moreover, recombinant wt NUDIX5 expressed in bacteria and likely unphosphorylated catalyzes the synthesis of ATP from ADP-ribose and PPi (Figure 3E).

Previous work suggested that ATP generated via ADPR might play a role in DNA damage and replication ^7^. It has been suggested that in response to stimuli, and prior to transcription the torsional stress endured by the chromatin is relieved by the generation of local DNA breaks ^27^. In addition an interaction of progesterone receptor with the DNA-dependent protein Kinase Ku has been reported ^28^, further supporting a link between hormone action and DNA repair. It is therefore possible that the important role of parylation in DNA repair is also related to need of nuclear energy for the extensive chromatin remodeling associated with the repair process.

Finally NUDIX5 overexpression in breast cancer patients correlates with a poor outcome in hormone receptor positive breast cancer compared to PR and ER negative cancer (Extended Figure 4I). This in addition to the observation that NUDIX5 overexpression is significant in multiple cancer types and correlates with high expression of PARP1 (Extended Figures 3D and 4J) suggests that perhaps this mechanism of energy rescue and ATP generation is a process exploited in cancer and hence NUDIX5 provides an exciting new target for PARP1 combinatorial drug therapy.

## Acknowledgements

We wish to thank Harm H. Kampinga, Department of Cell Biology, RuG & UMC Groningen, The Netherlands for the nuclear, cytoplasmic and mitochondrial targeted luciferase expressing plasmids, and Hiromi Imamura, The Hakubi Center for Advanced Research & Graduate School of Biostudies, Kyoto University, Japan for the Ateam ATP/ADP expressing plasmids. We also thank Sara Capdevila, and Juan Martin Caballero for their support with IVIS Bioluminescent imaging equipment, the CRG Microarray, Proteomics and Microscopy Facilities for expert help and Juan Valcarcel, CRG, for advice with the manuscript. We acknowledge financial support from the Spanish MEC (CSD2006-00049; BMC 2003-02902 and 2010-15313) and the Catalan Government (AGAUR).

## Materials and Methods

### ATP in vitro Assay

ATP production via ADP-ribose degradation was inferred via measurement of luciferase in vitro briefly as follows. Recombinant human NUDIX5 at varying concentrations (Extended methods), was incubated in 200uM PPi, 10mM MgCl2, 10mM tris-HCl, in the presence or absence of ADP-ribose (Sigma) at 37C for 20 minutes. Equal volumes of luciferase substrate (Roche HSII ATP kit) was then added and the luciferase measurements made using 1s integration time.

## Author Contributions

RHGW and MB designed the study and the strategy; RHGW performed the large majority of the experimental work; RHGW and FLD performed the light microscopy measurements; DS and AP helped with the bioinformatic analysis, JB and BO did the structural modeling, ASN and GPV helped with ChIP-, DNaseI- and MNase-seq analysis, MW performed the phosphoproteomic analysis, R.H.G.W. and MB discussed the results and wrote the paper.

**Extended Data Figure 1.**
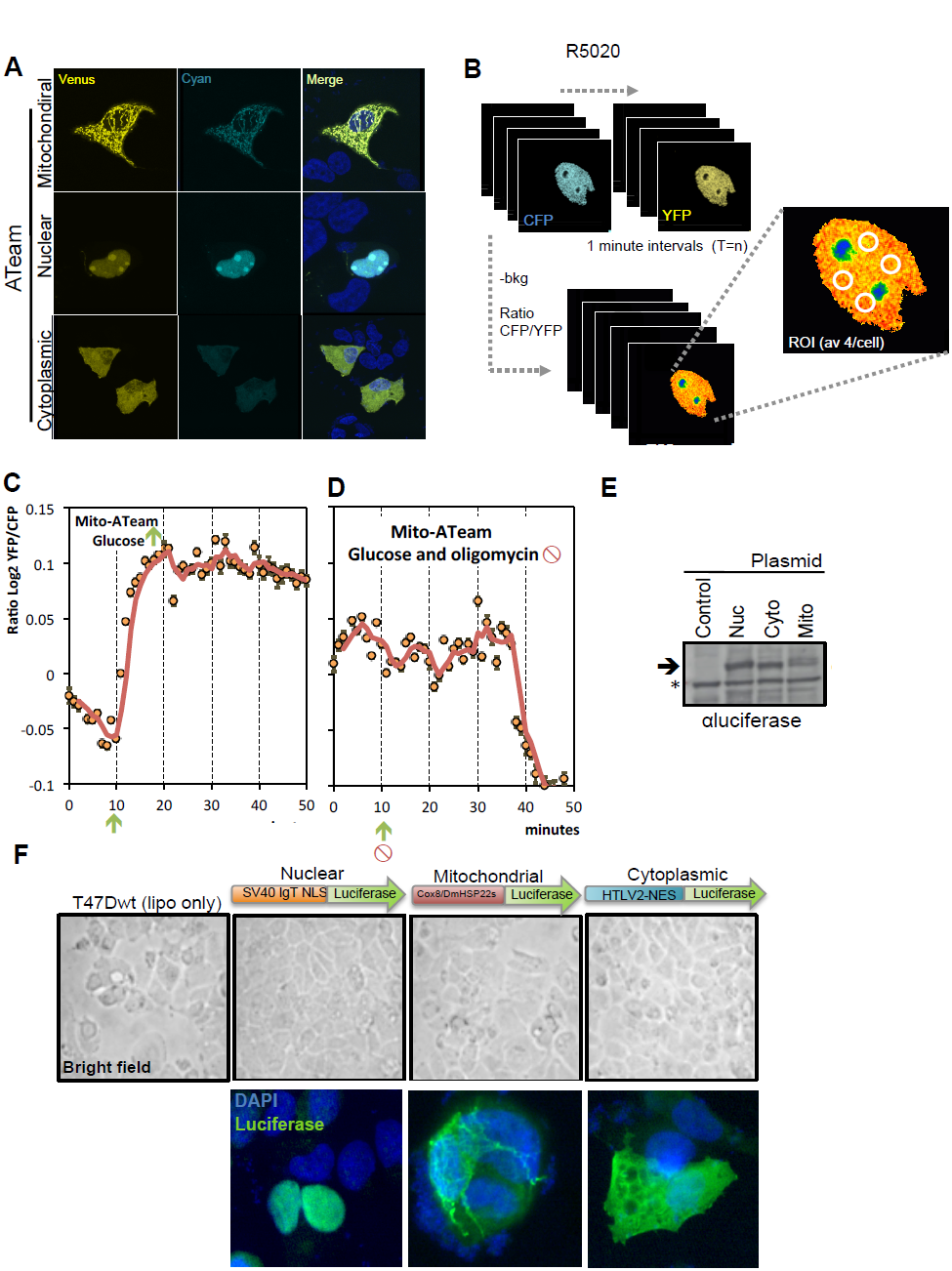
ATP levels increase in the nucleus of cells treated with hormone.

**A**. Representative immunofluorescence images showing the localization of Mitochondrial, Nuclear and Cytoplasmic ATeam ATP sensors in T47D cells.

**B**. Schematic of the experimental procedure used to record and measure ATP levels live in cells.

**C**. ATP/ADP ratio quantification of Mito-ATeam treated with glucose (10mM indicated by arrow). Data is presented as log_2_^YFP/CFP^.

**D**. ATP/ADP ratio quantification of Mito-ATeam treated with simultaneous treatment of glucose (10mM, indicated by arrow) and oligomycin (1ug/ml, indicated by red symbol). Data is presented as log_2_^YFP/CFP^.

**E**. T47Dwt cells transfected with nuclear/mitochondrial or cytoplasmic targeted luciferase constructs. The protein levels of each construct were detected by western blotting using luciferase specific antibody (* indicates non-specific antibody binding).

**F**. T47Dwt cells transfected with nuclear/mitochondrial or cytoplasmic targeted luciferase constructs (schematic indicated). Representative images of the cellular localisation of each construct comparing DAPI, nuclear staining and an anti-luciferase antibody is shown, no phenotypic abnormalities were observed following expression of the luciferase constructs (bright field).

**Extended Data Figure 2.**
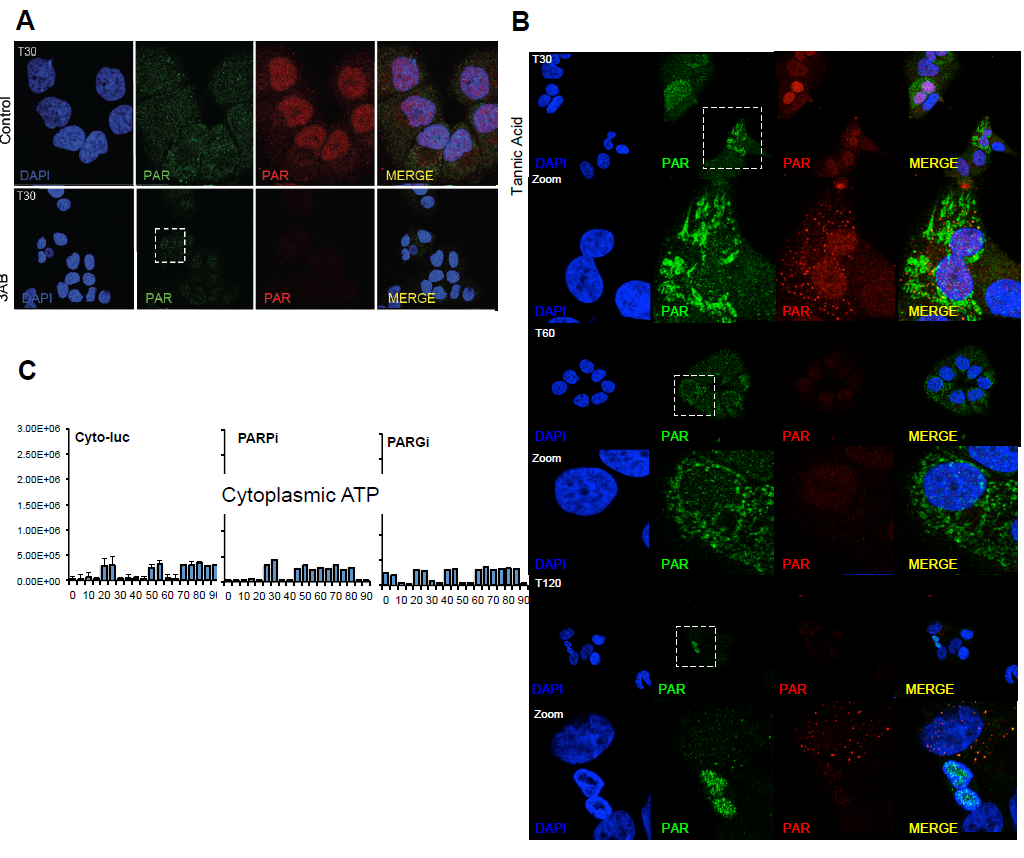
Specificity and sensitivity of ATeams. **A**. PAR levels were visualised by immunofluorescence using two different anti-PAR antibodies (monoclonal;*red*, polyclonal; *green*) in cells treated with hormone for 30 min in control (no inhibitor) or following inhibition of PARP (3AB, PARPi). **B**. PAR levels were visualised by immunofluorescence using two different anti-PAR antibodies (monoclonal;*red*, polyclonal; *green*) in cells treated with hormone for the indicated times following inhibition of PARG (TA, PARGi). **C**. Bioluminescent quantification of cytoplasmic ATP levels using the cytoplasmic targeted luciferase construct in T47Dwt cells treated with hormone following either solvent alone (control) or prior treatment with inhibitors of PARP (3AB, PARPi) or PARG (TA, PARGi).

**Extended Data Figure 3.**
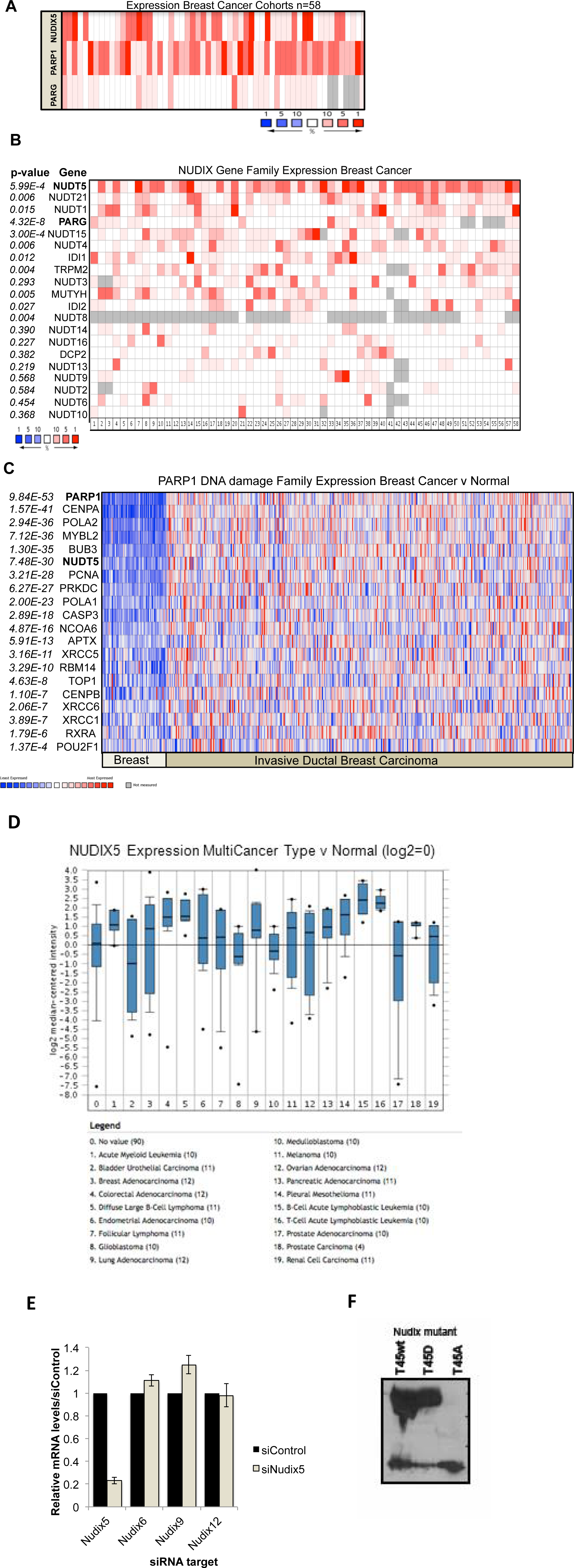
NUDIX5 overexpression correlates with PARP1 and PARP family expression in breast and other cancers. **A**. Expression of NUDIX5, PARP1 and PARG in 58 breast cancer cohorts (n=2456 patients). **B**. Expression of NUDT5, PARG and other annotated Nudix family members in the 58 breast cancer cohorts (**A**) (n=2456 patients). The p value indicates the significance of the observed difference; note that only the overexpression of NUDT5 and PARG but not other Nudix family members is significant. **C**. The expression of NUDT5, in conjunction with annotated PARP1 family members is shown for normal breast and invasive ductal breast carcinoma (n=1056 patients). The p value indicates the significance of the difference; note that NUDT5 overexpression correlates with PARP1 and DNA damage markers in cancer versus normal tissue. **D**. Expression of NUDIX5, in 19 different cancer types (log2 median centred). **E**. qRT-PCR analysis showing the relative mRNA expression of Nudix family members NUDIX6, NUDIX6 and NUDIX9 in the presence or absence of NUDI5 siRNA. **F**. *In vitro* expression of pANT-NUDIX5 plasmid in a transcription/translation assay generates GST-tagged NUDIX5 (as described in extended methods) detected by western blot with anti-GST antibody.

**Extended Data Figure 4.**
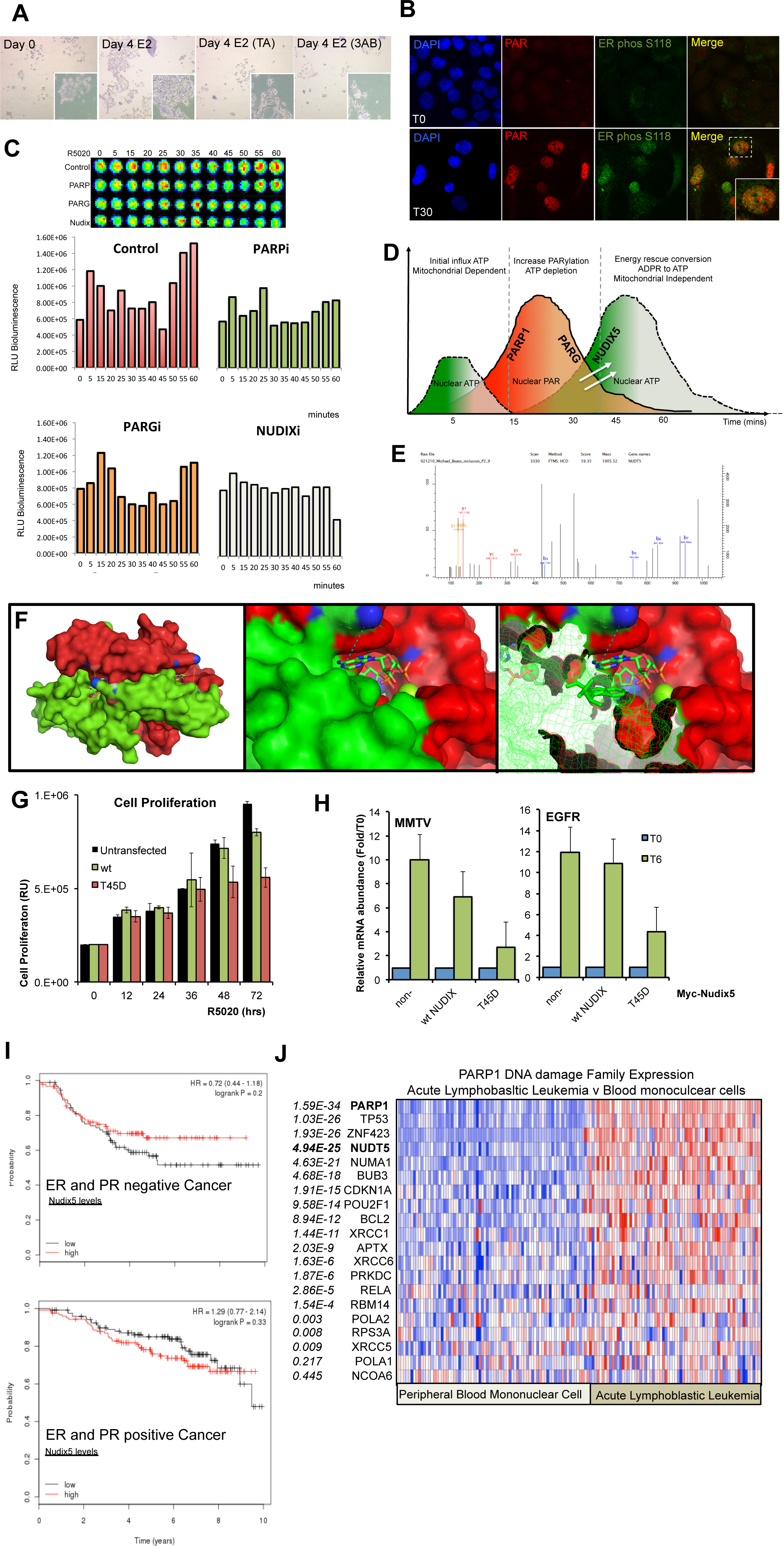
Cell growth and gene expression in response to oestrogen is dependent on the activities of PARP, PARG and NUDIX5. Mechanism of action **A**. Bright field images showing the proliferation of MCF7 cells 4 days post estrogen treatment in the presence or absence of PARP (3AB, PARPi) or PARG (TA, PARGi) inhibitors. **B**. Immunofluroescence images showing the levels of PAR (red) and phospho-ER (Serine 118, green) in MCF7 cells treated with oestrogen for 30 minutes. **C**. ATP levels were measured in MCF7 cells treated with estrogen for the time points indicated by bioluminescence (quantification **lower panel**, example scan **upper panel**), in the presence or absence of PARP (3AB, PARPi) or PARG (TA, PARGi) inhibitors or NUDIX5 targeted siRNA (NUDIX5i). **D**. Model of ATP economy in the nucleus in response to hormone. ATP dynamics can be separated into three distinct phases. First, energetic demands of the nucleus are met via mitochondrial dependent mechanisms. ATP is used for initial remodeling and for NAD synthesis. Second, transient increase in PARylation via the actions of PARP1 and PARG result in a depletion of cellular NAD, in concert with high energetic demand for chromatin remodeling and activation NUDIX5. Third, the combined action of PARG and NUDIX5 result in an increase in nuclear ATP independent of the mitochondria, in order to ensure the vast energetic demands and to preclude cell death via PARthanatos. **E**. MS2 spectrum for the phosphorylated Nudix5 peptide, showing the identification of Nudix5 T45 phosphorylation by high-resolution mass spectrometry. T47D cell extracts were prepared and proteins digested into peptides using trypsin. Phosphopeptides were enriched using strong cation exchange (SCX) and immobilized metal affinity (IMAC) chromatography. **F**. Far view with the two interactions that will be fixed by the phosphorylation in volume *(left*).View of the dimer from the active site visualizing both Nudix5 molecules in volume (middle) or one in mesh (right panel). ADPR and the Mg ion are visualized. **G**. Overexpression of the phosphomimetic NUDIX5T45D mutant in T47D cells acts as a dominant negative on cell proliferation in response to hormone. Data represents the mean relative mRNA abundance (FC T6hrs/T0hrs) of three independent experiments carried out in duplicate +/−SEM. **H**. Overexpression of NUDIX5T45D in T47D is dominant negative in progesterone induced gene expression analysis. Data represents the mean relative mRNA abundance (FC T6hrs/T0hrs) of three independent experiments carried out in duplicate +/−SEM. **I**. Kaplain-meyer survival graph showing the trend for both ER and PR positive (**top**) or negative (**bottom**) breast cancer samples. Data was separated according to the relative levels of Nudix (high (>75% percentile or low <25% percentile). Data was obtained from the TGVA cancer atlas. **J**. The expression of NUDIX5, in conjunction with annotated PARP1 family members is shown for normal peripheral blood mononuclear cells or acute lymphoblastic leukemia. The p value given indicates the significance of the difference obtained. Note that NUDIX5 overexpression clearly correlated with PARP1 and DNA damage proteins in cancer versus normal tissue.

