## Extended Materials and Methods and Tables for "ADP-ribose derived Nuclear ATP is Required for Chromatin Remodeling and Hormonal Gene Regulation (97 charact)"

**Extended Data**

**Extended Materials Methods**

**Cell culture; hormone and inhibitor treatment**

T47DM cells were used for all experiments unless otherwise stated. For hormone induction experiments, cells were grown in RPMI medium without Phenol Red, supplemented with 10% dextran-coated charcoal-treated FBS (DCC/FBS) after 24 h in serum-free conditions; cells were incubated with R5020 (10 nM) or vehicle (ethanol) as described (Vicent et al. 2011). For hormone induction experiments in MCF7 cells a similar procedure was performed; cells were grown in DMEM medium without Phenol Red, supplemented with 10% dextran-coated charcoal-treated FBS (DCC/FBS) after 24 h in serum-free conditions; cells were incubated with Estradiol (10 nM) or vehicle (ethanol). PARG and PARP inhibition were carried out via incubating cells with 5uM TA (tannic acid) or 10uM 3AB (3-amino-benzamide) respectively 1 hour prior to hormone treatment. All transfections were performed using Lipofectamine$^{2000}$ (Invitrogen) according to manufacturers instructions.

**PAR-capture ELISA**

Hormone and or inhibitor treatments were carried out as described, and sample preparation was carried out as follows: At the required time point, cells were washed twice with ice-cold PBS and scraped in lysis buffer (0.4 M NaCl, 1% Triton X-100) plus protease inhibitors. Cell suspensions were then incubated for 30 min on ice with periodic vortexing. The disrupted cell suspension was centrifuged at 10,000g for 10 min at 4°C, and the supernatant was recovered, snap-frozen, and stored at 80°C until required. Ninety-six-well black-walled plates were incubated with 2 ng/mL anti-PAR monoclonal antibody (Trevigen) in 50 mM sodium carbonate (pH 7.6) overnight at 4°C. The plates were washed once with PBS and blocked with blocking solution (5% semiskimmed milk powder, 0.1% Tween 20, 25 mM Tris at pH 7.4, 150 mM NaCl) for 1 h at room temperature. The plate was then washed four times in PBS and 0.1% Tween 20. One-hundred microliters of each sample (20 mg of total protein) or a PAR standard was applied and incubated for 1.5 h at room temperature. The plate was washed four times with PBS and 0.1% Tween 20 and incubated with anti-PAR rabbit polyclonal antibody (Trevigen) for 1.5 h at room temperature. The plate was washed four times with PBS and 0.1% Tween 20 and incubated with anti-rabbit HRP conjugate secondary antibody (5% semiskimmed dried milk) for 1.5 h at room temperature. The plate was washed six times with PBS and 0.1% Tween 20, and PAR was identified via incubation with TACs Sapphire (Trevigen) for 10 min in the dark. The reaction was terminated with 5% phosphoric acid, and the absorbance at 450 nm and 520 nm was recorded.

**NAD depletion assay**

$T47D^M$ cells were either left untreated or treated with the inhibitor indicated prior to R5020 addition as described. NAD concentration was determined as described (Wright et al 2012). At T = t, cells were washed with ice-cold PBS and incubated in 0.5 M PCA for 20 min on ice. An equal volume of KOH (1 M)/K2HPO4 (0.33 M) (pH 7.0) was added, and the samples were vortexed and incubated for 20 min on ice. Samples were spun at 10,000 rpm for 2 min at 4°C, and the supernatant was stored at _20°C. The quantification of NAD was carried out as follows: Samples were added to an equal volume of reaction buffer (200 mM Bicine at pH 7.8, 1 mg/mL BSA, 1MEtOH, 10mMEDTA at pH 8.0, 4mMphenazine). One tenth of the reaction volume of alcohol dehydrogenase was added and incubated for 30 min at 30°C. The reaction was terminated with a 0.33 sample volume of 12 mM iodoacetate, and the absorbance at 570 mM was measured.

**Modelling and evaluation of the phosphorylated and unphosphorylated T45 NUDIX5 homodimer.** In order to mimic the phosphorylation in T45 of NUDIX5 homodimer, we modeled the mutant form T45D with MODELLER (Fischer et al 2003). The dimer structure of NUDIX5 was taken from the crystal 2DSC (Zhong et al 2006) deposited in the Protein Data Bank (Berman et al 2000) and used as template for modeling. We used MODELLER to generate, by simulated annealing and distance restraints, two sets of 500 conformations of the native and T45D mutant form of NUDIX5 homodimer. These sets respectively mimicked the homodimer interaction of the phosphorylated and non-phosphorylated forms of NUDIX5.

The comparison of the homodimer interaction, phosphorylated and non-phosphorylated, was performed using the docking score potential of ZRANK(Pierce et al 2007). We calculated the distribution of the scores of docking of both sets.

**Chromatin Immunoprecipitation (ChIP) in cultured cells.** ChIP assays were performed as described (Vicent et al 2009) using the H2A specific antibody (Cell Signaling Technologies), or the H1 antibody (AE4 abcam). Quantification of chromatin immunoprecipitation was performed by qRT-PCR and the fold enrichment of target sequences in the immunoprecipitated (IP) compared to input (I) fractions were calculated using the comparative Ct method with ( $2^{Ct(IP)-Ct\ (Ref).}$). Values were corrected by the human b-globin gene and referred as relative abundance over time zero. Primers sequences are available on request

**RNA interference experiments.** All siRNAs were transfected into the T47D$^{M}$ cells using Lipofectamine 2000 (Invitrogen) as described (Vicent et al 2009). siRNA are sequences available on request.

**RNA extraction and RT-PCR.** Total RNA was prepared and cDNA generated as previously described (Wright et al. 2012). Quantification of gene products was performed by qRT-PCR. Each value calculated using the standard curve method was corrected by the human GAPDH and expressed as relative RNA abundance over time zero. Primer sequences are available on request.

**Protein Extract Preparation, western blotting/Co-immunoprecipitation assay.** Cells were prepared as previously described (Wright et al 2012), briefly cells were lysed (1% NP40, 150mM NaCl, 50mM Tris-HCl) and following total protein quantification, 2mg of cell extracts were incubated with protein G/A agarose beads previously coupled with 5μg of the corresponding antibodies or an unspecific control antibody as described (Vicent et al. 2009). Inputs and immunoprecipitated material was analyzed by western blot using; PARP1 (Cell Signalling technologies), PARG (Abcam ab169639), PAR (trevigen) and NUDIX5 (abcam) specific antibodies.

**Immunofluroescence.** T47D$^{M}$ or MCF7 cells were grown on coverslips treated with hormone for the desired length of time as described in previous sections prior to fixation using 4% paraformaldehyde in PBS for 15 min and permeabilized with PBS

0.2% Triton X-100 at room temperature. Coverslips were blocked with 5% skim-milk for 1 h at room temperature and incubated 2 hours with primary antibodies diluted in PBS 1% skim-milk at 1/500 PAR monoclonal and or polylonal (trevigen)), 1/1000 (PARP1 (Cell Signalling technologies), or 1/500 anti-phospho Serine 118 ER, anti-luciferase (Sigma L0159). After washes with PBS Tween 20 0.05%, samples were incubated with secondary antibodies (AlexaFluor 555 anti-rabbit and AlexaFluor 488 anti-mouse, Invitrogen-Molecular Probes) for 1 h at room temperature. After washes with PBS Tween 0.05% and DNA staining with DAPI, samples were mounted with mowiol. Images were acquire with a Leica TCS SP5 CFS confocal microscope.

**Microarray Design and Procedure.** The levels of NUDIX5 were reduced in $T47D^M$ cells using siRNA as described or the activity of PARP and PARG were inhibited by treating $T47D^M$ cells with either treated with 3AB or TA respectively. As a control, $T47D^M$ cells were treated with solvent (EtOH alone) prior to R5020 induction for 0 or 6 hours as described above. Total RNA was prepared and cDNA generated as described. Independently generated biological replicates hybridised against the human whole genome 44K expression platform (Agilent). Significantly induced, or repressed genes were considered as those with a $\log_2$ fold of $\geq 1.5$ and $\leq -1.5$ respectively and a q value of $\leq 5.00$.

**Site directed mutagenesis and detection of Myc-NUDIX5 phosphomimetic constructs for expression in mammalian cells.** Site-directed mutagenesis was carried out on the wild-type human PARP1 fused to Myc (pCMV-Myc NUDT5; genescript) as directed in the Quick Change mutagenesis kit. The mutagenic primers were as follows 5'-3'; T45D Fw; TGTACGTTTCACTGATTCCCAATCTCTAGTTTTACCAGTAGGATCC, T47D Rv; GGATCCTACTGGTAAAACTAGAGATTGGGAATCAGTGAAACGTACA. Following the mutagenesis procedure, mutations were confirmed by sequencing. Myc-wtNUDIX5 or Myc-NUDIX5-Thr/Asp mutant plasmids were transfected into $T47D^M$ cells using Lipofectamine 2000 (Invitrogen) as described. 48 hours later, cells were treated with hormone, cell proliferation, and gene or protein expression determined as described above.

**Cell proliferation assay**

$T47D^M$ / MCF7 cells transfected with control, NUDIX5 siRNAs or treated with the inhibitors 3AB/tannic acid as described above. Cells ($1 \times 10^4$) were plated in a 96-well plate in the presence or absence of R5020 ($T47D^M$) or estradiaol (MCF7). The cell proliferation ELISA BrdU Colorimetric assay (Roche) was performed according to the manufacturer's instructions. The experiments were performed in triplicate.

**Purification of Recombinant NUDT5 in vitro**

Recombinant NUDIXT5 was produced using 1-step Human high yield in vitro mammalian translation (IVT) system (Thermo Scientific). Briefly, IVT reactions containing (HeLa cell lysate, accessory proteins, reaction mix, 4ug NUDIX5 DNA (pANT7 GST-NUDT5 (DNASU, plasmid depository)) in 100ul was dialysed for 16 hours at 30°C according to manufacturers instructions. Recombinant NUDIX5 was then purified from the total reaction using GST beads (as described Wright et al 2012).

**DNA damage; Cell Survival Assay**

MCF7 cells were transfected with control, NUDIX5 siRNAs or treated with the inhibitors 3AB/tannic acid as described above. Cells were then plated in a 100mm plates at varying concentrations and subjected to DNA damage (H2O2) at varying concentrations (0.1, 0.2, 0.5mM). The medium was replaced every 3 days and the resultant surviving colonies stained using crystal violet solution (0.5% crystal violet, 10% methanol). Data is represented as a percentage surviving fraction compared to control (undamaged).

**PAR Mass Spec Peptide Enrichment analysis**

T47D cells were treated with hormone or ethanol for 30 minutes and the resultant lysates were immunoprecipted using anti-PAR antibody. Immunoprecipiated peptides were digested with trypsin and 1ug of each sample were analyzed by LCMSMS using a MEDI_CID method in the LTQ Orbitrap XL. To avoid carry over, samples were injected in increasing amount and BSA controls were included both in the digestion and LCMSMS analysis for quality control. Samples were searched against SwissProt_Human (January 2013) using an internal version of the search algorithm Mascot (http://www.matrixscience.com/). Data analysis was performed with Proteome Discoverer v1.3. Peptides have been filtered based on the FDR (False Discovery rate) for peptides with FDR better (lower) than 5%. Protein Discoverer gives an approximate estimation of protein amount with the parameter "Area" which is the average peak area of the 3 top peptides for a given protein. Immunoprecipitation was performed in two independent experiments for both control and hormone treated samples and the relative fold change compared to control was determined using the average area for the two experiments.

**Real time ATP visualisation using ATeam probes following hormone**

T47D cells were transfected with either Nuclear (Nuc) / Mitochondrial (Mito) or Cytoplasmic (Cyto) targeted A-team constructs in 35mm diameter cell culture plates using lipofectamine 2000 (Roche, following manufacturers instructions). 24 hours prior to hormone treatment cells were serum starved (RPMI 0% FCS, minus phenol red). For each individual experiment, 4 fields of view were selected containing between 2-5 cells/view. R5020 was added directly at the microscope prior to YFP and CFP emission detection at 1 minute intervals for the length of time required at selected FOV. For analysis; multiple regions of interest (ROI) of equal size were selected from each individual cell imaged and the ratio for each region of interest (ROI) was calculated following background subtraction according to protocol detailed in Kardash *et al.*, 2011. Results are represented as the mean +/-SEM of multiple ROIs.

**ATP visualisation using Bioluminescence Imaging**

T47D or MCF7 cells were seeded in RPMI or DMEM media respectively, as described above in black walled 96 well plates, were transfected using lipofectamine 2000 (Invitrogen, as described above) with the luciferase constructs (Nuc/Cyto/Mito-luciferase) as required, at a concentration of 50ng DNA/well. 24 hours prior to imaging the cells were starved with 0% FCS. R5020 or 17-estradiaol were added as described above for the length of time prior to the addition of 100ul of D-luciferin (1mg/ml, Sigma). The relative amounts of ATP were then visualised and quantitated using bioluminescence imaging IVIS system (Xenogen Corp).

**Solexa ChIP-seq analysis**

DNA was subjected to deep sequencing using the Solexa genome analyzer (Illumina Hi-Seq 2000). Single-ended sequences were trimmed to 50 bp and mapped to the human genome assembly hg19 using Bowtie (http//bowtie-bio.sourceforge.net<http://http//bowtie-bio.sourceforge.net>), an ultrafast short-read mapping program (Langmead et al. 2009), keeping only tags that mapped uniquely and with no more than two mismatches.

**Peak detection and annotation analysis**

The integrated software Macs (Zhang et al. 2008) was used to detect peaks that were enriched from background reads. The algorithm was applied by using the sliding window method to count the reads using the "nomodel" option, shift size set to 75, tag size set to 50, and a P-value threshold of 1 × 10−3 settings for transcription factors and chromatin remodelers; BPTF, HP1y, PARP1, CDK2, CTCF, PR, and PolII to identify differential binding between the two conditions by treating the T0 of the samples as the control. In the case of histone modifications, the algorithm was applied using "nomodel," "nolambda," and a P-value threshold of 1 × 10−3. In order to perform the average profiles, we used R programming language for plotting the mapped read signals over the regions of interest. All of the genome-wide annotations and statistics of protein–DNA interaction patterns from ChIP-seq data were performed with the standalone application CEAS (Cis-regulatory Element Annotation System). In the case of the ChiP-seq PR data, the signals of treated and untreated samples were compared using Poisson analysis (P-value < 0.001) (Ballare et al. 2013).

**GO Biological Process and Pathway Analysis**

GO biological process and pathway analysis was performed using MSigDB Collections database v4 gene set enrichment analysis online tools using a significance cut off value of $p < 0.05$.

### Extended References

Berman, H.M. *et al.* The Protein Data Bank. *Nucleic Acids Res* **28**, 235-242 (2000).

Fiser, A., Do, R.K. & Sali, A. Modeling of loops in protein structures. *Protein Sci* **9**, 1753-1773 (2000).

Pierce, B. and Weng, Z. (2007) ZRANK: reranking protein docking predictions with an optimized energy function. *Proteins*, **67**, 1078–1086.

Tao Ye, Arnaud R. Krebs, Mohamed-Amin Choukrallah, Celine Keime, Frederic Plewniak, Irwin Davidson, and Laszlo Tora. 2011. seqMINER: an integrated ChIP-seq data interpretation platform. Nucleic Acids Res. 2011 39: e35

Vicent, G.P. *et al.* 2009 Two chromatin remodeling activities cooperate during activation of hormone responsive promoters. *PLoS Genet* **5**, e1000567 (2009).

Vicent, G.P. *et al.* 2011 Four enzymes cooperate to displace histone H1 during the first minute of hormonal gene activation. *Genes Dev* **25**, 845-862.

Wright, R. H. G. *et al* 2012. CDK2-dependent activation of PARP-1 is required for hormonal gene regulation in breast cancer cells. *Genes Dev* **26**:1972-1983

Zha, M., Zhong, C., Peng, Y., Hu, H. and Ding, J. (2006) Crystal structures of human NUDT5 reveal insights into the structural basis of the substrate specificity. *J. Mol. Biol.*, **364**, 1021–1033.

Zhang, Y. *et al.* Model-based analysis of ChIP-Seq (MACS). *Genome Biol* **9**, R137 (2008).

**Extended Tables**

**Extended Table 1: Peptide Enrichment Before and after Hormone following ADPR enrichment.** Table lists accession number; description, and the peptide enrichment score for 2 independent experiments (A, B; hormone treated T30 and C, D Control T0) as described in methods above), results are represented as a log2(ratio) Hormone (T30)/Control (T0).

**Extended Table 2: PAR Mass Spec Functional Groups.** Table lists accession number; description, and the peptide ratio (hormone/control) for functionally related groups of proteins; histones, kinases, phosphatase and enzymes involved in NAD, ADPR metabolism and ATP synthesis.

**Extended Table 3: GO Biological Function Analysis Control (T0) PAR Mass Spec.** Significant ($p < 0.05$) GO biological processes enriched using proteins identified in control (T0) PAR mass spec analysis.

**Extended Table 4: GO Biological Function Analysis Control (T30) PAR Mass Spec.** Significant ($p < 0.05$) GO biological processes enriched using proteins identified in control (T30) PAR mass spec analysis.

**Extended Table 5: GO Biological Function Analysis for Genes Dependent on Nudix5.** Significant ($p < 0.05$) GO biological processes enriched using genes identified as Nudix5 dependent (Figure 4A).

**Extended Table 6: GO Biological Function Analysis for Genes Independent on Nudix5.** Significant ($p < 0.05$) GO biological processes enriched using genes identified as Nudix5 independent (Figure 4A).

**Extended Table 7: Pathway Analysis (KEGG/REACTOME for Genes Dependent or Independent on Nudix5.** Significant ($p < 0.05$) pathways enriched using genes identified as Nudix5 dependent or independent (Figure 4A).

Extended Table 1

| | | Area | | | | log (area) | | | | | | |
|---|---|---|---|---|---|---|---|---|---|---|---|---|
| | | Hormone_1 | Hormone_2 | Control 1 | Control 2 | Hormone_1 | Hormone_2 | Control 1 | Control 2 | Hormone | Control | Hormone/Control |
| Accession | Description | A3: Area | B3: Area | C3: Area | D3: Area | A3: Area | B3: Area | C3: Area | D3: Area | Average | Averag | Ratio |
| P56385 | ATP synthase subunit e, mitochondrial OS=Homo sapiens GN=ATP5I PE=1 SV=2 - [ATP5I_HUMAN] | 1.979E7 | 2.269E7 | | | 7.296 | 7.356 | | | 7.326 | 3 | 4.326 |
| C9IYN3 | Uncharacterized protein OS=Homo sapiens GN=HNRPLL PE=4 SV=1 - [C9IYN3_HUMAN] | | 2.103E7 | | | | 7.323 | | | 7.323 | 3 | 4.323 |
| Q15738 | Sterol-4-alpha-carboxylate 3-dehydrogenase, decarboxylating OS=Homo sapiens GN=NSDHL PE=1 SV=2 - [N | 2.166E7 | 1.241E7 | | | 7.336 | 7.094 | | | 7.215 | 3 | 4.215 |
| P04792 | Heat shock protein beta-1 OS=Homo sapiens GN=HSPB1 PE=1 SV=2 - [HSPB1_HUMAN] | 1.379E7 | | | | 7.140 | | | | 7.140 | 3 | 4.140 |
| O14561 | Acyl carrier protein, mitochondrial OS=Homo sapiens GN=NDUFAB1 PE=1 SV=3 - [ACPM_HUMAN] | 1.208E7 | 1.412E7 | | | 7.082 | 7.150 | | | 7.116 | 3 | 4.116 |
| E9PEJ4 | Uncharacterized protein OS=Homo sapiens GN=DLAT PE=3 SV=1 - [E9PEJ4_HUMAN] | 1.389E7 | 1.043E7 | | | 7.143 | 7.018 | | | 7.080 | 3 | 4.080 |
| D6RA32 | Uncharacterized protein OS=Homo sapiens GN=MAP1B PE=4 SV=2 - [D6RA32_HUMAN] | | 1.190E7 | | | | 7.076 | | | 7.076 | 3 | 4.076 |
| Q9BXP5-5 | Isoform 5 of Serrate RNA effector molecule homolog OS=Homo sapiens GN=SRRT - [SRRT_HUMAN] | | 1.109E7 | | | | 7.045 | | | 7.045 | 3 | 4.045 |
| F5GYG9 | Uncharacterized protein OS=Homo sapiens GN=AP2B1 PE=4 SV=1 - [F5GYG9_HUMAN] | 1.307E7 | 9.372E6 | | | 7.116 | 6.972 | | | 7.044 | 3 | 4.044 |
| Q9UNS2 | COP9 signalosome complex subunit 3 OS=Homo sapiens GN=COPS3 PE=1 SV=3 - [CSN3_HUMAN] | 1.209E7 | 8.448E6 | | | 7.082 | 6.927 | | | 7.005 | 3 | 4.005 |
| A3KFL1 | Exosome component 2 (Fragment) OS=Homo sapiens GN=EXOSC2 PE=4 SV=1 - [A3KFL1_HUMAN] | 9.318E6 | | | | 6.969 | | | | 6.969 | 3 | 3.969 |
| E9PDK5 | | 1.230E7 | 6.638E6 | | | 7.090 | 6.822 | | | 6.956 | 3 | 3.956 |
| E5RJ85 | Uncharacterized protein OS=Homo sapiens GN=CPNE3 PE=4 SV=1 - [E5RJ85_HUMAN] | 1.001E7 | 7.756E6 | | | 7.001 | 6.890 | | | 6.945 | 3 | 3.945 |
| Q9NUP9 | Protein lin-7 homolog C OS=Homo sapiens GN=LIN7C PE=1 SV=1 - [LIN7C_HUMAN] | | 8.699E6 | | | | 6.939 | | | 6.939 | 3 | 3.939 |
| P50897 | Palmitoyl-protein thioesterase 1 OS=Homo sapiens GN=PPT1 PE=1 SV=1 - [PPT1_HUMAN] | 1.154E7 | 6.333E6 | | | 7.062 | 6.802 | | | 6.932 | 3 | 3.932 |
| F8VR84 | Uncharacterized protein OS=Homo sapiens GN=C12orf10 PE=4 SV=1 - [F8VR84_HUMAN] | 7.966E6 | | | | 6.901 | | | | 6.901 | 3 | 3.901 |
| B4DZI8 | Coatomer protein complex, subunit beta 2 (Beta prime), isoform CRA_b OS=Homo sapiens GN=COPB2 PE=2 | 1.322E7 | 4.037E6 | | | 7.121 | 6.606 | | | 6.864 | 3 | 3.864 |
| Q5JS54 | Proteasome assembly chaperone 4 OS=Homo sapiens GN=PSMG4 PE=2 SV=2 - [PSMG4_HUMAN] | | 6.596E6 | | | | 6.819 | | | 6.819 | 3 | 3.819 |
| Q16851-2 | Isoform 2 of UTP--glucose-1-phosphate uridylyltransferase OS=Homo sapiens GN=UGP2 - [UGPA_HUMAN] | 9.629E6 | 3.282E6 | | | 6.984 | 6.516 | | | 6.750 | 3 | 3.750 |
| F5H170 | Uncharacterized protein OS=Homo sapiens GN=DNAJA4 PE=4 SV=1 - [F5H170_HUMAN] | 5.545E6 | | | | 6.744 | | | | 6.744 | 3 | 3.744 |
| Q92797-3 | Isoform 3 of Symplekin OS=Homo sapiens GN=SYMPK - [SYMPK_HUMAN] | | 5.455E6 | | | | 6.737 | | | 6.737 | 3 | 3.737 |
| O95182 | NADH dehydrogenase [ubiquinone] 1 alpha subcomplex subunit 7 OS=Homo sapiens GN=NDUFA7 PE=1 SV= | 7.846E6 | 2.817E6 | | | 6.895 | 6.450 | | | 6.672 | 3 | 3.672 |
| O94979-5 | Isoform 5 of Protein transport protein Sec31A OS=Homo sapiens GN=SEC31A - [SC31A_HUMAN] | 7.540E6 | 1.709E6 | | | 6.877 | 6.233 | | | 6.555 | 3 | 3.555 |
| B4DLG2 | Uncharacterized protein OS=Homo sapiens GN=ZC3H11A PE=2 SV=1 - [B4DLG2_HUMAN] | 3.315E6 | | 5.341E5 | 1.144E5 | 6.521 | | 5.728 | 5.059 | 6.521 | 5.393 | 1.127 |
| C9JZD1 | Uncharacterized protein OS=Homo sapiens GN=ARPC3 PE=4 SV=1 - [C9JZD1_HUMAN] | 5.007E7 | | 5.033E6 | | 7.700 | | 6.702 | | 7.700 | 6.702 | 0.998 |
| Q99497 | Protein DJ-1 OS=Homo sapiens GN=PARK7 PE=1 SV=2 - [PARK7_HUMAN] | 5.239E8 | 3.389E8 | 5.507E7 | 6.149E7 | 8.719 | 8.530 | 7.741 | 7.789 | 8.625 | 7.765 | 0.860 |
| Q9UNF0-2 | Isoform 2 of Protein kinase C and casein kinase substrate in neurons protein 2 OS=Homo sapiens GN=PACSI | 1.458E7 | 6.411E6 | | 1.415E6 | 7.164 | 6.807 | | 6.151 | 6.985 | 6.151 | 0.834 |
| Q9P0J0 | NADH dehydrogenase [ubiquinone] 1 alpha subcomplex subunit 13 OS=Homo sapiens GN=NDUFA13 PE=1 S | 1.385E7 | 1.215E7 | | 2.134E6 | 7.142 | 7.085 | | 6.329 | 7.113 | 6.329 | 0.784 |
| Q96C86 | Scavenger mRNA-decapping enzyme DcpS OS=Homo sapiens GN=DCPS PE=1 SV=2 - [DCPS_HUMAN] | 1.303E7 | | | 2.477E6 | 7.115 | | | 6.394 | 7.115 | 6.394 | 0.721 |
| C9JBI3 | Uncharacterized protein OS=Homo sapiens GN=PSPH PE=4 SV=1 - [C9JBI3_HUMAN] | 9.835E6 | | 6.403E5 | 6.269E6 | 6.993 | | 5.806 | 6.797 | 6.993 | 6.302 | 0.691 |
| Q8WWM7-6 | Isoform 6 of Ataxin-2-like protein OS=Homo sapiens GN=ATXN2L - [ATX2L_HUMAN] | 2.795E7 | 7.912E6 | 2.854E6 | 3.785E6 | 7.446 | 6.898 | 6.455 | 6.578 | 7.172 | 6.517 | 0.656 |
| P07355 | Annexin A2 OS=Homo sapiens GN=ANXA2 PE=1 SV=2 - [ANXA2_HUMAN] | 3.542E7 | 2.173E7 | 7.913E6 | 4.825E6 | 7.549 | 7.337 | 6.898 | 6.683 | 7.443 | 6.791 | 0.652 |
| P36871 | Phosphoglucomutase-1 OS=Homo sapiens GN=PGM1 PE=1 SV=3 - [PGM1_HUMAN] | 1.579E7 | 9.425E6 | 1.533E6 | 6.341E6 | 7.198 | 6.974 | 6.185 | 6.802 | 7.086 | 6.494 | 0.592 |
| E9PDV0 | Uncharacterized protein OS=Homo sapiens GN=CTH PE=3 SV=1 - [E9PDV0_HUMAN] | 1.822E7 | 7.752E6 | | 3.220E6 | 7.261 | 6.889 | | 6.508 | 7.075 | 6.508 | 0.567 |
| F5H8D7 | Uncharacterized protein OS=Homo sapiens GN=XRCC1 PE=4 SV=1 - [F5H8D7_HUMAN] | | 5.244E6 | 1.466E6 | | | 6.720 | 6.166 | | 6.720 | 6.166 | 0.553 |
| Q8N1F7 | Nuclear pore complex protein Nup93 OS=Homo sapiens GN=NUP93 PE=1 SV=2 - [NUP93_HUMAN] | 1.343E7 | | | 3.762E6 | 7.128 | | | 6.575 | 7.128 | 6.575 | 0.553 |
| C9JUE0 | Uncharacterized protein OS=Homo sapiens GN=TFG PE=4 SV=1 - [C9JUE0_HUMAN] | 1.068E7 | 5.124E7 | 7.172E6 | 6.169E6 | 7.028 | 7.710 | 6.856 | 6.790 | 7.369 | 6.823 | 0.546 |
| F5H3H2 | Uncharacterized protein OS=Homo sapiens GN=PFKL PE=4 SV=1 - [F5H3H2_HUMAN] | 2.007E7 | | | 5.732E6 | 7.303 | | | 6.758 | 7.303 | 6.758 | 0.544 |
| Q92922 | SWI/SNF complex subunit SMARCC1 OS=Homo sapiens GN=SMARCC1 PE=1 SV=3 - [SMRC1_HUMAN] | 2.696E7 | 7.960E6 | | 4.357E6 | 7.431 | 6.901 | | 6.639 | 7.166 | 6.639 | 0.527 |
| Q6R5J7 | 4.1G isoform OS=Homo sapiens GN=EPB41L2 PE=2 SV=1 - [Q6R5J7_HUMAN] | 1.861E7 | 8.718E6 | 3.983E6 | 3.697E6 | 7.270 | 6.940 | 6.600 | 6.568 | 7.105 | 6.584 | 0.521 |
| Q9BXW7-2 | Isoform 1 of Cat eye syndrome critical region protein 5 OS=Homo sapiens GN=CECR5 - [CECR5_HUMAN] | 1.339E7 | 9.270E6 | 3.356E6 | | 7.127 | 6.967 | 6.526 | | 7.047 | 6.526 | 0.521 |
| P20042 | Eukaryotic translation initiation factor 2 subunit 2 OS=Homo sapiens GN=EIF2S2 PE=1 SV=2 - [IF2B_HUMAN | 5.762E6 | 5.438E6 | | 1.734E6 | 6.761 | 6.735 | | 6.239 | 6.748 | 6.239 | 0.509 |
| Q9Y6G9 | Cytoplasmic dynein 1 light intermediate chain 1 OS=Homo sapiens GN=DYNC1LI1 PE=1 SV=3 - [DC1L1_HUMAN] | | 7.375E6 | 1.457E6 | 3.707E6 | | 6.868 | 6.163 | 6.569 | 6.868 | 6.366 | 0.502 |
| Q9UKY7 | Protein CDV3 homolog OS=Homo sapiens GN=CDV3 PE=1 SV=1 - [CDV3_HUMAN] | 6.656E7 | 2.291E7 | 7.688E6 | 1.970E7 | 7.823 | 7.360 | 6.886 | 7.295 | 7.592 | 7.090 | 0.501 |
| Q96CT7 | Coiled-coil domain-containing protein 124 OS=Homo sapiens GN=CCDC124 PE=1 SV=1 - [CC124_HUMAN] | | 1.907E7 | 3.324E6 | 1.124E7 | | 7.280 | 6.522 | 7.051 | 7.280 | 6.786 | 0.494 |
| P30419-2 | Isoform Short of Glycylpeptide N-tetradecanoyltransferase 1 OS=Homo sapiens GN=NMT1 - [NMT1_HUMAN] | 1.454E7 | 1.608E7 | 2.621E6 | 9.190E6 | 7.162 | 7.206 | 6.418 | 6.963 | 7.184 | 6.691 | 0.494 |
| A6NN80 | Annexin A6, isoform CRA_c OS=Homo sapiens GN=ANXA6 PE=3 SV=3 - [A6NN80_HUMAN] | 3.186E7 | 1.772E7 | 7.662E6 | | 7.503 | 7.248 | 6.884 | | 7.376 | 6.884 | 0.492 |
| B4DJ81 | Uncharacterized protein OS=Homo sapiens GN=NDUFS1 PE=2 SV=1 - [B4DJ81_HUMAN] | 1.736E7 | 7.347E6 | 4.286E6 | 3.274E6 | 7.240 | 6.866 | 6.632 | 6.515 | 7.053 | 6.574 | 0.479 |
| A8MUN4 | Uncharacterized protein OS=Homo sapiens GN=ATP6V1E1 PE=2 SV=1 - [A8MUN4_HUMAN] | 7.654E6 | 5.507E6 | 2.198E6 | | 6.884 | 6.741 | 6.342 | | 6.812 | 6.342 | 0.470 |
| A6NM98 | ARD1 homolog A, N-acetyltransferase (S. cerevisiae), isoform CRA_b OS=Homo sapiens GN=NAA10 PE=4 SV | 2.870E7 | 6.963E6 | 4.183E6 | 5.512E6 | 7.458 | 6.843 | 6.621 | 6.741 | 7.150 | 6.681 | 0.469 |
| B1AH89 | Tubulin tyrosine ligase-like family, member 12 OS=Homo sapiens GN=TTLL12 PE=4 SV=1 - [B1AH89_HUMAN | 1.329E7 | | 4.515E6 | | 7.123 | | 6.655 | | 7.123 | 6.655 | 0.469 |
| Q9ULC4-2 | Isoform 2 of Malignant T cell-amplified sequence 1 OS=Homo sapiens GN=MCTS1 - [MCTS1_HUMAN] | 2.245E7 | 2.525E7 | 3.211E6 | 2.194E7 | 7.351 | 7.402 | 6.507 | 7.341 | 7.377 | 6.924 | 0.453 |
| P35659 | Protein DEK OS=Homo sapiens GN=DEK PE=1 SV=1 - [DEK_HUMAN] | 2.286E7 | 1.501E7 | 4.896E6 | 8.747E6 | 7.359 | 7.176 | 6.690 | 6.942 | 7.268 | 6.816 | 0.452 |
| Q9BYN8 | 28S ribosomal protein S26, mitochondrial OS=Homo sapiens GN=MRPS26 PE=1 SV=1 - [RT26_HUMAN] | 7.242E6 | | 2.086E6 | 3.289E6 | 6.860 | | 6.319 | 6.517 | 6.860 | 6.418 | 0.442 |
| O43396 | Thioredoxin-like protein 1 OS=Homo sapiens GN=TXNL1 PE=1 SV=3 - [TXNL1_HUMAN] | 1.813E7 | | 8.030E6 | 5.371E6 | 7.258 | | 6.905 | 6.730 | 7.258 | 6.817 | 0.441 |
| Q13247-3 | Isoform SRP55-3 of Serine/arginine-rich splicing factor 6 OS=Homo sapiens GN=SRSF6 - [SRSF6_HUMAN] | 5.631E7 | 1.900E7 | 1.690E7 | 8.449E6 | 7.751 | 7.279 | 7.228 | 6.927 | 7.515 | 7.077 | 0.437 |
| P11413 | Glucose-6-phosphate 1-dehydrogenase OS=Homo sapiens GN=G6PD PE=1 SV=4 - [G6PD_HUMAN] | | 1.892E7 | 6.065E6 | 8.037E6 | | 7.277 | 6.783 | 6.905 | 7.277 | 6.844 | 0.433 |
| A8MZ71 | Uncharacterized protein OS=Homo sapiens GN=COX7A2 PE=4 SV=1 - [A8MZ71_HUMAN] | 3.596E7 | 2.951E7 | 9.632E6 | 1.577E7 | 7.556 | 7.470 | 6.984 | 7.198 | 7.513 | 7.091 | 0.422 |
| C9IZ80 | Uncharacterized protein OS=Homo sapiens GN=BZW1 PE=4 SV=1 - [C9IZ80_HUMAN] | 9.090E6 | | 6.079E6 | 2.044E6 | 6.959 | | 6.784 | 6.310 | 6.959 | 6.547 | 0.411 |
| P08621-2 | Isoform 2 of U1 small nuclear ribonucleoprotein 70 kDa OS=Homo sapiens GN=SNRNP70 - [RU17_HUMAN] | 2.621E7 | | | 1.046E7 | 7.419 | | | 7.019 | 7.419 | 7.019 | 0.399 |
| E9PJS4 | Uncharacterized protein OS=Homo sapiens GN=SRPR PE=4 SV=1 - [E9PJS4_HUMAN] | 2.183E6 | | | 8.729E5 | 6.339 | | | 5.941 | 6.339 | 5.941 | 0.398 |
| P45880-2 | Isoform 2 of Voltage-dependent anion-selective channel protein 2 OS=Homo sapiens GN=VDAC2 - [VDAC2_H | 1.195E8 | 8.456E7 | 4.016E7 | 4.120E7 | 8.077 | 7.927 | 7.604 | 7.615 | 8.002 | 7.609 | 0.393 |
| P21281 | V-type proton ATPase subunit B, brain isoform OS=Homo sapiens GN=ATP6V1B2 PE=1 SV=3 - [VATB2_HUM | 1.292E7 | 9.328E6 | 5.191E6 | 3.926E6 | 7.111 | 6.970 | 6.715 | 6.594 | 7.041 | 6.655 | 0.386 |
| O43488 | Aflatoxin B1 aldehyde reductase member 2 OS=Homo sapiens GN=AKR7A2 PE=1 SV=3 - [ARK72_HUMAN] | 1.942E7 | 7.736E6 | 3.233E6 | 7.995E6 | 7.288 | 6.889 | 6.510 | 6.903 | 7.088 | 6.706 | 0.382 |
| P19623 | Spermidine synthase OS=Homo sapiens GN=SRM PE=1 SV=1 - [SPEE_HUMAN] | 1.935E7 | | 1.207E7 | 5.358E6 | 7.287 | | 7.082 | 6.729 | 7.287 | 6.905 | 0.381 |
| Q15428 | Splicing factor 3A subunit 2 OS=Homo sapiens GN=SF3A2 PE=1 SV=2 - [SF3A2_HUMAN] | 3.326E7 | 1.521E7 | 9.121E6 | 9.691E6 | 7.522 | 7.182 | 6.960 | 6.986 | 7.352 | 6.973 | 0.379 |
| Q5LJ98 | TROVE domain family, member 2 OS=Homo sapiens GN=TROVE2 PE=2 SV=1 - [Q5LJ98_HUMAN] | 1.776E7 | 1.233E7 | 6.909E6 | 5.709E6 | 7.249 | 7.091 | 6.839 | 6.757 | 7.170 | 6.798 | 0.372 |
| Q99598 | Translin-associated protein X OS=Homo sapiens GN=TSNAX PE=1 SV=1 - [TSNAX_HUMAN] | | 1.894E7 | 4.519E6 | 1.434E7 | | 7.277 | 6.655 | 7.156 | 7.277 | 6.906 | 0.372 |
| Q15056-2 | Isoform Short of Eukaryotic translation initiation factor 4H OS=Homo sapiens GN=EIF4H - [IF4H_HUMAN] | 6.714E7 | 7.188E7 | 3.197E7 | 2.735E7 | 7.827 | 7.857 | 7.505 | 7.437 | 7.842 | 7.471 | 0.371 |
| P62805 | Histone H4 OS=Homo sapiens GN=HIST1H4A PE=1 SV=2 - [H4_HUMAN] | 3.810E7 | 2.604E7 | 1.365E7 | 1.322E7 | 7.581 | 7.416 | 7.135 | 7.121 | 7.498 | 7.128 | 0.370 |
| B0LM41 | Transcriptional coactivator CoAZ OS=Homo sapiens GN=RBM14/RBM4 fusion PE=2 SV=1 - [B0LM41_HUMAN | 1.444E7 | 5.028E6 | 4.980E6 | 2.730E6 | 7.159 | 6.701 | 6.697 | 6.436 | 6.930 | 6.567 | 0.364 |
| Q07021 | Complement component 1 Q subcomponent-binding protein, mitochondrial OS=Homo sapiens GN=C1QBP PE | 3.150E8 | 2.593E8 | 1.184E8 | 1.322E8 | 8.498 | 8.414 | 8.073 | 8.121 | 8.456 | 8.097 | 0.359 |
| F8VZ52 | Uncharacterized protein OS=Homo sapiens GN=DNM1L PE=3 SV=1 - [F8VZ52_HUMAN] | 2.183E7 | 1.640E7 | 8.134E6 | 8.670E6 | 7.339 | 7.215 | 6.910 | 6.938 | 7.277 | 6.924 | 0.353 |
| Q01081 | Splicing factor U2AF 35 kDa subunit OS=Homo sapiens GN=U2AF1 PE=1 SV=3 - [U2AF1_HUMAN] | 1.520E7 | 1.039E7 | 4.483E6 | 6.983E6 | 7.182 | 7.017 | 6.652 | 6.844 | 7.099 | 6.748 | 0.351 |
| E9PCI9 | Uncharacterized protein OS=Homo sapiens GN=FDPS PE=3 SV=1 - [E9PCI9_HUMAN] | 4.751E7 | 4.540E7 | | 2.072E7 | 7.677 | 7.657 | | 7.316 | 7.667 | 7.316 | 0.351 |
| Q9NR45 | Sialic acid synthase OS=Homo sapiens GN=NANS PE=1 SV=2 - [SIAS_HUMAN] | 3.057E7 | 1.022E7 | 8.225E6 | 7.586E6 | 7.485 | 7.009 | 6.915 | 6.880 | 7.247 | 6.898 | 0.350 |
| P36507 | Dual specificity mitogen-activated protein kinase kinase 2 OS=Homo sapiens GN=MAP2K2 PE=1 SV=1 - [MP2 | 1.744E7 | 5.530E6 | 4.432E6 | | 7.241 | 6.743 | 6.647 | | 6.992 | 6.647 | 0.346 |
| O95336 | 6-phosphogluconolactonase OS=Homo sapiens GN=PGLS PE=1 SV=2 - [6PGL_HUMAN] | 2.463E7 | 1.996E7 | | 1.014E7 | 7.391 | 7.300 | | 7.006 | 7.346 | 7.006 | 0.340 |
| P21796 | Voltage-dependent anion-selective channel protein 1 OS=Homo sapiens GN=VDAC1 PE=1 SV=2 - [VDAC1_HI | 1.316E8 | 5.926E7 | 5.254E7 | 3.121E7 | 8.119 | 7.773 | 7.720 | 7.494 | 7.946 | 7.607 | 0.339 |
| Q9H3K6 | BolA-like protein 2 OS=Homo sapiens GN=BOLA2 PE=1 SV=1 - [BOLA2_HUMAN] | 3.780E7 | 1.608E7 | 1.375E7 | 9.389E6 | 7.577 | 7.206 | 7.138 | 6.973 | 7.392 | 7.055 | 0.336 |
| Q9Y5J7 | Mitochondrial import inner membrane translocase subunit Tim9 OS=Homo sapiens GN=TIMM9 PE=1 SV=1 - | 3.481E7 | 1.564E7 | 9.939E6 | 1.170E7 | 7.542 | 7.194 | 6.997 | 7.068 | 7.368 | 7.033 | 0.335 |
| Q05CP4 | EIF2AK2 protein (Fragment) OS=Homo sapiens GN=EIF2AK2 PE=2 SV=1 - [Q05CP4_HUMAN] | | 4.878E6 | | 2.263E6 | | 6.688 | | 6.355 | 6.688 | 6.355 | 0.334 |
| O75874 | Isocitrate dehydrogenase [NADP] cytoplasmic OS=Homo sapiens GN=IDH1 PE=1 SV=2 - [IDHC_HUMAN] | 1.158E7 | 9.262E6 | 4.200E6 | 5.512E6 | 7.064 | 6.967 | 6.623 | 6.741 | 7.015 | 6.682 | 0.333 |
| Q96DI7 | U5 small nuclear ribonucleoprotein 40 kDa protein OS=Homo sapiens GN=SNRNP40 PE=1 SV=1 - [SNR40_HUMAN] | | 1.500E7 | 3.925E6 | 1.257E7 | | 7.176 | 6.594 | 7.099 | 7.176 | 6.847 | 0.330 |
| Q9Y520-2 | Isoform 2 of Protein PRRC2C OS=Homo sapiens GN=PRRC2C - [PRC2C_HUMAN] | 3.262E6 | 9.908E6 | 4.013E6 | 1.775E6 | 6.514 | 6.996 | 6.603 | 6.249 | 6.755 | 6.426 | 0.328 |
| P10253 | Lysosomal alpha-glucosidase OS=Homo sapiens GN=GAA PE=1 SV=4 - [LYAG_HUMAN] | 1.172E7 | 1.287E7 | 4.549E6 | 7.345E6 | 7.069 | 7.110 | 6.658 | 6.866 | 7.089 | 6.762 | 0.327 |
| E7ENK9 | Uncharacterized protein OS=Homo sapiens GN=CSE1L PE=4 SV=1 - [E7ENK9_HUMAN] | 1.026E8 | 6.546E7 | 3.103E7 | 4.811E7 | 8.011 | 7.816 | 7.492 | 7.682 | 7.914 | 7.587 | 0.327 |
| E7EWT5 | Uncharacterized protein OS=Homo sapiens GN=CIRBP PE=4 SV=1 - [E7EWT5_HUMAN] | 1.938E7 | | 9.166E6 | | 7.287 | | 6.962 | | 7.287 | 6.962 | 0.325 |
| Q9Y5K5-2 | Isoform 2 of Ubiquitin carboxyl-terminal hydrolase isozyme L5 OS=Homo sapiens GN=UCHL5 - [UCHL5_HUM | 1.528E7 | 7.622E6 | 5.176E6 | | 7.184 | 6.882 | 6.714 | | 7.033 | 6.714 | 0.319 |
| P27694 | Replication protein A 70 kDa DNA-binding subunit OS=Homo sapiens GN=RPA1 PE=1 SV=2 - [RFA1_HUMAN] | 2.551E7 | 2.421E7 | 1.090E7 | 1.320E7 | 7.407 | 7.384 | 7.037 | 7.121 | 7.395 | 7.079 | 0.316 |
| P42126-2 | Isoform 2 of Enoyl-CoA delta isomerase 1, mitochondrial OS=Homo sapiens GN=ECI1 - [ECI1_HUMAN] | 3.776E7 | 2.423E7 | 1.249E7 | 1.707E7 | 7.577 | 7.384 | 7.097 | 7.232 | 7.481 | 7.164 | 0.316 |
| P60900 | Proteasome subunit alpha type-6 OS=Homo sapiens GN=PSMA6 PE=1 SV=1 - [PSA6_HUMAN] | 6.933E7 | 4.406E7 | 2.531E7 | 2.815E7 | 7.841 | 7.644 | 7.403 | 7.449 | 7.742 | 7.426 | 0.316 |
| P53597 | Succinyl-CoA ligase [ADP/GDP-forming] subunit alpha, mitochondrial OS=Homo sapiens GN=SUCLG1 PE=1 S | 1.190E7 | 1.530E7 | 4.275E6 | 9.978E6 | 7.076 | 7.185 | 6.631 | 6.999 | 7.130 | 6.815 | 0.315 |
| P30043 | Flavin reductase (NADPH) OS=Homo sapiens GN=BLVRB PE=1 SV=3 - [BLVRB_HUMAN] | 1.055E7 | 1.498E7 | 4.910E6 | 7.608E6 | 7.023 | 7.175 | 6.691 | 6.881 | 7.099 | 6.786 | 0.313 |
| Q9UHD8-5 | Isoform 5 of Septin-9 OS=Homo sapiens GN=SEPT9 - [SEPT9_HUMAN] | 3.181E6 | 1.373E7 | 3.237E6 | 3.265E6 | 6.503 | 7.138 | 6.510 | 6.514 | 6.820 | 6.512 | 0.308 |
| P30040 | Endoplasmic reticulum resident protein 29 OS=Homo sapiens GN=ERP29 PE=1 SV=4 - [ERP29_HUMAN] | 3.182E7 | 2.852E7 | 1.558E7 | 1.411E7 | 7.503 | 7.455 | 7.193 | 7.150 | 7.479 | 7.171 | 0.308 |
| P78371 | T-complex protein 1 subunit beta OS=Homo sapiens GN=CCT2 PE=1 SV=4 - [TCPB_HUMAN] | 2.898E8 | 2.508E8 | 1.877E8 | 9.391E7 | 8.462 | 8.399 | 8.273 | 7.973 | 8.431 | 8.123 | 0.308 |
| P61586 | Transforming protein RhoA OS=Homo sapiens GN=RHOA PE=1 SV=1 - [RHOA_HUMAN] | 1.718E7 | 1.062E7 | 6.600E6 | 6.745E6 | 7.235 | 7.026 | 6.820 | 6.829 | 7.131 | 6.824 | 0.306 |
| P26368-2 | Isoform 2 of Splicing factor U2AF 65 kDa subunit OS=Homo sapiens GN=U2AF2 - [U2AF2_HUMAN] | 8.726E7 | 8.389E7 | 7.403E7 | 2.425E7 | 7.941 | 7.924 | 7.869 | 7.385 | 7.932 | 7.627 | 0.305 |
| G5E9U0 | Myo-inositol 1-phosphate synthase A1, isoform CRA_a OS=Homo sapiens GN=ISYNA1 PE=4 SV=1 - [G5E9U0 | 8.234E6 | 1.077E7 | | 4.665E6 | 6.916 | 7.032 | | 6.669 | 6.974 | 6.669 | 0.305 |
| Q9BX68 | Histidine triad nucleotide-binding protein 2, mitochondrial OS=Homo sapiens GN=HINT2 PE=1 SV=1 - [HINT | 4.986E7 | 3.815E7 | 2.003E7 | 2.337E7 | 7.698 | 7.581 | 7.302 | 7.369 | 7.640 | 7.335 | 0.304 |
| O95400 | CD2 antigen cytoplasmic tail-binding protein 2 OS=Homo sapiens GN=CD2BP2 PE=1 SV=1 - [CD2B2_HUMAN | 5.749E6 | 3.793E6 | | 2.319E6 | 6.760 | 6.579 | | 6.365 | 6.669 | 6.365 | 0.304 |
| O95347-2 | Isoform 2 of Structural maintenance of chromosomes protein 2 OS=Homo sapiens GN=SMC2 - [SMC2_HUMA | 1.646E7 | 1.170E7 | 7.338E6 | 6.497E6 | 7.216 | 7.068 | 6.866 | 6.813 | 7.142 | 6.839 | 0.303 |
| Q86VP6-2 | Isoform 2 of Cullin-associated NEDD8-dissociated protein 1 OS=Homo sapiens GN=CAND1 - [CAND1_HUMAN | 1.902E7 | 1.268E7 | 8.157E6 | 7.407E6 | 7.279 | 7.103 | 6.912 | 6.870 | 7.191 | 6.891 | 0.301 |
| P15927 | Replication protein A 32 kDa subunit OS=Homo sapiens GN=RPA2 PE=1 SV=1 - [RFA2_HUMAN] | 1.658E7 | 1.810E7 | 7.044E6 | 1.067E7 | 7.220 | 7.258 | 6.848 | 7.028 | 7.239 | 6.938 | 0.301 |
| B9A067 | Uncharacterized protein OS=Homo sapiens GN=IMMT PE=4 SV=2 - [B9A067_HUMAN] | 3.286E7 | 2.472E7 | 1.447E7 | 1.412E7 | 7.517 | 7.393 | 7.160 | 7.150 | 7.455 | 7.155 | 0.300 |
| P49411 | Elongation factor Tu, mitochondrial OS=Homo sapiens GN=TUFM PE=1 SV=2 - [EFTU_HUMAN] | 1.496E8 | 1.130E8 | 6.126E7 | 6.945E7 | 8.175 | 8.053 | 7.787 | 7.842 | 8.114 | 7.814 | 0.300 |
| P22392-2 | Isoform 3 of Nucleoside diphosphate kinase B OS=Homo sapiens GN=NME2 - [NDKB_HUMAN] | 5.621E8 | 3.140E8 | 1.617E8 | 2.774E8 | 8.750 | 8.497 | 8.209 | 8.443 | 8.623 | 8.326 | 0.297 |
| O75396 | Vesicle-trafficking protein SEC22b OS=Homo sapiens GN=SEC22B PE=1 SV=4 - [SC22B_HUMAN] | 2.233E7 | 1.646E7 | 1.076E7 | 8.692E6 | 7.349 | 7.216 | 7.032 | 6.939 | 7.283 | 6.985 | 0.297 |
| D6RBT8 | Uncharacterized protein OS=Homo sapiens GN=UCHL1 PE=4 SV=1 - [D6RBT8_HUMAN] | 7.233E7 | 4.856E7 | 2.304E7 | 3.885E7 | 7.859 | 7.686 | 7.363 | 7.589 | 7.773 | 7.476 | 0.297 |
| Q15942 | Zyxin OS=Homo sapiens GN=ZYX PE=1 SV=1 - [ZYX_HUMAN] | 2.945E7 | 1.520E7 | 1.267E7 | 9.026E6 | 7.469 | 7.182 | 7.103 | 6.955 | 7.325 | 7.029 | 0.296 |
| Q99798 | Aconitate hydratase, mitochondrial OS=Homo sapiens GN=ACO2 PE=1 SV=2 - [ACON_HUMAN] | 4.959E7 | 2.601E7 | 1.540E7 | 2.157E7 | 7.695 | 7.415 | 7.188 | 7.334 | 7.555 | 7.261 | 0.295 |
| O96008 | Mitochondrial import receptor subunit TOM40 homolog OS=Homo sapiens GN=TOMM40 PE=1 SV=1 - [TOM4 | 3.058E7 | 1.959E7 | 1.245E7 | | 7.485 | 7.292 | 7.095 | | 7.389 | 7.095 | 0.294 |
| B7Z1X8 | Uncharacterized protein OS=Homo sapiens GN=MAT2B PE=2 SV=1 - [B7Z1X8_HUMAN] | 9.693E6 | 1.108E7 | 6.794E6 | 4.100E6 | 6.986 | 7.044 | 6.832 | 6.613 | 7.015 | 6.722 | 0.293 |
| Q9HAV7 | GrpE protein homolog 1, mitochondrial OS=Homo sapiens GN=GRPEL1 PE=1 SV=2 - [GRPE1_HUMAN] | 3.015E7 | 2.102E7 | 9.170E6 | 1.793E7 | 7.479 | 7.323 | 6.962 | 7.254 | 7.401 | 7.108 | 0.293 |
| Q9Y3Y2-4 | Isoform 3 of Chromatin target of PRMT1 protein OS=Homo sapiens GN=CHTOP - [CHTOP_HUMAN] | 4.156E7 | 1.003E7 | 9.673E6 | 1.118E7 | 7.619 | 7.001 | 6.986 | 7.048 | 7.310 | 7.017 | 0.293 |
| E9PER8 | Uncharacterized protein OS=Homo sapiens GN=ARL6IP5 PE=4 SV=1 - [E9PER8_HUMAN] | | 1.316E7 | | 6.709E6 | | 7.119 | | 6.827 | 7.119 | 6.827 | 0.293 |
| B4DW74 | Uncharacterized protein OS=Homo sapiens GN=RAP1B PE=2 SV=1 - [B4DW74_HUMAN] | 1.137E7 | 1.008E7 | 5.689E6 | 5.261E6 | 7.056 | 7.004 | 6.755 | 6.721 | 7.030 | 6.738 | 0.292 |
| B7Z7G2 | Uncharacterized protein OS=Homo sapiens GN=FDXR PE=2 SV=1 - [B7Z7G2_HUMAN] | | 5.292E6 | 1.846E6 | 4.039E6 | | 6.724 | 6.266 | 6.606 | 6.724 | 6.436 | 0.287 |
| O75643 | U5 small nuclear ribonucleoprotein 200 kDa helicase OS=Homo sapiens GN=SNRNP200 PE=1 SV=2 - [U520_ | 1.093E7 | 8.448E6 | 5.822E6 | 4.225E6 | 7.039 | 6.927 | 6.765 | 6.626 | 6.983 | 6.695 | 0.287 |
| Q9BT78 | COP9 signalosome complex subunit 4 OS=Homo sapiens GN=COPS4 PE=1 SV=1 - [CSN4_HUMAN] | 1.422E7 | 8.967E6 | 4.280E6 | 7.950E6 | 7.153 | 6.953 | 6.631 | 6.900 | 7.053 | 6.766 | 0.287 |
| Q9H1E3 | Nuclear ubiquitous casein and cyclin-dependent kinases substrate OS=Homo sapiens GN=NUCKS1 PE=1 SV= | 1.038E8 | 5.142E7 | 3.617E7 | 3.978E7 | 8.016 | 7.711 | 7.558 | 7.600 | 7.864 | 7.579 | 0.285 |
| P07954-2 | Isoform Cytoplasmic of Fumarate hydratase, mitochondrial OS=Homo sapiens GN=FH - [FUMH_HUMAN] | 3.813E7 | 3.379E7 | 1.753E7 | 1.981E7 | 7.581 | 7.529 | 7.244 | 7.297 | 7.555 | 7.270 | 0.285 |
| Q9NVI7-2 | Isoform 2 of ATPase family AAA domain-containing protein 3A OS=Homo sapiens GN=ATAD3A - [ATD3A_HUN | 2.220E7 | 1.416E7 | 8.771E6 | 9.730E6 | 7.346 | 7.151 | 6.943 | 6.988 | 7.249 | 6.966 | 0.283 |
| P40939 | Trifunctional enzyme subunit alpha, mitochondrial OS=Homo sapiens GN=HADHA PE=1 SV=2 - [ECHA_HUM/ | 1.972E7 | 1.949E7 | 8.815E6 | 1.185E7 | 7.295 | 7.290 | 6.945 | 7.074 | 7.292 | 7.010 | 0.283 |
| B4E0P8 | Uncharacterized protein OS=Homo sapiens GN=UBXN1 PE=2 SV=1 - [B4E0P8_HUMAN] | 2.646E7 | | 1.382E7 | | 7.423 | | 7.141 | | 7.423 | 7.141 | 0.282 |
| Q14739 | Lamin-B receptor OS=Homo sapiens GN=LBR PE=1 SV=2 - [LBR_HUMAN] | 3.018E7 | 2.746E7 | | 1.504E7 | 7.480 | 7.439 | | 7.177 | 7.459 | 7.177 | 0.282 |
| Q9Y277 | Voltage-dependent anion-selective channel protein 3 OS=Homo sapiens GN=VDAC3 PE=1 SV=1 - [VDAC3_HI | 2.472E7 | 1.360E7 | 9.959E6 | 9.238E6 | 7.393 | 7.134 | 6.998 | 6.966 | 7.263 | 6.982 | 0.281 |
| Q96H99 | Cortactin OS=Homo sapiens GN=CTTN PE=2 SV=1 - [Q96H99_HUMAN] | 2.679E7 | 1.423E7 | 1.115E7 | 9.382E6 | 7.428 | 7.153 | 7.047 | 6.972 | 7.291 | 7.010 | 0.281 |
| P09874 | Poly [ADP-ribose] polymerase 1 OS=Homo sapiens GN=PARP1 PE=1 SV=4 - [PARP1_HUMAN] | 2.285E8 | 1.897E8 | 1.024E8 | 1.163E8 | 8.359 | 8.278 | 8.010 | 8.066 | 8.319 | 8.038 | 0.280 |
| E7EPC9 | Uncharacterized protein OS=Homo sapiens GN=EIF4B PE=4 SV=1 - [E7EPC9_HUMAN] | 3.471E7 | 2.072E7 | 1.124E7 | 1.783E7 | 7.540 | 7.316 | 7.051 | 7.251 | 7.428 | 7.151 | 0.277 |
| P00505 | Aspartate aminotransferase, mitochondrial OS=Homo sapiens GN=GOT2 PE=1 SV=3 - [AATM_HUMAN] | 5.288E7 | 4.168E7 | 2.218E7 | 2.835E7 | 7.723 | 7.620 | 7.346 | 7.453 | 7.672 | 7.399 | 0.272 |
| B4E0V0 | Uncharacterized protein OS=Homo sapiens GN=PNPO PE=2 SV=1 - [B4E0V0_HUMAN] | 7.000E6 | 1.046E7 | 4.574E6 | | 6.845 | 7.019 | 6.660 | | 6.932 | 6.660 | 0.272 |
| E7EMJ6 | Uncharacterized protein OS=Homo sapiens GN=SGTA PE=4 SV=1 - [E7EMJ6_HUMAN] | 1.563E7 | 1.062E7 | 5.363E6 | 8.923E6 | 7.194 | 7.026 | 6.729 | 6.951 | 7.110 | 6.840 | 0.270 |
| Q5JTZ9 | Alanyl-tRNA synthetase, mitochondrial OS=Homo sapiens GN=AARS2 PE=1 SV=1 - [SYAM_HUMAN] | 6.715E6 | | 2.988E6 | 4.358E6 | 6.827 | | 6.475 | 6.639 | 6.827 | 6.557 | 0.270 |
| Q07666-2 | Isoform 2 of KH domain-containing, RNA-binding, signal transduction-associated protein 1 OS=Homo sapiens | 4.456E7 | 2.935E7 | 2.080E7 | 1.817E7 | 7.649 | 7.468 | 7.318 | 7.259 | 7.558 | 7.289 | 0.270 |
| E9PBK9 | Uncharacterized protein OS=Homo sapiens GN=GTPBP1 PE=4 SV=1 - [E9PBK9_HUMAN] | 5.587E6 | 3.520E6 | | 2.393E6 | 6.747 | 6.547 | | 6.379 | 6.647 | 6.379 | 0.268 |
| O43615 | Mitochondrial import inner membrane translocase subunit TIM44 OS=Homo sapiens GN=TIMM44 PE=1 SV=2 | 2.325E7 | 7.682E6 | 6.360E6 | 8.196E6 | 7.366 | 6.885 | 6.803 | 6.914 | 7.126 | 6.859 | 0.267 |
| P47985 | Cytochrome b-c1 complex subunit Rieske, mitochondrial OS=Homo sapiens GN=UQCRFS1 PE=1 SV=2 - [UCF | 1.568E7 | 1.602E7 | 8.743E6 | 8.448E6 | 7.195 | 7.205 | 6.942 | 6.927 | 7.200 | 6.934 | 0.266 |

| | | | | | | | | | | | | |
|---|---|---|---|---|---|---|---|---|---|---|---|---|
| P00558 | Phosphoglycerate kinase 1 OS=Homo sapiens GN=PGK1 PE=1 SV=3 - [PGK1_HUMAN] | 3.133E8 | 2.327E8 | 1.753E8 | 1.245E8 | 8.496 | 8.367 | 8.244 | 8.095 | 8.431 | 8.169 | 0.262 |
| C9JB50 | Ribosomal protein L28, isoform CRA_c OS=Homo sapiens GN=RPL28 PE=4 SV=1 - [C9JB50_HUMAN] | 7.762E7 | 5.860E7 | 3.411E7 | 3.992E7 | 7.890 | 7.768 | 7.533 | 7.601 | 7.829 | 7.567 | 0.262 |
| Q9Y3C6 | Peptidyl-prolyl cis-trans isomerase-like 1 OS=Homo sapiens GN=PPIL1 PE=1 SV=1 - [PPIL1_HUMAN] | 3.088E7 | 6.819E6 | 8.507E6 | 7.425E6 | 7.490 | 6.834 | 6.930 | 6.871 | 7.162 | 6.900 | 0.261 |
| E7EPA1 | Uncharacterized protein OS=Homo sapiens GN=PRPSAP2 PE=4 SV=1 - [E7EPA1_HUMAN] | 1.676E7 | 8.609E6 | | 6.581E6 | 7.224 | 6.935 | | 6.818 | 7.080 | 6.818 | 0.261 |
| P62851 | 40S ribosomal protein S25 OS=Homo sapiens GN=RPS25 PE=1 SV=1 - [RS25_HUMAN] | 7.658E7 | 3.544E7 | 2.776E7 | 2.941E7 | 7.884 | 7.550 | 7.443 | 7.469 | 7.717 | 7.456 | 0.261 |
| P00403 | Cytochrome c oxidase subunit 2 OS=Homo sapiens GN=MT-CO2 PE=1 SV=1 - [COX2_HUMAN] | 4.696E7 | 3.650E7 | 2.311E7 | 2.238E7 | 7.672 | 7.562 | 7.364 | 7.350 | 7.617 | 7.357 | 0.260 |
| F8W7I9 | Uncharacterized protein OS=Homo sapiens GN=RANGAP1 PE=4 SV=1 - [F8W7I9_HUMAN] | 2.172E7 | 1.545E7 | 7.804E6 | 1.299E7 | 7.337 | 7.189 | 6.892 | 7.114 | 7.263 | 7.003 | 0.260 |
| E7EWF1 | Uncharacterized protein OS=Homo sapiens GN=RPL4 PE=4 SV=1 - [E7EWF1_HUMAN] | 1.512E8 | 9.160E7 | 7.084E7 | 5.949E7 | 8.180 | 7.962 | 7.850 | 7.774 | 8.071 | 7.812 | 0.258 |
| E7ES08 | Uncharacterized protein OS=Homo sapiens GN=HMGB3 PE=4 SV=1 - [E7ES08_HUMAN] | 4.383E7 | 2.926E7 | 2.048E7 | 1.906E7 | 7.642 | 7.466 | 7.311 | 7.280 | 7.554 | 7.296 | 0.258 |
| P52758 | Ribonuclease UK114 OS=Homo sapiens GN=HRSP12 PE=1 SV=1 - [UK114_HUMAN] | 1.869E7 | 1.073E7 | 6.233E6 | 9.817E6 | 7.272 | 7.031 | 6.795 | 6.992 | 7.151 | 6.893 | 0.258 |
| P13984 | General transcription factor IIF subunit 2 OS=Homo sapiens GN=GTF2F2 PE=1 SV=2 - [T2FB_HUMAN] | 6.102E6 | 1.013E6 | 1.462E6 | 1.291E6 | 6.785 | 6.006 | 6.165 | 6.111 | 6.396 | 6.138 | 0.258 |
| P04908 | Histone H2A type 1-B/E OS=Homo sapiens GN=HIST1H2AB PE=1 SV=2 - [H2A1B_HUMAN] | 1.070E8 | 9.208E7 | 5.099E7 | 5.910E7 | 8.030 | 7.964 | 7.707 | 7.772 | 7.997 | 7.740 | 0.257 |
| B4DWN1 | Uncharacterized protein OS=Homo sapiens GN=LMAN2 PE=2 SV=1 - [B4DWN1_HUMAN] | | 7.874E6 | 4.943E6 | 3.838E6 | | 6.896 | 6.694 | 6.584 | 6.896 | 6.639 | 0.257 |
| Q07065 | Cytoskeleton-associated protein 4 OS=Homo sapiens GN=CKAP4 PE=1 SV=2 - [CKAP4_HUMAN] | 2.045E7 | 1.278E7 | 8.958E6 | 8.975E6 | 7.311 | 7.107 | 6.952 | 6.953 | 7.209 | 6.953 | 0.256 |
| B8ZZT7 | Uncharacterized protein OS=Homo sapiens GN=HN1 PE=4 SV=1 - [B8ZZT7_HUMAN] | 3.751E7 | 2.920E7 | 1.925E7 | 1.750E7 | 7.574 | 7.465 | 7.285 | 7.243 | 7.520 | 7.264 | 0.256 |
| Q9Y262 | Eukaryotic translation initiation factor 3 subunit L OS=Homo sapiens GN=EIF3L PE=1 SV=1 - [EIF3L_HUMAN | 1.567E7 | 1.095E7 | 5.113E6 | 1.040E7 | 7.195 | 7.039 | 6.709 | 7.017 | 7.117 | 6.863 | 0.254 |
| P56537 | Eukaryotic translation initiation factor 6 OS=Homo sapiens GN=EIF6 PE=1 SV=1 - [IF6_HUMAN] | 3.121E7 | | 1.735E7 | 1.746E7 | 7.494 | | 7.239 | 7.242 | 7.494 | 7.241 | 0.254 |
| E7EQB9 | Uncharacterized protein OS=Homo sapiens GN=POLR1C PE=4 SV=1 - [E7EQB9_HUMAN] | 7.881E6 | | 4.424E6 | | 6.897 | | 6.646 | | 6.897 | 6.646 | 0.251 |
| B7Z6U8 | Uncharacterized protein OS=Homo sapiens GN=FHL1 PE=2 SV=1 - [B7Z6U8_HUMAN] | 3.226E7 | 1.639E7 | 1.247E7 | 1.347E7 | 7.509 | 7.215 | 7.096 | 7.129 | 7.362 | 7.113 | 0.249 |
| P49736 | DNA replication licensing factor MCM2 OS=Homo sapiens GN=MCM2 PE=1 SV=4 - [MCM2_HUMAN] | 1.871E7 | 1.848E7 | 6.964E6 | 1.579E7 | 7.272 | 7.267 | 6.843 | 7.198 | 7.269 | 7.021 | 0.249 |
| Q9Y2Z0-2 | Isoform 2 of Suppressor of G2 allele of SKP1 homolog OS=Homo sapiens GN=SUGT1 - [SUGT1_HUMAN] | 2.778E7 | 1.369E7 | 9.593E6 | 1.271E7 | 7.444 | 7.136 | 6.982 | 7.104 | 7.290 | 7.043 | 0.247 |
| E7EM48 | Uncharacterized protein OS=Homo sapiens GN=MCM4 PE=3 SV=1 - [E7EM48_HUMAN] | 3.181E7 | 2.809E7 | 1.422E7 | 2.015E7 | 7.503 | 7.449 | 7.153 | 7.304 | 7.476 | 7.229 | 0.247 |
| P62140 | Serine/threonine-protein phosphatase PP1-beta catalytic subunit OS=Homo sapiens GN=PPP1CB PE=1 SV=3 | 1.587E7 | 1.005E7 | 7.590E6 | 6.818E6 | 7.201 | 7.002 | 6.880 | 6.834 | 7.101 | 6.857 | 0.244 |
| E9PAQ6 | Uncharacterized protein OS=Homo sapiens GN=CCT3 PE=3 SV=1 - [E9PAQ6_HUMAN] | 1.304E8 | 1.100E8 | 5.183E7 | 8.986E7 | 8.115 | 8.041 | 7.715 | 7.954 | 8.078 | 7.834 | 0.244 |
| Q9UHV9 | Prefoldin subunit 2 OS=Homo sapiens GN=PFDN2 PE=1 SV=1 - [PFD2_HUMAN] | 6.975E7 | 3.071E7 | 2.392E7 | 2.911E7 | 7.844 | 7.487 | 7.379 | 7.464 | 7.665 | 7.421 | 0.244 |
| Q8IYB3-2 | Isoform 2 of Serine/arginine repetitive matrix protein 1 OS=Homo sapiens GN=SRRM1 - [SRRM1_HUMAN] | 3.103E7 | 1.271E7 | 9.854E6 | 1.303E7 | 7.492 | 7.104 | 6.994 | 7.115 | 7.298 | 7.054 | 0.244 |
| P68402 | Platelet-activating factor acetylhydrolase IB subunit beta OS=Homo sapiens GN=PAFAH1B2 PE=1 SV=1 - [PA | 2.361E7 | 2.301E7 | 1.246E7 | 1.434E7 | 7.373 | 7.362 | 7.096 | 7.156 | 7.368 | 7.126 | 0.241 |
| Q9C0C9 | Ubiquitin-conjugating enzyme E2 O OS=Homo sapiens GN=UBE2O PE=1 SV=3 - [UBE2O_HUMAN] | | 2.959E6 | | 1.699E6 | | 6.471 | | 6.230 | 6.471 | 6.230 | 0.241 |
| P45974-2 | Isoform Short of Ubiquitin carboxyl-terminal hydrolase 5 OS=Homo sapiens GN=USP5 - [UBP5_HUMAN] | 2.689E7 | 2.356E7 | 1.487E7 | 1.407E7 | 7.430 | 7.372 | 7.172 | 7.148 | 7.401 | 7.160 | 0.240 |
| P61024 | Cyclin-dependent kinases regulatory subunit 1 OS=Homo sapiens GN=CKS1B PE=1 SV=1 - [CKS1_HUMAN] | 1.135E7 | | 6.985E6 | 6.152E6 | 7.055 | | 6.844 | 6.789 | 7.055 | 6.817 | 0.238 |
| Q96AG4 | Leucine-rich repeat-containing protein 59 OS=Homo sapiens GN=LRRC59 PE=1 SV=1 - [LRC59_HUMAN] | 3.580E7 | 2.753E7 | 1.829E7 | 1.801E7 | 7.554 | 7.440 | 7.262 | 7.255 | 7.497 | 7.259 | 0.238 |
| P00441 | Superoxide dismutase [Cu-Zn] OS=Homo sapiens GN=SOD1 PE=1 SV=2 - [SODC_HUMAN] | 4.146E7 | 2.972E7 | 2.011E7 | 2.049E7 | 7.618 | 7.473 | 7.303 | 7.311 | 7.545 | 7.307 | 0.238 |
| Q13435 | Splicing factor 3B subunit 2 OS=Homo sapiens GN=SF3B2 PE=1 SV=2 - [SF3B2_HUMAN] | 6.443E7 | 3.727E7 | 2.416E7 | 3.328E7 | 7.809 | 7.571 | 7.383 | 7.522 | 7.690 | 7.453 | 0.238 |
| P39748 | Flap endonuclease 1 OS=Homo sapiens GN=FEN1 PE=1 SV=1 - [FEN1_HUMAN] | 2.329E7 | 1.568E7 | 1.084E7 | 1.137E7 | 7.367 | 7.195 | 7.035 | 7.056 | 7.281 | 7.045 | 0.236 |
| E9PLL6 | Uncharacterized protein OS=Homo sapiens GN=RPL27A PE=3 SV=1 - [E9PLL6_HUMAN] | 9.148E7 | 5.731E7 | 3.401E7 | 5.234E7 | 7.961 | 7.758 | 7.532 | 7.719 | 7.860 | 7.625 | 0.235 |
| Q9UNZ2-6 | Isoform 4 of NSFL1 cofactor p47 OS=Homo sapiens GN=NSFL1C - [NSF1C_HUMAN] | 1.855E7 | 1.655E7 | 1.107E7 | 9.415E6 | 7.268 | 7.219 | 7.044 | 6.974 | 7.244 | 7.009 | 0.235 |
| Q9Y295 | Developmentally-regulated GTP-binding protein 1 OS=Homo sapiens GN=DRG1 PE=1 SV=1 - [DRG1_HUMAN | 1.064E7 | 9.698E6 | 3.908E6 | 9.039E6 | 7.027 | 6.987 | 6.592 | 6.956 | 7.007 | 6.774 | 0.233 |
| O75880 | Protein SCO1 homolog, mitochondrial OS=Homo sapiens GN=SCO1 PE=1 SV=1 - [SCO1_HUMAN] | 1.435E7 | 5.881E6 | 5.972E6 | 4.846E6 | 7.157 | 6.769 | 6.776 | 6.685 | 6.963 | 6.731 | 0.232 |
| F5H1E1 | Uncharacterized protein OS=Homo sapiens GN=COPE PE=4 SV=1 - [F5H1E1_HUMAN] | 2.321E7 | 2.024E7 | 1.440E7 | 1.122E7 | 7.366 | 7.306 | 7.158 | 7.050 | 7.336 | 7.104 | 0.232 |
| P31930 | Cytochrome b-c1 complex subunit 1, mitochondrial OS=Homo sapiens GN=UQCRC1 PE=1 SV=3 - [QCR1_HU | 3.124E7 | 2.923E7 | 1.603E7 | 1.961E7 | 7.495 | 7.466 | 7.205 | 7.293 | 7.480 | 7.249 | 0.232 |
| P48444 | Coatomer subunit delta OS=Homo sapiens GN=ARCN1 PE=1 SV=1 - [COPD_HUMAN] | | 1.486E7 | 6.434E6 | 1.182E7 | | 7.172 | 6.808 | 7.073 | 7.172 | 6.941 | 0.231 |
| Q9Y490 | Talin-1 OS=Homo sapiens GN=TLN1 PE=1 SV=3 - [TLN1_HUMAN] | 4.292E7 | 3.468E7 | 2.893E7 | 1.783E7 | 7.633 | 7.540 | 7.461 | 7.251 | 7.586 | 7.356 | 0.230 |
| O75223-2 | Isoform 2 of Gamma-glutamylcyclotransferase OS=Homo sapiens GN=GGCT - [GGCT_HUMAN] | | 1.129E7 | 5.852E6 | 7.558E6 | | 7.053 | 6.767 | 6.878 | 7.053 | 6.823 | 0.230 |
| P36551 | Coproporphyrinogen-III oxidase, mitochondrial OS=Homo sapiens GN=CPOX PE=1 SV=3 - [HEM6_HUMAN] | 1.204E7 | | 6.427E6 | 7.827E6 | 7.081 | | 6.808 | 6.894 | 7.081 | 6.851 | 0.230 |
| Q9UQ35 | Serine/arginine repetitive matrix protein 2 OS=Homo sapiens GN=SRRM2 PE=1 SV=2 - [SRRM2_HUMAN] | 2.224E7 | 1.401E7 | 1.018E7 | 1.067E7 | 7.347 | 7.146 | 7.008 | 7.028 | 7.247 | 7.018 | 0.229 |
| P30050 | 60S ribosomal protein L12 OS=Homo sapiens GN=RPL12 PE=1 SV=1 - [RL12_HUMAN] | 1.261E8 | 8.483E7 | 5.026E7 | 7.435E7 | 8.101 | 7.929 | 7.701 | 7.871 | 8.015 | 7.786 | 0.228 |
| F5H2L4 | Uncharacterized protein OS=Homo sapiens GN=ACOT13 PE=4 SV=1 - [F5H2L4_HUMAN] | 2.161E7 | 1.508E7 | | 1.067E7 | 7.335 | 7.178 | | 7.028 | 7.256 | 7.028 | 0.228 |
| P19105 | Myosin regulatory light chain 12A OS=Homo sapiens GN=MYL12A PE=1 SV=2 - [ML12A_HUMAN] | 5.204E7 | 2.550E7 | 2.094E7 | 2.217E7 | 7.716 | 7.407 | 7.321 | 7.346 | 7.561 | 7.333 | 0.228 |
| P12956 | X-ray repair cross-complementing protein 6 OS=Homo sapiens GN=XRCC6 PE=1 SV=2 - [XRCC6_HUMAN] | 1.296E8 | 6.738E7 | 5.401E7 | 5.678E7 | 8.112 | 7.829 | 7.733 | 7.754 | 7.970 | 7.743 | 0.227 |
| Q9NSD9 | Phenylalanyl-tRNA synthetase beta chain OS=Homo sapiens GN=FARSB PE=1 SV=3 - [SYFB_HUMAN] | 1.087E7 | 3.413E6 | 3.622E6 | | 7.036 | 6.533 | 6.559 | | 6.785 | 6.559 | 0.226 |
| C9J8S3 | Uncharacterized protein OS=Homo sapiens GN=RAB7A PE=3 SV=1 - [C9J8S3_HUMAN] | 2.495E7 | 2.007E7 | 1.021E7 | 1.766E7 | 7.397 | 7.303 | 7.009 | 7.247 | 7.350 | 7.128 | 0.222 |
| P33993 | DNA replication licensing factor MCM7 OS=Homo sapiens GN=MCM7 PE=1 SV=4 - [MCM7_HUMAN] | 2.009E7 | 1.857E7 | 9.248E6 | 1.453E7 | 7.303 | 7.269 | 6.966 | 7.162 | 7.286 | 7.064 | 0.222 |
| Q00610-2 | Isoform 2 of Clathrin heavy chain 1 OS=Homo sapiens GN=CLTC - [CLH1_HUMAN] | 9.875E7 | 4.220E7 | 3.593E7 | 4.193E7 | 7.995 | 7.625 | 7.555 | 7.622 | 7.810 | 7.589 | 0.221 |
| P27816-6 | Isoform 6 of Microtubule-associated protein 4 OS=Homo sapiens GN=MAP4 - [MAP4_HUMAN] | 3.978E7 | 3.706E7 | 2.335E7 | 2.283E7 | 7.600 | 7.569 | 7.368 | 7.359 | 7.584 | 7.363 | 0.221 |
| O14828-2 | Isoform 2 of Secretory carrier-associated membrane protein 3 OS=Homo sapiens GN=SCAMP3 - [SCAM3_HUI | 1.599E7 | 8.526E6 | | 7.024E6 | 7.204 | 6.931 | | 6.847 | 7.067 | 6.847 | 0.221 |
| P61163 | Alpha-centractin OS=Homo sapiens GN=ACTR1A PE=1 SV=1 - [ACTZ_HUMAN] | 3.089E7 | 2.150E7 | 1.506E7 | 1.596E7 | 7.490 | 7.332 | 7.178 | 7.203 | 7.411 | 7.190 | 0.221 |
| O75531 | Barrier-to-autointegration factor OS=Homo sapiens GN=BANF1 PE=1 SV=1 - [BAF_HUMAN] | 1.245E8 | 7.205E7 | 6.414E7 | 5.063E7 | 8.095 | 7.858 | 7.807 | 7.704 | 7.976 | 7.756 | 0.221 |
| F8VY35 | Uncharacterized protein OS=Homo sapiens GN=NAP1L1 PE=3 SV=1 - [F8VY35_HUMAN] | 1.432E8 | 1.188E8 | 6.280E7 | 9.821E7 | 8.156 | 8.075 | 7.798 | 7.992 | 8.115 | 7.895 | 0.220 |
| P61204 | ADP-ribosylation factor 3 OS=Homo sapiens GN=ARF3 PE=1 SV=2 - [ARF3_HUMAN] | 4.907E7 | 5.594E7 | 2.235E7 | 4.457E7 | 7.691 | 7.748 | 7.349 | 7.649 | 7.719 | 7.499 | 0.220 |
| P63173 | 60S ribosomal protein L38 OS=Homo sapiens GN=RPL38 PE=1 SV=2 - [RL38_HUMAN] | 1.367E8 | 7.435E7 | 5.809E7 | 6.353E7 | 8.136 | 7.871 | 7.764 | 7.803 | 8.004 | 7.784 | 0.220 |
| Q9BZX2 | Uridine-cytidine kinase 2 OS=Homo sapiens GN=UCK2 PE=1 SV=1 - [UCK2_HUMAN] | 5.360E6 | | 3.231E6 | | 6.729 | | 6.509 | | 6.729 | 6.509 | 0.220 |
| P12004 | Proliferating cell nuclear antigen OS=Homo sapiens GN=PCNA PE=1 SV=1 - [PCNA_HUMAN] | 1.500E8 | 1.243E8 | 7.727E7 | 8.768E7 | 8.176 | 8.095 | 7.888 | 7.943 | 8.135 | 7.915 | 0.220 |
| Q5QPE4 | Sorting nexin 5 (Fragment) OS=Homo sapiens GN=SNX5 PE=2 SV=1 - [Q5QPE4_HUMAN] | 1.612E7 | 1.112E7 | 1.154E7 | 5.648E6 | 7.207 | 7.046 | 7.062 | 6.752 | 7.127 | 6.907 | 0.220 |
| B4DDD2 | Uncharacterized protein OS=Homo sapiens GN=DHFR PE=2 SV=1 - [B4DDD2_HUMAN] | 3.837E7 | 1.367E7 | 1.378E7 | 1.385E7 | 7.584 | 7.136 | 7.139 | 7.142 | 7.360 | 7.140 | 0.219 |
| P55072 | Transitional endoplasmic reticulum ATPase OS=Homo sapiens GN=VCP PE=1 SV=4 - [TERA_HUMAN] | 1.247E8 | 6.568E7 | 5.188E7 | 5.790E7 | 8.096 | 7.817 | 7.715 | 7.763 | 7.957 | 7.739 | 0.218 |
| Q9BSD7 | Cancer-related nucleoside-triphosphatase OS=Homo sapiens GN=NTPCR PE=1 SV=1 - [NTPCR_HUMAN] | 2.179E7 | 1.578E7 | 1.197E7 | 1.054E7 | 7.338 | 7.198 | 7.078 | 7.023 | 7.268 | 7.051 | 0.218 |
| Q01105 | Protein SET OS=Homo sapiens GN=SET PE=1 SV=3 - [SET_HUMAN] | 3.343E8 | 2.669E8 | 1.946E8 | 1.684E8 | 8.524 | 8.426 | 8.289 | 8.226 | 8.475 | 8.258 | 0.217 |
| P13693 | Translationally-controlled tumor protein OS=Homo sapiens GN=TPT1 PE=1 SV=1 - [TCTP_HUMAN] | 9.504E7 | 7.433E7 | 4.003E7 | 6.483E7 | 7.978 | 7.871 | 7.602 | 7.812 | 7.925 | 7.707 | 0.217 |
| Q8N1G4 | Leucine-rich repeat-containing protein 47 OS=Homo sapiens GN=LRRC47 PE=1 SV=1 - [LRC47_HUMAN] | 1.459E7 | 9.187E6 | 6.000E6 | 8.236E6 | 7.164 | 6.963 | 6.778 | 6.916 | 7.064 | 6.847 | 0.217 |
| E7EPK1 | Uncharacterized protein OS=Homo sapiens GN=SEPT7 PE=3 SV=1 - [E7EPK1_HUMAN] | 3.276E7 | 2.254E7 | 1.757E7 | 1.553E7 | 7.515 | 7.353 | 7.245 | 7.191 | 7.434 | 7.218 | 0.216 |
| P08579 | U2 small nuclear ribonucleoprotein B'' OS=Homo sapiens GN=SNRPB2 PE=1 SV=1 - [RU2B_HUMAN] | 1.773E7 | 1.245E7 | 7.506E6 | 1.088E7 | 7.249 | 7.095 | 6.875 | 7.037 | 7.172 | 6.956 | 0.216 |
| G3XAC6 | RNA-binding region (RNP1, RRM) containing 2, isoform CRA_f OS=Homo sapiens GN=RBM39 PE=4 SV=1 - [( | 2.027E7 | 8.700E6 | 8.332E6 | 7.843E6 | 7.307 | 6.940 | 6.921 | 6.894 | 7.123 | 6.908 | 0.216 |
| F5GYQ2 | Uncharacterized protein OS=Homo sapiens GN=TRMT112 PE=4 SV=1 - [F5GYQ2_HUMAN] | 2.482E7 | 1.299E7 | 9.865E6 | 1.214E7 | 7.395 | 7.114 | 6.994 | 7.084 | 7.254 | 7.039 | 0.215 |
| P55036 | 26S proteasome non-ATPase regulatory subunit 4 OS=Homo sapiens GN=PSMD4 PE=1 SV=1 - [PSMD4_HUM | 4.001E7 | 1.208E7 | 1.193E7 | 1.510E7 | 7.602 | 7.082 | 7.077 | 7.179 | 7.342 | 7.128 | 0.214 |
| P62314 | Small nuclear ribonucleoprotein Sm D1 OS=Homo sapiens GN=SNRPD1 PE=1 SV=1 - [SMD1_HUMAN] | 9.241E7 | 7.176E7 | 6.627E7 | 3.732E7 | 7.966 | 7.856 | 7.821 | 7.572 | 7.911 | 7.697 | 0.214 |
| Q14240 | Eukaryotic initiation factor 4A-II OS=Homo sapiens GN=EIF4A2 PE=1 SV=2 - [IF4A2_HUMAN] | 2.428E8 | 1.059E8 | 9.865E7 | 9.726E7 | 8.385 | 8.025 | 7.994 | 7.988 | 8.205 | 7.991 | 0.214 |
| C9JUG7 | Uncharacterized protein OS=Homo sapiens GN=CAPZA2 PE=4 SV=1 - [C9JUG7_HUMAN] | 4.951E7 | 2.424E7 | | 2.117E7 | 7.695 | 7.385 | | 7.326 | 7.540 | 7.326 | 0.214 |
| Q14157-4 | Isoform 4 of Ubiquitin-associated protein 2-like OS=Homo sapiens GN=UBAP2L - [UBP2L_HUMAN] | 2.309E7 | 1.726E7 | 1.213E7 | 1.227E7 | 7.363 | 7.237 | 7.084 | 7.089 | 7.300 | 7.086 | 0.214 |
| B4DXW1 | Uncharacterized protein OS=Homo sapiens GN=ACTR3 PE=2 SV=1 - [B4DXW1_HUMAN] | 1.067E7 | 6.999E6 | 5.040E6 | 5.552E6 | 7.028 | 6.845 | 6.702 | 6.744 | 6.937 | 6.723 | 0.213 |
| C9JFR7 | Uncharacterized protein OS=Homo sapiens GN=CYCS PE=3 SV=1 - [C9JFR7_HUMAN] | 2.960E7 | 2.921E7 | 1.730E7 | 1.884E7 | 7.471 | 7.466 | 7.238 | 7.275 | 7.468 | 7.257 | 0.212 |
| Q9BPW8 | Protein NipSnap homolog 1 OS=Homo sapiens GN=NIPSNAP1 PE=1 SV=1 - [NIPS1_HUMAN] | 2.721E7 | 1.896E7 | 1.343E7 | 1.450E7 | 7.435 | 7.278 | 7.128 | 7.161 | 7.356 | 7.145 | 0.211 |
| F5GZ27 | Lon protease homolog OS=Homo sapiens GN=LONP1 PE=3 SV=1 - [F5GZ27_HUMAN] | 3.276E7 | 3.220E7 | 2.352E7 | 1.695E7 | 7.515 | 7.508 | 7.372 | 7.229 | 7.512 | 7.300 | 0.211 |
| E7EU96 | Uncharacterized protein OS=Homo sapiens GN=CSNK2A1 PE=4 SV=1 - [E7EU96_HUMAN] | 3.309E7 | 1.962E7 | 1.736E7 | 1.427E7 | 7.520 | 7.293 | 7.239 | 7.154 | 7.406 | 7.197 | 0.209 |
| P23526 | Adenosylhomocysteinase OS=Homo sapiens GN=AHCY PE=1 SV=4 - [SAHH_HUMAN] | 6.171E7 | 3.971E7 | 2.673E7 | 3.507E7 | 7.790 | 7.599 | 7.427 | 7.545 | 7.695 | 7.486 | 0.209 |
| P30837 | Aldehyde dehydrogenase X, mitochondrial OS=Homo sapiens GN=ALDH1B1 PE=1 SV=3 - [AL1B1_HUMAN] | 2.046E7 | 1.567E7 | 9.851E6 | 1.248E7 | 7.311 | 7.195 | 6.993 | 7.096 | 7.253 | 7.045 | 0.208 |
| E5RHG8 | Uncharacterized protein OS=Homo sapiens GN=TCEB1 PE=4 SV=1 - [E5RHG8_HUMAN] | 2.900E7 | 1.391E7 | 1.657E7 | 9.347E6 | 7.462 | 7.143 | 7.219 | 6.971 | 7.303 | 7.095 | 0.208 |
| B4DVY1 | Uncharacterized protein OS=Homo sapiens GN=EIF3D PE=2 SV=1 - [B4DVY1_HUMAN] | 3.773E7 | 2.827E7 | 2.592E7 | 1.583E7 | 7.577 | 7.451 | 7.414 | 7.200 | 7.514 | 7.307 | 0.207 |
| P46940 | Ras GTPase-activating-like protein IQGAP1 OS=Homo sapiens GN=IQGAP1 PE=1 SV=1 - [IQGA1_HUMAN] | 1.523E7 | 1.045E7 | 1.054E7 | 5.821E6 | 7.183 | 7.019 | 7.023 | 6.765 | 7.101 | 6.894 | 0.207 |
| B4DVU3 | Uncharacterized protein OS=Homo sapiens GN=EIF3C PE=2 SV=1 - [B4DVU3_HUMAN] | 2.573E7 | 1.873E7 | 1.254E7 | 1.488E7 | 7.410 | 7.273 | 7.098 | 7.173 | 7.342 | 7.136 | 0.206 |
| Q07812-5 | Isoform Epsilon of Apoptosis regulator BAX OS=Homo sapiens GN=BAX - [BAX_HUMAN] | 1.967E7 | 1.233E7 | 9.619E6 | 9.811E6 | 7.294 | 7.091 | 6.983 | 6.992 | 7.192 | 6.987 | 0.205 |
| Q92945 | Far upstream element-binding protein 2 OS=Homo sapiens GN=KHSRP PE=1 SV=4 - [FUBP2_HUMAN] | 1.537E8 | 1.010E8 | 7.986E7 | 7.593E7 | 8.187 | 8.004 | 7.902 | 7.880 | 8.095 | 7.891 | 0.204 |
| Q16822 | Phosphoenolpyruvate carboxykinase [GTP], mitochondrial OS=Homo sapiens GN=PCK2 PE=1 SV=3 - [PCKGM | 1.815E7 | 1.515E7 | 8.874E6 | 1.212E7 | 7.259 | 7.180 | 6.948 | 7.083 | 7.220 | 7.016 | 0.204 |
| P24666 | Low molecular weight phosphotyrosine protein phosphatase OS=Homo sapiens GN=ACP1 PE=1 SV=3 - [PPA( | 1.357E7 | 1.020E7 | 6.300E6 | 8.597E6 | 7.133 | 7.009 | 6.799 | 6.934 | 7.071 | 6.867 | 0.204 |
| P61353 | 60S ribosomal protein L27 OS=Homo sapiens GN=RPL27 PE=1 SV=2 - [RL27_HUMAN] | 8.842E7 | 5.041E7 | 3.888E7 | 4.487E7 | 7.947 | 7.703 | 7.590 | 7.652 | 7.825 | 7.621 | 0.204 |
| O95433 | Activator of 90 kDa heat shock protein ATPase homolog 1 OS=Homo sapiens GN=AHSA1 PE=1 SV=1 - [AHSA | 1.428E7 | | 7.117E6 | 1.123E7 | 7.155 | | 6.852 | 7.051 | 7.155 | 6.951 | 0.203 |
| P04632 | Calpain small subunit 1 OS=Homo sapiens GN=CAPNS1 PE=1 SV=1 - [CPNS1_HUMAN] | 1.673E7 | 1.808E7 | 1.032E7 | 1.155E7 | 7.224 | 7.257 | 7.014 | 7.063 | 7.240 | 7.038 | 0.202 |
| E7EVE6 | Uncharacterized protein OS=Homo sapiens GN=CMPK1 PE=3 SV=1 - [E7EVE6_HUMAN] | 1.209E7 | 1.116E7 | 6.417E6 | 8.324E6 | 7.083 | 7.048 | 6.807 | 6.920 | 7.065 | 6.864 | 0.201 |
| Q05639 | Elongation factor 1-alpha 2 OS=Homo sapiens GN=EEF1A2 PE=1 SV=1 - [EF1A2_HUMAN] | 9.863E8 | 5.910E8 | 4.607E8 | 5.014E8 | 8.994 | 8.772 | 8.663 | 8.700 | 8.883 | 8.682 | 0.201 |
| P30084 | Enoyl-CoA hydratase, mitochondrial OS=Homo sapiens GN=ECHS1 PE=1 SV=4 - [ECHM_HUMAN] | 2.644E7 | 2.428E7 | 1.347E7 | 1.891E7 | 7.422 | 7.385 | 7.129 | 7.277 | 7.404 | 7.203 | 0.201 |
| P05388 | 60S acidic ribosomal protein P0 OS=Homo sapiens GN=RPLP0 PE=1 SV=1 - [RLA0_HUMAN] | 9.150E7 | 1.486E8 | 8.301E7 | 6.515E7 | 7.961 | 8.172 | 7.919 | 7.814 | 8.067 | 7.867 | 0.200 |
| P35579 | Myosin-9 OS=Homo sapiens GN=MYH9 PE=1 SV=4 - [MYH9_HUMAN] | 5.289E7 | 3.227E7 | 3.091E7 | 2.197E7 | 7.723 | 7.509 | 7.490 | 7.342 | 7.616 | 7.416 | 0.200 |
| P62263 | 40S ribosomal protein S14 OS=Homo sapiens GN=RPS14 PE=1 SV=3 - [RS14_HUMAN] | 9.901E7 | 7.016E7 | 5.128E7 | 5.413E7 | 7.996 | 7.846 | 7.710 | 7.733 | 7.921 | 7.722 | 0.199 |
| F8W651 | Uncharacterized protein OS=Homo sapiens GN=COPZ1 PE=4 SV=1 - [F8W651_HUMAN] | 3.678E7 | 2.920E7 | 2.383E7 | 1.802E7 | 7.566 | 7.465 | 7.377 | 7.256 | 7.516 | 7.316 | 0.199 |
| Q15637-4 | Isoform 4 of Splicing factor 1 OS=Homo sapiens GN=SF1 - [SF01_HUMAN] | 4.001E7 | 3.759E7 | 3.100E7 | 1.941E7 | 7.602 | 7.575 | 7.491 | 7.288 | 7.589 | 7.390 | 0.199 |
| Q13428-2 | Isoform 2 of Treacle protein OS=Homo sapiens GN=TCOF1 - [TCOF_HUMAN] | 1.285E7 | 6.304E6 | | 5.700E6 | 7.109 | 6.800 | | 6.756 | 6.954 | 6.756 | 0.198 |
| P31689 | DnaJ homolog subfamily A member 1 OS=Homo sapiens GN=DNAJA1 PE=1 SV=2 - [DNJA1_HUMAN] | 2.160E7 | 1.276E7 | 7.639E6 | 1.447E7 | 7.335 | 7.106 | 6.883 | 7.160 | 7.220 | 7.022 | 0.198 |
| P15880 | 40S ribosomal protein S2 OS=Homo sapiens GN=RPS2 PE=1 SV=2 - [RS2_HUMAN] | 1.354E8 | 8.110E7 | 5.665E7 | 7.774E7 | 8.132 | 7.909 | 7.753 | 7.891 | 8.020 | 7.822 | 0.198 |
| Q99536 | Synaptic vesicle membrane protein VAT-1 homolog OS=Homo sapiens GN=VAT1 PE=1 SV=2 - [VAT1_HUMAN | 3.010E7 | | 1.907E7 | | 7.479 | | 7.280 | | 7.479 | 7.280 | 0.198 |
| P42704 | Leucine-rich PPR motif-containing protein, mitochondrial OS=Homo sapiens GN=LRPPRC PE=1 SV=3 - [LPPR( | 3.202E7 | 2.709E7 | 2.202E7 | 1.582E7 | 7.505 | 7.433 | 7.343 | 7.199 | 7.469 | 7.271 | 0.198 |
| P23284 | Peptidyl-prolyl cis-trans isomerase B OS=Homo sapiens GN=PPIB PE=1 SV=2 - [PPIB_HUMAN] | 8.181E7 | 5.222E7 | 4.331E7 | 3.963E7 | 7.913 | 7.718 | 7.637 | 7.598 | 7.815 | 7.617 | 0.198 |
| Q8NCW5-2 | Isoform 2 of Apolipoprotein A-I-binding protein OS=Homo sapiens GN=APOA1BP - [AIBP_HUMAN] | 2.193E7 | 1.637E7 | 1.143E7 | 1.263E7 | 7.341 | 7.214 | 7.058 | 7.101 | 7.278 | 7.080 | 0.198 |
| P16152 | Carbonyl reductase [NADPH] 1 OS=Homo sapiens GN=CBR1 PE=1 SV=3 - [CBR1_HUMAN] | 3.385E7 | 2.482E7 | 1.809E7 | 1.871E7 | 7.529 | 7.395 | 7.258 | 7.272 | 7.462 | 7.265 | 0.197 |
| O60869-2 | Isoform 2 of Endothelial differentiation-related factor 1 OS=Homo sapiens GN=EDF1 - [EDF1_HUMAN] | 2.489E7 | 1.642E7 | 1.101E7 | 1.495E7 | 7.396 | 7.215 | 7.042 | 7.175 | 7.306 | 7.108 | 0.197 |
| P07900 | Heat shock protein HSP 90-alpha OS=Homo sapiens GN=HSP90AA1 PE=1 SV=5 - [HS90A_HUMAN] | 7.309E8 | 4.860E8 | 3.693E8 | 3.876E8 | 8.864 | 8.687 | 8.567 | 8.588 | 8.775 | 8.578 | 0.197 |
| F8VYE8 | Serine/threonine-protein phosphatase OS=Homo sapiens GN=PPP1CC PE=3 SV=1 - [F8VYE8_HUMAN] | 1.493E7 | 9.948E6 | 6.823E6 | 8.782E6 | 7.174 | 6.998 | 6.834 | 6.944 | 7.086 | 6.889 | 0.197 |
| Q96B49 | Mitochondrial import receptor subunit TOM6 homolog OS=Homo sapiens GN=TOMM6 PE=1 SV=1 - [TOM6_HUMAN] | | 3.456E7 | 2.861E7 | 1.686E7 | | 7.539 | 7.456 | 7.227 | 7.539 | 7.342 | 0.197 |
| O76094 | Signal recognition particle 72 kDa protein OS=Homo sapiens GN=SRP72 PE=1 SV=3 - [SRP72_HUMAN] | 1.141E7 | 9.247E6 | 6.660E6 | 6.404E6 | 7.057 | 6.966 | 6.823 | 6.806 | 7.012 | 6.815 | 0.197 |
| Q9Y3I0 | tRNA-splicing ligase RtcB homolog OS=Homo sapiens GN=C22orf28 PE=1 SV=1 - [RTCB_HUMAN] | 3.022E7 | 2.867E7 | 1.562E7 | 2.244E7 | 7.480 | 7.457 | 7.194 | 7.351 | 7.469 | 7.272 | 0.196 |
| Q13155 | Aminoacyl tRNA synthase complex-interacting multifunctional protein 2 OS=Homo sapiens GN=AIMP2 PE=1 S | 3.995E7 | 1.823E7 | 1.794E7 | 1.643E7 | 7.602 | 7.261 | 7.254 | 7.216 | 7.431 | 7.235 | 0.196 |
| A8KA46 | Ras suppressor protein 1 OS=Homo sapiens GN=RSU1 PE=2 SV=1 - [A8KA46_HUMAN] | 1.480E7 | 1.388E7 | 8.949E6 | 9.313E6 | 7.170 | 7.142 | 6.952 | 6.969 | 7.156 | 6.960 | 0.196 |
| Q15233 | Non-POU domain-containing octamer-binding protein OS=Homo sapiens GN=NONO PE=1 SV=4 - [NONO_HL | 1.477E8 | 1.281E8 | 8.156E7 | 9.414E7 | 8.169 | 8.107 | 7.911 | 7.974 | 8.138 | 7.943 | 0.196 |
| P07814 | Bifunctional aminoacyl-tRNA synthetase OS=Homo sapiens GN=EPRS PE=1 SV=5 - [SYEP_HUMAN] | 5.475E7 | 3.957E7 | 2.792E7 | 3.153E7 | 7.738 | 7.597 | 7.446 | 7.499 | 7.668 | 7.472 | 0.196 |
| B4DUN2 | Malate dehydrogenase OS=Homo sapiens GN=MDH1 PE=2 SV=1 - [B4DUN2_HUMAN] | 1.596E8 | 1.046E8 | 8.976E7 | 7.572E7 | 8.203 | 8.019 | 7.953 | 7.879 | 8.111 | 7.916 | 0.195 |
| F6SBX2 | Uncharacterized protein OS=Homo sapiens GN=IARS2 PE=3 SV=1 - [F6SBX2_HUMAN] | 1.711E7 | 1.594E7 | 1.055E7 | | 7.233 | 7.203 | 7.023 | | 7.218 | 7.023 | 0.195 |
| Q15365 | Poly(rC)-binding protein 1 OS=Homo sapiens GN=PCBP1 PE=1 SV=2 - [PCBP1_HUMAN] | 1.098E8 | 9.010E7 | 6.064E7 | 6.663E7 | 8.041 | 7.955 | 7.783 | 7.824 | 7.998 | 7.803 | 0.194 |
| Q16777 | Histone H2A type 2-C OS=Homo sapiens GN=HIST2H2AC PE=1 SV=4 - [H2A2C_HUMAN] | 9.763E7 | 9.244E7 | 5.695E7 | 6.480E7 | 7.990 | 7.966 | 7.756 | 7.812 | 7.978 | 7.784 | 0.194 |
| P42167 | Lamina-associated polypeptide 2, isoforms beta/gamma OS=Homo sapiens GN=TMPO PE=1 SV=2 - [LAP2B_ | 6.868E7 | 4.145E7 | 3.774E7 | 3.085E7 | 7.837 | 7.617 | 7.577 | 7.489 | 7.727 | 7.533 | 0.194 |
| B7Z3X5 | Uncharacterized protein OS=Homo sapiens GN=PDHA1 PE=2 SV=1 - [B7Z3X5_HUMAN] | 2.439E7 | 8.330E6 | 9.356E6 | 8.907E6 | 7.387 | 6.921 | 6.971 | 6.950 | 7.154 | 6.960 | 0.194 |
| Q02790 | Peptidyl-prolyl cis-trans isomerase FKBP4 OS=Homo sapiens GN=FKBP4 PE=1 SV=3 - [FKBP4_HUMAN] | 9.335E7 | 7.108E7 | 5.016E7 | 5.443E7 | 7.970 | 7.852 | 7.700 | 7.736 | 7.911 | 7.718 | 0.193 |
| F8W1A4 | Uncharacterized protein OS=Homo sapiens GN=AK2 PE=3 SV=1 - [F8W1A4_HUMAN] | 1.783E8 | 1.130E8 | 9.006E7 | 9.254E7 | 8.251 | 8.053 | 7.955 | 7.966 | 8.152 | 7.960 | 0.192 |
| Q15046 | Lysyl-tRNA synthetase OS=Homo sapiens GN=KARS PE=1 SV=3 - [SYK_HUMAN] | 4.348E7 | 2.823E7 | 2.178E7 | 2.332E7 | 7.638 | 7.451 | 7.338 | 7.368 | 7.544 | 7.353 | 0.192 |
| P61981 | 14-3-3 protein gamma OS=Homo sapiens GN=YWHAG PE=1 SV=2 - [1433G_HUMAN] | 2.676E8 | 1.643E8 | 1.368E8 | 1.331E8 | 8.428 | 8.216 | 8.136 | 8.124 | 8.322 | 8.130 | 0.192 |
| P12270 | Nucleoprotein TPR OS=Homo sapiens GN=TPR PE=1 SV=3 - [TPR_HUMAN] | 1.486E7 | 1.280E7 | 9.854E6 | 7.997E6 | 7.172 | 7.107 | 6.994 | 6.903 | 7.140 | 6.948 | 0.191 |
| Q9BZZ5-2 | Isoform 2 of Apoptosis inhibitor 5 OS=Homo sapiens GN=API5 - [API5_HUMAN] | 2.461E7 | 1.974E7 | 1.303E7 | 1.546E7 | 7.391 | 7.295 | 7.115 | 7.189 | 7.343 | 7.152 | 0.191 |
| Q9BUF5 | Tubulin beta-6 chain OS=Homo sapiens GN=TUBB6 PE=1 SV=1 - [TBB6_HUMAN] | 6.172E8 | 4.731E8 | 3.357E8 | 3.610E8 | 8.790 | 8.675 | 8.526 | 8.558 | 8.733 | 8.542 | 0.191 |
| B4E1K7 | Uncharacterized protein OS=Homo sapiens GN=STOML2 PE=2 SV=1 - [B4E1K7_HUMAN] | 5.250E7 | 4.428E7 | 2.965E7 | 3.260E7 | 7.720 | 7.646 | 7.472 | 7.513 | 7.683 | 7.493 | 0.191 |
| B4DMT5 | Uncharacterized protein OS=Homo sapiens GN=EIF3F PE=2 SV=1 - [B4DMT5_HUMAN] | 1.786E7 | 1.636E7 | 8.675E6 | 1.405E7 | 7.252 | 7.214 | 6.938 | 7.148 | 7.233 | 7.043 | 0.190 |
| P62249 | 40S ribosomal protein S16 OS=Homo sapiens GN=RPS16 PE=1 SV=2 - [RS16_HUMAN] | 9.129E7 | 5.851E7 | 4.610E7 | 4.834E7 | 7.960 | 7.767 | 7.664 | 7.684 | 7.864 | 7.674 | 0.190 |
| P29401 | Transketolase OS=Homo sapiens GN=TKT PE=1 SV=3 - [TKT_HUMAN] | 2.285E8 | 1.739E8 | 1.193E8 | 1.391E8 | 8.359 | 8.240 | 8.077 | 8.143 | 8.299 | 8.110 | 0.190 |
| O43175 | D-3-phosphoglycerate dehydrogenase OS=Homo sapiens GN=PHGDH PE=1 SV=4 - [SERA_HUMAN] | 2.602E8 | 1.606E8 | 1.319E8 | 1.324E8 | 8.415 | 8.206 | 8.120 | 8.122 | 8.311 | 8.121 | 0.190 |
| B1AJY5 | Proteasome (Prosome, macropain) 26S subunit, non-ATPase, 10 OS=Homo sapiens GN=PSMD10 PE=4 SV=1 | 9.807E6 | 1.099E7 | 6.713E6 | | 6.992 | 7.041 | 6.827 | | 7.016 | 6.827 | 0.189 |
| O00151 | PDZ and LIM domain protein 1 OS=Homo sapiens GN=PDLIM1 PE=1 SV=4 - [PDLI1_HUMAN] | 3.019E7 | 2.633E7 | 1.883E7 | 1.773E7 | 7.480 | 7.420 | 7.275 | 7.249 | 7.450 | 7.262 | 0.188 |
| O00487 | 26S proteasome non-ATPase regulatory subunit 14 OS=Homo sapiens GN=PSMD14 PE=1 SV=1 - [PSDE_HUI | 4.165E7 | 3.730E7 | 2.106E7 | 3.101E7 | 7.620 | 7.572 | 7.324 | 7.491 | 7.596 | 7.408 | 0.188 |
| P62847-2 | Isoform 2 of 40S ribosomal protein S24 OS=Homo sapiens GN=RPS24 - [RS24_HUMAN] | 1.259E8 | 1.030E8 | 7.026E7 | 7.767E7 | 8.100 | 8.013 | 7.847 | 7.890 | 8.057 | 7.868 | 0.188 |
| P54577 | Tyrosyl-tRNA synthetase, cytoplasmic OS=Homo sapiens GN=YARS PE=1 SV=4 - [SYYC_HUMAN] | 9.409E7 | 7.357E7 | 5.379E7 | 5.424E7 | 7.974 | 7.867 | 7.731 | 7.734 | 7.920 | 7.733 | 0.188 |

| | | | | | | | | | | | | |
|---|---|---|---|---|---|---|---|---|---|---|---|---|
| Q9Y371 | Endophilin-B1 OS=Homo sapiens GN=SH3GLB1 PE=1 SV=1 - [SHLB1_HUMAN] | 1.070E7 | | | 6.950E6 | 7.029 | | | 6.842 | 7.029 | 6.842 | 0.187 |
| O00425 | Insulin-like growth factor 2 mRNA-binding protein 3 OS=Homo sapiens GN=IGF2BP3 PE=1 SV=2 - [IF2B3_HU | 5.843E7 | 3.975E7 | 3.258E7 | 3.010E7 | 7.767 | 7.599 | 7.513 | 7.479 | 7.683 | 7.496 | 0.187 |
| E9PQK4 | Uncharacterized protein OS=Homo sapiens GN=CELF1 PE=4 SV=1 - [E9PQK4_HUMAN] | 7.520E6 | 4.367E6 | 3.881E6 | 3.572E6 | 6.876 | 6.640 | 6.589 | 6.553 | 6.758 | 6.571 | 0.187 |
| B3KPW0 | G-rich RNA sequence binding factor 1, isoform CRA_a OS=Homo sapiens GN=GRSF1 PE=2 SV=1 - [B3KPW0_ | 3.623E7 | 3.191E7 | 2.155E7 | 2.265E7 | 7.559 | 7.504 | 7.333 | 7.355 | 7.531 | 7.344 | 0.187 |
| P39656 | Dolichyl-diphosphooligosaccharide--protein glycosyltransferase 48 kDa subunit OS=Homo sapiens GN=DDOST | 3.299E7 | 3.600E7 | 2.168E7 | 2.323E7 | 7.518 | 7.556 | 7.336 | 7.366 | 7.537 | 7.351 | 0.186 |
| O95373 | Importin-7 OS=Homo sapiens GN=IPO7 PE=1 SV=1 - [IPO7_HUMAN] | 1.976E7 | 1.430E7 | 1.143E7 | 1.049E7 | 7.296 | 7.155 | 7.058 | 7.021 | 7.225 | 7.039 | 0.186 |
| O95831-3 | Isoform 3 of Apoptosis-inducing factor 1, mitochondrial OS=Homo sapiens GN=AIFM1 - [AIFM1_HUMAN] | 6.673E7 | 5.067E7 | 3.761E7 | 3.817E7 | 7.824 | 7.705 | 7.575 | 7.582 | 7.765 | 7.579 | 0.186 |
| P28066 | Proteasome subunit alpha type-5 OS=Homo sapiens GN=PSMA5 PE=1 SV=3 - [PSA5_HUMAN] | 1.222E8 | 7.584E7 | 6.164E7 | 6.383E7 | 8.087 | 7.880 | 7.790 | 7.805 | 7.983 | 7.797 | 0.186 |
| P14174 | Macrophage migration inhibitory factor OS=Homo sapiens GN=MIF PE=1 SV=4 - [MIF_HUMAN] | 8.262E8 | 6.288E8 | 4.482E8 | 4.944E8 | 8.917 | 8.798 | 8.652 | 8.694 | 8.858 | 8.673 | 0.185 |
| F2Z2J9 | Phosphoglycerate mutase OS=Homo sapiens PE=3 SV=1 - [F2Z2J9_HUMAN] | 3.317E8 | 1.814E8 | 1.628E8 | 1.579E8 | 8.521 | 8.259 | 8.212 | 8.198 | 8.390 | 8.205 | 0.185 |
| P35527 | Keratin, type I cytoskeletal 9 OS=Homo sapiens GN=KRT9 PE=1 SV=3 - [K1C9_HUMAN] | 2.770E8 | 2.133E8 | 1.585E8 | 1.594E8 | 8.442 | 8.329 | 8.200 | 8.202 | 8.386 | 8.201 | 0.184 |
| P08238 | Heat shock protein HSP 90-beta OS=Homo sapiens GN=HSP90AB1 PE=1 SV=4 - [HS90B_HUMAN] | 7.631E8 | 4.882E8 | 3.876E8 | 4.113E8 | 8.883 | 8.689 | 8.588 | 8.614 | 8.786 | 8.601 | 0.184 |
| P04264 | Keratin, type II cytoskeletal 1 OS=Homo sapiens GN=KRT1 PE=1 SV=6 - [K2C1_HUMAN] | 7.419E8 | 5.210E8 | 3.790E8 | 4.376E8 | 8.870 | 8.717 | 8.579 | 8.641 | 8.794 | 8.610 | 0.184 |
| Q5JYR6 | Ribophorin II OS=Homo sapiens GN=RPN2 PE=2 SV=1 - [Q5JYR6_HUMAN] | 2.958E7 | 2.333E7 | 1.586E7 | 1.868E7 | 7.471 | 7.368 | 7.200 | 7.271 | 7.419 | 7.236 | 0.184 |
| Q9H9B4 | Sideroflexin-1 OS=Homo sapiens GN=SFXN1 PE=1 SV=4 - [SFXN1_HUMAN] | 3.933E7 | 1.681E7 | 1.302E7 | 2.189E7 | 7.595 | 7.225 | 7.115 | 7.340 | 7.410 | 7.227 | 0.183 |
| B4DR31 | Uncharacterized protein OS=Homo sapiens GN=DPYSL2 PE=2 SV=1 - [B4DR31_HUMAN] | 2.016E7 | 1.468E7 | 9.597E6 | 1.331E7 | 7.305 | 7.167 | 6.982 | 7.124 | 7.236 | 7.053 | 0.183 |
| Q8WUM4 | Programmed cell death 6-interacting protein OS=Homo sapiens GN=PDCD6IP PE=1 SV=1 - [PDC6I_HUMAN] | 2.423E7 | 1.736E7 | 1.345E7 | 1.351E7 | 7.384 | 7.240 | 7.129 | 7.131 | 7.312 | 7.130 | 0.182 |
| Q99623 | Prohibitin-2 OS=Homo sapiens GN=PHB2 PE=1 SV=2 - [PHB2_HUMAN] | 1.391E8 | 9.298E7 | 7.792E7 | 7.181E7 | 8.143 | 7.968 | 7.892 | 7.856 | 8.056 | 7.874 | 0.182 |
| Q00341 | Vigilin OS=Homo sapiens GN=HDLBP PE=1 SV=2 - [VIGLN_HUMAN] | 2.203E7 | 1.911E7 | 1.224E7 | 1.490E7 | 7.343 | 7.281 | 7.088 | 7.173 | 7.312 | 7.130 | 0.182 |
| Q15102 | Platelet-activating factor acetylhydrolase IB subunit gamma OS=Homo sapiens GN=PAFAH1B3 PE=1 SV=1 - [ | 1.977E7 | 1.276E7 | 1.213E7 | 9.015E6 | 7.296 | 7.106 | 7.084 | 6.955 | 7.201 | 7.019 | 0.182 |
| E7EPE0 | Uncharacterized protein OS=Homo sapiens GN=ACAD9 PE=4 SV=1 - [E7EPE0_HUMAN] | 7.453E6 | 7.361E6 | 4.709E6 | 5.071E6 | 6.872 | 6.867 | 6.673 | 6.705 | 6.870 | 6.689 | 0.181 |
| O95777 | N-alpha-acetyltransferase 38, NatC auxiliary subunit OS=Homo sapiens GN=NAA38 PE=1 SV=3 - [NAA38_HU | 1.624E7 | 1.447E7 | 1.075E7 | 9.522E6 | 7.211 | 7.160 | 7.031 | 6.979 | 7.186 | 7.005 | 0.181 |
| Q86UE4 | Protein LYRIC OS=Homo sapiens GN=MTDH PE=1 SV=2 - [LYRIC_HUMAN] | 1.801E7 | 1.095E7 | | 9.270E6 | 7.256 | 7.040 | | 6.967 | 7.148 | 6.967 | 0.180 |
| F5H897 | Uncharacterized protein OS=Homo sapiens GN=TRAP1 PE=3 SV=1 - [F5H897_HUMAN] | 3.519E8 | 2.437E8 | 1.897E8 | 1.972E8 | 8.546 | 8.387 | 8.278 | 8.295 | 8.467 | 8.287 | 0.180 |
| P38117 | Electron transfer flavoprotein subunit beta OS=Homo sapiens GN=ETFB PE=1 SV=3 - [ETFB_HUMAN] | 4.525E7 | 2.999E7 | 2.433E7 | 2.439E7 | 7.656 | 7.477 | 7.386 | 7.387 | 7.566 | 7.387 | 0.180 |
| B7Z3U6 | ATPase, Na+/K+ transporting, alpha 1 polypeptide, isoform CRA_a OS=Homo sapiens GN=ATP1A1 PE=2 SV= | 3.907E7 | 2.850E7 | 1.848E7 | 2.637E7 | 7.592 | 7.455 | 7.267 | 7.421 | 7.523 | 7.344 | 0.179 |
| F5H2G1 | Uncharacterized protein OS=Homo sapiens GN=CHD4 PE=4 SV=1 - [F5H2G1_HUMAN] | 6.483E6 | 9.027E6 | 5.062E6 | | 6.812 | 6.956 | 6.704 | | 6.884 | 6.704 | 0.179 |
| P23921 | Ribonucleoside-diphosphate reductase large subunit OS=Homo sapiens GN=RRM1 PE=1 SV=1 - [RIR1_HUM/ | 1.545E7 | 1.128E7 | | 8.743E6 | 7.189 | 7.052 | | 6.942 | 7.121 | 6.942 | 0.179 |
| Q86V81 | THO complex subunit 4 OS=Homo sapiens GN=THOC4 PE=1 SV=3 - [THOC4_HUMAN] | 8.184E7 | 7.468E7 | 7.859E7 | 3.412E7 | 7.913 | 7.873 | 7.895 | 7.533 | 7.893 | 7.714 | 0.179 |
| P08195-2 | Isoform 2 of 4F2 cell-surface antigen heavy chain OS=Homo sapiens GN=SLC3A2 - [4F2_HUMAN] | 4.714E7 | 2.766E7 | 2.067E7 | 2.770E7 | 7.673 | 7.442 | 7.315 | 7.442 | 7.558 | 7.379 | 0.179 |
| B7Z9I1 | Uncharacterized protein OS=Homo sapiens GN=ACADM PE=2 SV=1 - [B7Z9I1_HUMAN] | 3.359E7 | 1.837E7 | 1.717E7 | 1.580E7 | 7.526 | 7.264 | 7.235 | 7.199 | 7.395 | 7.217 | 0.179 |
| P21333-2 | Isoform 2 of Filamin-A OS=Homo sapiens GN=FLNA - [FLNA_HUMAN] | 3.383E7 | 2.526E7 | 1.813E7 | 2.080E7 | 7.529 | 7.402 | 7.258 | 7.318 | 7.466 | 7.288 | 0.178 |
| Q16658 | Fascin OS=Homo sapiens GN=FSCN1 PE=1 SV=3 - [FSCN1_HUMAN] | 1.355E7 | 1.524E7 | 7.926E6 | 1.151E7 | 7.132 | 7.183 | 6.899 | 7.061 | 7.158 | 6.980 | 0.177 |
| P31040 | Succinate dehydrogenase [ubiquinone] flavoprotein subunit, mitochondrial OS=Homo sapiens GN=SDHA PE= | 4.029E7 | 3.626E7 | 2.988E7 | 2.164E7 | 7.605 | 7.559 | 7.475 | 7.335 | 7.582 | 7.405 | 0.177 |
| P06753-2 | Isoform 2 of Tropomyosin alpha-3 chain OS=Homo sapiens GN=TPM3 - [TPM3_HUMAN] | 2.687E8 | 9.470E7 | 1.126E8 | 1.003E8 | 8.429 | 7.976 | 8.051 | 8.001 | 8.203 | 8.026 | 0.176 |
| Q15836 | Vesicle-associated membrane protein 3 OS=Homo sapiens GN=VAMP3 PE=1 SV=3 - [VAMP3_HUMAN] | 9.117E6 | 3.703E6 | 3.733E6 | 4.017E6 | 6.960 | 6.569 | 6.572 | 6.604 | 6.764 | 6.588 | 0.176 |
| F8VV40 | Uncharacterized protein OS=Homo sapiens GN=LTA4H PE=4 SV=1 - [F8VV40_HUMAN] | 3.036E7 | 2.132E7 | 1.566E7 | 1.840E7 | 7.482 | 7.329 | 7.195 | 7.265 | 7.406 | 7.230 | 0.176 |
| C9JD32 | Uncharacterized protein OS=Homo sapiens GN=RPL23 PE=3 SV=1 - [C9JD32_HUMAN] | 6.198E7 | 3.733E7 | 3.284E7 | 3.137E7 | 7.792 | 7.572 | 7.516 | 7.496 | 7.682 | 7.506 | 0.176 |
| P41252 | Isoleucyl-tRNA synthetase, cytoplasmic OS=Homo sapiens GN=IARS PE=1 SV=2 - [SYIC_HUMAN] | 3.999E7 | 2.727E7 | 2.201E7 | 2.211E7 | 7.602 | 7.436 | 7.343 | 7.345 | 7.519 | 7.344 | 0.175 |
| P06493 | Cyclin-dependent kinase 1 OS=Homo sapiens GN=CDK1 PE=1 SV=3 - [CDK1_HUMAN] | 2.082E7 | 1.312E7 | 9.574E6 | 1.274E7 | 7.318 | 7.118 | 6.981 | 7.105 | 7.218 | 7.043 | 0.175 |
| P15374 | Ubiquitin carboxyl-terminal hydrolase isozyme L3 OS=Homo sapiens GN=UCHL3 PE=1 SV=1 - [UCHL3_HUMA | 3.662E7 | 2.544E7 | 1.935E7 | 2.150E7 | 7.564 | 7.406 | 7.287 | 7.333 | 7.485 | 7.310 | 0.175 |
| P24534 | Elongation factor 1-beta OS=Homo sapiens GN=EEF1B2 PE=1 SV=3 - [EF1B_HUMAN] | 9.080E7 | 6.361E7 | 4.863E7 | 5.315E7 | 7.958 | 7.804 | 7.687 | 7.725 | 7.881 | 7.706 | 0.175 |
| O60814 | Histone H2B type 1-K OS=Homo sapiens GN=HIST1H2BK PE=1 SV=3 - [H2B1K_HUMAN] | 4.957E7 | 4.739E7 | 3.681E7 | 2.858E7 | 7.695 | 7.676 | 7.566 | 7.456 | 7.685 | 7.511 | 0.174 |
| P62191 | 26S protease regulatory subunit 4 OS=Homo sapiens GN=PSMC1 PE=1 SV=1 - [PRS4_HUMAN] | 2.790E7 | 2.093E7 | 1.674E7 | 1.563E7 | 7.446 | 7.321 | 7.224 | 7.194 | 7.383 | 7.209 | 0.174 |
| Q9GZT3 | SRA stem-loop-interacting RNA-binding protein, mitochondrial OS=Homo sapiens GN=SLIRP PE=1 SV=1 - [SI | 3.025E7 | 2.625E7 | 1.793E7 | 1.988E7 | 7.481 | 7.419 | 7.254 | 7.298 | 7.450 | 7.276 | 0.174 |
| P00568 | Adenylate kinase isoenzyme 1 OS=Homo sapiens GN=AK1 PE=1 SV=3 - [KAD1_HUMAN] | 1.309E7 | 1.205E7 | 9.289E6 | 7.638E6 | 7.117 | 7.081 | 6.968 | 6.883 | 7.099 | 6.925 | 0.174 |
| Q15417 | Calponin-3 OS=Homo sapiens GN=CNN3 PE=1 SV=1 - [CNN3_HUMAN] | 2.744E7 | 1.036E7 | 8.440E6 | 1.515E7 | 7.438 | 7.015 | 6.926 | 7.180 | 7.227 | 7.053 | 0.173 |
| P09429 | High mobility group protein B1 OS=Homo sapiens GN=HMGB1 PE=1 SV=3 - [HMGB1_HUMAN] | 2.800E8 | 1.827E8 | 1.521E8 | 1.513E8 | 8.447 | 8.262 | 8.182 | 8.180 | 8.354 | 8.181 | 0.173 |
| P13010 | X-ray repair cross-complementing protein 5 OS=Homo sapiens GN=XRCC5 PE=1 SV=3 - [XRCC5_HUMAN] | 1.090E8 | 7.127E7 | 5.840E7 | 6.004E7 | 8.037 | 7.853 | 7.766 | 7.778 | 7.945 | 7.772 | 0.173 |
| F5H6N3 | Uncharacterized protein OS=Homo sapiens GN=CLTA PE=4 SV=1 - [F5H6N3_HUMAN] | 2.871E7 | 2.183E7 | 1.937E7 | 1.462E7 | 7.458 | 7.339 | 7.287 | 7.165 | 7.399 | 7.226 | 0.172 |
| P23246 | Splicing factor, proline- and glutamine-rich OS=Homo sapiens GN=SFPQ PE=1 SV=2 - [SFPQ_HUMAN] | 1.661E8 | 1.181E8 | 8.844E7 | 1.003E8 | 8.220 | 8.072 | 7.947 | 8.001 | 8.146 | 7.974 | 0.172 |
| P50395 | Rab GDP dissociation inhibitor beta OS=Homo sapiens GN=GDI2 PE=1 SV=2 - [GDIB_HUMAN] | 5.869E7 | 4.584E7 | 3.394E7 | 3.585E7 | 7.769 | 7.661 | 7.531 | 7.554 | 7.715 | 7.543 | 0.172 |
| P36542 | ATP synthase subunit gamma, mitochondrial OS=Homo sapiens GN=ATP5C1 PE=1 SV=1 - [ATPG_HUMAN] | 3.313E7 | 2.581E7 | 1.929E7 | 2.006E7 | 7.520 | 7.412 | 7.285 | 7.302 | 7.466 | 7.294 | 0.172 |
| P54727 | UV excision repair protein RAD23 homolog B OS=Homo sapiens GN=RAD23B PE=1 SV=1 - [RD23B_HUMAN] | 2.618E7 | 1.808E7 | 1.322E7 | 1.622E7 | 7.418 | 7.257 | 7.121 | 7.210 | 7.338 | 7.166 | 0.172 |
| P06733 | Alpha-enolase OS=Homo sapiens GN=ENO1 PE=1 SV=2 - [ENOA_HUMAN] | 1.419E9 | 1.054E9 | 8.468E8 | 8.012E8 | 9.152 | 9.023 | 8.928 | 8.904 | 9.087 | 8.916 | 0.172 |
| F5H721 | Uncharacterized protein OS=Homo sapiens GN=WBP11 PE=4 SV=1 - [F5H721_HUMAN] | | 1.902E6 | 1.372E6 | 1.196E6 | | 6.279 | 6.137 | 6.078 | 6.279 | 6.108 | 0.172 |
| Q9H0U4 | Ras-related protein Rab-1B OS=Homo sapiens GN=RAB1B PE=1 SV=1 - [RAB1B_HUMAN] | 3.174E7 | 2.156E7 | 1.772E7 | 1.753E7 | 7.502 | 7.334 | 7.248 | 7.244 | 7.418 | 7.246 | 0.172 |
| F5GWR2 | Uncharacterized protein OS=Homo sapiens GN=HSPD1 PE=3 SV=1 - [F5GWR2_HUMAN] | 5.747E8 | 4.564E8 | 3.377E8 | 3.530E8 | 8.759 | 8.659 | 8.529 | 8.548 | 8.709 | 8.538 | 0.171 |
| Q01082 | Spectrin beta chain, brain 1 OS=Homo sapiens GN=SPTBN1 PE=1 SV=2 - [SPTB2_HUMAN] | 1.104E7 | 3.666E6 | 4.657E6 | 3.957E6 | 7.043 | 6.564 | 6.668 | 6.597 | 6.804 | 6.633 | 0.171 |
| P06576 | ATP synthase subunit beta, mitochondrial OS=Homo sapiens GN=ATP5B PE=1 SV=3 - [ATPB_HUMAN] | 4.041E8 | 2.926E8 | 2.262E8 | 2.379E8 | 8.606 | 8.466 | 8.355 | 8.376 | 8.536 | 8.366 | 0.171 |
| B4E241 | Splicing factor, arginine/serine-rich 3, isoform CRA_a OS=Homo sapiens GN=SFRS3 PE=2 SV=1 - [B4E241_H | 9.660E7 | 6.292E7 | 6.929E7 | 3.997E7 | 7.985 | 7.799 | 7.841 | 7.602 | 7.892 | 7.721 | 0.171 |
| P11142 | Heat shock cognate 71 kDa protein OS=Homo sapiens GN=HSPA8 PE=1 SV=1 - [HSP7C_HUMAN] | 1.017E9 | 7.201E8 | 5.159E8 | 6.470E8 | 9.007 | 8.857 | 8.713 | 8.811 | 8.932 | 8.762 | 0.171 |
| E9PPH5 | Uncharacterized protein OS=Homo sapiens GN=ANP32E PE=4 SV=1 - [E9PPH5_HUMAN] | 3.782E7 | 2.055E7 | 1.840E7 | 1.926E7 | 7.578 | 7.313 | 7.265 | 7.285 | 7.445 | 7.275 | 0.170 |
| E9PC52 | Uncharacterized protein OS=Homo sapiens GN=RBBP7 PE=4 SV=1 - [E9PC52_HUMAN] | 6.694E7 | 5.509E7 | 4.047E7 | 4.160E7 | 7.826 | 7.741 | 7.607 | 7.619 | 7.783 | 7.613 | 0.170 |
| P25705 | ATP synthase subunit alpha, mitochondrial OS=Homo sapiens GN=ATP5A1 PE=1 SV=1 - [ATPA_HUMAN] | 2.658E8 | 1.426E8 | 1.474E8 | 1.175E8 | 8.425 | 8.154 | 8.169 | 8.070 | 8.289 | 8.119 | 0.170 |
| B4DDD6 | Uncharacterized protein OS=Homo sapiens GN=DBNL PE=2 SV=1 - [B4DDD6_HUMAN] | 1.680E7 | | 1.032E7 | 1.250E7 | 7.225 | | 7.014 | 7.097 | 7.225 | 7.055 | 0.170 |
| Q15008 | 26S proteasome non-ATPase regulatory subunit 6 OS=Homo sapiens GN=PSMD6 PE=1 SV=1 - [PSMD6_HUM | 3.144E7 | 2.198E7 | 1.827E7 | 1.731E7 | 7.497 | 7.342 | 7.262 | 7.238 | 7.420 | 7.250 | 0.170 |
| P38919 | Eukaryotic initiation factor 4A-III OS=Homo sapiens GN=EIF4A3 PE=1 SV=4 - [IF4A3_HUMAN] | 1.393E8 | 7.588E7 | 7.018E7 | 6.895E7 | 8.144 | 7.880 | 7.846 | 7.839 | 8.012 | 7.842 | 0.170 |
| P50402 | Emerin OS=Homo sapiens GN=EMD PE=1 SV=1 - [EMD_HUMAN] | 2.057E7 | 2.049E7 | 1.349E7 | 1.432E7 | 7.313 | 7.312 | 7.130 | 7.156 | 7.312 | 7.143 | 0.169 |
| Q86SX6 | Glutaredoxin-related protein 5, mitochondrial OS=Homo sapiens GN=GLRX5 PE=1 SV=2 - [GLRX5_HUMAN] | 1.215E7 | 7.089E6 | | 6.285E6 | 7.085 | 6.851 | | 6.798 | 6.968 | 6.798 | 0.169 |
| P39019 | 40S ribosomal protein S19 OS=Homo sapiens GN=RPS19 PE=1 SV=2 - [RS19_HUMAN] | 1.918E8 | 1.145E8 | 1.039E8 | 9.699E7 | 8.283 | 8.059 | 8.016 | 7.987 | 8.171 | 8.002 | 0.169 |
| Q9Y4Z0 | U6 snRNA-associated Sm-like protein LSm4 OS=Homo sapiens GN=LSM4 PE=1 SV=1 - [LSM4_HUMAN] | 2.072E7 | 1.970E7 | 1.161E7 | 1.620E7 | 7.316 | 7.295 | 7.065 | 7.210 | 7.306 | 7.137 | 0.168 |
| O14929-2 | Isoform B of Histone acetyltransferase type B catalytic subunit OS=Homo sapiens GN=HAT1 - [HAT1_HUMAN | 2.078E7 | 1.258E7 | 9.475E6 | 1.272E7 | 7.318 | 7.100 | 6.977 | 7.104 | 7.209 | 7.040 | 0.168 |
| P00491 | Purine nucleoside phosphorylase OS=Homo sapiens GN=PNP PE=1 SV=2 - [PNPH_HUMAN] | 3.252E7 | 1.825E7 | 1.780E7 | 1.539E7 | 7.512 | 7.261 | 7.250 | 7.187 | 7.387 | 7.219 | 0.168 |
| Q12904 | Aminoacyl tRNA synthase complex-interacting multifunctional protein 1 OS=Homo sapiens GN=AIMP1 PE=1 S | 1.309E7 | 1.576E7 | 7.409E6 | 1.286E7 | 7.117 | 7.198 | 6.870 | 7.109 | 7.157 | 6.989 | 0.168 |
| P49915 | GMP synthase [glutamine-hydrolyzing] OS=Homo sapiens GN=GMPS PE=1 SV=1 - [GUAA_HUMAN] | 4.594E7 | 4.207E7 | 2.859E7 | 3.121E7 | 7.662 | 7.624 | 7.456 | 7.494 | 7.643 | 7.475 | 0.168 |
| Q8TEX9 | Importin-4 OS=Homo sapiens GN=IPO4 PE=1 SV=2 - [IPO4_HUMAN] | 1.121E7 | 8.590E6 | 6.981E6 | 6.372E6 | 7.050 | 6.934 | 6.844 | 6.804 | 6.992 | 6.824 | 0.168 |
| A8K7X6 | Poly(RC) binding protein 2, isoform CRA_b OS=Homo sapiens GN=PCBP2 PE=2 SV=1 - [A8K7X6_HUMAN] | 9.158E7 | 9.666E7 | 5.866E7 | 6.973E7 | 7.962 | 7.985 | 7.768 | 7.843 | 7.974 | 7.806 | 0.168 |
| Q9Y230 | RuvB-like 2 OS=Homo sapiens GN=RUVBL2 PE=1 SV=3 - [RUVB2_HUMAN] | 6.197E7 | 4.383E7 | 3.426E7 | 3.666E7 | 7.792 | 7.642 | 7.535 | 7.564 | 7.717 | 7.549 | 0.168 |
| O00299 | Chloride intracellular channel protein 1 OS=Homo sapiens GN=CLIC1 PE=1 SV=4 - [CLIC1_HUMAN] | 6.346E7 | 5.040E7 | 3.876E7 | 3.824E7 | 7.802 | 7.702 | 7.588 | 7.583 | 7.752 | 7.585 | 0.167 |
| Q04917 | 14-3-3 protein eta OS=Homo sapiens GN=YWHAH PE=1 SV=4 - [1433F_HUMAN] | 1.642E8 | 1.036E8 | 9.354E7 | 8.436E7 | 8.215 | 8.015 | 7.971 | 7.926 | 8.115 | 7.949 | 0.167 |
| Q9Y383 | Putative RNA-binding protein Luc7-like 2 OS=Homo sapiens GN=LUC7L2 PE=1 SV=2 - [LC7L2_HUMAN] | 2.904E7 | 2.344E7 | 1.692E7 | 1.866E7 | 7.463 | 7.370 | 7.228 | 7.271 | 7.416 | 7.250 | 0.167 |
| P30086 | Phosphatidylethanolamine-binding protein 1 OS=Homo sapiens GN=PEBP1 PE=1 SV=3 - [PEBP1_HUMAN] | 2.181E8 | 1.439E8 | 1.163E8 | 1.257E8 | 8.339 | 8.158 | 8.065 | 8.099 | 8.248 | 8.082 | 0.166 |
| Q8N163 | Protein KIAA1967 OS=Homo sapiens GN=KIAA1967 PE=1 SV=2 - [K1967_HUMAN] | 1.033E7 | 1.062E7 | | 7.158E6 | 7.014 | 7.026 | | 6.855 | 7.020 | 6.855 | 0.165 |
| P35268 | 60S ribosomal protein L22 OS=Homo sapiens GN=RPL22 PE=1 SV=2 - [RL22_HUMAN] | 1.492E8 | 1.147E8 | 8.993E7 | 8.888E7 | 8.174 | 8.059 | 7.954 | 7.949 | 8.117 | 7.951 | 0.165 |
| E9PGQ7 | Uncharacterized protein OS=Homo sapiens GN=SNW1 PE=4 SV=2 - [E9PGQ7_HUMAN] | 1.290E7 | | 9.352E6 | 8.335E6 | 7.111 | | 6.971 | 6.921 | 7.111 | 6.946 | 0.165 |
| P20674 | Cytochrome c oxidase subunit 5A, mitochondrial OS=Homo sapiens GN=COX5A PE=1 SV=2 - [COX5A_HUMA | 7.288E7 | 6.653E7 | 4.508E7 | 5.043E7 | 7.863 | 7.823 | 7.654 | 7.703 | 7.843 | 7.678 | 0.164 |
| Q15819 | Ubiquitin-conjugating enzyme E2 variant 2 OS=Homo sapiens GN=UBE2V2 PE=1 SV=4 - [UB2V2_HUMAN] | 4.331E7 | 3.653E7 | 2.565E7 | 2.895E7 | 7.637 | 7.563 | 7.409 | 7.462 | 7.600 | 7.435 | 0.164 |
| O60841 | Eukaryotic translation initiation factor 5B OS=Homo sapiens GN=EIF5B PE=1 SV=4 - [IF2P_HUMAN] | 2.640E7 | 2.079E7 | 1.889E7 | 1.365E7 | 7.422 | 7.318 | 7.276 | 7.135 | 7.370 | 7.206 | 0.164 |
| P63000 | Ras-related C3 botulinum toxin substrate 1 OS=Homo sapiens GN=RAC1 PE=1 SV=1 - [RAC1_HUMAN] | | 7.713E6 | 4.898E6 | 5.704E6 | | 6.887 | 6.690 | 6.756 | 6.887 | 6.723 | 0.164 |
| O76003 | Glutaredoxin-3 OS=Homo sapiens GN=GLRX3 PE=1 SV=2 - [GLRX3_HUMAN] | 2.642E7 | 1.808E7 | 1.355E7 | 1.657E7 | 7.422 | 7.257 | 7.132 | 7.219 | 7.340 | 7.176 | 0.164 |
| P30046 | D-dopachrome decarboxylase OS=Homo sapiens GN=DDT PE=1 SV=3 - [DOPD_HUMAN] | 1.299E8 | 9.340E7 | 7.333E7 | 7.782E7 | 8.114 | 7.970 | 7.865 | 7.891 | 8.042 | 7.878 | 0.164 |
| P35232 | Prohibitin OS=Homo sapiens GN=PHB PE=1 SV=1 - [PHB_HUMAN] | 1.570E8 | 1.147E8 | 9.024E7 | 9.433E7 | 8.196 | 8.059 | 7.955 | 7.975 | 8.128 | 7.965 | 0.163 |
| P04843 | Dolichyl-diphosphooligosaccharide--protein glycosyltransferase subunit 1 OS=Homo sapiens GN=RPN1 PE=1 | 5.195E7 | 3.741E7 | 3.001E7 | 3.069E7 | 7.716 | 7.573 | 7.477 | 7.487 | 7.644 | 7.482 | 0.162 |
| O94826 | Mitochondrial import receptor subunit TOM70 OS=Homo sapiens GN=TOMM70A PE=1 SV=1 - [TOM70_HUM/ | 2.536E7 | 8.195E6 | 1.263E7 | 7.811E6 | 7.404 | 6.914 | 7.101 | 6.893 | 7.159 | 6.997 | 0.162 |
| Q9NZI8 | Insulin-like growth factor 2 mRNA-binding protein 1 OS=Homo sapiens GN=IGF2BP1 PE=1 SV=2 - [IF2B1_HU | 1.035E8 | 7.686E7 | 6.163E7 | 6.124E7 | 8.015 | 7.886 | 7.790 | 7.787 | 7.950 | 7.788 | 0.162 |
| Q15370 | Transcription elongation factor B polypeptide 2 OS=Homo sapiens GN=TCEB2 PE=1 SV=1 - [ELOB_HUMAN] | 9.751E6 | 1.250E7 | 7.534E6 | 7.684E6 | 6.989 | 7.097 | 6.877 | 6.886 | 7.043 | 6.881 | 0.162 |
| C9J8H9 | Uncharacterized protein OS=Homo sapiens GN=ATP5J2 PE=4 SV=1 - [C9J8H9_HUMAN] | 1.519E7 | 8.051E6 | 9.291E6 | 6.249E6 | 7.181 | 6.906 | 6.968 | 6.796 | 7.044 | 6.882 | 0.162 |
| P07737 | Profilin-1 OS=Homo sapiens GN=PFN1 PE=1 SV=2 - [PROF1_HUMAN] | 5.424E8 | 3.719E8 | 3.085E8 | 3.107E8 | 8.734 | 8.570 | 8.489 | 8.492 | 8.652 | 8.491 | 0.162 |
| P04350 | Tubulin beta-4A chain OS=Homo sapiens GN=TUBB4A PE=1 SV=2 - [TBB4A_HUMAN] | 1.246E9 | 9.728E8 | 7.487E8 | 7.703E8 | 9.096 | 8.988 | 8.874 | 8.887 | 9.042 | 8.880 | 0.161 |
| Q9Y5J9 | Mitochondrial import inner membrane translocase subunit Tim8 B OS=Homo sapiens GN=TIMM8B PE=1 SV=1 - [TIM8B_HUMAI | 1.153E7 | 8.335E6 | 7.586E6 | | | 7.062 | 6.921 | 6.880 | 7.062 | 6.900 | 0.161 |
| Q13200 | 26S proteasome non-ATPase regulatory subunit 2 OS=Homo sapiens GN=PSMD2 PE=1 SV=3 - [PSMD2_HUM | 3.201E7 | 1.768E7 | 1.799E7 | 1.498E7 | 7.505 | 7.247 | 7.255 | 7.176 | 7.376 | 7.215 | 0.161 |
| Q9Y333 | U6 snRNA-associated Sm-like protein LSm2 OS=Homo sapiens GN=LSM2 PE=1 SV=1 - [LSM2_HUMAN] | 2.059E7 | 2.035E7 | 1.136E7 | 1.757E7 | 7.314 | 7.308 | 7.055 | 7.245 | 7.311 | 7.150 | 0.161 |
| P62195 | 26S protease regulatory subunit 8 OS=Homo sapiens GN=PSMC5 PE=1 SV=1 - [PRS8_HUMAN] | 3.253E7 | 2.185E7 | 2.118E7 | 1.599E7 | 7.512 | 7.339 | 7.326 | 7.204 | 7.426 | 7.265 | 0.161 |
| P41091 | Eukaryotic translation initiation factor 2 subunit 3 OS=Homo sapiens GN=EIF2S3 PE=1 SV=3 - [IF2G_HUMAN | 3.456E7 | 2.156E7 | 2.001E7 | 1.778E7 | 7.539 | 7.334 | 7.301 | 7.250 | 7.436 | 7.276 | 0.161 |
| P11177-2 | Isoform 2 of Pyruvate dehydrogenase E1 component subunit beta, mitochondrial OS=Homo sapiens GN=PDH | 5.167E7 | 3.799E7 | 2.887E7 | 3.247E7 | 7.713 | 7.580 | 7.460 | 7.511 | 7.646 | 7.486 | 0.161 |
| Q13148 | TAR DNA-binding protein 43 OS=Homo sapiens GN=TARDBP PE=1 SV=1 - [TADBP_HUMAN] | 3.323E7 | 2.581E7 | 2.085E7 | 1.967E7 | 7.522 | 7.412 | 7.319 | 7.294 | 7.467 | 7.306 | 0.160 |
| Q13283 | Ras GTPase-activating protein-binding protein 1 OS=Homo sapiens GN=G3BP1 PE=1 SV=1 - [G3BP1_HUMAN | 4.577E7 | 3.221E7 | 2.424E7 | 2.909E7 | 7.661 | 7.508 | 7.384 | 7.464 | 7.584 | 7.424 | 0.160 |
| Q14204 | Cytoplasmic dynein 1 heavy chain 1 OS=Homo sapiens GN=DYNC1H1 PE=1 SV=5 - [DYHC1_HUMAN] | 3.548E7 | 2.367E7 | 2.010E7 | 2.006E7 | 7.550 | 7.374 | 7.303 | 7.302 | 7.462 | 7.303 | 0.159 |
| B7Z9L0 | T-complex protein 1 subunit delta OS=Homo sapiens GN=CCT4 PE=2 SV=1 - [B7Z9L0_HUMAN] | 1.217E8 | 9.652E7 | 7.344E7 | 7.696E7 | 8.085 | 7.985 | 7.866 | 7.886 | 8.035 | 7.876 | 0.159 |
| Q8WXF1-2 | Isoform 2 of Paraspeckle component 1 OS=Homo sapiens GN=PSPC1 - [PSPC1_HUMAN] | 3.505E7 | 2.552E7 | 2.135E7 | 2.020E7 | 7.545 | 7.407 | 7.329 | 7.305 | 7.476 | 7.317 | 0.158 |
| P31150 | Rab GDP dissociation inhibitor alpha OS=Homo sapiens GN=GDI1 PE=1 SV=2 - [GDIA_HUMAN] | 5.443E7 | 4.705E7 | 3.519E7 | 3.509E7 | 7.736 | 7.673 | 7.546 | 7.545 | 7.704 | 7.546 | 0.158 |
| G5E9B2 | Chaperonin containing TCP1, subunit 8 (Theta), isoform CRA_a OS=Homo sapiens GN=CCT8 PE=4 SV=1 - [C | 1.623E8 | 9.674E7 | 8.729E7 | 8.677E7 | 8.210 | 7.986 | 7.941 | 7.938 | 8.098 | 7.940 | 0.158 |
| O75947-2 | Isoform 2 of ATP synthase subunit d, mitochondrial OS=Homo sapiens GN=ATP5H - [ATP5H_HUMAN] | 5.502E7 | 3.476E7 | 2.824E7 | 3.270E7 | 7.740 | 7.541 | 7.451 | 7.515 | 7.641 | 7.483 | 0.158 |
| Q9Y2L1 | Exosome complex exonuclease RRP44 OS=Homo sapiens GN=DIS3 PE=1 SV=2 - [RRP44_HUMAN] | 1.596E7 | 9.455E6 | 9.530E6 | 7.649E6 | 7.203 | 6.976 | 6.979 | 6.884 | 7.089 | 6.931 | 0.158 |
| P27348 | 14-3-3 protein theta OS=Homo sapiens GN=YWHAQ PE=1 SV=1 - [1433T_HUMAN] | 1.998E8 | 1.218E8 | 1.107E8 | 1.062E8 | 8.301 | 8.086 | 8.044 | 8.026 | 8.193 | 8.035 | 0.158 |
| P68104 | Elongation factor 1-alpha 1 OS=Homo sapiens GN=EEF1A1 PE=1 SV=1 - [EF1A1_HUMAN] | 1.956E9 | 1.513E9 | 1.180E9 | 1.213E9 | 9.291 | 9.180 | 9.072 | 9.084 | 9.236 | 9.078 | 0.158 |
| [illegible] | [illegible] | [illegible] | [illegible] | [illegible] | [illegible] | [illegible] | [illegible] | [illegible] | [illegible] | [illegible] | [illegible] | [illegible] |
| P11908 | Ribose-phosphate pyrophosphokinase 2 OS=Homo sapiens GN=PRPS2 PE=1 SV=2 - [PRPS2_HUMAN] | 4.140E7 | 2.752E7 | 2.368E7 | 2.334E7 | 7.617 | 7.440 | 7.374 | 7.368 | 7.528 | 7.371 | 0.157 |
| P61970 | Nuclear transport factor 2 OS=Homo sapiens GN=NUTF2 PE=1 SV=1 - [NTF2_HUMAN] | 1.943E7 | 1.182E7 | 1.085E7 | 1.028E7 | 7.289 | 7.073 | 7.035 | 7.012 | 7.181 | 7.024 | 0.157 |
| P10809 | 60 kDa heat shock protein, mitochondrial OS=Homo sapiens GN=HSPD1 PE=1 SV=2 - [CH60_HUMAN] | 7.551E8 | 5.482E8 | 4.402E8 | 4.566E8 | 8.878 | 8.739 | 8.644 | 8.660 | 8.808 | 8.652 | 0.157 |
| P24752 | Acetyl-CoA acetyltransferase, mitochondrial OS=Homo sapiens GN=ACAT1 PE=1 SV=1 - [THIL_HUMAN] | 1.101E8 | 9.068E7 | 6.749E7 | 7.190E7 | 8.042 | 7.958 | 7.829 | 7.857 | 8.000 | 7.843 | 0.157 |
| B7Z254 | Uncharacterized protein OS=Homo sapiens GN=PDIA6 PE=2 SV=1 - [B7Z254_HUMAN] | 8.125E7 | 5.767E7 | 4.430E7 | 5.151E7 | 7.910 | 7.761 | 7.646 | 7.712 | 7.835 | 7.679 | 0.156 |
| E9PFH8 | Uncharacterized protein OS=Homo sapiens GN=SSB PE=4 SV=1 - [E9PFH8_HUMAN] | 1.024E8 | 8.725E7 | 6.443E7 | 6.754E7 | 8.010 | 7.941 | 7.809 | 7.830 | 7.976 | 7.819 | 0.156 |
| P60842 | Eukaryotic initiation factor 4A-I OS=Homo sapiens GN=EIF4A1 PE=1 SV=1 - [IF4A1_HUMAN] | 2.614E8 | 1.579E8 | 1.387E8 | 1.450E8 | 8.417 | 8.199 | 8.142 | 8.161 | 8.308 | 8.152 | 0.156 |
| C9J0K6 | Uncharacterized protein OS=Homo sapiens GN=SRI PE=4 SV=1 - [C9J0K6_HUMAN] | 9.511E6 | 6.832E6 | 4.940E6 | 6.421E6 | 6.978 | 6.835 | 6.694 | 6.808 | 6.906 | 6.751 | 0.156 |
| P05141 | ADP/ATP translocase 2 OS=Homo sapiens GN=SLC25A5 PE=1 SV=7 - [ADT2_HUMAN] | 1.152E8 | 8.059E7 | 6.493E7 | 6.986E7 | 8.062 | 7.906 | 7.812 | 7.844 | 7.984 | 7.828 | 0.156 |
| P62258 | 14-3-3 protein epsilon OS=Homo sapiens GN=YWHAE PE=1 SV=1 - [1433E_HUMAN] | 5.914E8 | 3.964E8 | 3.317E8 | 3.455E8 | 8.772 | 8.598 | 8.521 | 8.538 | 8.685 | 8.530 | 0.155 |
| P62861 | 40S ribosomal protein S30 OS=Homo sapiens GN=FAU PE=1 SV=1 - [RS30_HUMAN] | 6.820E7 | 4.026E7 | 3.354E7 | 4.006E7 | 7.834 | 7.605 | 7.526 | 7.603 | 7.719 | 7.564 | 0.155 |
| P31939 | Bifunctional purine biosynthesis protein PURH OS=Homo sapiens GN=ATIC PE=1 SV=3 - [PUR9_HUMAN] | 7.175E7 | 5.211E7 | 4.208E7 | 4.353E7 | 7.856 | 7.717 | 7.624 | 7.639 | 7.786 | 7.631 | 0.155 |
| P22695 | Cytochrome b-c1 complex subunit 2, mitochondrial OS=Homo sapiens GN=UQCRC2 PE=1 SV=3 - [QCR2_HU | 3.697E7 | 2.994E7 | 2.167E7 | 2.504E7 | 7.568 | 7.476 | 7.336 | 7.399 | 7.522 | 7.367 | 0.155 |
| D6RGI3 | Septin 11, isoform CRA_b OS=Homo sapiens GN=SEPT11 PE=3 SV=1 - [D6RGI3_HUMAN] | 3.601E7 | 2.395E7 | 1.967E7 | 2.152E7 | 7.556 | 7.379 | 7.294 | 7.333 | 7.468 | 7.313 | 0.155 |
| C9JSA3 | Uncharacterized protein OS=Homo sapiens GN=PDCD10 PE=4 SV=1 - [C9JSA3_HUMAN] | 1.590E7 | 6.953E6 | 9.462E6 | 5.734E6 | 7.201 | 6.842 | 6.976 | 6.758 | 7.022 | 6.867 | 0.155 |
| P49591 | Seryl-tRNA synthetase, cytoplasmic OS=Homo sapiens GN=SARS PE=1 SV=3 - [SYSC_HUMAN] | 1.659E7 | 1.569E7 | 1.128E7 | 1.133E7 | 7.220 | 7.196 | 7.052 | 7.054 | 7.208 | 7.053 | 0.154 |
| D6PSQ9 | Selenophosphate synthetase 1 +E9 variant OS=Homo sapiens GN=SEPHS1 PE=2 SV=1 - [D6PSQ9_HUMAN] | 1.110E7 | 1.388E7 | | 8.709E6 | 7.045 | 7.142 | | 6.940 | 7.094 | 6.940 | 0.154 |
| A6NGP5 | Uncharacterized protein OS=Homo sapiens GN=HN1L PE=2 SV=2 - [A6NGP5_HUMAN] | 3.824E7 | 2.510E7 | 2.037E7 | 2.321E7 | 7.583 | 7.400 | 7.309 | 7.366 | 7.491 | 7.337 | 0.154 |
| P13073 | Cytochrome c oxidase subunit 4 isoform 1, mitochondrial OS=Homo sapiens GN=COX4I1 PE=1 SV=1 - [COX4 | 4.850E7 | 2.271E7 | | 2.330E7 | 7.686 | 7.356 | | 7.367 | 7.521 | 7.367 | 0.154 |
| P42766 | 60S ribosomal protein L35 OS=Homo sapiens GN=RPL35 PE=1 SV=2 - [RL35_HUMAN] | 7.175E7 | 4.335E7 | 3.711E7 | 4.133E7 | 7.856 | 7.637 | 7.569 | 7.616 | 7.746 | 7.593 | 0.154 |
| F5H7U0 | 6-phosphogluconate dehydrogenase, decarboxylating OS=Homo sapiens GN=PGD PE=3 SV=1 - [F5H7U0_HU | 2.531E7 | 2.278E7 | 1.735E7 | 1.639E7 | 7.403 | 7.358 | 7.239 | 7.215 | 7.380 | 7.227 | 0.153 |
| Q00059 | Transcription factor A, mitochondrial OS=Homo sapiens GN=TFAM PE=1 SV=1 - [TFAM_HUMAN] | 1.280E7 | 1.247E7 | 8.283E6 | 9.506E6 | 7.107 | 7.096 | 6.918 | 6.978 | 7.101 | 6.948 | 0.153 |
| B4DT43 | Uncharacterized protein OS=Homo sapiens GN=ETFA PE=2 SV=1 - [B4DT43_HUMAN] | 8.672E7 | 7.160E7 | 5.443E7 | 5.630E7 | 7.938 | 7.855 | 7.736 | 7.750 | 7.897 | 7.743 | 0.153 |
| Q96I24 | Far upstream element-binding protein 3 OS=Homo sapiens GN=FUBP3 PE=1 SV=2 - [FUBP3_HUMAN] | 1.271E7 | 9.821E6 | 9.056E6 | 6.809E6 | 7.104 | 6.992 | 6.957 | 6.833 | 7.048 | 6.895 | 0.153 |
| O43809 | Cleavage and polyadenylation specificity factor subunit 5 OS=Homo sapiens GN=NUDT21 PE=1 SV=1 - [CPSI | 2.230E7 | 1.823E7 | 1.611E7 | 1.247E7 | 7.348 | 7.261 | 7.207 | 7.096 | 7.304 | 7.151 | 0.153 |
| E7EUT8 | Uncharacterized protein OS=Homo sapiens GN=SF3A3 PE=4 SV=1 - [E7EUT8_HUMAN] | 1.002E7 | 1.306E7 | 7.511E6 | 8.623E6 | 7.001 | 7.116 | 6.876 | 6.936 | 7.059 | 6.906 | 0.153 |
| P00492 | Hypoxanthine-guanine phosphoribosyltransferase OS=Homo sapiens GN=HPRT1 PE=1 SV=2 - [HPRT_HUMAI | 7.030E7 | 4.385E7 | 3.758E7 | 4.061E7 | 7.847 | 7.642 | 7.575 | 7.609 | 7.744 | 7.592 | 0.153 |
| Q9GZS3 | WD repeat-containing protein 61 OS=Homo sapiens GN=WDR61 PE=1 SV=1 - [WDR61_HUMAN] | 3.041E7 | 2.485E7 | 2.151E7 | 1.741E7 | 7.483 | 7.395 | 7.333 | 7.241 | 7.439 | 7.287 | 0.153 |

| | | | | | | | | | | | | |
|---|---|---|---|---|---|---|---|---|---|---|---|---|
| P51610-2 | Isoform 2 of Host cell factor 1 OS=Homo sapiens GN=HCFC1 - [HCFC1_HUMAN] | 1.327E7 | 8.548E6 | 7.935E6 | 7.084E6 | 7.123 | 6.932 | 6.900 | 6.850 | 7.027 | 6.875 | 0.152 |
| Q00688 | Peptidyl-prolyl cis-trans isomerase FKBP3 OS=Homo sapiens GN=FKBP3 PE=1 SV=1 - [FKBP3_HUMAN] | 7.271E7 | 3.587E7 | 3.998E7 | 3.234E7 | 7.862 | 7.555 | 7.602 | 7.510 | 7.708 | 7.556 | 0.152 |
| Q9Y266 | Nuclear migration protein nudC OS=Homo sapiens GN=NUDC PE=1 SV=1 - [NUDC_HUMAN] | 6.653E7 | 3.882E7 | 3.537E7 | 3.623E7 | 7.823 | 7.589 | 7.549 | 7.559 | 7.706 | 7.554 | 0.152 |
| P26196 | Probable ATP-dependent RNA helicase DDX6 OS=Homo sapiens GN=DDX6 PE=1 SV=2 - [DDX6_HUMAN] | 1.212E7 | 1.212E7 | 8.367E6 | 8.719E6 | 7.083 | 7.083 | 6.923 | 6.940 | 7.083 | 6.932 | 0.152 |
| F5GYN4 | Uncharacterized protein OS=Homo sapiens GN=OTUB1 PE=4 SV=1 - [F5GYN4_HUMAN] | 3.586E7 | 2.354E7 | 1.786E7 | 2.349E7 | 7.555 | 7.372 | 7.252 | 7.371 | 7.463 | 7.311 | 0.152 |
| Q9Y5L4 | Mitochondrial import inner membrane translocase subunit Tim13 OS=Homo sapiens GN=TIMM13 PE=1 SV=1 | 5.037E7 | 5.098E7 | 3.012E7 | 4.241E7 | 7.702 | 7.707 | 7.479 | 7.627 | 7.705 | 7.553 | 0.152 |
| P62753 | 40S ribosomal protein S6 OS=Homo sapiens GN=RPS6 PE=1 SV=1 - [RS6_HUMAN] | 1.855E8 | 1.274E8 | 1.057E8 | 1.115E8 | 8.268 | 8.105 | 8.024 | 8.047 | 8.187 | 8.036 | 0.151 |
| P27824 | Calnexin OS=Homo sapiens GN=CANX PE=1 SV=2 - [CALX_HUMAN] | 7.905E7 | 6.317E7 | 4.922E7 | 5.061E7 | 7.898 | 7.801 | 7.692 | 7.704 | 7.849 | 7.698 | 0.151 |
| B4E0X8 | Uncharacterized protein OS=Homo sapiens GN=FUBP1 PE=2 SV=1 - [B4E0X8_HUMAN] | 7.467E7 | 5.499E7 | 4.390E7 | 4.666E7 | 7.873 | 7.740 | 7.642 | 7.669 | 7.807 | 7.656 | 0.151 |
| P51148 | Ras-related protein Rab-5C OS=Homo sapiens GN=RAB5C PE=1 SV=2 - [RAB5C_HUMAN] | 1.434E7 | 1.630E7 | 1.136E7 | 1.026E7 | 7.156 | 7.212 | 7.055 | 7.011 | 7.184 | 7.033 | 0.151 |
| P13639 | Elongation factor 2 OS=Homo sapiens GN=EEF2 PE=1 SV=4 - [EF2_HUMAN] | 2.734E8 | 2.400E8 | 1.792E8 | 1.827E8 | 8.437 | 8.380 | 8.253 | 8.262 | 8.408 | 8.258 | 0.151 |
| P17174 | Aspartate aminotransferase, cytoplasmic OS=Homo sapiens GN=GOT1 PE=1 SV=3 - [AATC_HUMAN] | 2.546E7 | 2.001E7 | 1.494E7 | 1.706E7 | 7.406 | 7.301 | 7.174 | 7.232 | 7.354 | 7.203 | 0.150 |
| F5H698 | Uncharacterized protein OS=Homo sapiens GN=LARS PE=3 SV=1 - [F5H698_HUMAN] | 3.224E7 | 2.204E7 | 2.078E7 | 1.711E7 | 7.508 | 7.343 | 7.318 | 7.233 | 7.426 | 7.275 | 0.150 |
| F5H048 | Uncharacterized protein OS=Homo sapiens GN=SF3A1 PE=4 SV=1 - [F5H048_HUMAN] | 1.747E7 | 1.632E7 | 1.154E7 | 1.237E7 | 7.242 | 7.213 | 7.062 | 7.092 | 7.228 | 7.077 | 0.150 |
| Q15019 | Septin-2 OS=Homo sapiens GN=SEPT2 PE=1 SV=1 - [SEPT2_HUMAN] | 3.996E7 | 2.956E7 | 2.484E7 | 2.381E7 | 7.602 | 7.471 | 7.395 | 7.377 | 7.536 | 7.386 | 0.150 |
| P35998 | 26S protease regulatory subunit 7 OS=Homo sapiens GN=PSMC2 PE=1 SV=3 - [PRS7_HUMAN] | 2.812E7 | 2.261E7 | 1.998E7 | 1.594E7 | 7.449 | 7.354 | 7.301 | 7.203 | 7.402 | 7.252 | 0.150 |
| P50502 | Hsc70-interacting protein OS=Homo sapiens GN=ST13 PE=1 SV=2 - [F10A1_HUMAN] | 4.877E7 | 6.109E7 | 5.208E7 | 2.868E7 | 7.688 | 7.786 | 7.717 | 7.458 | 7.737 | 7.587 | 0.150 |
| P60866 | 40S ribosomal protein S20 OS=Homo sapiens GN=RPS20 PE=1 SV=1 - [RS20_HUMAN] | 1.570E8 | 7.982E7 | 7.737E7 | 8.144E7 | 8.196 | 7.902 | 7.889 | 7.911 | 8.049 | 7.900 | 0.149 |
| P07237 | Protein disulfide-isomerase OS=Homo sapiens GN=P4HB PE=1 SV=3 - [PDIA1_HUMAN] | 7.461E7 | 3.291E7 | 3.197E7 | 3.872E7 | 7.873 | 7.517 | 7.505 | 7.588 | 7.695 | 7.546 | 0.149 |
| P62937 | Peptidyl-prolyl cis-trans isomerase A OS=Homo sapiens GN=PPIA PE=1 SV=2 - [PPIA_HUMAN] | 7.176E8 | 6.016E8 | 4.304E8 | 5.056E8 | 8.856 | 8.779 | 8.634 | 8.704 | 8.818 | 8.669 | 0.149 |
| P24539 | ATP synthase subunit b, mitochondrial OS=Homo sapiens GN=ATP5F1 PE=1 SV=2 - [AT5F1_HUMAN] | 3.315E7 | 1.734E7 | 1.476E7 | 1.965E7 | 7.521 | 7.239 | 7.169 | 7.293 | 7.380 | 7.231 | 0.149 |
| P43490 | Nicotinamide phosphoribosyltransferase OS=Homo sapiens GN=NAMPT PE=1 SV=1 - [NAMPT_HUMAN] | 2.598E7 | 1.318E7 | 1.138E7 | 1.521E7 | 7.415 | 7.120 | 7.056 | 7.182 | 7.267 | 7.119 | 0.148 |
| P08107 | Heat shock 70 kDa protein 1A/1B OS=Homo sapiens GN=HSPA1A PE=1 SV=5 - [HSP71_HUMAN] | 1.066E9 | 7.669E8 | 6.453E8 | 6.412E8 | 9.028 | 8.885 | 8.810 | 8.807 | 8.956 | 8.808 | 0.148 |
| P37840-2 | Isoform 2-4 of Alpha-synuclein OS=Homo sapiens GN=SNCA - [SYUA_HUMAN] | 9.200E6 | 5.558E6 | 5.590E6 | 4.635E6 | 6.964 | 6.745 | 6.747 | 6.666 | 6.854 | 6.707 | 0.148 |
| P46781 | 40S ribosomal protein S9 OS=Homo sapiens GN=RPS9 PE=1 SV=3 - [RS9_HUMAN] | 3.392E7 | 1.877E7 | 1.630E7 | 1.984E7 | 7.530 | 7.273 | 7.212 | 7.298 | 7.402 | 7.255 | 0.147 |
| P43487 | Ran-specific GTPase-activating protein OS=Homo sapiens GN=RANBP1 PE=1 SV=1 - [RANG_HUMAN] | 1.444E8 | 9.791E7 | 7.967E7 | 9.017E7 | 8.159 | 7.991 | 7.901 | 7.955 | 8.075 | 7.928 | 0.147 |
| Q9NRX4 | 14 kDa phosphohistidine phosphatase OS=Homo sapiens GN=PHPT1 PE=1 SV=1 - [PHP14_HUMAN] | 5.811E7 | 4.934E7 | 3.740E7 | 3.900E7 | 7.764 | 7.693 | 7.573 | 7.591 | 7.729 | 7.582 | 0.147 |
| B4DWJ2 | Uncharacterized protein OS=Homo sapiens GN=QARS PE=2 SV=1 - [B4DWJ2_HUMAN] | 1.970E7 | 1.686E7 | 1.325E7 | 1.278E7 | 7.294 | 7.227 | 7.122 | 7.107 | 7.261 | 7.114 | 0.146 |
| P61106 | Ras-related protein Rab-14 OS=Homo sapiens GN=RAB14 PE=1 SV=4 - [RAB14_HUMAN] | 4.697E7 | 2.192E7 | 3.098E7 | 1.699E7 | 7.672 | 7.341 | 7.491 | 7.230 | 7.506 | 7.361 | 0.146 |
| F8W181 | 60S ribosomal protein L6 OS=Homo sapiens GN=RPL6 PE=3 SV=1 - [F8W181_HUMAN] | 1.491E8 | 7.753E7 | 7.233E7 | 8.178E7 | 8.173 | 7.889 | 7.859 | 7.913 | 8.031 | 7.886 | 0.145 |
| G3V2B8 | Uncharacterized protein OS=Homo sapiens GN=MTHFD1 PE=3 SV=1 - [G3V2B8_HUMAN] | 8.560E7 | 7.583E7 | 5.850E7 | 5.681E7 | 7.932 | 7.880 | 7.767 | 7.754 | 7.906 | 7.761 | 0.145 |
| B4DR70 | Uncharacterized protein OS=Homo sapiens GN=FUS PE=2 SV=1 - [B4DR70_HUMAN] | 5.295E7 | 2.958E7 | 2.934E7 | 2.735E7 | 7.724 | 7.471 | 7.467 | 7.437 | 7.597 | 7.452 | 0.145 |
| Q14257 | Reticulocalbin-2 OS=Homo sapiens GN=RCN2 PE=1 SV=1 - [RCN2_HUMAN] | | 8.962E6 | | 6.417E6 | | 6.952 | | 6.807 | 6.952 | 6.807 | 0.145 |
| P68371 | Tubulin beta-4B chain OS=Homo sapiens GN=TUBB4B PE=1 SV=1 - [TBB4B_HUMAN] | 1.379E9 | 1.104E9 | 8.816E8 | 8.864E8 | 9.139 | 9.043 | 8.945 | 8.948 | 9.091 | 8.946 | 0.145 |
| B7Z1R5 | Uncharacterized protein OS=Homo sapiens GN=ATP6V1A PE=2 SV=1 - [B7Z1R5_HUMAN] | 2.513E7 | 1.718E7 | 1.456E7 | 1.523E7 | 7.400 | 7.235 | 7.163 | 7.183 | 7.318 | 7.173 | 0.145 |
| Q13011 | Delta(3,5)-Delta(2,4)-dienoyl-CoA isomerase, mitochondrial OS=Homo sapiens GN=ECH1 PE=1 SV=2 - [ECH | 1.938E7 | 1.942E7 | 1.367E7 | 1.419E7 | 7.287 | 7.288 | 7.136 | 7.152 | 7.288 | 7.144 | 0.144 |
| Q9H0C8 | Integrin-linked kinase-associated serine/threonine phosphatase 2C OS=Homo sapiens GN=ILKAP PE=1 SV=1 | 1.161E7 | 7.815E6 | | 6.844E6 | 7.065 | 6.893 | | 6.835 | 6.979 | 6.835 | 0.144 |
| F8VPF3 | Uncharacterized protein OS=Homo sapiens GN=MYL6 PE=4 SV=1 - [F8VPF3_HUMAN] | 1.096E8 | 6.513E7 | 5.854E7 | 6.301E7 | 8.040 | 7.814 | 7.767 | 7.799 | 7.927 | 7.783 | 0.143 |
| Q04637-5 | Isoform D of Eukaryotic translation initiation factor 4 gamma 1 OS=Homo sapiens GN=EIF4G1 - [IF4G1_HUM | 4.181E7 | 2.827E7 | 2.668E7 | 2.290E7 | 7.621 | 7.451 | 7.426 | 7.360 | 7.536 | 7.393 | 0.143 |
| Q7Z784 | DNAJC7 protein OS=Homo sapiens GN=DNAJC7 PE=2 SV=1 - [Q7Z784_HUMAN] | 1.027E7 | 7.129E6 | 6.318E6 | 5.993E6 | 7.012 | 6.853 | 6.801 | 6.778 | 6.932 | 6.789 | 0.143 |
| Q15181 | Inorganic pyrophosphatase OS=Homo sapiens GN=PPA1 PE=1 SV=2 - [IPYR_HUMAN] | 1.799E8 | 1.368E8 | 1.117E8 | 1.141E8 | 8.255 | 8.136 | 8.048 | 8.057 | 8.196 | 8.052 | 0.143 |
| Q9NTJ3-2 | Isoform 2 of Structural maintenance of chromosomes protein 4 OS=Homo sapiens GN=SMC4 - [SMC4_HUMA | 2.309E7 | 1.935E7 | 1.560E7 | 1.482E7 | 7.363 | 7.287 | 7.193 | 7.171 | 7.325 | 7.182 | 0.143 |
| Q09028-3 | Isoform 3 of Histone-binding protein RBBP4 OS=Homo sapiens GN=RBBP4 - [RBBP4_HUMAN] | 7.788E7 | 5.278E7 | 4.595E7 | 4.633E7 | 7.891 | 7.722 | 7.662 | 7.666 | 7.807 | 7.664 | 0.143 |
| E9PLK3 | Uncharacterized protein OS=Homo sapiens GN=NPEPPS PE=4 SV=1 - [E9PLK3_HUMAN] | 2.791E7 | 1.822E7 | 1.467E7 | 1.797E7 | 7.446 | 7.261 | 7.166 | 7.255 | 7.353 | 7.210 | 0.143 |
| A6NKB8 | Uncharacterized protein OS=Homo sapiens GN=RNPEP PE=4 SV=1 - [A6NKB8_HUMAN] | 2.517E7 | 1.957E7 | 1.620E7 | 1.578E7 | 7.401 | 7.292 | 7.210 | 7.198 | 7.346 | 7.204 | 0.142 |
| P33316-2 | Isoform 2 of Deoxyuridine 5'-triphosphate nucleotidohydrolase, mitochondrial OS=Homo sapiens GN=DUT - [ | 5.029E7 | 2.585E7 | 2.466E7 | 2.743E7 | 7.701 | 7.412 | 7.392 | 7.438 | 7.557 | 7.415 | 0.142 |
| P30044-2 | Isoform Cytoplasmic+peroxisomal of Peroxiredoxin-5, mitochondrial OS=Homo sapiens GN=PRDX5 - [PRDX5_ | 5.176E7 | 4.808E7 | 3.159E7 | 4.101E7 | 7.714 | 7.682 | 7.500 | 7.613 | 7.698 | 7.556 | 0.142 |
| Q9BU61 | NADH dehydrogenase [ubiquinone] 1 alpha subcomplex assembly factor 3 OS=Homo sapiens GN=NDUFAF3 I | 1.234E7 | 8.474E6 | 6.735E6 | 8.082E6 | 7.091 | 6.928 | 6.828 | 6.908 | 7.010 | 6.868 | 0.142 |
| O75369-6 | Isoform 6 of Filamin-B OS=Homo sapiens GN=FLNB - [FLNB_HUMAN] | 1.619E7 | 1.571E7 | 1.207E7 | 1.097E7 | 7.209 | 7.196 | 7.082 | 7.040 | 7.203 | 7.061 | 0.142 |
| E7EQQ1 | Uncharacterized protein OS=Homo sapiens GN=MSH2 PE=4 SV=1 - [E7EQQ1_HUMAN] | 8.883E6 | 5.092E6 | 4.648E6 | 5.068E6 | 6.949 | 6.707 | 6.667 | 6.705 | 6.828 | 6.686 | 0.142 |
| O95292 | Vesicle-associated membrane protein-associated protein B/C OS=Homo sapiens GN=VAPB PE=1 SV=3 - [VAP | 1.954E7 | 6.618E6 | 8.806E6 | 7.650E6 | 7.291 | 6.821 | 6.945 | 6.884 | 7.056 | 6.914 | 0.142 |
| P22314 | Ubiquitin-like modifier-activating enzyme 1 OS=Homo sapiens GN=UBA1 PE=1 SV=3 - [UBA1_HUMAN] | 1.816E8 | 1.331E8 | 1.003E8 | 1.256E8 | 8.259 | 8.124 | 8.001 | 8.099 | 8.192 | 8.050 | 0.142 |
| P35580 | Myosin-10 OS=Homo sapiens GN=MYH10 PE=1 SV=3 - [MYH10_HUMAN] | 2.693E7 | 1.489E7 | 1.313E7 | 1.596E7 | 7.430 | 7.173 | 7.118 | 7.203 | 7.302 | 7.161 | 0.141 |
| P51659 | Peroxisomal multifunctional enzyme type 2 OS=Homo sapiens GN=HSD17B4 PE=1 SV=3 - [DHB4_HUMAN] | 2.112E7 | 2.199E7 | 1.460E7 | 1.664E7 | 7.325 | 7.342 | 7.164 | 7.221 | 7.334 | 7.193 | 0.141 |
| P52272-2 | Isoform 2 of Heterogeneous nuclear ribonucleoprotein M OS=Homo sapiens GN=HNRNPM - [HNRPM_HUMAN | 1.421E8 | 9.851E7 | 8.029E7 | 9.121E7 | 8.152 | 7.993 | 7.905 | 7.960 | 8.073 | 7.932 | 0.141 |
| O00232 | 26S proteasome non-ATPase regulatory subunit 12 OS=Homo sapiens GN=PSMD12 PE=1 SV=3 - [PSD12_HU | 1.865E7 | | 1.284E7 | 1.418E7 | 7.271 | | 7.109 | 7.152 | 7.271 | 7.130 | 0.140 |
| P00338 | L-lactate dehydrogenase A chain OS=Homo sapiens GN=LDHA PE=1 SV=2 - [LDHA_HUMAN] | 2.060E8 | 1.466E8 | 1.207E8 | 1.312E8 | 8.314 | 8.166 | 8.082 | 8.118 | 8.240 | 8.100 | 0.140 |
| O14737 | Programmed cell death protein 5 OS=Homo sapiens GN=PDCD5 PE=1 SV=3 - [PDCD5_HUMAN] | 6.128E7 | 5.319E7 | 3.411E7 | 5.008E7 | 7.787 | 7.726 | 7.533 | 7.700 | 7.757 | 7.616 | 0.140 |
| Q9H2U2-3 | Isoform 3 of Inorganic pyrophosphatase 2, mitochondrial OS=Homo sapiens GN=PPA2 - [IPYR2_HUMAN] | 3.743E7 | 3.418E7 | 2.214E7 | 3.030E7 | 7.573 | 7.534 | 7.345 | 7.482 | 7.554 | 7.413 | 0.140 |
| P61247 | 40S ribosomal protein S3a OS=Homo sapiens GN=RPS3A PE=1 SV=2 - [RS3A_HUMAN] | 1.050E8 | 7.854E7 | 6.872E7 | 6.290E7 | 8.021 | 7.895 | 7.837 | 7.799 | 7.958 | 7.818 | 0.140 |
| P58546 | Myotrophin OS=Homo sapiens GN=MTPN PE=1 SV=2 - [MTPN_HUMAN] | 6.544E7 | 4.857E7 | 3.906E7 | 4.270E7 | 7.816 | 7.686 | 7.592 | 7.630 | 7.751 | 7.611 | 0.140 |
| P56192 | Methionyl-tRNA synthetase, cytoplasmic OS=Homo sapiens GN=MARS PE=1 SV=2 - [SYMC_HUMAN] | 3.546E7 | 3.364E7 | 2.491E7 | 2.519E7 | 7.550 | 7.527 | 7.396 | 7.401 | 7.538 | 7.399 | 0.140 |
| P60228 | Eukaryotic translation initiation factor 3 subunit E OS=Homo sapiens GN=EIF3E PE=1 SV=1 - [EIF3E_HUMAN | 2.204E7 | 1.606E7 | 1.102E7 | 1.691E7 | 7.343 | 7.206 | 7.042 | 7.228 | 7.275 | 7.135 | 0.139 |
| P48047 | ATP synthase subunit O, mitochondrial OS=Homo sapiens GN=ATP5O PE=1 SV=1 - [ATPO_HUMAN] | 4.109E7 | 2.712E7 | 2.683E7 | 2.189E7 | 7.614 | 7.433 | 7.429 | 7.340 | 7.524 | 7.384 | 0.139 |
| Q9BUB1 | PRKAR2A protein OS=Homo sapiens GN=PRKAR2A PE=2 SV=1 - [Q9BUB1_HUMAN] | 1.573E7 | 1.376E7 | 1.068E7 | 1.070E7 | 7.197 | 7.139 | 7.029 | 7.029 | 7.168 | 7.029 | 0.139 |
| Q7KZF4 | Staphylococcal nuclease domain-containing protein 1 OS=Homo sapiens GN=SND1 PE=1 SV=1 - [SND1_HUM | 5.951E7 | 3.858E7 | 3.397E7 | 3.573E7 | 7.775 | 7.586 | 7.531 | 7.553 | 7.681 | 7.542 | 0.138 |
| F8W7D6 | Uncharacterized protein OS=Homo sapiens GN=GPHN PE=4 SV=1 - [F8W7D6_HUMAN] | 8.724E6 | 6.524E6 | 4.298E6 | 7.017E6 | 6.941 | 6.815 | 6.633 | 6.846 | 6.878 | 6.740 | 0.138 |
| E9PID8 | Uncharacterized protein OS=Homo sapiens GN=CSTF2 PE=4 SV=1 - [E9PID8_HUMAN] | 1.982E7 | 1.745E7 | 1.057E7 | 1.735E7 | 7.297 | 7.242 | 7.024 | 7.239 | 7.269 | 7.132 | 0.138 |
| P62714 | Serine/threonine-protein phosphatase 2A catalytic subunit beta isoform OS=Homo sapiens GN=PPP2CB PE=1 | 2.189E7 | 7.664E6 | 1.024E7 | 8.690E6 | 7.340 | 6.884 | 7.010 | 6.939 | 7.112 | 6.975 | 0.138 |
| F5H0F5 | Uncharacterized protein OS=Homo sapiens GN=ZC3H15 PE=4 SV=1 - [F5H0F5_HUMAN] | 1.090E7 | 9.513E6 | 7.901E6 | 6.966E6 | 7.037 | 6.978 | 6.898 | 6.843 | 7.008 | 6.870 | 0.138 |
| P49721 | Proteasome subunit beta type-2 OS=Homo sapiens GN=PSMB2 PE=1 SV=1 - [PSB2_HUMAN] | 1.610E7 | 1.426E7 | 1.109E7 | 1.100E7 | 7.207 | 7.154 | 7.045 | 7.041 | 7.180 | 7.043 | 0.137 |
| Q9UBQ7 | Glyoxylate reductase/hydroxypyruvate reductase OS=Homo sapiens GN=GRHPR PE=1 SV=1 - [GRHPR_HUM/ | 1.803E7 | 1.944E7 | 1.478E7 | 1.260E7 | 7.256 | 7.289 | 7.170 | 7.100 | 7.272 | 7.135 | 0.137 |
| P42771 | Cyclin-dependent kinase inhibitor 2A, isoforms 1/2/3 OS=Homo sapiens GN=CDKN2A PE=1 SV=2 - [CD2A1_I | 2.472E7 | 1.632E7 | 1.414E7 | 1.517E7 | 7.393 | 7.213 | 7.150 | 7.181 | 7.303 | 7.166 | 0.137 |
| B4DDF7 | Uncharacterized protein OS=Homo sapiens GN=PPP2R1A PE=2 SV=1 - [B4DDF7_HUMAN] | 3.318E7 | 2.834E7 | 2.118E7 | 2.362E7 | 7.521 | 7.452 | 7.326 | 7.373 | 7.487 | 7.350 | 0.137 |
| P54886-2 | Isoform Short of Delta-1-pyrroline-5-carboxylate synthase OS=Homo sapiens GN=ALDH18A1 - [P5CS_HUMAN | 4.189E7 | 3.767E7 | 2.543E7 | 3.302E7 | 7.622 | 7.576 | 7.405 | 7.519 | 7.599 | 7.462 | 0.137 |
| P13667 | Protein disulfide-isomerase A4 OS=Homo sapiens GN=PDIA4 PE=1 SV=2 - [PDIA4_HUMAN] | 7.064E7 | 5.019E7 | 4.194E7 | 4.500E7 | 7.849 | 7.701 | 7.623 | 7.653 | 7.775 | 7.638 | 0.137 |
| Q13185 | Chromobox protein homolog 3 OS=Homo sapiens GN=CBX3 PE=1 SV=4 - [CBX3_HUMAN] | 1.641E7 | 1.522E7 | 1.241E7 | 1.072E7 | 7.215 | 7.182 | 7.094 | 7.030 | 7.199 | 7.062 | 0.137 |
| Q15393 | Splicing factor 3B subunit 3 OS=Homo sapiens GN=SF3B3 PE=1 SV=4 - [SF3B3_HUMAN] | 3.348E7 | 2.270E7 | 1.996E7 | 2.027E7 | 7.525 | 7.356 | 7.300 | 7.307 | 7.440 | 7.304 | 0.137 |
| P06744 | Glucose-6-phosphate isomerase OS=Homo sapiens GN=GPI PE=1 SV=4 - [G6PI_HUMAN] | 1.505E8 | 1.258E8 | 9.782E7 | 1.035E8 | 8.178 | 8.100 | 7.990 | 8.015 | 8.139 | 8.003 | 0.136 |
| B7Z4V2 | Uncharacterized protein OS=Homo sapiens GN=HSPA9 PE=2 SV=1 - [B7Z4V2_HUMAN] | 2.231E8 | 1.751E8 | 1.408E8 | 1.486E8 | 8.348 | 8.243 | 8.149 | 8.172 | 8.296 | 8.160 | 0.135 |
| P52907 | F-actin-capping protein subunit alpha-1 OS=Homo sapiens GN=CAPZA1 PE=1 SV=3 - [CAZA1_HUMAN] | 5.742E7 | 3.834E7 | 3.388E7 | 3.484E7 | 7.759 | 7.584 | 7.530 | 7.542 | 7.671 | 7.536 | 0.135 |
| Q53GQ0 | Estradiol 17-beta-dehydrogenase 12 OS=Homo sapiens GN=HSD17B12 PE=1 SV=2 - [DHB12_HUMAN] | 1.280E7 | 1.042E7 | 8.015E6 | 8.940E6 | 7.107 | 7.018 | 6.904 | 6.951 | 7.062 | 6.928 | 0.135 |
| Q9Y617 | Phosphoserine aminotransferase OS=Homo sapiens GN=PSAT1 PE=1 SV=2 - [SERC_HUMAN] | 7.519E7 | 5.315E7 | 4.528E7 | 4.755E7 | 7.876 | 7.726 | 7.656 | 7.677 | 7.801 | 7.667 | 0.134 |
| Q08945 | FACT complex subunit SSRP1 OS=Homo sapiens GN=SSRP1 PE=1 SV=1 - [SSRP1_HUMAN] | 2.725E7 | 2.450E7 | 1.849E7 | 1.945E7 | 7.435 | 7.389 | 7.267 | 7.289 | 7.412 | 7.278 | 0.134 |
| Q15691 | Microtubule-associated protein RP/EB family member 1 OS=Homo sapiens GN=MAPRE1 PE=1 SV=3 - [MARE | 5.294E7 | 3.154E7 | 2.916E7 | 3.091E7 | 7.724 | 7.499 | 7.465 | 7.490 | 7.611 | 7.477 | 0.134 |
| Q13126 | S-methyl-5'-thioadenosine phosphorylase OS=Homo sapiens GN=MTAP PE=1 SV=2 - [MTAP_HUMAN] | 2.213E7 | 2.233E7 | 2.121E7 | 1.258E7 | 7.345 | 7.349 | 7.327 | 7.100 | 7.347 | 7.213 | 0.134 |
| P18206-2 | Isoform 1 of Vinculin OS=Homo sapiens GN=VCL - [VINC_HUMAN] | 3.117E7 | 2.253E7 | 1.852E7 | 2.048E7 | 7.494 | 7.353 | 7.268 | 7.311 | 7.423 | 7.290 | 0.134 |
| P12268 | Inosine-5'-monophosphate dehydrogenase 2 OS=Homo sapiens GN=IMPDH2 PE=1 SV=2 - [IMDH2_HUMAN] | 1.121E8 | 7.005E7 | 6.903E7 | 6.147E7 | 8.050 | 7.845 | 7.839 | 7.789 | 7.948 | 7.814 | 0.134 |
| E9PKD1 | Uncharacterized protein OS=Homo sapiens GN=TOR1AIP1 PE=4 SV=1 - [E9PKD1_HUMAN] | 1.452E7 | 9.192E6 | 8.498E6 | | 7.162 | 6.963 | 6.929 | | 7.063 | 6.929 | 0.133 |
| Q9Y2W1 | Thyroid hormone receptor-associated protein 3 OS=Homo sapiens GN=THRAP3 PE=1 SV=2 - [TR150_HUMAI | 1.040E7 | 3.379E6 | 5.005E6 | 3.803E6 | 7.017 | 6.529 | 6.699 | 6.580 | 6.773 | 6.640 | 0.133 |
| B4DJV2 | Citrate synthase OS=Homo sapiens GN=CS PE=2 SV=1 - [B4DJV2_HUMAN] | 6.631E7 | 6.385E7 | 4.493E7 | 5.107E7 | 7.822 | 7.805 | 7.653 | 7.708 | 7.813 | 7.680 | 0.133 |
| Q13838 | Spliceosome RNA helicase DDX39B OS=Homo sapiens GN=DDX39B PE=1 SV=1 - [DX39B_HUMAN] | 1.571E8 | 1.091E8 | 9.440E7 | 9.843E7 | 8.196 | 8.038 | 7.975 | 7.993 | 8.117 | 7.984 | 0.133 |
| O60664 | Perilipin-3 OS=Homo sapiens GN=PLIN3 PE=1 SV=3 - [PLIN3_HUMAN] | 1.275E7 | 8.083E6 | 7.087E6 | 7.890E6 | 7.106 | 6.908 | 6.850 | 6.897 | 7.007 | 6.874 | 0.133 |
| P23381 | Tryptophanyl-tRNA synthetase, cytoplasmic OS=Homo sapiens GN=WARS PE=1 SV=2 - [SYWC_HUMAN] | 4.285E7 | 3.139E7 | 3.027E7 | 2.412E7 | 7.632 | 7.497 | 7.481 | 7.382 | 7.564 | 7.432 | 0.133 |
| P53621 | Coatomer subunit alpha OS=Homo sapiens GN=COPA PE=1 SV=2 - [COPA_HUMAN] | 1.996E7 | 1.874E7 | 1.356E7 | 1.502E7 | 7.300 | 7.273 | 7.132 | 7.177 | 7.287 | 7.154 | 0.132 |
| Q6IBS0 | Twinfilin-2 OS=Homo sapiens GN=TWF2 PE=1 SV=2 - [TWF2_HUMAN] | 1.766E7 | 1.569E7 | 1.207E7 | 1.251E7 | 7.247 | 7.196 | 7.082 | 7.097 | 7.221 | 7.089 | 0.132 |
| P32322 | Pyrroline-5-carboxylate reductase 1, mitochondrial OS=Homo sapiens GN=PYCR1 PE=1 SV=2 - [P5CR1_HUM | 1.648E7 | 1.650E7 | 9.979E6 | 1.485E7 | 7.217 | 7.218 | 6.999 | 7.172 | 7.217 | 7.085 | 0.132 |
| F5H4I0 | Dihydrolipoyl dehydrogenase OS=Homo sapiens GN=DLD PE=3 SV=1 - [F5H4I0_HUMAN] | 3.514E7 | 2.772E7 | 2.077E7 | 2.556E7 | 7.546 | 7.443 | 7.317 | 7.408 | 7.494 | 7.363 | 0.132 |
| P28070 | Proteasome subunit beta type-4 OS=Homo sapiens GN=PSMB4 PE=1 SV=4 - [PSB4_HUMAN] | 3.787E7 | 3.125E7 | 2.411E7 | 2.681E7 | 7.578 | 7.495 | 7.382 | 7.428 | 7.537 | 7.405 | 0.131 |
| P62913 | 60S ribosomal protein L11 OS=Homo sapiens GN=RPL11 PE=1 SV=2 - [RL11_HUMAN] | 1.676E8 | 1.090E8 | 9.632E7 | 1.037E8 | 8.224 | 8.038 | 7.984 | 8.016 | 8.131 | 8.000 | 0.131 |
| Q15843 | NEDD8 OS=Homo sapiens GN=NEDD8 PE=1 SV=1 - [NEDD8_HUMAN] | 8.847E7 | 7.329E7 | 5.956E7 | 5.952E7 | 7.947 | 7.865 | 7.775 | 7.775 | 7.906 | 7.775 | 0.131 |
| P61289 | Proteasome activator complex subunit 3 OS=Homo sapiens GN=PSME3 PE=1 SV=1 - [PSME3_HUMAN] | 2.520E7 | 1.684E7 | 1.453E7 | 1.598E7 | 7.401 | 7.226 | 7.162 | 7.204 | 7.314 | 7.183 | 0.131 |
| Q9BVA1 | Tubulin beta-2B chain OS=Homo sapiens GN=TUBB2B PE=1 SV=1 - [TBB2B_HUMAN] | 1.440E9 | 1.214E9 | 9.844E8 | 9.736E8 | 9.158 | 9.084 | 8.993 | 8.988 | 9.121 | 8.991 | 0.131 |
| F8VW92 | Uncharacterized protein OS=Homo sapiens GN=TUBB PE=3 SV=1 - [F8VW92_HUMAN] | 1.440E9 | 1.214E9 | 9.844E8 | 9.736E8 | 9.158 | 9.084 | 8.993 | 8.988 | 9.121 | 8.991 | 0.131 |
| Q7Z6Z7-2 | Isoform 2 of E3 ubiquitin-protein ligase HUWE1 OS=Homo sapiens GN=HUWE1 - [HUWE1_HUMAN] | 1.295E7 | 1.232E7 | 9.018E6 | 9.705E6 | 7.112 | 7.090 | 6.955 | 6.987 | 7.101 | 6.971 | 0.130 |
| P00918 | Carbonic anhydrase 2 OS=Homo sapiens GN=CA2 PE=1 SV=2 - [CAH2_HUMAN] | 1.053E8 | 7.998E7 | 6.558E7 | 7.049E7 | 8.022 | 7.903 | 7.817 | 7.848 | 7.963 | 7.832 | 0.130 |
| P84098 | 60S ribosomal protein L19 OS=Homo sapiens GN=RPL19 PE=1 SV=1 - [RL19_HUMAN] | 5.791E7 | 4.210E7 | 3.517E7 | 3.813E7 | 7.763 | 7.624 | 7.546 | 7.581 | 7.694 | 7.564 | 0.130 |
| P20700 | Lamin-B1 OS=Homo sapiens GN=LMNB1 PE=1 SV=2 - [LMNB1_HUMAN] | 1.324E7 | 1.305E7 | 1.066E7 | 8.916E6 | 7.122 | 7.116 | 7.028 | 6.950 | 7.119 | 6.989 | 0.130 |
| P62304 | Small nuclear ribonucleoprotein E OS=Homo sapiens GN=SNRPE PE=1 SV=1 - [RUXE_HUMAN] | 9.905E7 | 8.714E7 | 6.779E7 | 7.007E7 | 7.996 | 7.940 | 7.831 | 7.846 | 7.968 | 7.838 | 0.130 |
| F5H5Y2 | Uncharacterized protein OS=Homo sapiens GN=SSR1 PE=4 SV=1 - [F5H5Y2_HUMAN] | 1.047E7 | 8.402E6 | 7.037E6 | 6.886E6 | 7.020 | 6.924 | 6.847 | 6.838 | 6.972 | 6.843 | 0.130 |
| Q16543 | Hsp90 co-chaperone Cdc37 OS=Homo sapiens GN=CDC37 PE=1 SV=1 - [CDC37_HUMAN] | 4.358E7 | 3.416E7 | 2.739E7 | 2.994E7 | 7.639 | 7.534 | 7.438 | 7.476 | 7.586 | 7.457 | 0.129 |
| B1AK87 | Capping protein (Actin filament) muscle Z-line, beta OS=Homo sapiens GN=CAPZB PE=4 SV=1 - [B1AK87_HU | 2.273E7 | 1.267E7 | 1.207E7 | 1.315E7 | 7.357 | 7.103 | 7.082 | 7.119 | 7.230 | 7.100 | 0.129 |
| P20290-2 | Isoform 2 of Transcription factor BTF3 OS=Homo sapiens GN=BTF3 - [BTF3_HUMAN] | 1.021E8 | 6.948E7 | 5.817E7 | 6.724E7 | 8.009 | 7.842 | 7.765 | 7.828 | 7.926 | 7.796 | 0.129 |
| E9PKG1 | Uncharacterized protein OS=Homo sapiens GN=PRMT1 PE=4 SV=1 - [E9PKG1_HUMAN] | 1.900E7 | 1.078E7 | 1.076E7 | 1.050E7 | 7.279 | 7.032 | 7.032 | 7.021 | 7.156 | 7.027 | 0.129 |
| P40227 | T-complex protein 1 subunit zeta OS=Homo sapiens GN=CCT6A PE=1 SV=3 - [TCPZ_HUMAN] | 1.649E8 | 1.208E8 | 1.078E8 | 1.021E8 | 8.217 | 8.082 | 8.033 | 8.009 | 8.150 | 8.021 | 0.129 |
| P11940-2 | Isoform 2 of Polyadenylate-binding protein 1 OS=Homo sapiens GN=PABPC1 - [PABP1_HUMAN] | 9.749E7 | 7.216E7 | 5.901E7 | 6.596E7 | 7.989 | 7.858 | 7.771 | 7.819 | 7.924 | 7.795 | 0.129 |
| P25787 | Proteasome subunit alpha type-2 OS=Homo sapiens GN=PSMA2 PE=1 SV=2 - [PSA2_HUMAN] | 6.444E7 | 5.155E7 | 4.239E7 | 4.339E7 | 7.809 | 7.712 | 7.627 | 7.637 | 7.761 | 7.632 | 0.128 |
| E9PMI6 | Uncharacterized protein OS=Homo sapiens GN=CLNS1A PE=4 SV=1 - [E9PMI6_HUMAN] | 6.013E7 | 5.340E7 | 4.004E7 | 4.442E7 | 7.779 | 7.728 | 7.602 | 7.648 | 7.753 | 7.625 | 0.128 |
| P17980 | 26S protease regulatory subunit 6A OS=Homo sapiens GN=PSMC3 PE=1 SV=3 - [PRS6A_HUMAN] | 3.289E7 | 2.574E7 | 2.244E7 | 2.091E7 | 7.517 | 7.411 | 7.351 | 7.320 | 7.464 | 7.336 | 0.128 |
| B3KQT9 | Uncharacterized protein OS=Homo sapiens GN=PDIA3 PE=2 SV=1 - [B3KQT9_HUMAN] | 1.145E8 | 9.029E7 | 7.474E7 | 7.696E7 | 8.059 | 7.956 | 7.874 | 7.886 | 8.007 | 7.880 | 0.127 |
| Q96CX2 | BTB/POZ domain-containing protein KCTD12 OS=Homo sapiens GN=KCTD12 PE=1 SV=1 - [KCD12_HUMAN] | 3.084E7 | 1.982E7 | 1.677E7 | 2.031E7 | 7.489 | 7.297 | 7.224 | 7.308 | 7.393 | 7.266 | 0.127 |
| P14625 | Endoplasmin OS=Homo sapiens GN=HSP90B1 PE=1 SV=1 - [ENPL_HUMAN] | 1.289E8 | 8.073E7 | 8.322E7 | 6.973E7 | 8.110 | 7.907 | 7.920 | 7.843 | 8.009 | 7.882 | 0.127 |
| Q13596-2 | Isoform 1A of Sorting nexin-1 OS=Homo sapiens GN=SNX1 - [SNX1_HUMAN] | 1.218E7 | 1.184E7 | | 8.971E6 | 7.086 | 7.073 | | 6.953 | 7.080 | 6.953 | 0.127 |
| Q96C90 | Protein phosphatase 1 regulatory subunit 14B OS=Homo sapiens GN=PPP1R14B PE=1 SV=3 - [PP14B_HUMAN] | | 2.463E7 | 1.366E7 | 2.479E7 | | 7.392 | 7.135 | 7.394 | 7.392 | 7.265 | 0.127 |
| Q96EP5-2 | Isoform 2 of DAZ-associated protein 1 OS=Homo sapiens GN=DAZAP1 - [DAZP1_HUMAN] | 1.596E7 | 1.739E7 | 1.355E7 | 1.144E7 | 7.203 | 7.240 | 7.132 | 7.058 | 7.222 | 7.095 | 0.127 |
| P04406 | Glyceraldehyde-3-phosphate dehydrogenase OS=Homo sapiens GN=GAPDH PE=1 SV=3 - [G3P_HUMAN] | 1.334E9 | 1.057E9 | 8.559E8 | 9.203E8 | 9.125 | 9.024 | 8.932 | 8.964 | 9.075 | 8.948 | 0.126 |
| P12236 | ADP/ATP translocase 3 OS=Homo sapiens GN=SLC25A6 PE=1 SV=4 - [ADT3_HUMAN] | 5.522E7 | 3.784E7 | 3.227E7 | 3.619E7 | 7.742 | 7.578 | 7.509 | 7.559 | 7.660 | 7.534 | 0.126 |
| Q5JR95 | Ribosomal protein S8 OS=Homo sapiens GN=RPS8 PE=4 SV=1 - [Q5JR95_HUMAN] | 1.889E8 | 1.294E8 | 1.097E8 | 1.249E8 | 8.276 | 8.112 | 8.040 | 8.097 | 8.194 | 8.068 | 0.126 |
| P31948 | Stress-induced-phosphoprotein 1 OS=Homo sapiens GN=STIP1 PE=1 SV=1 - [STIP1_HUMAN] | 1.121E8 | 1.095E8 | 6.915E7 | 9.954E7 | 8.049 | 8.040 | 7.840 | 7.998 | 8.045 | 7.919 | 0.126 |
| P04075 | Fructose-bisphosphate aldolase A OS=Homo sapiens GN=ALDOA PE=1 SV=2 - [ALDOA_HUMAN] | 3.770E8 | 2.875E8 | 2.440E8 | 2.491E8 | 8.576 | 8.459 | 8.387 | 8.396 | 8.517 | 8.392 | 0.126 |
| P28838-2 | Isoform 2 of Cytosol aminopeptidase OS=Homo sapiens GN=LAP3 - [AMPL_HUMAN] | 1.431E7 | 1.387E7 | 1.044E7 | 1.066E7 | 7.155 | 7.142 | 7.019 | 7.028 | 7.149 | 7.023 | 0.125 |
| P48643 | T-complex protein 1 subunit epsilon OS=Homo sapiens GN=CCT5 PE=1 SV=1 - [TCPE_HUMAN] | 1.092E8 | 8.672E7 | 7.130E7 | 7.469E7 | 8.038 | 7.938 | 7.853 | 7.873 | 7.988 | 7.863 | 0.125 |
| E7ETA0 | Uncharacterized protein OS=Homo sapiens GN=YBX1 PE=4 SV=1 - [E7ETA0_HUMAN] | 1.880E8 | 9.751E7 | 1.016E8 | 1.015E8 | 8.274 | 7.989 | 8.007 | 8.007 | 8.132 | 8.007 | 0.125 |
| E9PH29 | Uncharacterized protein OS=Homo sapiens GN=PRDX3 PE=4 SV=1 - [E9PH29_HUMAN] | 1.679E8 | 1.253E8 | 1.086E8 | 1.092E8 | 8.225 | 8.098 | 8.036 | 8.038 | 8.162 | 8.037 | 0.124 |
| E9PN51 | Uncharacterized protein OS=Homo sapiens GN=NDUFS8 PE=4 SV=1 - [E9PN51_HUMAN] | | 7.823E6 | | 5.877E6 | | 6.893 | | 6.769 | 6.893 | 6.769 | 0.124 |
| Q9BUJ2-2 | Isoform 2 of Heterogeneous nuclear ribonucleoprotein U-like protein 1 OS=Homo sapiens GN=HNRNPUL1 - [I | 1.667E7 | 1.174E7 | 1.185E7 | 9.334E6 | 7.222 | 7.070 | 7.074 | 6.970 | 7.146 | 7.022 | 0.124 |
| Q9UKK9 | ADP-sugar pyrophosphatase OS=Homo sapiens GN=NUDT5 PE=1 SV=1 - [NUDT5_HUMAN] | 4.218E7 | 6.205E7 | 1.744E7 | 2.784E7 | 7.625 | 7.793 | 7.242 | 7.445 | 7.709 | 7.343 | 0.366 |
| P46778 | 60S ribosomal protein L21 OS=Homo sapiens GN=RPL21 PE=1 SV=2 - [RL21_HUMAN] | 1.131E8 | 8.838E7 | 7.289E7 | 7.800E7 | 8.054 | 7.946 | 7.863 | 7.892 | 8.000 | 7.877 | 0.123 |
| P63244 | Guanine nucleotide-binding protein subunit beta-2-like 1 OS=Homo sapiens GN=GNB2L1 PE=1 SV=3 - [GBLP | 7.920E7 | 6.068E7 | 5.365E7 | 5.099E7 | 7.899 | 7.783 | 7.730 | 7.707 | 7.841 | 7.719 | 0.122 |
| P12532 | Creatine kinase U-type, mitochondrial OS=Homo sapiens GN=CKMT1A PE=1 SV=1 - [KCRU_HUMAN] | 4.476E7 | 4.743E7 | 3.080E7 | 3.925E7 | 7.651 | 7.676 | 7.489 | 7.594 | 7.663 | 7.541 | 0.122 |
| O15067 | Phosphoribosylformylglycinamidine synthase OS=Homo sapiens GN=PFAS PE=1 SV=4 - [PUR4_HUMAN] | 3.205E7 | 2.464E7 | 2.092E7 | 2.150E7 | 7.506 | 7.392 | 7.321 | 7.332 | 7.449 | 7.326 | 0.122 |
| G3V580 | Uncharacterized protein OS=Homo sapiens GN=PRMT5 PE=4 SV=1 - [G3V580_HUMAN] | 7.422E6 | | 6.816E6 | 4.620E6 | 6.871 | | 6.834 | 6.665 | 6.871 | 6.749 | 0.121 |
| B3KUK2 | Superoxide dismutase OS=Homo sapiens GN=SOD2 PE=2 SV=1 - [B3KUK2_HUMAN] | 2.218E7 | 2.601E7 | 1.575E7 | 2.096E7 | 7.346 | 7.415 | 7.197 | 7.321 | 7.381 | 7.259 | 0.121 |
| Q52LJ0 | Protein FAM98B OS=Homo sapiens GN=FAM98B PE=1 SV=1 - [FA98B_HUMAN] | 1.169E7 | 8.152E6 | 7.395E6 | | 7.068 | 6.911 | 6.869 | | 6.990 | 6.869 | 0.121 |
| P61604 | 10 kDa heat shock protein, mitochondrial OS=Homo sapiens GN=HSPE1 PE=1 SV=2 - [CH10_HUMAN] | 3.478E8 | 2.240E8 | 2.033E8 | 2.204E8 | 8.541 | 8.350 | 8.308 | 8.343 | 8.446 | 8.326 | 0.120 |

| | | | | | | | | | | | | |
|---|---|---|---|---|---|---|---|---|---|---|---|---|
| Q86X55-2 | Isoform 2 of Histone-arginine methyltransferase CARM1 OS=Homo sapiens GN=CARM1 - [CARM1_HUMAN] | 1.095E7 | 1.638E7 | 8.722E6 | 1.183E7 | 7.039 | 7.214 | 6.941 | 7.073 | 7.127 | 7.007 | 0.120 |
| F2Z3D0 | Uncharacterized protein OS=Homo sapiens GN=SUMO2 PE=4 SV=1 - [F2Z3D0_HUMAN] | 1.899E8 | 1.355E8 | 9.483E7 | 1.560E8 | 8.279 | 8.132 | 7.977 | 8.193 | 8.205 | 8.085 | 0.120 |
| F2Z3H3 | Uncharacterized protein OS=Homo sapiens GN=ANP32A PE=4 SV=2 - [F2Z3H3_HUMAN] | 9.028E7 | 5.831E7 | 5.345E7 | 5.666E7 | 7.956 | 7.766 | 7.728 | 7.753 | 7.861 | 7.741 | 0.120 |
| C9JW90 | Uncharacterized protein OS=Homo sapiens GN=PITPNB PE=4 SV=1 - [C9JW90_HUMAN] | 2.020E7 | 1.491E7 | 1.198E7 | 1.452E7 | 7.305 | 7.174 | 7.078 | 7.162 | 7.239 | 7.120 | 0.119 |
| B5MC59 | Replication protein A3, 14kDa, isoform CRA_a OS=Homo sapiens GN=RPA3 PE=4 SV=1 - [B5MC59_HUMAN] | 9.629E6 | 2.559E7 | 9.399E6 | 1.515E7 | 6.984 | 7.408 | 6.973 | 7.180 | 7.196 | 7.077 | 0.119 |
| P41250 | Glycyl-tRNA synthetase OS=Homo sapiens GN=GARS PE=1 SV=3 - [SYG_HUMAN] | 4.511E7 | 4.825E7 | 3.570E7 | 3.531E7 | 7.654 | 7.683 | 7.553 | 7.548 | 7.669 | 7.550 | 0.119 |
| P25205 | DNA replication licensing factor MCM3 OS=Homo sapiens GN=MCM3 PE=1 SV=3 - [MCM3_HUMAN] | 3.938E7 | 3.016E7 | 2.581E7 | 2.668E7 | 7.595 | 7.479 | 7.412 | 7.426 | 7.537 | 7.419 | 0.118 |
| Q6P2Q9 | Pre-mRNA-processing-splicing factor 8 OS=Homo sapiens GN=PRPF8 PE=1 SV=2 - [PRP8_HUMAN] | 1.526E7 | 1.573E7 | 9.058E6 | 1.543E7 | 7.184 | 7.197 | 6.957 | 7.188 | 7.190 | 7.073 | 0.117 |
| E7EUR9 | Uncharacterized protein OS=Homo sapiens GN=CALD1 PE=4 SV=1 - [E7EUR9_HUMAN] | 1.842E7 | 1.314E7 | | 1.187E7 | 7.265 | 7.119 | | 7.075 | 7.192 | 7.075 | 0.117 |
| Q14974 | Importin subunit beta-1 OS=Homo sapiens GN=KPNB1 PE=1 SV=2 - [IMB1_HUMAN] | 7.735E7 | 6.494E7 | 5.337E7 | 5.486E7 | 7.888 | 7.812 | 7.727 | 7.739 | 7.850 | 7.733 | 0.117 |
| Q9Y3F4 | Serine-threonine kinase receptor-associated protein OS=Homo sapiens GN=STRAP PE=1 SV=1 - [STRAP_HUI | 3.278E7 | 2.727E7 | 2.160E7 | 2.416E7 | 7.516 | 7.436 | 7.334 | 7.383 | 7.476 | 7.359 | 0.117 |
| Q92522 | Histone H1x OS=Homo sapiens GN=H1FX PE=1 SV=1 - [H1X_HUMAN] | 4.095E7 | 2.930E7 | 2.731E7 | 2.566E7 | 7.612 | 7.467 | 7.436 | 7.409 | 7.540 | 7.423 | 0.117 |
| Q13405 | 39S ribosomal protein L49, mitochondrial OS=Homo sapiens GN=MRPL49 PE=1 SV=1 - [RM49_HUMAN] | 1.435E7 | 1.167E7 | 9.092E6 | 1.076E7 | 7.157 | 7.067 | 6.959 | 7.032 | 7.112 | 6.995 | 0.117 |
| Q06203 | Amidophosphoribosyltransferase OS=Homo sapiens GN=PPAT PE=1 SV=1 - [PUR1_HUMAN] | 1.873E7 | 1.344E7 | 1.233E7 | 1.192E7 | 7.273 | 7.128 | 7.091 | 7.076 | 7.200 | 7.084 | 0.117 |
| P16949 | Stathmin OS=Homo sapiens GN=STMN1 PE=1 SV=3 - [STMN1_HUMAN] | 1.210E8 | 9.536E7 | 8.195E7 | 8.243E7 | 8.083 | 7.979 | 7.914 | 7.916 | 8.031 | 7.915 | 0.116 |
| P32119 | Peroxiredoxin-2 OS=Homo sapiens GN=PRDX2 PE=1 SV=5 - [PRDX2_HUMAN] | 2.651E8 | 1.887E8 | 1.676E8 | 1.748E8 | 8.423 | 8.276 | 8.224 | 8.242 | 8.350 | 8.233 | 0.116 |
| Q13347 | Eukaryotic translation initiation factor 3 subunit I OS=Homo sapiens GN=EIF3I PE=1 SV=1 - [EIF3I_HUMAN] | 3.652E7 | 2.566E7 | 2.155E7 | 2.551E7 | 7.563 | 7.409 | 7.333 | 7.407 | 7.486 | 7.370 | 0.116 |
| P12814-2 | Isoform 2 of Alpha-actinin-1 OS=Homo sapiens GN=ACTN1 - [ACTN1_HUMAN] | 3.899E7 | 2.276E7 | 2.229E7 | 2.345E7 | 7.591 | 7.357 | 7.348 | 7.370 | 7.474 | 7.359 | 0.115 |
| P61086-2 | Isoform 2 of Ubiquitin-conjugating enzyme E2 K OS=Homo sapiens GN=UBE2K - [UBE2K_HUMAN] | 9.197E6 | 7.835E6 | 6.977E6 | 6.084E6 | 6.964 | 6.894 | 6.844 | 6.784 | 6.929 | 6.814 | 0.115 |
| P49720 | Proteasome subunit beta type-3 OS=Homo sapiens GN=PSMB3 PE=1 SV=2 - [PSB3_HUMAN] | 2.917E7 | 1.907E7 | 1.422E7 | 2.306E7 | 7.465 | 7.280 | 7.153 | 7.363 | 7.373 | 7.258 | 0.115 |
| P26583 | High mobility group protein B2 OS=Homo sapiens GN=HMGB2 PE=1 SV=2 - [HMGB2_HUMAN] | 8.842E7 | 5.848E7 | 6.203E7 | 4.921E7 | 7.947 | 7.767 | 7.793 | 7.692 | 7.857 | 7.742 | 0.114 |
| Q8WXC6 | Myeloma-overexpressed gene 2 protein OS=Homo sapiens GN=MYEOV2 PE=2 SV=3 - [MYOV2_HUMAN] | 1.688E7 | 1.244E7 | 1.065E7 | 1.165E7 | 7.227 | 7.095 | 7.027 | 7.066 | 7.161 | 7.047 | 0.114 |
| P62081 | 40S ribosomal protein S7 OS=Homo sapiens GN=RPS7 PE=1 SV=1 - [RS7_HUMAN] | 1.836E8 | 1.627E8 | 1.239E8 | 1.426E8 | 8.264 | 8.211 | 8.093 | 8.154 | 8.238 | 8.124 | 0.114 |
| Q9UL46 | Proteasome activator complex subunit 2 OS=Homo sapiens GN=PSME2 PE=1 SV=4 - [PSME2_HUMAN] | 1.309E7 | 8.135E6 | 7.485E6 | 8.409E6 | 7.117 | 6.910 | 6.874 | 6.925 | 7.014 | 6.899 | 0.114 |
| P17812 | CTP synthase 1 OS=Homo sapiens GN=CTPS PE=1 SV=2 - [PYRG1_HUMAN] | 2.944E7 | 1.418E7 | 1.558E7 | 1.587E7 | 7.469 | 7.152 | 7.193 | 7.201 | 7.310 | 7.197 | 0.114 |
| Q9UMX0 | Ubiquilin-1 OS=Homo sapiens GN=UBQLN1 PE=1 SV=2 - [UBQL1_HUMAN] | 2.123E7 | 1.736E7 | 1.544E7 | 1.414E7 | 7.327 | 7.239 | 7.189 | 7.150 | 7.283 | 7.170 | 0.114 |
| Q9NP79 | Vacuolar protein sorting-associated protein VTA1 homolog OS=Homo sapiens GN=VTA1 PE=1 SV=1 - [VTA1_ | 2.482E7 | 2.445E7 | 1.818E7 | 1.980E7 | 7.395 | 7.388 | 7.259 | 7.297 | 7.391 | 7.278 | 0.113 |
| P06280 | Alpha-galactosidase A OS=Homo sapiens GN=GLA PE=1 SV=1 - [AGAL_HUMAN] | 1.089E7 | | | 8.392E6 | 7.037 | | | 6.924 | 7.037 | 6.924 | 0.113 |
| P35080-2 | Isoform IIb of Profilin-2 OS=Homo sapiens GN=PFN2 - [PROF2_HUMAN] | 2.144E7 | 1.473E7 | 1.304E7 | 1.442E7 | 7.331 | 7.168 | 7.115 | 7.159 | 7.250 | 7.137 | 0.113 |
| Q9BRA2 | Thioredoxin domain-containing protein 17 OS=Homo sapiens GN=TXNDC17 PE=1 SV=1 - [TXD17_HUMAN] | 2.123E7 | 1.058E7 | 1.105E7 | 1.213E7 | 7.327 | 7.025 | 7.044 | 7.084 | 7.176 | 7.064 | 0.112 |
| E5RI99 | Uncharacterized protein OS=Homo sapiens GN=RPL30 PE=3 SV=1 - [E5RI99_HUMAN] | 4.185E7 | 4.730E7 | 3.092E7 | 3.825E7 | 7.622 | 7.675 | 7.490 | 7.583 | 7.648 | 7.536 | 0.112 |
| P62424 | 60S ribosomal protein L7a OS=Homo sapiens GN=RPL7A PE=1 SV=2 - [RL7A_HUMAN] | 1.868E8 | 1.126E8 | 1.078E8 | 1.167E8 | 8.271 | 8.052 | 8.032 | 8.067 | 8.161 | 8.050 | 0.112 |
| P49588 | Alanyl-tRNA synthetase, cytoplasmic OS=Homo sapiens GN=AARS PE=1 SV=2 - [SYAC_HUMAN] | 3.618E7 | 3.312E7 | 2.665E7 | 2.690E7 | 7.558 | 7.520 | 7.426 | 7.430 | 7.539 | 7.428 | 0.112 |
| P78527 | DNA-dependent protein kinase catalytic subunit OS=Homo sapiens GN=PRKDC PE=1 SV=3 - [PRKDC_HUMAI | 2.179E7 | 2.007E7 | 1.542E7 | 1.700E7 | 7.338 | 7.303 | 7.188 | 7.230 | 7.320 | 7.209 | 0.111 |
| O43660-2 | Isoform 2 of Pleiotropic regulator 1 OS=Homo sapiens GN=PLRG1 - [PLRG1_HUMAN] | 5.138E6 | 2.312E6 | | 2.668E6 | 6.711 | 6.364 | | 6.426 | 6.537 | 6.426 | 0.111 |
| P62495 | Eukaryotic peptide chain release factor subunit 1 OS=Homo sapiens GN=ETF1 PE=1 SV=3 - [ERF1_HUMAN] | 3.619E7 | 2.645E7 | 2.263E7 | 2.536E7 | 7.559 | 7.422 | 7.355 | 7.404 | 7.491 | 7.379 | 0.111 |
| P62701 | 40S ribosomal protein S4, X isoform OS=Homo sapiens GN=RPS4X PE=1 SV=2 - [RS4X_HUMAN] | 8.987E7 | 6.918E7 | 6.171E7 | 6.042E7 | 7.954 | 7.840 | 7.790 | 7.781 | 7.897 | 7.786 | 0.111 |
| P14868 | Aspartyl-tRNA synthetase, cytoplasmic OS=Homo sapiens GN=DARS PE=1 SV=2 - [SYDC_HUMAN] | 2.164E7 | 9.254E6 | 1.068E7 | 1.125E7 | 7.335 | 6.966 | 7.029 | 7.051 | 7.151 | 7.040 | 0.111 |
| Q9NP97 | Dynein light chain roadblock-type 1 OS=Homo sapiens GN=DYNLRB1 PE=1 SV=3 - [DLRB1_HUMAN] | 2.971E7 | 3.002E7 | 2.104E7 | 2.546E7 | 7.473 | 7.477 | 7.323 | 7.406 | 7.475 | 7.364 | 0.111 |
| P51665 | 26S proteasome non-ATPase regulatory subunit 7 OS=Homo sapiens GN=PSMD7 PE=1 SV=2 - [PSD7_HUMA | 3.645E7 | 3.048E7 | 2.529E7 | 2.640E7 | 7.562 | 7.484 | 7.403 | 7.422 | 7.523 | 7.412 | 0.111 |
| P08758 | Annexin A5 OS=Homo sapiens GN=ANXA5 PE=1 SV=2 - [ANXA5_HUMAN] | 1.044E8 | 9.617E7 | 7.546E7 | 7.997E7 | 8.019 | 7.983 | 7.878 | 7.903 | 8.001 | 7.890 | 0.110 |
| C9JA36 | Uncharacterized protein OS=Homo sapiens GN=ATP1B3 PE=3 SV=1 - [C9JA36_HUMAN] | | 1.002E7 | 7.421E6 | 8.138E6 | | 7.001 | 6.870 | 6.911 | 7.001 | 6.890 | 0.110 |
| O00231 | 26S proteasome non-ATPase regulatory subunit 11 OS=Homo sapiens GN=PSMD11 PE=1 SV=3 - [PSD11_HU | 3.086E7 | 2.710E7 | 2.075E7 | 2.428E7 | 7.489 | 7.433 | 7.317 | 7.385 | 7.461 | 7.351 | 0.110 |
| Q99733 | Nucleosome assembly protein 1-like 4 OS=Homo sapiens GN=NAP1L4 PE=1 SV=1 - [NP1L4_HUMAN] | 5.099E7 | 3.270E7 | 3.102E7 | 3.241E7 | 7.708 | 7.514 | 7.492 | 7.511 | 7.611 | 7.501 | 0.110 |
| Q9Y3C8 | Ubiquitin-fold modifier-conjugating enzyme 1 OS=Homo sapiens GN=UFC1 PE=1 SV=3 - [UFC1_HUMAN] | | 6.105E6 | 4.743E6 | | | 6.786 | 6.676 | | 6.786 | 6.676 | 0.110 |
| B4E0R6 | Uncharacterized protein OS=Homo sapiens GN=IPO5 PE=2 SV=1 - [B4E0R6_HUMAN] | 1.935E7 | 7.209E6 | 1.235E7 | 6.828E6 | 7.287 | 6.858 | 7.092 | 6.834 | 7.072 | 6.963 | 0.109 |
| P34949-2 | Isoform 2 of Mannose-6-phosphate isomerase OS=Homo sapiens GN=MPI - [MPI_HUMAN] | 1.492E7 | 1.269E7 | 1.245E7 | 9.198E6 | 7.174 | 7.103 | 7.095 | 6.964 | 7.139 | 7.029 | 0.109 |
| Q3MHD2 | Protein LSM12 homolog OS=Homo sapiens GN=LSM12 PE=1 SV=2 - [LSM12_HUMAN] | 1.662E7 | 5.420E7 | 9.894E6 | 5.509E7 | 7.221 | 7.734 | 6.995 | 7.741 | 7.477 | 7.368 | 0.109 |
| B7Z5F6 | Uncharacterized protein OS=Homo sapiens GN=UBA3 PE=2 SV=1 - [B7Z5F6_HUMAN] | 1.774E7 | | 1.348E7 | 1.415E7 | 7.249 | | 7.130 | 7.151 | 7.249 | 7.140 | 0.109 |
| P34932 | Heat shock 70 kDa protein 4 OS=Homo sapiens GN=HSPA4 PE=1 SV=4 - [HSP74_HUMAN] | 6.885E7 | 5.750E7 | 4.661E7 | 5.158E7 | 7.838 | 7.760 | 7.668 | 7.712 | 7.799 | 7.690 | 0.108 |
| Q8N183 | Mimitin, mitochondrial OS=Homo sapiens GN=NDUFAF2 PE=1 SV=1 - [MIMIT_HUMAN] | 1.089E7 | 8.321E6 | 8.319E6 | 6.624E6 | 7.037 | 6.920 | 6.920 | 6.821 | 6.979 | 6.871 | 0.108 |
| P67936 | Tropomyosin alpha-4 chain OS=Homo sapiens GN=TPM4 PE=1 SV=3 - [TPM4_HUMAN] | 1.388E8 | 7.758E7 | 9.194E7 | 7.128E7 | 8.142 | 7.890 | 7.964 | 7.853 | 8.016 | 7.908 | 0.108 |
| Q92973-2 | Isoform 2 of Transportin-1 OS=Homo sapiens GN=TNPO1 - [TNPO1_HUMAN] | 1.353E7 | 1.286E7 | 1.032E7 | 1.027E7 | 7.131 | 7.109 | 7.014 | 7.011 | 7.120 | 7.013 | 0.108 |
| P49773 | Histidine triad nucleotide-binding protein 1 OS=Homo sapiens GN=HINT1 PE=1 SV=2 - [HINT1_HUMAN] | 7.307E7 | 7.222E7 | 5.563E7 | 5.780E7 | 7.864 | 7.859 | 7.745 | 7.762 | 7.861 | 7.754 | 0.108 |
| O43852 | Calumenin OS=Homo sapiens GN=CALU PE=1 SV=2 - [CALU_HUMAN] | 3.206E7 | 1.645E7 | 1.803E7 | 1.784E7 | 7.506 | 7.216 | 7.256 | 7.251 | 7.361 | 7.254 | 0.107 |
| Q07955 | Serine/arginine-rich splicing factor 1 OS=Homo sapiens GN=SRSF1 PE=1 SV=2 - [SRSF1_HUMAN] | 4.343E7 | 3.375E7 | 3.521E7 | 2.541E7 | 7.638 | 7.528 | 7.547 | 7.405 | 7.583 | 7.476 | 0.107 |
| P61160 | Actin-related protein 2 OS=Homo sapiens GN=ACTR2 PE=1 SV=1 - [ARP2_HUMAN] | 2.261E7 | 8.305E6 | 8.429E6 | 1.363E7 | 7.354 | 6.919 | 6.926 | 7.134 | 7.137 | 7.030 | 0.107 |
| P49321 | Nuclear autoantigenic sperm protein OS=Homo sapiens GN=NASP PE=1 SV=2 - [NASP_HUMAN] | 7.403E7 | 6.835E7 | 5.028E7 | 6.166E7 | 7.869 | 7.835 | 7.701 | 7.790 | 7.852 | 7.746 | 0.106 |
| P78347-2 | Isoform 2 of General transcription factor II-I OS=Homo sapiens GN=GTF2I - [GTF2I_HUMAN] | 1.420E7 | 1.496E7 | 1.038E7 | 1.254E7 | 7.152 | 7.175 | 7.016 | 7.098 | 7.164 | 7.057 | 0.106 |
| Q8N4V1 | Membrane magnesium transporter 1 OS=Homo sapiens GN=MMGT1 PE=1 SV=1 - [MMGT1_HUMAN] | 2.123E7 | 7.441E6 | | 9.845E6 | 7.327 | 6.872 | | 6.993 | 7.099 | 6.993 | 0.106 |
| O43684-2 | Isoform 2 of Mitotic checkpoint protein BUB3 OS=Homo sapiens GN=BUB3 - [BUB3_HUMAN] | 2.174E7 | 1.666E7 | 1.471E7 | 1.513E7 | 7.337 | 7.222 | 7.168 | 7.180 | 7.280 | 7.174 | 0.106 |
| P13647 | Keratin, type II cytoskeletal 5 OS=Homo sapiens GN=KRT5 PE=1 SV=3 - [K2C5_HUMAN] | 1.042E8 | 8.028E7 | 6.385E7 | 8.053E7 | 8.018 | 7.905 | 7.805 | 7.906 | 7.961 | 7.856 | 0.106 |
| P10768 | S-formylglutathione hydrolase OS=Homo sapiens GN=ESD PE=1 SV=2 - [ESTD_HUMAN] | 3.737E7 | 3.370E7 | 2.933E7 | 2.643E7 | 7.573 | 7.528 | 7.467 | 7.422 | 7.550 | 7.445 | 0.105 |
| E7EPB3 | Uncharacterized protein OS=Homo sapiens GN=RPL14 PE=4 SV=1 - [E7EPB3_HUMAN] | 1.115E8 | 6.156E7 | 7.221E7 | 5.864E7 | 8.047 | 7.789 | 7.859 | 7.768 | 7.918 | 7.813 | 0.105 |
| B4E2W0 | Uncharacterized protein OS=Homo sapiens GN=HADHB PE=2 SV=1 - [B4E2W0_HUMAN] | 1.234E7 | 1.238E7 | 9.706E6 | 9.729E6 | 7.091 | 7.093 | 6.987 | 6.988 | 7.092 | 6.988 | 0.105 |
| P43686 | 26S protease regulatory subunit 6B OS=Homo sapiens GN=PSMC4 PE=1 SV=2 - [PRS6B_HUMAN] | 4.235E7 | 3.761E7 | 3.065E7 | 3.213E7 | 7.627 | 7.575 | 7.486 | 7.507 | 7.601 | 7.497 | 0.104 |
| P26641 | Elongation factor 1-gamma OS=Homo sapiens GN=EEF1G PE=1 SV=3 - [EF1G_HUMAN] | 1.852E8 | 1.429E8 | 1.227E8 | 1.335E8 | 8.268 | 8.155 | 8.089 | 8.126 | 8.211 | 8.107 | 0.104 |
| P62269 | 40S ribosomal protein S18 OS=Homo sapiens GN=RPS18 PE=1 SV=3 - [RS18_HUMAN] | 9.786E7 | 5.582E7 | 5.591E7 | 6.053E7 | 7.991 | 7.747 | 7.747 | 7.782 | 7.869 | 7.765 | 0.104 |
| P09211 | Glutathione S-transferase P OS=Homo sapiens GN=GSTP1 PE=1 SV=2 - [GSTP1_HUMAN] | 2.905E8 | 2.268E8 | 1.986E8 | 2.057E8 | 8.463 | 8.356 | 8.298 | 8.313 | 8.409 | 8.306 | 0.104 |
| P30041 | Peroxiredoxin-6 OS=Homo sapiens GN=PRDX6 PE=1 SV=3 - [PRDX6_HUMAN] | 1.633E8 | 1.581E8 | 1.226E8 | 1.309E8 | 8.213 | 8.199 | 8.088 | 8.117 | 8.206 | 8.103 | 0.103 |
| B7Z9F1 | Serine hydroxymethyltransferase OS=Homo sapiens GN=SHMT2 PE=2 SV=1 - [B7Z9F1_HUMAN] | 9.250E7 | 7.019E7 | 6.279E7 | 6.430E7 | 7.966 | 7.846 | 7.798 | 7.808 | 7.906 | 7.803 | 0.103 |
| O14818 | Proteasome subunit alpha type-7 OS=Homo sapiens GN=PSMA7 PE=1 SV=1 - [PSA7_HUMAN] | 6.469E7 | 4.597E7 | 4.414E7 | 4.189E7 | 7.811 | 7.662 | 7.645 | 7.622 | 7.737 | 7.633 | 0.103 |
| P52732 | Kinesin-like protein KIF11 OS=Homo sapiens GN=KIF11 PE=1 SV=2 - [KIF11_HUMAN] | 3.721E6 | 4.113E6 | | 3.085E6 | 6.571 | 6.614 | | 6.489 | 6.592 | 6.489 | 0.103 |
| P82979 | SAP domain-containing ribonucleoprotein OS=Homo sapiens GN=SARNP PE=1 SV=3 - [SARNP_HUMAN] | 1.977E7 | 1.305E7 | 1.261E7 | 1.274E7 | 7.296 | 7.116 | 7.101 | 7.105 | 7.206 | 7.103 | 0.103 |
| Q9Y2Q3-4 | Isoform 4 of Glutathione S-transferase kappa 1 OS=Homo sapiens GN=GSTK1 - [GSTK1_HUMAN] | 1.678E7 | 1.576E7 | 1.271E7 | 1.296E7 | 7.225 | 7.197 | 7.104 | 7.113 | 7.211 | 7.108 | 0.103 |
| Q9NTK5 | Obg-like ATPase 1 OS=Homo sapiens GN=OLA1 PE=1 SV=2 - [OLA1_HUMAN] | 2.787E7 | 2.602E7 | 2.438E7 | 1.853E7 | 7.445 | 7.415 | 7.387 | 7.268 | 7.430 | 7.327 | 0.103 |
| P52565 | Rho GDP-dissociation inhibitor 1 OS=Homo sapiens GN=ARHGDIA PE=1 SV=3 - [GDIR1_HUMAN] | 1.366E8 | 1.089E8 | 8.117E7 | 1.142E8 | 8.135 | 8.037 | 7.909 | 8.058 | 8.086 | 7.984 | 0.103 |
| F5H668 | Uncharacterized protein OS=Homo sapiens GN=RAB6A PE=3 SV=1 - [F5H668_HUMAN] | 6.076E7 | 1.467E7 | 3.098E7 | 1.794E7 | 7.784 | 7.166 | 7.491 | 7.254 | 7.475 | 7.372 | 0.103 |
| Q92499 | ATP-dependent RNA helicase DDX1 OS=Homo sapiens GN=DDX1 PE=1 SV=2 - [DDX1_HUMAN] | 3.069E7 | 2.756E7 | 2.314E7 | 2.279E7 | 7.487 | 7.440 | 7.364 | 7.358 | 7.464 | 7.361 | 0.103 |
| Q5VYK3 | Proteasome-associated protein ECM29 homolog OS=Homo sapiens GN=ECM29 PE=1 SV=2 - [ECM29_HUMAI | 4.974E6 | | 2.811E6 | 5.499E6 | 6.697 | | 6.449 | 6.740 | 6.697 | 6.595 | 0.102 |
| F5GX33 | Uncharacterized protein OS=Homo sapiens GN=SLC25A13 PE=3 SV=1 - [F5GX33_HUMAN] | 7.800E6 | 1.177E7 | 7.064E6 | 8.122E6 | 6.892 | 7.071 | 6.849 | 6.910 | 6.981 | 6.879 | 0.102 |
| Q0VDF9 | Heat shock 70 kDa protein 14 OS=Homo sapiens GN=HSPA14 PE=1 SV=1 - [HSP7E_HUMAN] | 1.492E7 | 1.200E7 | 1.039E7 | 1.079E7 | 7.174 | 7.079 | 7.016 | 7.033 | 7.127 | 7.025 | 0.102 |
| Q15293 | Reticulocalbin-1 OS=Homo sapiens GN=RCN1 PE=1 SV=1 - [RCN1_HUMAN] | 2.201E7 | 2.025E7 | 1.685E7 | 1.655E7 | 7.343 | 7.306 | 7.227 | 7.219 | 7.325 | 7.223 | 0.102 |
| F5H399 | Uncharacterized protein OS=Homo sapiens GN=RAB2A PE=3 SV=1 - [F5H399_HUMAN] | | 8.697E6 | 7.218E6 | 6.578E6 | | 6.939 | 6.858 | 6.818 | 6.939 | 6.838 | 0.101 |
| P62333 | 26S protease regulatory subunit 10B OS=Homo sapiens GN=PSMC6 PE=1 SV=1 - [PRS10_HUMAN] | 3.244E7 | 2.674E7 | 2.186E7 | 2.497E7 | 7.511 | 7.427 | 7.340 | 7.397 | 7.469 | 7.369 | 0.101 |
| P14678-2 | Isoform SM-B of Small nuclear ribonucleoprotein-associated proteins B and B' OS=Homo sapiens GN=SNRPB | 6.876E7 | 5.568E7 | 4.253E7 | 5.665E7 | 7.837 | 7.746 | 7.629 | 7.753 | 7.791 | 7.691 | 0.101 |
| A6NLM8 | Uncharacterized protein OS=Homo sapiens GN=SSR4 PE=4 SV=1 - [A6NLM8_HUMAN] | 2.856E7 | 2.374E7 | 2.110E7 | 2.025E7 | 7.456 | 7.376 | 7.324 | 7.307 | 7.416 | 7.315 | 0.100 |
| Q96C36 | Pyrroline-5-carboxylate reductase 2 OS=Homo sapiens GN=PYCR2 PE=1 SV=1 - [P5CR2_HUMAN] | 2.951E7 | 2.569E7 | 2.127E7 | 2.252E7 | 7.470 | 7.410 | 7.328 | 7.353 | 7.440 | 7.340 | 0.100 |
| P62158 | Calmodulin OS=Homo sapiens GN=CALM1 PE=1 SV=2 - [CALM_HUMAN] | 4.880E8 | 3.857E8 | 3.311E8 | 3.593E8 | 8.688 | 8.586 | 8.520 | 8.555 | 8.637 | 8.538 | 0.100 |
| B4DK30 | Uncharacterized protein OS=Homo sapiens GN=EFTUD2 PE=2 SV=1 - [B4DK30_HUMAN] | 1.997E7 | 1.383E7 | 1.261E7 | 1.387E7 | 7.300 | 7.141 | 7.101 | 7.142 | 7.221 | 7.121 | 0.099 |
| Q9Y224 | UPF0568 protein C14orf166 OS=Homo sapiens GN=C14orf166 PE=1 SV=1 - [CN166_HUMAN] | 3.246E7 | 2.134E7 | 2.038E7 | 2.152E7 | 7.511 | 7.329 | 7.309 | 7.333 | 7.420 | 7.321 | 0.099 |
| Q99426 | Tubulin-folding cofactor B OS=Homo sapiens GN=TBCB PE=1 SV=2 - [TBCB_HUMAN] | 1.713E7 | 1.077E7 | 1.297E7 | 9.008E6 | 7.234 | 7.032 | 7.113 | 6.955 | 7.133 | 7.034 | 0.099 |
| Q9C005 | Protein dpy-30 homolog OS=Homo sapiens GN=DPY30 PE=1 SV=1 - [DPY30_HUMAN] | 1.916E7 | 2.322E7 | 1.687E7 | 1.671E7 | 7.282 | 7.366 | 7.227 | 7.223 | 7.324 | 7.225 | 0.099 |
| P63104 | 14-3-3 protein zeta/delta OS=Homo sapiens GN=YWHAZ PE=1 SV=1 - [1433Z_HUMAN] | 2.267E8 | 1.871E8 | 1.594E8 | 1.686E8 | 8.355 | 8.272 | 8.202 | 8.227 | 8.314 | 8.215 | 0.099 |
| Q16836 | Hydroxyacyl-coenzyme A dehydrogenase, mitochondrial OS=Homo sapiens GN=HADH PE=1 SV=3 - [HCDH_I | 3.621E7 | 4.539E7 | 3.079E7 | 3.385E7 | 7.559 | 7.657 | 7.488 | 7.530 | 7.608 | 7.509 | 0.099 |
| Q9Y265 | RuvB-like 1 OS=Homo sapiens GN=RUVBL1 PE=1 SV=1 - [RUVB1_HUMAN] | 7.253E7 | 4.463E7 | 4.921E7 | 4.172E7 | 7.861 | 7.650 | 7.692 | 7.620 | 7.755 | 7.656 | 0.099 |
| P46783 | 40S ribosomal protein S10 OS=Homo sapiens GN=RPS10 PE=1 SV=1 - [RS10_HUMAN] | 8.236E7 | 6.222E7 | 5.503E7 | 5.908E7 | 7.916 | 7.794 | 7.741 | 7.771 | 7.855 | 7.756 | 0.099 |
| Q96K17 | Transcription factor BTF3 homolog 4 OS=Homo sapiens GN=BTF3L4 PE=1 SV=1 - [BT3L4_HUMAN] | 4.031E7 | 3.723E7 | 3.754E7 | 2.536E7 | 7.605 | 7.571 | 7.575 | 7.404 | 7.588 | 7.489 | 0.099 |
| P35613-2 | Isoform 2 of Basigin OS=Homo sapiens GN=BSG - [BASI_HUMAN] | 3.794E7 | 9.548E6 | 1.350E7 | 1.704E7 | 7.579 | 6.980 | 7.130 | 7.232 | 7.279 | 7.181 | 0.098 |
| P50213 | Isocitrate dehydrogenase [NAD] subunit alpha, mitochondrial OS=Homo sapiens GN=IDH3A PE=1 SV=1 - [II | 1.152E7 | 8.560E6 | 8.174E6 | 7.674E6 | 7.061 | 6.932 | 6.912 | 6.885 | 6.997 | 6.899 | 0.098 |
| P16403 | Histone H1.2 OS=Homo sapiens GN=HIST1H1C PE=1 SV=2 - [H12_HUMAN] | 1.307E8 | 9.073E7 | 8.372E7 | 9.022E7 | 8.116 | 7.958 | 7.923 | 7.955 | 8.037 | 7.939 | 0.098 |
| Q86UP2-2 | Isoform 2 of Kinectin OS=Homo sapiens GN=KTN1 - [KTN1_HUMAN] | 6.865E6 | 3.807E6 | 4.081E6 | | 6.837 | 6.581 | 6.611 | | 6.709 | 6.611 | 0.098 |
| F5H5L4 | Uncharacterized protein OS=Homo sapiens GN=MRPL1 PE=4 SV=1 - [F5H5L4_HUMAN] | | 5.269E6 | | 4.209E6 | | 6.722 | | 6.624 | 6.722 | 6.624 | 0.097 |
| P62750 | 60S ribosomal protein L23a OS=Homo sapiens GN=RPL23A PE=1 SV=1 - [RL23A_HUMAN] | 1.926E8 | 1.126E8 | 1.123E8 | 1.235E8 | 8.285 | 8.052 | 8.050 | 8.092 | 8.168 | 8.071 | 0.097 |
| B4DWS4 | Uncharacterized protein OS=Homo sapiens GN=SLC1A5 PE=2 SV=1 - [B4DWS4_HUMAN] | 3.275E7 | 2.495E7 | 2.224E7 | 2.353E7 | 7.515 | 7.397 | 7.347 | 7.372 | 7.456 | 7.359 | 0.097 |
| P26599 | Polypyrimidine tract-binding protein 1 OS=Homo sapiens GN=PTBP1 PE=1 SV=1 - [PTBP1_HUMAN] | 9.261E7 | 9.045E7 | 7.330E7 | 7.328E7 | 7.967 | 7.956 | 7.865 | 7.865 | 7.962 | 7.865 | 0.096 |
| P22102 | Trifunctional purine biosynthetic protein adenosine-3 OS=Homo sapiens GN=GART PE=1 SV=1 - [PUR2_HUM | 5.290E7 | 4.450E7 | 3.827E7 | 3.956E7 | 7.723 | 7.648 | 7.583 | 7.597 | 7.686 | 7.590 | 0.096 |
| P49189 | 4-trimethylaminobutyraldehyde dehydrogenase OS=Homo sapiens GN=ALDH9A1 PE=1 SV=3 - [AL9A1_HUM/ | 2.635E7 | 1.450E7 | 1.808E7 | 1.361E7 | 7.421 | 7.161 | 7.257 | 7.134 | 7.291 | 7.196 | 0.096 |
| Q5T6H7 | Uncharacterized protein OS=Homo sapiens GN=XPNPEP1 PE=2 SV=1 - [Q5T6H7_HUMAN] | 1.280E7 | 8.694E6 | | 8.467E6 | 7.107 | 6.939 | | 6.928 | 7.023 | 6.928 | 0.095 |
| Q9UHX1-4 | Isoform 4 of Poly(U)-binding-splicing factor PUF60 OS=Homo sapiens GN=PUF60 - [PUF60_HUMAN] | 5.219E7 | 5.039E7 | 4.018E7 | 4.222E7 | 7.718 | 7.702 | 7.604 | 7.625 | 7.710 | 7.615 | 0.095 |
| P26639 | Threonyl-tRNA synthetase, cytoplasmic OS=Homo sapiens GN=TARS PE=1 SV=3 - [SYTC_HUMAN] | 1.322E7 | 1.245E7 | 8.320E6 | 1.283E7 | 7.121 | 7.095 | 6.920 | 7.108 | 7.108 | 7.014 | 0.094 |
| P37108 | Signal recognition particle 14 kDa protein OS=Homo sapiens GN=SRP14 PE=1 SV=2 - [SRP14_HUMAN] | 3.322E7 | 2.747E7 | | 2.436E7 | 7.521 | 7.439 | | 7.387 | 7.480 | 7.387 | 0.093 |
| Q96QK1 | Vacuolar protein sorting-associated protein 35 OS=Homo sapiens GN=VPS35 PE=1 SV=2 - [VPS35_HUMAN] | 1.834E7 | 1.658E7 | 1.318E7 | 1.500E7 | 7.263 | 7.219 | 7.120 | 7.176 | 7.241 | 7.148 | 0.093 |
| B4DIR6 | Uncharacterized protein OS=Homo sapiens GN=CHMP5 PE=2 SV=1 - [B4DIR6_HUMAN] | 1.712E7 | 2.825E7 | 1.279E7 | 2.463E7 | 7.234 | 7.451 | 7.107 | 7.391 | 7.342 | 7.249 | 0.093 |
| F5H345 | Uncharacterized protein OS=Homo sapiens GN=HMBS PE=4 SV=1 - [F5H345_HUMAN] | 1.449E7 | 1.091E7 | 1.110E7 | 9.291E6 | 7.161 | 7.038 | 7.045 | 6.968 | 7.099 | 7.007 | 0.093 |
| Q13526 | Peptidyl-prolyl cis-trans isomerase NIMA-interacting 1 OS=Homo sapiens GN=PIN1 PE=1 SV=1 - [PIN1_HUM. | 1.813E7 | 1.638E7 | 1.411E7 | 1.375E7 | 7.258 | 7.214 | 7.149 | 7.138 | 7.236 | 7.144 | 0.093 |
| P63241 | Eukaryotic translation initiation factor 5A-1 OS=Homo sapiens GN=EIF5A PE=1 SV=2 - [IF5A1_HUMAN] | 1.488E8 | 1.368E8 | 1.157E8 | 1.150E8 | 8.173 | 8.136 | 8.063 | 8.061 | 8.154 | 8.062 | 0.092 |
| O95202 | LETM1 and EF-hand domain-containing protein 1, mitochondrial OS=Homo sapiens GN=LETM1 PE=1 SV=1 - | 1.456E7 | 1.545E7 | 1.190E7 | 1.238E7 | 7.163 | 7.189 | 7.075 | 7.093 | 7.176 | 7.084 | 0.092 |
| E9PAV3 | Uncharacterized protein OS=Homo sapiens GN=NACA PE=4 SV=1 - [E9PAV3_HUMAN] | 1.139E8 | 8.787E7 | 7.594E7 | 8.647E7 | 8.056 | 7.944 | 7.880 | 7.937 | 8.000 | 7.909 | 0.091 |
| Q99714 | 3-hydroxyacyl-CoA dehydrogenase type-2 OS=Homo sapiens GN=HSD17B10 PE=1 SV=3 - [HCD2_HUMAN] | 6.292E7 | 5.078E7 | 4.671E7 | 4.492E7 | 7.799 | 7.706 | 7.669 | 7.652 | 7.752 | 7.661 | 0.091 |
| Q6YN16-2 | Isoform 2 of Hydroxysteroid dehydrogenase-like protein 2 OS=Homo sapiens GN=HSDL2 - [HSDL2_HUMAN] | 1.124E7 | 9.687E6 | 8.301E6 | 8.621E6 | 7.051 | 6.986 | 6.919 | 6.936 | 7.018 | 6.927 | 0.091 |
| Q06830 | Peroxiredoxin-1 OS=Homo sapiens GN=PRDX1 PE=1 SV=1 - [PRDX1_HUMAN] | 4.413E8 | 3.131E8 | 2.929E8 | 3.106E8 | 8.645 | 8.496 | 8.467 | 8.492 | 8.570 | 8.479 | 0.091 |
| P60891 | Ribose-phosphate pyrophosphokinase 1 OS=Homo sapiens GN=PRPS1 PE=1 SV=2 - [PRPS1_HUMAN] | 3.843E7 | 2.752E7 | 2.987E7 | 2.334E7 | 7.585 | 7.440 | 7.475 | 7.368 | 7.512 | 7.422 | 0.091 |
| P49207 | 60S ribosomal protein L34 OS=Homo sapiens GN=RPL34 PE=1 SV=3 - [RL34_HUMAN] | 3.086E7 | 1.971E7 | 1.870E7 | 2.146E7 | 7.489 | 7.295 | 7.272 | 7.332 | 7.392 | 7.302 | 0.090 |
| Q8NC51-2 | Isoform 2 of Plasminogen activator inhibitor 1 RNA-binding protein OS=Homo sapiens GN=SERBP1 - [PAIRB_ | 1.164E8 | 8.694E7 | 8.160E7 | 8.182E7 | 8.066 | 7.939 | 7.912 | 7.913 | 8.002 | 7.912 | 0.090 |
| E5RIW3 | Uncharacterized protein OS=Homo sapiens GN=TBCA PE=4 SV=1 - [E5RIW3_HUMAN] | 1.329E8 | 9.987E7 | 9.452E7 | 9.288E7 | 8.124 | 7.999 | 7.976 | 7.968 | 8.061 | 7.972 | 0.090 |
| P26373 | 60S ribosomal protein L13 OS=Homo sapiens GN=RPL13 PE=1 SV=4 - [RL13_HUMAN] | 1.110E8 | 9.654E7 | 8.427E7 | 8.436E7 | 8.045 | 7.985 | 7.926 | 7.926 | 8.015 | 7.926 | 0.089 |
| B1ANR0 | Poly(A) binding protein, cytoplasmic 4 (Inducible form) OS=Homo sapiens GN=PABPC4 PE=4 SV=1 - [B1ANR | 7.411E7 | 6.919E7 | 5.446E7 | 6.262E7 | 7.870 | 7.840 | 7.736 | 7.797 | 7.855 | 7.766 | 0.089 |
| P31937 | 3-hydroxyisobutyrate dehydrogenase, mitochondrial OS=Homo sapiens GN=HIBADH PE=1 SV=2 - [3HIDH_H | 1.448E7 | 1.153E7 | 7.991E6 | 1.391E7 | 7.161 | 7.062 | 6.903 | 7.143 | 7.111 | 7.023 | 0.088 |
| P83731 | 60S ribosomal protein L24 OS=Homo sapiens GN=RPL24 PE=1 SV=1 - [RL24_HUMAN] | 7.857E7 | 6.362E7 | 5.706E7 | 5.844E7 | 7.895 | 7.804 | 7.756 | 7.767 | 7.849 | 7.762 | 0.088 |
| P17987 | T-complex protein 1 subunit alpha OS=Homo sapiens GN=TCP1 PE=1 SV=1 - [TCPA_HUMAN] | 9.094E7 | 8.437E7 | 7.225E7 | 7.103E7 | 7.959 | 7.926 | 7.859 | 7.851 | 7.942 | 7.855 | 0.087 |
| P12277 | Creatine kinase B-type OS=Homo sapiens GN=CKB PE=1 SV=1 - [KCRB_HUMAN] | 7.189E8 | 6.682E8 | 4.959E8 | 6.486E8 | 8.857 | 8.825 | 8.695 | 8.812 | 8.841 | 8.754 | 0.087 |
| P17844 | Probable ATP-dependent RNA helicase DDX5 OS=Homo sapiens GN=DDX5 PE=1 SV=1 - [DDX5_HUMAN] | 4.155E7 | 2.688E7 | 2.475E7 | 3.023E7 | 7.619 | 7.429 | 7.394 | 7.480 | 7.524 | 7.437 | 0.087 |
| O43447 | Peptidyl-prolyl cis-trans isomerase H OS=Homo sapiens GN=PPIH PE=1 SV=1 - [PPIH_HUMAN] | 1.895E7 | 1.659E7 | 1.444E7 | 1.459E7 | 7.278 | 7.220 | 7.160 | 7.164 | 7.249 | 7.162 | 0.087 |
| P09012 | U1 small nuclear ribonucleoprotein A OS=Homo sapiens GN=SNRPA PE=1 SV=3 - [SNRPA_HUMAN] | 2.951E7 | | 2.291E7 | 2.549E7 | 7.470 | | 7.360 | 7.406 | 7.470 | 7.383 | 0.087 |
| P09234 | U1 small nuclear ribonucleoprotein C OS=Homo sapiens GN=SNRPC PE=1 SV=1 - [RU1C_HUMAN] | 3.419E7 | 1.775E7 | 2.459E7 | 1.664E7 | 7.534 | 7.249 | 7.391 | 7.221 | 7.392 | 7.306 | 0.086 |
| Q14697 | Neutral alpha-glucosidase AB OS=Homo sapiens GN=GANAB PE=1 SV=3 - [GANAB_HUMAN] | 6.688E7 | 4.502E7 | 4.318E7 | 4.708E7 | 7.825 | 7.653 | 7.635 | 7.673 | 7.739 | 7.654 | 0.085 |
| B4DV79 | Uncharacterized protein OS=Homo sapiens GN=EIF3B PE=2 SV=1 - [B4DV79_HUMAN] | 2.265E7 | 2.142E7 | 1.695E7 | 1.937E7 | 7.355 | 7.331 | 7.229 | 7.287 | 7.343 | 7.258 | 0.085 |
| E9PM92 | Uncharacterized protein OS=Homo sapiens GN=C11orf58 PE=4 SV=1 - [E9PM92_HUMAN] | | 4.340E7 | 3.395E7 | 3.757E7 | | 7.637 | 7.531 | 7.575 | 7.637 | 7.553 | 0.085 |
| Q96A33-2 | Isoform 2 of Coiled-coil domain-containing protein 47 OS=Homo sapiens GN=CCDC47 - [CCD47_HUMAN] | 8.202E6 | 1.076E7 | 6.942E6 | 8.630E6 | 6.914 | 7.032 | 6.842 | 6.936 | 6.973 | 6.889 | 0.084 |
| P52292 | Importin subunit alpha-2 OS=Homo sapiens GN=KPNA2 PE=1 SV=1 - [IMA2_HUMAN] | 2.355E7 | 1.799E7 | 1.674E7 | 1.719E7 | 7.372 | 7.255 | 7.224 | 7.235 | 7.314 | 7.230 | 0.084 |
| F8VPD4 | Uncharacterized protein OS=Homo sapiens GN=CAD PE=3 SV=1 - [F8VPD4_HUMAN] | 2.156E7 | 2.689E7 | 1.960E7 | 2.011E7 | 7.334 | 7.430 | 7.292 | 7.303 | 7.382 | 7.298 | 0.084 |
| P07741 | Adenine phosphoribosyltransferase OS=Homo sapiens GN=APRT PE=1 SV=2 - [APT_HUMAN] | 5.207E7 | 4.094E7 | 3.981E7 | 3.652E7 | 7.717 | 7.612 | 7.600 | 7.562 | 7.664 | 7.581 | 0.083 |
| P37802 | Transgelin-2 OS=Homo sapiens GN=TAGLN2 PE=1 SV=3 - [TAGL2_HUMAN] | 1.042E8 | 8.449E7 | 7.722E7 | 7.798E7 | 8.018 | 7.927 | 7.888 | 7.892 | 7.972 | 7.890 | 0.083 |
| P36957 | Dihydrolipoyllysine-residue succinyltransferase component of 2-oxoglutarate dehydrogenase complex, mitoch | 1.640E7 | 1.054E7 | 8.811E6 | 1.342E7 | 7.215 | 7.023 | 6.945 | 7.128 | 7.119 | 7.036 | 0.082 |

| | | | | | | | | | | | | |
|---|---|---|---|---|---|---|---|---|---|---|---|---|
| Q9BXJ9 | N-alpha-acetyltransferase 15, NatA auxiliary subunit OS=Homo sapiens GN=NAA15 PE=1 SV=1 - [NAA15_HUMAN] | | 3.919E6 | 3.363E6 | 3.128E6 | | 6.593 | 6.527 | 6.495 | 6.593 | 6.511 | 0.082 |
| P30566 | Adenylosuccinate lyase OS=Homo sapiens GN=ADSL PE=1 SV=2 - [PUR8_HUMAN] | 2.349E7 | 9.293E6 | 1.110E7 | 1.348E7 | 7.371 | 6.968 | 7.045 | 7.130 | 7.170 | 7.087 | 0.082 |
| Q14683 | Structural maintenance of chromosomes protein 1A OS=Homo sapiens GN=SMC1A PE=1 SV=2 - [SMC1A_HU | 5.151E6 | 4.733E6 | 4.010E6 | 4.168E6 | 6.712 | 6.675 | 6.603 | 6.620 | 6.693 | 6.612 | 0.082 |
| E9PN17 | Uncharacterized protein OS=Homo sapiens GN=ATP5L PE=4 SV=1 - [E9PN17_HUMAN] | 1.417E7 | 8.817E6 | 8.783E6 | 9.759E6 | 7.151 | 6.945 | 6.944 | 6.989 | 7.048 | 6.967 | 0.082 |
| P10412 | Histone H1.4 OS=Homo sapiens GN=HIST1H1E PE=1 SV=2 - [H14_HUMAN] | 1.307E8 | 1.046E8 | 9.086E7 | 1.032E8 | 8.116 | 8.019 | 7.958 | 8.014 | 8.068 | 7.986 | 0.082 |
| O43776 | Asparaginyl-tRNA synthetase, cytoplasmic OS=Homo sapiens GN=NARS PE=1 SV=1 - [SYNC_HUMAN] | 2.225E7 | 2.064E7 | 1.751E7 | 1.802E7 | 7.347 | 7.315 | 7.243 | 7.256 | 7.331 | 7.249 | 0.082 |
| G3XAL7 | Putative uncharacterized protein LOC133619 OS=Homo sapiens GN=PRRC1 PE=4 SV=1 - [G3XAL7_HUMAN] | 8.050E6 | 7.962E6 | 7.260E6 | 6.072E6 | 6.906 | 6.901 | 6.861 | 6.783 | 6.903 | 6.822 | 0.081 |
| P54136 | Arginyl-tRNA synthetase, cytoplasmic OS=Homo sapiens GN=RARS PE=1 SV=2 - [SYRC_HUMAN] | 1.834E7 | 1.774E7 | 1.430E7 | 1.578E7 | 7.263 | 7.249 | 7.155 | 7.198 | 7.256 | 7.177 | 0.079 |
| P49755 | Transmembrane emp24 domain-containing protein 10 OS=Homo sapiens GN=TMED10 PE=1 SV=2 - [TMEDA | 2.010E7 | 1.555E7 | 1.451E7 | 1.495E7 | 7.303 | 7.192 | 7.162 | 7.175 | 7.247 | 7.168 | 0.079 |
| Q15287-3 | Isoform 3 of RNA-binding protein with serine-rich domain 1 OS=Homo sapiens GN=RNPS1 - [RNPS1_HUMAN | 1.921E7 | 1.412E7 | 1.304E7 | 1.447E7 | 7.283 | 7.150 | 7.115 | 7.160 | 7.217 | 7.138 | 0.079 |
| P62854 | 40S ribosomal protein S26 OS=Homo sapiens GN=RPS26 PE=1 SV=3 - [RS26_HUMAN] | 9.018E7 | 5.593E7 | 5.963E7 | 5.883E7 | 7.955 | 7.748 | 7.775 | 7.770 | 7.851 | 7.773 | 0.079 |
| A8K1H6 | COP9 constitutive photomorphogenic homolog subunit 8 (Arabidopsis), isoform CRA_b OS=Homo sapiens GN | 3.788E7 | 6.441E6 | 1.304E7 | | 7.578 | 6.809 | 7.115 | | 7.194 | 7.115 | 0.078 |
| P23396 | 40S ribosomal protein S3 OS=Homo sapiens GN=RPS3 PE=1 SV=2 - [RS3_HUMAN] | 1.714E8 | 1.037E8 | 1.113E8 | 1.116E8 | 8.234 | 8.016 | 8.046 | 8.048 | 8.125 | 8.047 | 0.078 |
| P62826 | GTP-binding nuclear protein Ran OS=Homo sapiens GN=RAN PE=1 SV=3 - [RAN_HUMAN] | 2.897E8 | 1.814E8 | 1.889E8 | 1.944E8 | 8.462 | 8.259 | 8.276 | 8.289 | 8.360 | 8.282 | 0.078 |
| P00387-2 | Isoform 2 of NADH-cytochrome b5 reductase 3 OS=Homo sapiens GN=CYB5R3 - [NB5R3_HUMAN] | 1.901E7 | 2.768E7 | 1.830E7 | 2.010E7 | 7.279 | 7.442 | 7.262 | 7.303 | 7.361 | 7.283 | 0.078 |
| Q9NX55 | Huntingtin-interacting protein K OS=Homo sapiens GN=HYPK PE=1 SV=2 - [HYPK_HUMAN] | 1.510E7 | 1.292E7 | 1.028E7 | 1.327E7 | 7.179 | 7.111 | 7.012 | 7.123 | 7.145 | 7.067 | 0.078 |
| B4DZP5 | Uncharacterized protein OS=Homo sapiens GN=DDB1 PE=2 SV=1 - [B4DZP5_HUMAN] | 2.754E7 | 2.579E7 | 2.171E7 | 2.290E7 | 7.440 | 7.411 | 7.337 | 7.360 | 7.426 | 7.348 | 0.077 |
| Q9Y5B9 | FACT complex subunit SPT16 OS=Homo sapiens GN=SUPT16H PE=1 SV=1 - [SP16H_HUMAN] | 1.807E7 | 2.248E7 | 1.791E7 | 1.592E7 | 7.257 | 7.352 | 7.253 | 7.202 | 7.304 | 7.227 | 0.077 |
| P14618 | Pyruvate kinase isozymes M1/M2 OS=Homo sapiens GN=PKM2 PE=1 SV=4 - [KPYM_HUMAN] | 4.179E8 | 3.524E8 | 3.226E8 | 3.220E8 | 8.621 | 8.547 | 8.509 | 8.508 | 8.584 | 8.508 | 0.076 |
| P55327-2 | Isoform 2 of Tumor protein D52 OS=Homo sapiens GN=TPD52 - [TPD52_HUMAN] | 1.010E7 | 1.010E7 | 8.477E6 | 8.529E6 | 7.004 | 7.004 | 6.928 | 6.931 | 7.004 | 6.930 | 0.075 |
| F2Z3A5 | Uncharacterized protein OS=Homo sapiens GN=RPL3 PE=3 SV=1 - [F2Z3A5_HUMAN] | 1.598E8 | 1.244E8 | 1.183E8 | 1.193E8 | 8.204 | 8.095 | 8.073 | 8.077 | 8.149 | 8.075 | 0.074 |
| Q969H8 | UPF0556 protein C19orf10 OS=Homo sapiens GN=C19orf10 PE=1 SV=1 - [CS010_HUMAN] | 2.591E7 | 1.280E7 | 1.301E7 | 1.813E7 | 7.414 | 7.107 | 7.114 | 7.258 | 7.260 | 7.186 | 0.074 |
| C9JLK5 | Uncharacterized protein OS=Homo sapiens GN=EEF1E1 PE=4 SV=1 - [C9JLK5_HUMAN] | 2.928E7 | 2.877E7 | 2.883E7 | 2.083E7 | 7.467 | 7.459 | 7.460 | 7.319 | 7.463 | 7.389 | 0.073 |
| Q86U42-2 | Isoform 2 of Polyadenylate-binding protein 2 OS=Homo sapiens GN=PABPN1 - [PABP2_HUMAN] | 3.409E7 | 1.452E7 | 2.702E7 | 1.307E7 | 7.533 | 7.162 | 7.432 | 7.116 | 7.347 | 7.274 | 0.073 |
| P23528 | Cofilin-1 OS=Homo sapiens GN=CFL1 PE=1 SV=3 - [COF1_HUMAN] | 4.574E8 | 4.126E8 | 3.617E8 | 3.727E8 | 8.660 | 8.616 | 8.558 | 8.571 | 8.638 | 8.565 | 0.073 |
| P62491 | Ras-related protein Rab-11A OS=Homo sapiens GN=RAB11A PE=1 SV=3 - [RB11A_HUMAN] | 3.539E7 | 2.077E7 | 1.991E7 | 2.637E7 | 7.549 | 7.317 | 7.299 | 7.421 | 7.433 | 7.360 | 0.073 |
| Q99829 | Copine-1 OS=Homo sapiens GN=CPNE1 PE=1 SV=1 - [CPNE1_HUMAN] | 1.289E7 | 9.955E6 | 9.828E6 | 9.331E6 | 7.110 | 6.998 | 6.992 | 6.970 | 7.054 | 6.981 | 0.073 |
| Q9UJY4 | ADP-ribosylation factor-binding protein GGA2 OS=Homo sapiens GN=GGA2 PE=1 SV=3 - [GGA2_HUMAN] | 3.416E6 | 8.435E6 | 4.721E6 | 4.367E6 | 6.533 | 6.926 | 6.674 | 6.640 | 6.730 | 6.657 | 0.073 |
| C9JGE3 | Ewing sarcoma breakpoint region 1, isoform CRA_e OS=Homo sapiens GN=EWSR1 PE=4 SV=1 - [C9JGE3_HI | 1.692E7 | 1.719E7 | 1.169E7 | 1.784E7 | 7.229 | 7.235 | 7.068 | 7.251 | 7.232 | 7.160 | 0.072 |
| O00264 | Membrane-associated progesterone receptor component 1 OS=Homo sapiens GN=PGRMC1 PE=1 SV=3 - [PG | 7.299E7 | 4.655E7 | 4.922E7 | 4.947E7 | 7.863 | 7.668 | 7.692 | 7.694 | 7.766 | 7.693 | 0.072 |
| B7Z403 | Uncharacterized protein OS=Homo sapiens GN=GLOD4 PE=2 SV=1 - [B7Z403_HUMAN] | 3.556E7 | 1.403E7 | | 1.893E7 | 7.551 | 7.147 | | 7.277 | 7.349 | 7.277 | 0.072 |
| P60174 | Triosephosphate isomerase OS=Homo sapiens GN=TPI1 PE=1 SV=2 - [TPIS_HUMAN] | 6.116E8 | 3.262E8 | 3.685E8 | 3.887E8 | 8.786 | 8.513 | 8.566 | 8.590 | 8.650 | 8.578 | 0.072 |
| P13645 | Keratin, type I cytoskeletal 10 OS=Homo sapiens GN=KRT10 PE=1 SV=6 - [K1C10_HUMAN] | 4.228E8 | 3.885E8 | 3.422E8 | 3.450E8 | 8.626 | 8.589 | 8.534 | 8.538 | 8.608 | 8.536 | 0.072 |
| P40926 | Malate dehydrogenase, mitochondrial OS=Homo sapiens GN=MDH2 PE=1 SV=3 - [MDHM_HUMAN] | 3.904E8 | 3.292E8 | 3.030E8 | 3.048E8 | 8.592 | 8.517 | 8.482 | 8.484 | 8.554 | 8.483 | 0.072 |
| P68032 | Actin, alpha cardiac muscle 1 OS=Homo sapiens GN=ACTC1 PE=1 SV=1 - [ACTC_HUMAN] | 9.818E8 | 7.885E8 | 7.325E8 | 7.623E8 | 8.992 | 8.897 | 8.865 | 8.882 | 8.944 | 8.873 | 0.071 |
| P06748-2 | Isoform 2 of Nucleophosmin OS=Homo sapiens GN=NPM1 - [NPM_HUMAN] | 6.497E8 | 6.124E8 | 5.221E8 | 5.502E8 | 8.813 | 8.787 | 8.718 | 8.741 | 8.800 | 8.729 | 0.071 |
| D6RAF8 | Uncharacterized protein OS=Homo sapiens GN=HNRNPD PE=4 SV=1 - [D6RAF8_HUMAN] | 1.790E8 | 1.157E8 | 1.277E8 | 1.171E8 | 8.253 | 8.063 | 8.106 | 8.069 | 8.158 | 8.087 | 0.071 |
| O15372 | Eukaryotic translation initiation factor 3 subunit H OS=Homo sapiens GN=EIF3H PE=1 SV=1 - [EIF3H_HUMAI | 1.856E7 | 1.792E7 | 1.518E7 | 1.584E7 | 7.268 | 7.253 | 7.181 | 7.200 | 7.261 | 7.191 | 0.070 |
| E7ERZ4 | Uncharacterized protein OS=Homo sapiens GN=PMPCB PE=3 SV=1 - [E7ERZ4_HUMAN] | 1.917E7 | 1.656E7 | 1.566E7 | 1.467E7 | 7.283 | 7.219 | 7.195 | 7.166 | 7.251 | 7.181 | 0.070 |
| P62266 | 40S ribosomal protein S23 OS=Homo sapiens GN=RPS23 PE=1 SV=3 - [RS23_HUMAN] | 6.681E7 | 5.228E7 | 4.772E7 | 5.317E7 | 7.825 | 7.718 | 7.679 | 7.726 | 7.772 | 7.702 | 0.069 |
| Q9Y3U8 | 60S ribosomal protein L36 OS=Homo sapiens GN=RPL36 PE=1 SV=3 - [RL36_HUMAN] | 4.925E7 | 3.255E7 | 3.406E7 | 3.423E7 | 7.692 | 7.512 | 7.532 | 7.534 | 7.602 | 7.533 | 0.069 |
| Q9Y3A5 | Ribosome maturation protein SBDS OS=Homo sapiens GN=SBDS PE=1 SV=4 - [SBDS_HUMAN] | | 1.577E7 | 1.448E7 | 1.249E7 | | 7.198 | 7.161 | 7.097 | 7.198 | 7.129 | 0.069 |
| Q92688-2 | Isoform 2 of Acidic leucine-rich nuclear phosphoprotein 32 family member B OS=Homo sapiens GN=ANP32B | 6.693E7 | 5.843E7 | 5.083E7 | 5.598E7 | 7.826 | 7.767 | 7.706 | 7.748 | 7.796 | 7.727 | 0.069 |
| P25398 | 40S ribosomal protein S12 OS=Homo sapiens GN=RPS12 PE=1 SV=3 - [RS12_HUMAN] | 1.095E8 | 1.056E8 | 8.524E7 | 9.897E7 | 8.040 | 8.023 | 7.931 | 7.995 | 8.032 | 7.963 | 0.068 |
| Q5T760 | Splicing factor, arginine/serine-rich 11 (Fragment) OS=Homo sapiens GN=SFRS11 PE=2 SV=1 - [Q5T760_HU | 2.201E7 | 2.183E7 | 1.845E7 | 1.900E7 | 7.343 | 7.339 | 7.266 | 7.279 | 7.341 | 7.272 | 0.068 |
| O60888-3 | Isoform C of Protein CutA OS=Homo sapiens GN=CUTA - [CUTA_HUMAN] | 4.990E7 | 3.180E7 | 3.271E7 | 3.546E7 | 7.698 | 7.502 | 7.515 | 7.550 | 7.600 | 7.532 | 0.068 |
| P53041 | Serine/threonine-protein phosphatase 5 OS=Homo sapiens GN=PPP5C PE=1 SV=1 - [PPP5_HUMAN] | | 1.600E7 | 1.381E7 | 1.356E7 | | 7.204 | 7.140 | 7.132 | 7.204 | 7.136 | 0.068 |
| P62906 | 60S ribosomal protein L10a OS=Homo sapiens GN=RPL10A PE=1 SV=2 - [RL10A_HUMAN] | 9.066E7 | 7.058E7 | 6.748E7 | 6.935E7 | 7.957 | 7.849 | 7.829 | 7.841 | 7.903 | 7.835 | 0.068 |
| F5GYQ4 | Glutamate dehydrogenase OS=Homo sapiens GN=GLUD1 PE=3 SV=1 - [F5GYQ4_HUMAN] | 2.568E7 | 2.936E7 | 2.226E7 | 2.478E7 | 7.410 | 7.468 | 7.347 | 7.394 | 7.439 | 7.371 | 0.068 |
| Q9Y3C4 | TP53RK-binding protein OS=Homo sapiens GN=TPRKB PE=1 SV=1 - [TPRKB_HUMAN] | | 4.595E6 | 4.627E6 | 3.339E6 | | 6.662 | 6.665 | 6.524 | 6.662 | 6.594 | 0.068 |
| Q9P258 | Protein RCC2 OS=Homo sapiens GN=RCC2 PE=1 SV=2 - [RCC2_HUMAN] | 2.517E7 | 1.778E7 | 1.952E7 | 1.682E7 | 7.401 | 7.250 | 7.290 | 7.226 | 7.325 | 7.258 | 0.067 |
| Q8WXX5 | DnaJ homolog subfamily C member 9 OS=Homo sapiens GN=DNAJC9 PE=1 SV=1 - [DNJC9_HUMAN] | 1.743E7 | 2.380E7 | 2.158E7 | 1.411E7 | 7.241 | 7.376 | 7.334 | 7.150 | 7.309 | 7.242 | 0.067 |
| P11021 | 78 kDa glucose-regulated protein OS=Homo sapiens GN=HSPA5 PE=1 SV=2 - [GRP78_HUMAN] | 2.334E8 | 1.819E8 | 1.787E8 | 1.748E8 | 8.368 | 8.260 | 8.252 | 8.243 | 8.314 | 8.247 | 0.067 |
| P08708 | 40S ribosomal protein S17 OS=Homo sapiens GN=RPS17 PE=1 SV=2 - [RS17_HUMAN] | 1.596E8 | 7.982E7 | 7.209E7 | 1.302E8 | 8.203 | 7.902 | 7.858 | 8.115 | 8.053 | 7.986 | 0.066 |
| Q8N8S7-2 | Isoform 2 of Protein enabled homolog OS=Homo sapiens GN=ENAH - [ENAH_HUMAN] | 1.746E7 | 1.142E7 | 1.418E7 | 1.037E7 | 7.242 | 7.058 | 7.152 | 7.016 | 7.150 | 7.084 | 0.066 |
| B4DEK4 | Uncharacterized protein OS=Homo sapiens GN=SNX2 PE=2 SV=1 - [B4DEK4_HUMAN] | 1.271E7 | 9.844E6 | 8.595E6 | 1.074E7 | 7.104 | 6.993 | 6.934 | 7.031 | 7.049 | 6.983 | 0.066 |
| P20618 | Proteasome subunit beta type-1 OS=Homo sapiens GN=PSMB1 PE=1 SV=2 - [PSB1_HUMAN] | 2.987E7 | 2.624E7 | 2.435E7 | 2.377E7 | 7.475 | 7.419 | 7.386 | 7.376 | 7.447 | 7.381 | 0.066 |
| O14980 | Exportin-1 OS=Homo sapiens GN=XPO1 PE=1 SV=1 - [XPO1_HUMAN] | 1.176E7 | 1.213E7 | 1.076E7 | 9.804E6 | 7.071 | 7.084 | 7.032 | 6.991 | 7.077 | 7.012 | 0.066 |
| F5H816 | Phosphorylase OS=Homo sapiens GN=PYGL PE=3 SV=1 - [F5H816_HUMAN] | 1.934E7 | 2.612E7 | 1.884E7 | 1.989E7 | 7.286 | 7.417 | 7.275 | 7.299 | 7.352 | 7.287 | 0.065 |
| Q15427 | Splicing factor 3B subunit 4 OS=Homo sapiens GN=SF3B4 PE=1 SV=1 - [SF3B4_HUMAN] | 1.220E7 | 1.074E7 | 1.109E7 | 8.772E6 | 7.086 | 7.031 | 7.045 | 6.943 | 7.059 | 6.994 | 0.065 |
| P28074 | Proteasome subunit beta type-5 OS=Homo sapiens GN=PSMB5 PE=1 SV=3 - [PSB5_HUMAN] | 1.669E7 | 1.763E7 | 1.368E7 | 1.598E7 | 7.222 | 7.246 | 7.136 | 7.203 | 7.234 | 7.170 | 0.064 |
| P62820 | Ras-related protein Rab-1A OS=Homo sapiens GN=RAB1A PE=1 SV=3 - [RAB1A_HUMAN] | 3.738E7 | 2.396E7 | 3.098E7 | 2.150E7 | 7.573 | 7.380 | 7.491 | 7.332 | 7.476 | 7.412 | 0.064 |
| P09661 | U2 small nuclear ribonucleoprotein A' OS=Homo sapiens GN=SNRPA1 PE=1 SV=2 - [RU2A_HUMAN] | 2.914E7 | 2.352E7 | 1.993E7 | 2.563E7 | 7.465 | 7.371 | 7.299 | 7.409 | 7.418 | 7.354 | 0.064 |
| Q9H2W6 | 39S ribosomal protein L46, mitochondrial OS=Homo sapiens GN=MRPL46 PE=1 SV=1 - [RM46_HUMAN] | 1.720E7 | 1.207E7 | | 1.245E7 | 7.236 | 7.082 | | 7.095 | 7.159 | 7.095 | 0.063 |
| P27797 | Calreticulin OS=Homo sapiens GN=CALR PE=1 SV=1 - [CALR_HUMAN] | 1.694E8 | 1.222E8 | 1.231E8 | 1.257E8 | 8.229 | 8.087 | 8.090 | 8.099 | 8.158 | 8.095 | 0.063 |
| D3YTB1 | Uncharacterized protein OS=Homo sapiens GN=RPL32 PE=4 SV=1 - [D3YTB1_HUMAN] | 1.125E8 | 7.632E7 | 8.713E7 | 7.389E7 | 8.051 | 7.883 | 7.940 | 7.869 | 7.967 | 7.904 | 0.063 |
| P46777 | 60S ribosomal protein L5 OS=Homo sapiens GN=RPL5 PE=1 SV=3 - [RL5_HUMAN] | 1.381E8 | 8.820E7 | 8.988E7 | 1.016E8 | 8.140 | 7.945 | 7.954 | 8.007 | 8.043 | 7.980 | 0.063 |
| Q9BWD1 | Acetyl-CoA acetyltransferase, cytosolic OS=Homo sapiens GN=ACAT2 PE=1 SV=2 - [THIC_HUMAN] | 2.906E7 | 2.981E7 | 2.621E7 | 2.486E7 | 7.463 | 7.474 | 7.419 | 7.395 | 7.469 | 7.407 | 0.062 |
| E9PDE8 | Uncharacterized protein OS=Homo sapiens GN=HSPA4L PE=3 SV=1 - [E9PDE8_HUMAN] | 2.405E7 | 1.594E7 | 1.614E7 | 1.788E7 | 7.381 | 7.203 | 7.208 | 7.252 | 7.292 | 7.230 | 0.062 |
| B3KW71 | Quinoid dihydropteridine reductase, isoform CRA_e OS=Homo sapiens GN=QDPR PE=2 SV=1 - [B3KW71_HUMAN] | | 8.014E6 | 7.102E6 | 6.816E6 | | 6.904 | 6.851 | 6.834 | 6.904 | 6.842 | 0.061 |
| Q9UQ80 | Proliferation-associated protein 2G4 OS=Homo sapiens GN=PA2G4 PE=1 SV=3 - [PA2G4_HUMAN] | 9.168E7 | 1.119E8 | 8.580E7 | 9.022E7 | 7.962 | 8.049 | 7.934 | 7.955 | 8.006 | 7.944 | 0.061 |
| B4E049 | Uncharacterized protein OS=Homo sapiens GN=SEPT6 PE=2 SV=1 - [B4E049_HUMAN] | 1.237E7 | 7.920E6 | 7.128E6 | 1.038E7 | 7.093 | 6.899 | 6.853 | 7.016 | 6.996 | 6.935 | 0.061 |
| P46782 | 40S ribosomal protein S5 OS=Homo sapiens GN=RPS5 PE=1 SV=4 - [RS5_HUMAN] | 8.007E7 | 5.898E7 | 5.961E7 | 5.992E7 | 7.903 | 7.771 | 7.775 | 7.778 | 7.837 | 7.776 | 0.061 |
| F8W7K3 | Uncharacterized protein OS=Homo sapiens GN=SPTAN1 PE=4 SV=1 - [F8W7K3_HUMAN] | 9.680E6 | 7.849E6 | 7.770E6 | 7.405E6 | 6.986 | 6.895 | 6.890 | 6.869 | 6.940 | 6.880 | 0.060 |
| Q96AB3 | Isochorismatase domain-containing protein 2, mitochondrial OS=Homo sapiens GN=ISOC2 PE=1 SV=1 - [ISOC2_HUMAN] | | 1.236E7 | 1.139E7 | 1.018E7 | | 7.092 | 7.056 | 7.008 | 7.092 | 7.032 | 0.060 |
| B4DJ66 | Proteasome (Prosome, macropain) 26S subunit, non-ATPase, 13, isoform CRA_d OS=Homo sapiens GN=PSMI | 1.265E7 | 7.942E6 | 9.344E6 | 8.168E6 | 7.102 | 6.900 | 6.971 | 6.912 | 7.001 | 6.941 | 0.060 |
| P63313 | Thymosin beta-10 OS=Homo sapiens GN=TMSB10 PE=1 SV=2 - [TYB10_HUMAN] | 1.863E8 | 1.318E8 | 1.374E8 | 1.360E8 | 8.270 | 8.120 | 8.138 | 8.133 | 8.195 | 8.136 | 0.059 |
| P09669 | Cytochrome c oxidase subunit 6C OS=Homo sapiens GN=COX6C PE=1 SV=2 - [COX6C_HUMAN] | 1.043E7 | 1.329E7 | 9.840E6 | 1.074E7 | 7.018 | 7.123 | 6.993 | 7.031 | 7.071 | 7.012 | 0.059 |
| P07195 | L-lactate dehydrogenase B chain OS=Homo sapiens GN=LDHB PE=1 SV=2 - [LDHB_HUMAN] | 3.473E8 | 1.868E8 | 2.252E8 | 2.203E8 | 8.541 | 8.271 | 8.353 | 8.343 | 8.406 | 8.348 | 0.058 |
| E9PMD7 | Serine/threonine-protein phosphatase OS=Homo sapiens GN=PPP1CA PE=3 SV=1 - [E9PMD7_HUMAN] | 9.048E6 | 8.927E6 | 7.179E6 | 8.616E6 | 6.957 | 6.951 | 6.856 | 6.935 | 6.954 | 6.896 | 0.058 |
| P68036 | Ubiquitin-conjugating enzyme E2 L3 OS=Homo sapiens GN=UBE2L3 PE=1 SV=1 - [UB2L3_HUMAN] | 3.436E7 | 4.049E7 | 3.389E7 | 3.143E7 | 7.536 | 7.607 | 7.530 | 7.497 | 7.572 | 7.514 | 0.058 |
| Q8NAK9 | Uncharacterized protein OS=Homo sapiens GN=SRSF2 PE=2 SV=1 - [Q8NAK9_HUMAN] | 8.731E7 | 7.144E7 | 6.691E7 | 7.151E7 | 7.941 | 7.854 | 7.826 | 7.854 | 7.897 | 7.840 | 0.058 |
| P53396 | ATP-citrate synthase OS=Homo sapiens GN=ACLY PE=1 SV=3 - [ACLY_HUMAN] | 4.942E7 | 4.177E7 | 3.862E7 | 4.109E7 | 7.694 | 7.621 | 7.587 | 7.614 | 7.657 | 7.600 | 0.057 |
| Q13263 | Transcription intermediary factor 1-beta OS=Homo sapiens GN=TRIM28 PE=1 SV=5 - [TIF1B_HUMAN] | 1.372E8 | 1.594E8 | 1.299E8 | 1.295E8 | 8.137 | 8.202 | 8.113 | 8.112 | 8.170 | 8.113 | 0.057 |
| O43598 | Deoxyribonucleoside 5'-monophosphate N-glycosidase OS=Homo sapiens GN=RCL PE=1 SV=1 - [RCL_HUMA | 4.238E7 | 4.634E7 | 3.819E7 | 3.957E7 | 7.627 | 7.666 | 7.582 | 7.597 | 7.647 | 7.590 | 0.057 |
| P15311 | Ezrin OS=Homo sapiens GN=EZR PE=1 SV=4 - [EZRI_HUMAN] | 5.714E7 | 4.375E7 | 4.320E7 | 4.465E7 | 7.757 | 7.641 | 7.635 | 7.650 | 7.699 | 7.643 | 0.056 |
| Q92900-2 | Isoform 2 of Regulator of nonsense transcripts 1 OS=Homo sapiens GN=UPF1 - [RENT1_HUMAN] | 5.337E6 | 6.251E6 | 4.606E6 | 5.588E6 | 6.727 | 6.796 | 6.663 | 6.747 | 6.762 | 6.705 | 0.056 |
| Q92841-4 | Isoform 4 of Probable ATP-dependent RNA helicase DDX17 OS=Homo sapiens GN=DDX17 - [DDX17_HUMAN | 4.637E7 | 2.663E7 | 2.949E7 | 3.231E7 | 7.666 | 7.425 | 7.470 | 7.509 | 7.546 | 7.490 | 0.056 |
| F8WJN3 | Uncharacterized protein OS=Homo sapiens GN=CPSF6 PE=4 SV=1 - [F8WJN3_HUMAN] | 1.317E7 | 9.500E6 | 9.789E6 | 9.910E6 | 7.120 | 6.978 | 6.991 | 6.996 | 7.049 | 6.993 | 0.055 |
| Q9UNE7-2 | Isoform 2 of E3 ubiquitin-protein ligase CHIP OS=Homo sapiens GN=STUB1 - [CHIP_HUMAN] | 8.046E6 | 5.163E6 | 5.678E6 | 5.681E6 | 6.906 | 6.713 | 6.754 | 6.754 | 6.809 | 6.754 | 0.055 |
| P25788-2 | Isoform 2 of Proteasome subunit alpha type-3 OS=Homo sapiens GN=PSMA3 - [PSA3_HUMAN] | 3.073E7 | 2.132E7 | 2.244E7 | 2.273E7 | 7.488 | 7.329 | 7.351 | 7.357 | 7.408 | 7.354 | 0.054 |
| P16989-3 | Isoform 3 of DNA-binding protein A OS=Homo sapiens GN=CSDA - [DBPA_HUMAN] | 3.982E7 | 2.948E7 | 2.954E7 | 3.099E7 | 7.600 | 7.470 | 7.470 | 7.491 | 7.535 | 7.481 | 0.054 |
| P05387 | 60S acidic ribosomal protein P2 OS=Homo sapiens GN=RPLP2 PE=1 SV=1 - [RLA2_HUMAN] | 3.206E8 | 2.439E8 | 2.365E8 | 2.588E8 | 8.506 | 8.387 | 8.374 | 8.413 | 8.447 | 8.393 | 0.053 |
| Q7Z7K0 | COX assembly mitochondrial protein homolog OS=Homo sapiens GN=CMC1 PE=1 SV=1 - [COXAM_HUMAN] | 1.005E7 | 5.709E6 | 7.377E6 | 6.108E6 | 7.002 | 6.757 | 6.868 | 6.786 | 6.879 | 6.827 | 0.053 |
| P27635 | 60S ribosomal protein L10 OS=Homo sapiens GN=RPL10 PE=1 SV=4 - [RL10_HUMAN] | 6.276E7 | 4.456E7 | 5.149E7 | 4.265E7 | 7.798 | 7.649 | 7.712 | 7.630 | 7.723 | 7.671 | 0.053 |
| O15355 | Protein phosphatase 1G OS=Homo sapiens GN=PPM1G PE=1 SV=1 - [PPM1G_HUMAN] | 1.651E7 | 1.900E7 | 1.580E7 | 1.560E7 | 7.218 | 7.279 | 7.199 | 7.193 | 7.248 | 7.196 | 0.052 |
| G3V5X9 | Uncharacterized protein OS=Homo sapiens GN=SNX6 PE=4 SV=1 - [G3V5X9_HUMAN] | 1.013E7 | | 6.815E6 | 1.184E7 | 7.006 | | 6.833 | 7.073 | 7.006 | 6.953 | 0.052 |
| P25789 | Proteasome subunit alpha type-4 OS=Homo sapiens GN=PSMA4 PE=1 SV=1 - [PSA4_HUMAN] | 1.724E7 | 1.684E7 | 1.428E7 | 1.598E7 | 7.237 | 7.226 | 7.155 | 7.204 | 7.231 | 7.179 | 0.052 |
| Q14320 | Protein FAM50A OS=Homo sapiens GN=FAM50A PE=1 SV=2 - [FA50A_HUMAN] | 1.340E7 | 6.182E6 | 8.261E6 | 7.890E6 | 7.127 | 6.791 | 6.917 | 6.897 | 6.959 | 6.907 | 0.052 |
| C9JU56 | Uncharacterized protein OS=Homo sapiens GN=RPL31 PE=4 SV=1 - [C9JU56_HUMAN] | 1.040E8 | 6.887E7 | 7.465E7 | 7.559E7 | 8.017 | 7.838 | 7.873 | 7.878 | 7.928 | 7.876 | 0.052 |
| B4DFA2 | Uncharacterized protein OS=Homo sapiens GN=NSF PE=2 SV=1 - [B4DFA2_HUMAN] | | 4.105E6 | 4.467E6 | 2.972E6 | | 6.613 | 6.650 | 6.473 | 6.613 | 6.562 | 0.052 |
| O75937 | DnaJ homolog subfamily C member 8 OS=Homo sapiens GN=DNAJC8 PE=1 SV=2 - [DNJC8_HUMAN] | 5.622E7 | 3.112E7 | 3.823E7 | 3.629E7 | 7.750 | 7.493 | 7.582 | 7.560 | 7.621 | 7.571 | 0.050 |
| P28340 | DNA polymerase delta catalytic subunit OS=Homo sapiens GN=POLD1 PE=1 SV=2 - [DPOD1_HUMAN] | | 1.291E7 | 1.201E7 | 1.103E7 | | 7.111 | 7.079 | 7.043 | 7.111 | 7.061 | 0.050 |
| P51858 | Hepatoma-derived growth factor OS=Homo sapiens GN=HDGF PE=1 SV=1 - [HDGF_HUMAN] | 4.981E7 | 4.334E7 | 4.201E7 | 4.099E7 | 7.697 | 7.637 | 7.623 | 7.613 | 7.667 | 7.618 | 0.049 |
| P18124 | 60S ribosomal protein L7 OS=Homo sapiens GN=RPL7 PE=1 SV=1 - [RL7_HUMAN] | 1.213E8 | 8.292E7 | 9.458E7 | 8.494E7 | 8.084 | 7.919 | 7.976 | 7.929 | 8.001 | 7.952 | 0.049 |
| C9J5D1 | Uncharacterized protein OS=Homo sapiens GN=NAA50 PE=4 SV=1 - [C9J5D1_HUMAN] | 1.729E7 | 8.045E6 | 1.202E7 | 9.262E6 | 7.238 | 6.905 | 7.080 | 6.967 | 7.072 | 7.023 | 0.048 |
| O00154-6 | Isoform 6 of Cytosolic acyl coenzyme A thioester hydrolase OS=Homo sapiens GN=ACOT7 - [BACH_HUMAN] | 1.606E7 | 1.307E7 | 1.333E7 | 1.262E7 | 7.206 | 7.116 | 7.125 | 7.101 | 7.161 | 7.113 | 0.048 |
| Q96A49 | Synapse-associated protein 1 OS=Homo sapiens GN=SYAP1 PE=1 SV=1 - [SYAP1_HUMAN] | 1.334E7 | 8.182E6 | | 9.383E6 | 7.125 | 6.913 | | 6.972 | 7.019 | 6.972 | 0.047 |
| Q13561 | Dynactin subunit 2 OS=Homo sapiens GN=DCTN2 PE=1 SV=4 - [DCTN2_HUMAN] | 1.929E7 | 1.398E7 | 1.838E7 | 1.189E7 | 7.285 | 7.146 | 7.264 | 7.075 | 7.216 | 7.170 | 0.046 |
| P22234 | Multifunctional protein ADE2 OS=Homo sapiens GN=PAICS PE=1 SV=3 - [PUR6_HUMAN] | 1.263E8 | 1.216E8 | 1.150E8 | 1.084E8 | 8.102 | 8.085 | 8.061 | 8.035 | 8.093 | 8.048 | 0.045 |
| P62244 | 40S ribosomal protein S15a OS=Homo sapiens GN=RPS15A PE=1 SV=2 - [RS15A_HUMAN] | 3.243E7 | 4.023E7 | 3.373E7 | 3.149E7 | 7.511 | 7.605 | 7.528 | 7.498 | 7.558 | 7.513 | 0.045 |
| Q5T6W5 | Heterogeneous nuclear ribonucleoprotein K OS=Homo sapiens GN=HNRNPK PE=2 SV=1 - [Q5T6W5_HUMAN | 2.858E8 | 1.958E8 | 2.124E8 | 2.157E8 | 8.456 | 8.292 | 8.327 | 8.334 | 8.374 | 8.331 | 0.043 |
| Q9BQ67 | Glutamate-rich WD repeat-containing protein 1 OS=Homo sapiens GN=GRWD1 PE=1 SV=1 - [GRWD1_HUMAN] | | 8.805E6 | 7.403E6 | 8.612E6 | | 6.945 | 6.869 | 6.935 | 6.945 | 6.902 | 0.042 |
| P06132 | Uroporphyrinogen decarboxylase OS=Homo sapiens GN=UROD PE=1 SV=2 - [DCUP_HUMAN] | 1.362E7 | 1.149E7 | 1.075E7 | 1.198E7 | 7.134 | 7.060 | 7.031 | 7.078 | 7.097 | 7.055 | 0.042 |
| Q9Y4Y9 | U6 snRNA-associated Sm-like protein LSm5 OS=Homo sapiens GN=LSM5 PE=1 SV=3 - [LSM5_HUMAN] | 2.022E7 | 1.754E7 | | 1.710E7 | 7.306 | 7.244 | | 7.233 | 7.275 | 7.233 | 0.042 |
| A8MX47 | Uncharacterized protein OS=Homo sapiens GN=PEPD PE=2 SV=2 - [A8MX47_HUMAN] | 2.270E7 | 1.890E7 | 2.234E7 | 1.584E7 | 7.356 | 7.277 | 7.349 | 7.200 | 7.316 | 7.275 | 0.042 |
| Q9UQE7 | Structural maintenance of chromosomes protein 3 OS=Homo sapiens GN=SMC3 PE=1 SV=2 - [SMC3_HUMAI | 4.906E6 | | 4.668E6 | 4.276E6 | 6.691 | | 6.669 | 6.631 | 6.691 | 6.650 | 0.041 |
| Q16222-2 | Isoform AGX1 of UDP-N-acetylhexosamine pyrophosphorylase OS=Homo sapiens GN=UAP1 - [UAP1_HUMAN] | 8.882E6 | 8.501E6 | | 7.914E6 | 6.949 | 6.929 | | 6.898 | 6.939 | 6.898 | 0.041 |
| Q7L014 | Probable ATP-dependent RNA helicase DDX46 OS=Homo sapiens GN=DDX46 PE=1 SV=2 - [DDX46_HUMAN] | | 4.617E6 | 4.006E6 | 4.429E6 | | 6.664 | 6.603 | 6.646 | 6.664 | 6.625 | 0.040 |
| Q9HB71 | Calcyclin-binding protein OS=Homo sapiens GN=CACYBP PE=1 SV=2 - [CYBP_HUMAN] | 1.616E8 | 1.486E8 | 1.369E8 | 1.474E8 | 8.209 | 8.172 | 8.136 | 8.168 | 8.190 | 8.152 | 0.038 |
| Q16629-3 | Isoform 3 of Serine/arginine-rich splicing factor 7 OS=Homo sapiens GN=SRSF7 - [SRSF7_HUMAN] | 2.324E7 | 2.117E7 | 2.218E7 | 1.864E7 | 7.366 | 7.326 | 7.346 | 7.271 | 7.346 | 7.308 | 0.038 |
| O00273 | DNA fragmentation factor subunit alpha OS=Homo sapiens GN=DFFA PE=1 SV=1 - [DFFA_HUMAN] | 1.516E7 | 1.328E7 | 1.312E7 | 1.297E7 | 7.181 | 7.123 | 7.118 | 7.113 | 7.152 | 7.116 | 0.037 |
| Q96A72 | Protein mago nashi homolog 2 OS=Homo sapiens GN=MAGOHB PE=1 SV=1 - [MGN2_HUMAN] | 1.556E7 | 9.416E6 | 1.120E7 | 1.106E7 | 7.192 | 6.974 | 7.049 | 7.044 | 7.083 | 7.047 | 0.036 |
| Q9H0D6-2 | Isoform 2 of 5'-3' exoribonuclease 2 OS=Homo sapiens GN=XRN2 - [XRN2_HUMAN] | 1.456E7 | 1.155E7 | 1.304E7 | 1.091E7 | 7.163 | 7.062 | 7.115 | 7.038 | 7.113 | 7.077 | 0.036 |
| Q99832 | T-complex protein 1 subunit eta OS=Homo sapiens GN=CCT7 PE=1 SV=2 - [TCPH_HUMAN] | 5.172E7 | 3.916E7 | 4.115E7 | 4.220E7 | 7.714 | 7.593 | 7.614 | 7.625 | 7.653 | 7.620 | 0.033 |
| Q01518 | Adenylyl cyclase-associated protein 1 OS=Homo sapiens GN=CAP1 PE=1 SV=5 - [CAP1_HUMAN] | 2.182E7 | 2.015E7 | 2.349E7 | 1.606E7 | 7.339 | 7.304 | 7.371 | 7.206 | 7.322 | 7.288 | 0.033 |
| Q12905 | Interleukin enhancer-binding factor 2 OS=Homo sapiens GN=ILF2 PE=1 SV=2 - [ILF2_HUMAN] | 5.305E7 | 4.031E7 | 4.263E7 | 4.321E7 | 7.725 | 7.605 | 7.630 | 7.636 | 7.665 | 7.633 | 0.032 |
| E9PL22 | Uncharacterized protein OS=Homo sapiens GN=HYOU1 PE=3 SV=1 - [E9PL22_HUMAN] | 1.766E7 | 1.579E7 | 1.458E7 | 1.651E7 | 7.247 | 7.198 | 7.164 | 7.218 | 7.223 | 7.191 | 0.032 |
| Q96P70 | Importin-9 OS=Homo sapiens GN=IPO9 PE=1 SV=3 - [IPO9_HUMAN] | 1.786E7 | 1.415E7 | 1.192E7 | 1.835E7 | 7.252 | 7.151 | 7.076 | 7.264 | 7.201 | 7.170 | 0.031 |
| B4E363 | Uncharacterized protein OS=Homo sapiens GN=FARSA PE=2 SV=1 - [B4E363_HUMAN] | 2.407E7 | 1.322E7 | 1.673E7 | 1.648E7 | 7.381 | 7.121 | 7.224 | 7.217 | 7.251 | 7.220 | 0.031 |
| F8W0A9 | Uncharacterized protein OS=Homo sapiens GN=ALDH2 PE=3 SV=1 - [F8W0A9_HUMAN] | 1.309E7 | 3.209E6 | 6.015E6 | 6.056E6 | 7.117 | 6.506 | 6.779 | 6.782 | 6.812 | 6.781 | 0.031 |
| P84090 | Enhancer of rudimentary homolog OS=Homo sapiens GN=ERH PE=1 SV=1 - [ERH_HUMAN] | 6.148E7 | 4.980E7 | 4.917E7 | 5.400E7 | 7.789 | 7.697 | 7.692 | 7.732 | 7.743 | 7.712 | 0.031 |
| O75822 | Eukaryotic translation initiation factor 3 subunit J OS=Homo sapiens GN=EIF3J PE=1 SV=2 - [EIF3J_HUMAN] | 4.705E7 | 1.000E7 | 1.188E7 | 3.442E7 | 7.673 | 7.000 | 7.075 | 7.537 | 7.336 | 7.306 | 0.030 |
| O43707 | Alpha-actinin-4 OS=Homo sapiens GN=ACTN4 PE=1 SV=2 - [ACTN4_HUMAN] | 3.266E7 | 2.778E7 | 2.784E7 | 2.835E7 | 7.514 | 7.444 | 7.445 | 7.453 | 7.479 | 7.449 | 0.030 |
| P32969 | 60S ribosomal protein L9 OS=Homo sapiens GN=RPL9 PE=1 SV=1 - [RL9_HUMAN] | 8.706E7 | 6.365E7 | 6.870E7 | 7.044E7 | 7.940 | 7.804 | 7.837 | 7.848 | 7.872 | 7.842 | 0.029 |
| P49327 | Fatty acid synthase OS=Homo sapiens GN=FASN PE=1 SV=3 - [FAS_HUMAN] | 7.355E7 | 5.824E7 | 6.062E7 | 6.177E7 | 7.867 | 7.765 | 7.783 | 7.791 | 7.816 | 7.787 | 0.029 |
| P63165 | Small ubiquitin-related modifier 1 OS=Homo sapiens GN=SUMO1 PE=1 SV=1 - [SUMO1_HUMAN] | 1.557E7 | 8.828E6 | 9.413E6 | 1.278E7 | 7.192 | 6.946 | 6.974 | 7.107 | 7.069 | 7.040 | 0.029 |
| Q8WVM8 | Sec1 family domain-containing protein 1 OS=Homo sapiens GN=SCFD1 PE=1 SV=4 - [SCFD1_HUMAN] | 1.127E7 | 1.039E7 | 8.676E6 | 1.183E7 | 7.052 | 7.017 | 6.938 | 7.073 | 7.034 | 7.006 | 0.029 |
| Q13151 | Heterogeneous nuclear ribonucleoprotein A0 OS=Homo sapiens GN=HNRNPA0 PE=1 SV=1 - [ROA0_HUMAN] | 2.219E7 | 2.483E7 | 2.682E7 | 1.808E7 | 7.346 | 7.395 | 7.428 | 7.257 | 7.371 | 7.343 | 0.028 |
| G3V153 | GPI-anchored membrane protein 1, isoform CRA_d OS=Homo sapiens GN=CAPRIN1 PE=4 SV=1 - [G3V153_I | 2.109E7 | 1.598E7 | 1.896E7 | 1.567E7 | 7.324 | 7.204 | 7.278 | 7.195 | 7.264 | 7.236 | 0.027 |
| O75533 | Splicing factor 3B subunit 1 OS=Homo sapiens GN=SF3B1 PE=1 SV=3 - [SF3B1_HUMAN] | 1.323E7 | 1.330E7 | 1.272E7 | 1.220E7 | 7.122 | 7.124 | 7.104 | 7.087 | 7.123 | 7.095 | 0.027 |
| F5H6K0 | Uncharacterized protein OS=Homo sapiens GN=DHX15 PE=4 SV=1 - [F5H6K0_HUMAN] | 5.680E6 | 6.284E6 | 6.012E6 | 5.236E6 | 6.754 | 6.798 | 6.779 | 6.719 | 6.776 | 6.749 | 0.027 |
| Q9UBT2 | SUMO-activating enzyme subunit 2 OS=Homo sapiens GN=UBA2 PE=1 SV=2 - [SAE2_HUMAN] | 3.548E7 | 7.711E6 | 1.826E7 | 1.335E7 | 7.550 | 6.887 | 7.262 | 7.126 | 7.219 | 7.194 | 0.025 |
| C9J4Z3 | Ribosomal protein L37a, isoform CRA_c OS=Homo sapiens GN=RPL37A PE=4 SV=1 - [C9J4Z3_HUMAN] | 3.798E7 | 3.062E7 | 2.728E7 | 3.801E7 | 7.580 | 7.486 | 7.436 | 7.580 | 7.533 | 7.508 | 0.025 |
| O43464-2 | Isoform 2 of Serine protease HTRA2, mitochondrial OS=Homo sapiens GN=HTRA2 - [HTRA2_HUMAN] | 6.324E6 | | 5.993E6 | | 6.801 | | 6.778 | | 6.801 | 6.778 | 0.023 |

| | | | | | | | | | | | | |
|---|---|---|---|---|---|---|---|---|---|---|---|---|
| P31153 | S-adenosylmethionine synthase isoform type-2 OS=Homo sapiens GN=MAT2A PE=1 SV=1 - [METK2_HUMAN | 2.370E7 | 2.161E7 | 2.001E7 | 2.323E7 | 7.375 | 7.335 | 7.301 | 7.366 | 7.355 | 7.334 | 0.021 |
| E7ERW8 | Uncharacterized protein OS=Homo sapiens GN=DIAPH1 PE=4 SV=1 - [E7ERW8_HUMAN] | 4.619E6 | 3.780E6 | 3.867E6 | 4.123E6 | 6.665 | 6.577 | 6.587 | 6.615 | 6.621 | 6.601 | 0.020 |
| B4DJP7 | Uncharacterized protein OS=Homo sapiens GN=SNRPD3 PE=2 SV=1 - [B4DJP7_HUMAN] | 1.170E8 | 7.927E7 | 9.980E7 | 8.507E7 | 8.068 | 7.899 | 7.999 | 7.930 | 7.984 | 7.964 | 0.019 |
| P13797 | Plastin-3 OS=Homo sapiens GN=PLS3 PE=1 SV=4 - [PLST_HUMAN] | 4.447E7 | 5.307E7 | 4.739E7 | 4.567E7 | 7.648 | 7.725 | 7.676 | 7.660 | 7.686 | 7.668 | 0.019 |
| Q9NUU7 | ATP-dependent RNA helicase DDX19A OS=Homo sapiens GN=DDX19A PE=1 SV=1 - [DD19A_HUMAN] | 1.737E7 | 1.925E7 | 1.720E7 | 1.783E7 | 7.240 | 7.284 | 7.236 | 7.251 | 7.262 | 7.243 | 0.019 |
| Q9BS26 | Endoplasmic reticulum resident protein 44 OS=Homo sapiens GN=ERP44 PE=1 SV=1 - [ERP44_HUMAN] | | 8.061E6 | 8.194E6 | 7.281E6 | | 6.906 | 6.913 | 6.862 | 6.906 | 6.888 | 0.019 |
| Q49AN9 | SNRPG protein OS=Homo sapiens GN=SNRPG PE=4 SV=1 - [Q49AN9_HUMAN] | 4.339E7 | 4.875E7 | 4.514E7 | 4.394E7 | 7.637 | 7.688 | 7.655 | 7.643 | 7.663 | 7.649 | 0.014 |
| C9J9K3 | Uncharacterized protein OS=Homo sapiens GN=RPSA PE=3 SV=1 - [C9J9K3_HUMAN] | 3.380E8 | 2.495E8 | 2.643E8 | 3.015E8 | 8.529 | 8.397 | 8.422 | 8.479 | 8.463 | 8.451 | 0.012 |
| E9PDQ8 | Uncharacterized protein OS=Homo sapiens GN=SUCLG2 PE=4 SV=1 - [E9PDQ8_HUMAN] | 7.597E6 | 4.672E6 | | 5.798E6 | 6.881 | 6.670 | | 6.763 | 6.775 | 6.763 | 0.012 |
| Q9UMS4 | Pre-mRNA-processing factor 19 OS=Homo sapiens GN=PRPF19 PE=1 SV=1 - [PRP19_HUMAN] | 2.702E7 | 4.501E7 | 3.483E7 | 3.308E7 | 7.432 | 7.653 | 7.542 | 7.520 | 7.543 | 7.531 | 0.012 |
| B4DFL2 | Isocitrate dehydrogenase [NADP] OS=Homo sapiens GN=IDH2 PE=2 SV=1 - [B4DFL2_HUMAN] | 5.385E6 | 9.763E6 | 4.808E6 | 1.037E7 | 6.731 | 6.990 | 6.682 | 7.016 | 6.860 | 6.849 | 0.012 |
| P26038 | Moesin OS=Homo sapiens GN=MSN PE=1 SV=3 - [MOES_HUMAN] | 5.276E7 | 3.800E7 | 4.320E7 | 4.413E7 | 7.722 | 7.580 | 7.635 | 7.645 | 7.651 | 7.640 | 0.011 |
| P35241 | Radixin OS=Homo sapiens GN=RDX PE=1 SV=1 - [RADI_HUMAN] | 5.276E7 | 3.800E7 | 4.320E7 | 4.413E7 | 7.722 | 7.580 | 7.635 | 7.645 | 7.651 | 7.640 | 0.011 |
| P19338 | Nucleolin OS=Homo sapiens GN=NCL PE=1 SV=3 - [NUCL_HUMAN] | 4.382E8 | 3.313E8 | 3.702E8 | 3.745E8 | 8.642 | 8.520 | 8.568 | 8.573 | 8.581 | 8.571 | 0.010 |
| B7WP09 | Target of myb1 (Chicken) OS=Homo sapiens GN=TOM1 PE=4 SV=2 - [B7WP09_HUMAN] | | 3.535E6 | 3.223E6 | 3.710E6 | | 6.548 | 6.508 | 6.569 | 6.548 | 6.539 | 0.010 |
| Q92616 | Translational activator GCN1 OS=Homo sapiens GN=GCN1L1 PE=1 SV=6 - [GCN1L_HUMAN] | 1.064E7 | 6.773E6 | 6.658E6 | 1.037E7 | 7.027 | 6.831 | 6.823 | 7.016 | 6.929 | 6.920 | 0.009 |
| E7EQZ3 | Uncharacterized protein OS=Homo sapiens GN=GSPT1 PE=4 SV=1 - [E7EQZ3_HUMAN] | 1.166E7 | 1.729E7 | 1.261E7 | 1.534E7 | 7.067 | 7.238 | 7.101 | 7.186 | 7.152 | 7.143 | 0.009 |
| Q9H444 | Charged multivesicular body protein 4b OS=Homo sapiens GN=CHMP4B PE=1 SV=1 - [CHM4B_HUMAN] | 1.512E7 | 8.413E6 | 1.099E7 | 1.113E7 | 7.180 | 6.925 | 7.041 | 7.047 | 7.052 | 7.044 | 0.008 |
| Q04837 | Single-stranded DNA-binding protein, mitochondrial OS=Homo sapiens GN=SSBP1 PE=1 SV=1 - [SSBP_HUM/ | 1.079E8 | 8.308E7 | 9.188E7 | 9.387E7 | 8.033 | 7.920 | 7.963 | 7.973 | 7.976 | 7.968 | 0.008 |
| P98179 | Putative RNA-binding protein 3 OS=Homo sapiens GN=RBM3 PE=1 SV=1 - [RBM3_HUMAN] | 4.090E7 | 2.234E7 | 2.947E7 | 2.986E7 | 7.612 | 7.349 | 7.469 | 7.475 | 7.480 | 7.472 | 0.008 |
| O00571 | ATP-dependent RNA helicase DDX3X OS=Homo sapiens GN=DDX3X PE=1 SV=3 - [DDX3X_HUMAN] | 2.310E7 | 2.652E7 | 2.346E7 | 2.518E7 | 7.364 | 7.423 | 7.370 | 7.401 | 7.394 | 7.386 | 0.008 |
| Q96K98 | Uncharacterized protein OS=Homo sapiens GN=SRP68 PE=2 SV=1 - [Q96K98_HUMAN] | 1.371E7 | 1.275E7 | 1.717E7 | 9.832E6 | 7.137 | 7.105 | 7.235 | 6.993 | 7.121 | 7.114 | 0.008 |
| B7Z2D6 | Uncharacterized protein OS=Homo sapiens GN=CBS PE=2 SV=1 - [B7Z2D6_HUMAN] | 2.015E7 | 1.721E7 | 1.902E7 | 1.761E7 | 7.304 | 7.236 | 7.279 | 7.246 | 7.270 | 7.263 | 0.008 |
| P37837 | Transaldolase OS=Homo sapiens GN=TALDO1 PE=1 SV=2 - [TALDO_HUMAN] | 3.687E7 | 1.462E7 | 2.523E7 | 2.076E7 | 7.567 | 7.165 | 7.402 | 7.317 | 7.366 | 7.360 | 0.006 |
| B4E2M8 | Uncharacterized protein OS=Homo sapiens GN=FAF2 PE=2 SV=1 - [B4E2M8_HUMAN] | | 1.122E7 | 9.064E6 | 1.356E7 | | 7.050 | 6.957 | 7.132 | 7.050 | 7.045 | 0.005 |
| O95881 | Thioredoxin domain-containing protein 12 OS=Homo sapiens GN=TXNDC12 PE=1 SV=1 - [TXD12_HUMAN] | 1.395E7 | 1.343E7 | | 1.353E7 | 7.145 | 7.128 | | 7.131 | 7.136 | 7.131 | 0.005 |
| Q9NYU2-2 | Isoform 2 of UDP-glucose:glycoprotein glucosyltransferase 1 OS=Homo sapiens GN=UGGT1 - [UGGG1_HUMAN] | | 9.217E6 | 8.533E6 | 9.753E6 | | 6.965 | 6.931 | 6.989 | 6.965 | 6.960 | 0.004 |
| P49419-2 | Isoform 2 of Alpha-aminoadipic semialdehyde dehydrogenase OS=Homo sapiens GN=ALDH7A1 - [AL7A1_HU | 1.107E7 | 2.602E7 | 2.361E7 | 1.202E7 | 7.044 | 7.415 | 7.373 | 7.080 | 7.230 | 7.227 | 0.003 |
| E9PGT1 | Uncharacterized protein OS=Homo sapiens GN=TSN PE=4 SV=1 - [E9PGT1_HUMAN] | 8.125E6 | 8.914E6 | 5.556E6 | 1.286E7 | 6.910 | 6.950 | 6.745 | 7.109 | 6.930 | 6.927 | 0.003 |
| Q14152 | Eukaryotic translation initiation factor 3 subunit A OS=Homo sapiens GN=EIF3A PE=1 SV=1 - [EIF3A_HUMAN | 2.458E7 | 3.006E7 | 2.823E7 | 2.587E7 | 7.391 | 7.478 | 7.451 | 7.413 | 7.434 | 7.432 | 0.002 |
| Q00325-2 | Isoform B of Phosphate carrier protein, mitochondrial OS=Homo sapiens GN=SLC25A3 - [MPCP_HUMAN] | 4.013E7 | 2.602E7 | 2.577E7 | 4.014E7 | 7.603 | 7.415 | 7.411 | 7.604 | 7.509 | 7.507 | 0.002 |
| Q04760-2 | Isoform 2 of Lactoylglutathione lyase OS=Homo sapiens GN=GLO1 - [LGUL_HUMAN] | 1.170E8 | 5.249E7 | 5.707E7 | 1.069E8 | 8.068 | 7.720 | 7.756 | 8.029 | 7.894 | 7.893 | 0.001 |
| Q12906-4 | Isoform 4 of Interleukin enhancer-binding factor 3 OS=Homo sapiens GN=ILF3 - [ILF3_HUMAN] | 6.980E7 | 4.047E7 | 4.977E7 | 5.664E7 | 7.844 | 7.607 | 7.697 | 7.753 | 7.725 | 7.725 | 0.000 |
| Q9BY44-2 | Isoform 2 of Eukaryotic translation initiation factor 2A OS=Homo sapiens GN=EIF2A - [EIF2A_HUMAN] | | 5.301E6 | 5.637E6 | 4.996E6 | | 6.724 | 6.751 | 6.699 | 6.724 | 6.725 | -0.001 |
| O43252 | Bifunctional 3'-phosphoadenosine 5'-phosphosulfate synthase 1 OS=Homo sapiens GN=PAPSS1 PE=1 SV=2 - | 1.069E7 | 7.355E6 | 8.596E6 | 9.189E6 | 7.029 | 6.867 | 6.934 | 6.963 | 6.948 | 6.949 | -0.001 |
| P67870 | Casein kinase II subunit beta OS=Homo sapiens GN=CSNK2B PE=1 SV=1 - [CSK2B_HUMAN] | 1.116E7 | 1.391E7 | 1.129E7 | 1.382E7 | 7.048 | 7.143 | 7.053 | 7.140 | 7.095 | 7.097 | -0.001 |
| O60271-7 | Isoform 7 of C-Jun-amino-terminal kinase-interacting protein 4 OS=Homo sapiens GN=SPAG9 - [JIP4_HUMAN | 2.337E6 | 2.496E6 | | 2.431E6 | 6.369 | 6.397 | | 6.386 | 6.383 | 6.386 | -0.003 |
| P25786 | Proteasome subunit alpha type-1 OS=Homo sapiens GN=PSMA1 PE=1 SV=1 - [PSA1_HUMAN] | 3.910E7 | 3.507E7 | 3.682E7 | 3.794E7 | 7.592 | 7.545 | 7.566 | 7.579 | 7.569 | 7.573 | -0.004 |
| O60506-3 | Isoform 3 of Heterogeneous nuclear ribonucleoprotein Q OS=Homo sapiens GN=SYNCRIP - [HNRPQ_HUMAN | 6.057E7 | 3.816E7 | 4.407E7 | 5.362E7 | 7.782 | 7.582 | 7.644 | 7.729 | 7.682 | 7.687 | -0.005 |
| P10599 | Thioredoxin OS=Homo sapiens GN=TXN PE=1 SV=3 - [THIO_HUMAN] | 3.811E7 | 2.377E7 | 2.513E7 | 3.687E7 | 7.581 | 7.376 | 7.400 | 7.567 | 7.479 | 7.483 | -0.005 |
| Q6FI81 | Anamorsin OS=Homo sapiens GN=CIAPIN1 PE=1 SV=2 - [CPIN1_HUMAN] | 1.493E7 | 1.116E7 | 1.345E7 | 1.268E7 | 7.174 | 7.048 | 7.129 | 7.103 | 7.111 | 7.116 | -0.005 |
| P61758 | Prefoldin subunit 3 OS=Homo sapiens GN=VBP1 PE=1 SV=3 - [PFD3_HUMAN] | 2.669E7 | 1.990E7 | 2.253E7 | 2.415E7 | 7.426 | 7.299 | 7.353 | 7.383 | 7.363 | 7.368 | -0.005 |
| P04181 | Ornithine aminotransferase, mitochondrial OS=Homo sapiens GN=OAT PE=1 SV=1 - [OAT_HUMAN] | 3.761E7 | 3.469E7 | 3.180E7 | 4.207E7 | 7.575 | 7.540 | 7.502 | 7.624 | 7.558 | 7.563 | -0.005 |
| P53999 | Activated RNA polymerase II transcriptional coactivator p15 OS=Homo sapiens GN=SUB1 PE=1 SV=3 - [TCP4 | 3.340E7 | 1.371E8 | 7.144E7 | 6.575E7 | 7.524 | 8.137 | 7.854 | 7.818 | 7.830 | 7.836 | -0.006 |
| D6RFH4 | Uncharacterized protein OS=Homo sapiens GN=CYB5B PE=3 SV=1 - [D6RFH4_HUMAN] | 3.367E7 | 3.279E7 | 3.270E7 | 3.477E7 | 7.527 | 7.516 | 7.515 | 7.541 | 7.521 | 7.528 | -0.006 |
| F8W9D6 | Adenylosuccinate synthetase OS=Homo sapiens GN=ADSS PE=3 SV=1 - [F8W9D6_HUMAN] | | 1.428E7 | 1.440E7 | 1.462E7 | | 7.155 | 7.159 | 7.165 | 7.155 | 7.162 | -0.007 |
| O15116 | U6 snRNA-associated Sm-like protein LSm1 OS=Homo sapiens GN=LSM1 PE=1 SV=1 - [LSM1_HUMAN] | 4.797E6 | 7.637E6 | | 6.167E6 | 6.681 | 6.883 | | 6.790 | 6.782 | 6.790 | -0.008 |
| P10606 | Cytochrome c oxidase subunit 5B, mitochondrial OS=Homo sapiens GN=COX5B PE=1 SV=2 - [COX5B_HUMA | 4.252E7 | 4.214E7 | | 4.319E7 | 7.629 | 7.625 | | 7.635 | 7.627 | 7.635 | -0.009 |
| Q13867 | Bleomycin hydrolase OS=Homo sapiens GN=BLMH PE=1 SV=1 - [BLMH_HUMAN] | 1.623E7 | 1.124E7 | 1.523E7 | 1.254E7 | 7.210 | 7.051 | 7.183 | 7.098 | 7.131 | 7.141 | -0.010 |
| P62312 | U6 snRNA-associated Sm-like protein LSm6 OS=Homo sapiens GN=LSM6 PE=1 SV=1 - [LSM6_HUMAN] | 6.125E6 | 5.266E6 | | 5.823E6 | 6.787 | 6.721 | | 6.765 | 6.754 | 6.765 | -0.011 |
| B4DM74 | Uncharacterized protein OS=Homo sapiens GN=RPL18A PE=2 SV=1 - [B4DM74_HUMAN] | 1.773E7 | 1.193E7 | 1.525E7 | 1.460E7 | 7.249 | 7.077 | 7.183 | 7.164 | 7.163 | 7.174 | -0.011 |
| Q9NRR5 | Ubiquilin-4 OS=Homo sapiens GN=UBQLN4 PE=1 SV=2 - [UBQL4_HUMAN] | 8.439E6 | 1.502E7 | 1.112E7 | 1.207E7 | 6.926 | 7.177 | 7.046 | 7.082 | 7.052 | 7.064 | -0.013 |
| P62857 | 40S ribosomal protein S28 OS=Homo sapiens GN=RPS28 PE=1 SV=1 - [RS28_HUMAN] | 4.589E7 | 3.500E7 | 3.911E7 | 4.372E7 | 7.662 | 7.544 | 7.592 | 7.641 | 7.603 | 7.616 | -0.014 |
| A8K318 | Uncharacterized protein OS=Homo sapiens GN=PRKCSH PE=2 SV=1 - [A8K318_HUMAN] | 4.891E7 | 3.849E7 | 4.636E7 | 4.344E7 | 7.689 | 7.585 | 7.666 | 7.638 | 7.637 | 7.652 | -0.015 |
| C9JWK3 | Uncharacterized protein OS=Homo sapiens GN=HARS PE=4 SV=1 - [C9JWK3_HUMAN] | 1.378E7 | 1.448E7 | 1.450E7 | 1.472E7 | 7.139 | 7.161 | 7.161 | 7.168 | 7.150 | 7.165 | -0.015 |
| A2ABK4 | RD RNA binding protein OS=Homo sapiens GN=RDBP PE=4 SV=1 - [A2ABK4_HUMAN] | 6.635E6 | 2.262E6 | 4.017E6 | | 6.822 | 6.355 | 6.604 | | 6.588 | 6.604 | -0.016 |
| P62310 | U6 snRNA-associated Sm-like protein LSm3 OS=Homo sapiens GN=LSM3 PE=1 SV=2 - [LSM3_HUMAN] | 1.421E7 | 1.792E7 | 1.635E7 | 1.675E7 | 7.152 | 7.253 | 7.213 | 7.224 | 7.203 | 7.219 | -0.016 |
| F5GZ76 | Uncharacterized protein OS=Homo sapiens GN=USO1 PE=4 SV=1 - [F5GZ76_HUMAN] | 6.165E6 | 3.071E6 | 4.891E6 | 4.162E6 | 6.790 | 6.487 | 6.689 | 6.619 | 6.639 | 6.654 | -0.016 |
| Q8WVC2 | 40S ribosomal protein S21 OS=Homo sapiens GN=RPS21 PE=2 SV=1 - [Q8WVC2_HUMAN] | 9.438E7 | 8.091E7 | 9.011E7 | 9.152E7 | 7.975 | 7.908 | 7.955 | 7.962 | 7.941 | 7.958 | -0.017 |
| P62277 | 40S ribosomal protein S13 OS=Homo sapiens GN=RPS13 PE=1 SV=2 - [RS13_HUMAN] | 7.328E7 | 6.116E7 | 6.651E7 | 7.277E7 | 7.865 | 7.786 | 7.823 | 7.862 | 7.826 | 7.842 | -0.017 |
| O75534-2 | Isoform Short of Cold shock domain-containing protein E1 OS=Homo sapiens GN=CSDE1 - [CSDE1_HUMAN] | 1.077E7 | 5.433E6 | 8.750E6 | 7.242E6 | 7.032 | 6.735 | 6.942 | 6.860 | 6.884 | 6.901 | -0.017 |
| P48637 | Glutathione synthetase OS=Homo sapiens GN=GSS PE=1 SV=1 - [GSHB_HUMAN] | 6.947E6 | | 7.419E6 | 7.069E6 | 6.842 | | 6.870 | 6.849 | 6.842 | 6.860 | -0.018 |
| D6RD66 | Uncharacterized protein OS=Homo sapiens GN=WDR1 PE=4 SV=1 - [D6RD66_HUMAN] | | 6.059E6 | 4.933E6 | 8.131E6 | | 6.782 | 6.693 | 6.910 | 6.782 | 6.802 | -0.019 |
| P08779 | Keratin, type I cytoskeletal 16 OS=Homo sapiens GN=KRT16 PE=1 SV=4 - [K1C16_HUMAN] | 6.871E7 | 5.215E7 | 6.273E7 | 6.293E7 | 7.837 | 7.717 | 7.797 | 7.799 | 7.777 | 7.798 | -0.021 |
| Q08211 | ATP-dependent RNA helicase A OS=Homo sapiens GN=DHX9 PE=1 SV=4 - [DHX9_HUMAN] | 1.943E7 | 1.304E7 | 1.418E7 | 1.974E7 | 7.288 | 7.115 | 7.152 | 7.295 | 7.202 | 7.223 | -0.022 |
| Q96HS1-2 | Isoform 2 of Serine/threonine-protein phosphatase PGAM5, mitochondrial OS=Homo sapiens GN=PGAM5 - [F | 5.547E6 | | | 5.832E6 | 6.744 | | | 6.766 | 6.744 | 6.766 | -0.022 |
| P05386 | 60S acidic ribosomal protein P1 OS=Homo sapiens GN=RPLP1 PE=1 SV=1 - [RLA1_HUMAN] | 3.619E8 | 2.749E8 | 2.758E8 | 3.996E8 | 8.559 | 8.439 | 8.441 | 8.602 | 8.499 | 8.521 | -0.022 |
| Q99460-2 | Isoform 2 of 26S proteasome non-ATPase regulatory subunit 1 OS=Homo sapiens GN=PSMD1 - [PSMD1_HUMAN] | | 9.567E6 | 8.568E6 | 1.190E7 | | 6.981 | 6.933 | 7.075 | 6.981 | 7.004 | -0.023 |
| Q9HDC9-2 | Isoform 2 of Adipocyte plasma membrane-associated protein OS=Homo sapiens GN=APMAP - [APMAP_HUM/ | 1.426E7 | 1.542E7 | 1.184E7 | 2.073E7 | 7.154 | 7.188 | 7.073 | 7.317 | 7.171 | 7.195 | -0.024 |
| Q92598-2 | Isoform Beta of Heat shock protein 105 kDa OS=Homo sapiens GN=HSPH1 - [HS105_HUMAN] | 1.780E7 | 2.391E7 | 2.119E7 | 2.256E7 | 7.250 | 7.379 | 7.326 | 7.353 | 7.314 | 7.340 | -0.025 |
| G3V4F7 | Uncharacterized protein OS=Homo sapiens GN=SRP54 PE=4 SV=1 - [G3V4F7_HUMAN] | 6.553E6 | 9.036E6 | 8.493E6 | 7.890E6 | 6.816 | 6.956 | 6.929 | 6.897 | 6.886 | 6.913 | -0.027 |
| P84243 | Histone H3.3 OS=Homo sapiens GN=H3F3A PE=1 SV=2 - [H33_HUMAN] | 9.619E6 | 1.092E7 | | 1.090E7 | 6.983 | 7.038 | | 7.038 | 7.011 | 7.038 | -0.027 |
| F8WD96 | Uncharacterized protein OS=Homo sapiens GN=CTSD PE=3 SV=1 - [F8WD96_HUMAN] | 1.288E7 | 1.693E7 | 1.415E7 | 1.745E7 | 7.110 | 7.229 | 7.151 | 7.242 | 7.169 | 7.196 | -0.027 |
| P78417 | Glutathione S-transferase omega-1 OS=Homo sapiens GN=GSTO1 PE=1 SV=2 - [GSTO1_HUMAN] | 1.161E7 | 6.265E6 | | 9.081E6 | 7.065 | 6.797 | | 6.958 | 6.931 | 6.958 | -0.027 |
| P62942 | Peptidyl-prolyl cis-trans isomerase FKBP1A OS=Homo sapiens GN=FKBP1A PE=1 SV=2 - [FKB1A_HUMAN] | 7.280E7 | 4.440E7 | 4.784E7 | 7.667E7 | 7.862 | 7.647 | 7.680 | 7.885 | 7.755 | 7.782 | -0.027 |
| G3V1K1 | Coiled-coil-helix-coiled-coil-helix domain containing 3, isoform CRA_b OS=Homo sapiens GN=CHCHD3 PE=4 S | 7.639E6 | 6.401E6 | 6.856E6 | 8.102E6 | 6.883 | 6.806 | 6.836 | 6.909 | 6.845 | 6.872 | -0.028 |
| Q8WW12-2 | Isoform 2 of PEST proteolytic signal-containing nuclear protein OS=Homo sapiens GN=PCNP - [PCNP_HUMAN | 1.167E7 | 1.054E7 | 1.108E7 | 1.262E7 | 7.067 | 7.023 | 7.045 | 7.101 | 7.045 | 7.073 | -0.028 |
| P14927 | Cytochrome b-c1 complex subunit 7 OS=Homo sapiens GN=UQCRB PE=1 SV=2 - [QCR7_HUMAN] | 5.084E7 | 3.507E7 | 3.478E7 | 5.848E7 | 7.706 | 7.545 | 7.541 | 7.767 | 7.626 | 7.654 | -0.029 |
| Q5T4L4 | Ribosomal protein S27 OS=Homo sapiens GN=RPS27 PE=4 SV=1 - [Q5T4L4_HUMAN] | 6.436E7 | 2.704E7 | 4.445E7 | 4.485E7 | 7.809 | 7.432 | 7.648 | 7.652 | 7.620 | 7.650 | -0.030 |
| B0V043 | Uncharacterized protein OS=Homo sapiens GN=VARS PE=3 SV=1 - [B0V043_HUMAN] | 2.048E7 | 1.431E7 | 2.277E7 | 1.482E7 | 7.311 | 7.156 | 7.357 | 7.171 | 7.234 | 7.264 | -0.031 |
| E1B6W5 | Uncharacterized protein OS=Homo sapiens GN=SCP2 PE=4 SV=1 - [E1B6W5_HUMAN] | 1.759E7 | 1.078E7 | 1.585E7 | 1.378E7 | 7.245 | 7.033 | 7.200 | 7.139 | 7.139 | 7.170 | -0.031 |
| O43399-2 | Isoform 2 of Tumor protein D54 OS=Homo sapiens GN=TPD52L2 - [TPD54_HUMAN] | 2.822E7 | 1.287E7 | 2.152E7 | 1.947E7 | 7.451 | 7.110 | 7.333 | 7.289 | 7.280 | 7.311 | -0.031 |
| P81605 | Dermcidin OS=Homo sapiens GN=DCD PE=1 SV=2 - [DCD_HUMAN] | | 1.678E7 | 1.473E7 | 2.208E7 | | 7.225 | 7.168 | 7.344 | 7.225 | 7.256 | -0.031 |
| F6RFD5 | Uncharacterized protein OS=Homo sapiens GN=DSTN PE=4 SV=1 - [F6RFD5_HUMAN] | 1.943E7 | 9.052E6 | 1.499E7 | 1.356E7 | 7.289 | 6.957 | 7.176 | 7.132 | 7.123 | 7.154 | -0.031 |
| E9PCX6 | Uncharacterized protein OS=Homo sapiens GN=ASNS PE=4 SV=1 - [E9PCX6_HUMAN] | 2.022E7 | 2.229E7 | 2.105E7 | 2.492E7 | 7.306 | 7.348 | 7.323 | 7.396 | 7.327 | 7.360 | -0.033 |
| F5H8I7 | Uncharacterized protein OS=Homo sapiens GN=YTHDF2 PE=4 SV=1 - [F5H8I7_HUMAN] | 1.221E7 | 8.528E6 | 1.037E7 | 1.180E7 | 7.087 | 6.931 | 7.016 | 7.072 | 7.009 | 7.044 | -0.035 |
| G5E9H4 | Dynactin 1 (P150, glued homolog, Drosophila), isoform CRA_a OS=Homo sapiens GN=DCTN1 PE=4 SV=1 - [ | 1.588E6 | 5.385E6 | 3.176E6 | | 6.201 | 6.731 | 6.502 | | 6.466 | 6.502 | -0.036 |
| P31943 | Heterogeneous nuclear ribonucleoprotein H OS=Homo sapiens GN=HNRNPH1 PE=1 SV=4 - [HNRH1_HUMAN | 1.176E8 | 9.549E7 | 1.114E8 | 1.190E8 | 8.070 | 7.980 | 8.047 | 8.075 | 8.025 | 8.061 | -0.036 |
| P53985 | Monocarboxylate transporter 1 OS=Homo sapiens GN=SLC16A1 PE=1 SV=3 - [MOT1_HUMAN] | 1.333E7 | 1.012E7 | 1.262E7 | | 7.125 | 7.005 | 7.101 | | 7.065 | 7.101 | -0.036 |
| P18621 | 60S ribosomal protein L17 OS=Homo sapiens GN=RPL17 PE=1 SV=3 - [RL17_HUMAN] | 4.886E7 | 4.592E7 | 5.117E7 | 5.180E7 | 7.689 | 7.662 | 7.709 | 7.714 | 7.675 | 7.712 | -0.036 |
| F5H8K4 | Uncharacterized protein OS=Homo sapiens GN=PSMD3 PE=4 SV=1 - [F5H8K4_HUMAN] | 1.904E7 | 1.261E7 | 2.015E7 | 1.426E7 | 7.280 | 7.101 | 7.304 | 7.154 | 7.190 | 7.229 | -0.039 |
| P52597 | Heterogeneous nuclear ribonucleoprotein F OS=Homo sapiens GN=HNRNPF PE=1 SV=3 - [HNRPF_HUMAN] | 7.441E7 | 6.398E7 | 7.129E7 | 8.087E7 | 7.872 | 7.806 | 7.853 | 7.908 | 7.839 | 7.880 | -0.042 |
| P62280 | 40S ribosomal protein S11 OS=Homo sapiens GN=RPS11 PE=1 SV=3 - [RS11_HUMAN] | 2.963E7 | 1.821E7 | 2.407E7 | 2.738E7 | 7.472 | 7.260 | 7.382 | 7.437 | 7.366 | 7.409 | -0.043 |
| P49257 | Protein ERGIC-53 OS=Homo sapiens GN=LMAN1 PE=1 SV=2 - [LMAN1_HUMAN] | 8.145E6 | 5.379E6 | 7.317E6 | | 6.911 | 6.731 | 6.864 | | 6.821 | 6.864 | -0.044 |
| O14602 | Eukaryotic translation initiation factor 1A, Y-chromosomal OS=Homo sapiens GN=EIF1AY PE=1 SV=4 - [IF1A\ | 3.237E7 | 3.060E7 | 3.330E7 | 3.639E7 | 7.510 | 7.486 | 7.522 | 7.561 | 7.498 | 7.542 | -0.044 |
| A6NIT8 | Uncharacterized protein OS=Homo sapiens GN=HNRNPL PE=2 SV=1 - [A6NIT8_HUMAN] | 4.219E7 | 2.803E7 | 3.991E7 | 3.645E7 | 7.625 | 7.448 | 7.601 | 7.562 | 7.536 | 7.581 | -0.045 |
| P26358 | DNA (cytosine-5)-methyltransferase 1 OS=Homo sapiens GN=DNMT1 PE=1 SV=2 - [DNMT1_HUMAN] | | 4.081E6 | 4.166E6 | 4.948E6 | | 6.611 | 6.620 | 6.694 | 6.611 | 6.657 | -0.046 |
| Q96II6 | NADH dehydrogenase (Ubiquinone) 1 beta subcomplex, 10, 22kDa, isoform CRA_a OS=Homo sapiens GN=NDUFB10 PE=2 SV= | | 7.049E6 | 7.971E6 | | | 6.848 | 6.902 | | 6.848 | 6.902 | -0.053 |
| F8VWC5 | Ribosomal protein L18 OS=Homo sapiens GN=RPL18 PE=3 SV=1 - [F8VWC5_HUMAN] | 7.067E7 | 5.303E7 | 6.026E7 | 7.958E7 | 7.849 | 7.724 | 7.780 | 7.901 | 7.787 | 7.840 | -0.054 |
| Q99471 | Prefoldin subunit 5 OS=Homo sapiens GN=PFDN5 PE=1 SV=2 - [PFD5_HUMAN] | 6.284E6 | 4.702E6 | 5.624E6 | 6.764E6 | 6.798 | 6.672 | 6.750 | 6.830 | 6.735 | 6.790 | -0.055 |
| P61221 | ATP-binding cassette sub-family E member 1 OS=Homo sapiens GN=ABCE1 PE=1 SV=1 - [ABCE1_HUMAN] | 2.843E7 | 3.684E6 | 7.406E6 | 1.822E7 | 7.454 | 6.566 | 6.870 | 7.261 | 7.010 | 7.065 | -0.055 |
| B1AHB1 | MCM5 minichromosome maintenance deficient 5, cell division cycle 46 (S. cerevisiae), isoform CRA_c OS=Hor | 8.296E6 | 1.030E7 | 1.077E7 | 1.032E7 | 6.919 | 7.013 | 7.032 | 7.014 | 6.966 | 7.023 | -0.057 |
| P62917 | 60S ribosomal protein L8 OS=Homo sapiens GN=RPL8 PE=1 SV=2 - [RL8_HUMAN] | 7.568E7 | 4.242E7 | 6.457E7 | 6.510E7 | 7.879 | 7.628 | 7.810 | 7.814 | 7.753 | 7.812 | -0.059 |
| P18859 | ATP synthase-coupling factor 6, mitochondrial OS=Homo sapiens GN=ATP5J PE=1 SV=1 - [ATP5J_HUMAN] | | 1.404E7 | 1.493E7 | 1.731E7 | | 7.147 | 7.174 | 7.238 | 7.147 | 7.206 | -0.059 |
| P23193 | Transcription elongation factor A protein 1 OS=Homo sapiens GN=TCEA1 PE=1 SV=2 - [TCEA1_HUMAN] | 1.759E7 | 1.068E7 | 1.205E7 | 2.056E7 | 7.245 | 7.029 | 7.081 | 7.313 | 7.137 | 7.197 | -0.060 |
| O75934 | Pre-mRNA-splicing factor SPF27 OS=Homo sapiens GN=BCAS2 PE=1 SV=1 - [SPF27_HUMAN] | 2.925E7 | 3.628E7 | 3.757E7 | 3.766E7 | 7.466 | 7.560 | 7.575 | 7.576 | 7.513 | 7.575 | -0.062 |
| P08670 | Vimentin OS=Homo sapiens GN=VIM PE=1 SV=4 - [VIME_HUMAN] | 4.643E7 | 2.917E7 | 4.176E7 | 4.392E7 | 7.667 | 7.465 | 7.621 | 7.643 | 7.566 | 7.632 | -0.066 |
| Q9UHD1 | Cysteine and histidine-rich domain-containing protein 1 OS=Homo sapiens GN=CHORDC1 PE=1 SV=2 - [CHRD1_HUMAN] | | 1.182E7 | 1.055E7 | 1.795E7 | | 7.073 | 7.023 | 7.254 | 7.073 | 7.139 | -0.066 |
| B3KXI2 | Uncharacterized protein OS=Homo sapiens GN=SEC23A PE=2 SV=1 - [B3KXI2_HUMAN] | | 5.821E6 | 7.540E6 | 6.105E6 | | 6.765 | 6.877 | 6.786 | 6.765 | 6.832 | -0.067 |
| Q9UBE0 | SUMO-activating enzyme subunit 1 OS=Homo sapiens GN=SAE1 PE=1 SV=1 - [SAE1_HUMAN] | 1.512E7 | 3.166E7 | 2.640E7 | 2.469E7 | 7.180 | 7.500 | 7.422 | 7.393 | 7.340 | 7.407 | -0.067 |
| E7ERH2 | Uncharacterized protein OS=Homo sapiens GN=SKP1 PE=4 SV=1 - [E7ERH2_HUMAN] | 3.101E7 | 5.691E7 | 4.738E7 | 5.076E7 | 7.491 | 7.755 | 7.676 | 7.705 | 7.623 | 7.691 | -0.067 |
| P11387 | DNA topoisomerase 1 OS=Homo sapiens GN=TOP1 PE=1 SV=2 - [TOP1_HUMAN] | | 4.187E6 | 3.424E6 | 7.053E6 | | 6.622 | 6.535 | 6.848 | 6.622 | 6.691 | -0.070 |
| P33176 | Kinesin-1 heavy chain OS=Homo sapiens GN=KIF5B PE=1 SV=1 - [KINH_HUMAN] | 6.895E6 | 8.348E6 | 9.247E6 | 8.636E6 | 6.839 | 6.922 | 6.966 | 6.936 | 6.880 | 6.951 | -0.071 |
| P55795 | Heterogeneous nuclear ribonucleoprotein H2 OS=Homo sapiens GN=HNRNPH2 PE=1 SV=1 - [HNRH2_HUMA | 9.719E7 | 6.871E7 | 9.401E7 | 9.905E7 | 7.988 | 7.837 | 7.973 | 7.996 | 7.912 | 7.985 | -0.072 |
| P82932 | 28S ribosomal protein S6, mitochondrial OS=Homo sapiens GN=MRPS6 PE=1 SV=3 - [RT06_HUMAN] | 5.641E6 | 8.578E6 | 7.914E6 | 8.621E6 | 6.751 | 6.933 | 6.898 | 6.936 | 6.842 | 6.917 | -0.075 |
| P51572 | B-cell receptor-associated protein 31 OS=Homo sapiens GN=BCAP31 PE=1 SV=3 - [BAP31_HUMAN] | 1.456E7 | 1.023E7 | 1.550E7 | 1.358E7 | 7.163 | 7.010 | 7.190 | 7.133 | 7.086 | 7.162 | -0.075 |
| B7Z1N6 | Fructose-bisphosphate aldolase OS=Homo sapiens GN=ALDOC PE=2 SV=1 - [B7Z1N6_HUMAN] | 2.182E7 | 5.475E7 | 3.543E7 | 4.796E7 | 7.339 | 7.738 | 7.549 | 7.681 | 7.539 | 7.615 | -0.077 |
| O14979-3 | Isoform 3 of Heterogeneous nuclear ribonucleoprotein D-like OS=Homo sapiens GN=HNRPDL - [HNRDL_HUM | 3.157E7 | 2.411E7 | 3.258E7 | 3.325E7 | 7.499 | 7.382 | 7.513 | 7.522 | 7.441 | 7.517 | -0.077 |
| B4DWV7 | Uncharacterized protein OS=Homo sapiens GN=DDX20 PE=2 SV=1 - [B4DWV7_HUMAN] | 1.838E7 | 1.497E7 | 1.906E7 | 2.074E7 | 7.264 | 7.175 | 7.280 | 7.317 | 7.220 | 7.298 | -0.079 |
| P62316 | Small nuclear ribonucleoprotein Sm D2 OS=Homo sapiens GN=SNRPD2 PE=1 SV=1 - [SMD2_HUMAN] | 5.516E7 | 5.006E6 | 4.681E7 | 8.629E6 | 7.742 | 6.700 | 7.670 | 6.936 | 7.221 | 7.303 | -0.083 |
| P18754 | Regulator of chromosome condensation OS=Homo sapiens GN=RCC1 PE=1 SV=1 - [RCC1_HUMAN] | 2.802E7 | 2.994E7 | 2.411E7 | 5.166E7 | 7.447 | 7.476 | 7.382 | 7.713 | 7.462 | 7.548 | -0.086 |
| P13798 | Acylamino-acid-releasing enzyme OS=Homo sapiens GN=APEH PE=1 SV=4 - [ACPH_HUMAN] | 6.677E6 | 7.347E6 | 7.531E6 | 9.688E6 | 6.825 | 6.866 | 6.877 | 6.986 | 6.845 | 6.932 | -0.086 |
| O95232 | Luc7-like protein 3 OS=Homo sapiens GN=LUC7L3 PE=1 SV=2 - [LC7L3_HUMAN] | 5.957E6 | 1.155E6 | 3.027E6 | 3.383E6 | 6.775 | 6.062 | 6.481 | 6.529 | 6.419 | 6.505 | -0.086 |
| B4DLN1 | Uncharacterized protein OS=Homo sapiens GN=SLC25A10 PE=2 SV=1 - [B4DLN1_HUMAN] | 1.692E7 | 4.124E7 | 3.200E7 | 3.261E7 | 7.228 | 7.615 | 7.505 | 7.513 | 7.422 | 7.509 | -0.087 |
| Q00839-2 | Isoform Short of Heterogeneous nuclear ribonucleoprotein U OS=Homo sapiens GN=HNRNPU - [HNRPU_HUN | 2.009E8 | 1.532E8 | 2.129E8 | 2.197E8 | 8.303 | 8.185 | 8.328 | 8.342 | 8.244 | 8.335 | -0.091 |
| P61313 | 60S ribosomal protein L15 OS=Homo sapiens GN=RPL15 PE=1 SV=2 - [RL15_HUMAN] | 2.838E7 | 2.248E7 | 3.115E7 | 3.124E7 | 7.453 | 7.352 | 7.493 | 7.495 | 7.402 | 7.494 | -0.092 |
| Q9H3N1 | Thioredoxin-related transmembrane protein 1 OS=Homo sapiens GN=TMX1 PE=1 SV=1 - [TMX1_HUMAN] | 5.444E6 | | 5.621E6 | 8.059E6 | 6.736 | | 6.750 | 6.906 | 6.736 | 6.828 | -0.092 |
| E9PBK7 | Uncharacterized protein OS=Homo sapiens GN=MAPK3 PE=3 SV=1 - [E9PBK7_HUMAN] | | 1.215E7 | 1.297E7 | 1.746E7 | | 7.085 | 7.113 | 7.242 | 7.085 | 7.177 | -0.093 |
| F8VYH9 | Uncharacterized protein OS=Homo sapiens GN=CDK4 PE=3 SV=1 - [F8VYH9_HUMAN] | 1.247E7 | 9.167E6 | 1.276E7 | 1.394E7 | 7.096 | 6.962 | 7.106 | 7.144 | 7.029 | 7.125 | -0.096 |
| Q13242 | Serine/arginine-rich splicing factor 9 OS=Homo sapiens GN=SRSF9 PE=1 SV=1 - [SRSF9_HUMAN] | | 8.081E6 | | 1.018E7 | | 6.907 | | 7.008 | 6.907 | 7.008 | -0.100 |
| P62306 | Small nuclear ribonucleoprotein F OS=Homo sapiens GN=SNRPF PE=1 SV=1 - [RUXF_HUMAN] | 1.724E7 | | 2.467E7 | 1.939E7 | 7.237 | | 7.392 | 7.288 | 7.237 | 7.340 | -0.103 |
| O75368 | SH3 domain-binding glutamic acid-rich-like protein OS=Homo sapiens GN=SH3BGRL PE=1 SV=1 - [SH3L1_H | 5.832E6 | 1.170E7 | 1.164E7 | 9.612E6 | 6.766 | 7.068 | 7.066 | 6.983 | 6.917 | 7.024 | -0.107 |
| P05198 | Eukaryotic translation initiation factor 2 subunit 1 OS=Homo sapiens GN=EIF2S1 PE=1 SV=3 - [IF2A_HUMAN | 2.094E7 | 1.201E7 | 1.977E7 | 2.087E7 | 7.321 | 7.079 | 7.296 | 7.319 | 7.200 | 7.308 | -0.107 |
| P0CB43 | Protein FAM203B OS=Homo sapiens GN=FAM203B PE=3 SV=1 - [F203B_HUMAN] | 3.471E6 | 4.544E6 | | 5.121E6 | 6.540 | 6.657 | | 6.709 | 6.599 | 6.709 | -0.110 |
| P30519 | Heme oxygenase 2 OS=Homo sapiens GN=HMOX2 PE=1 SV=2 - [HMOX2_HUMAN] | 9.393E6 | 1.214E7 | 1.363E7 | 1.394E7 | 6.973 | 7.084 | 7.134 | 7.144 | 7.029 | 7.139 | -0.111 |
| Q9UK45 | U6 snRNA-associated Sm-like protein LSm7 OS=Homo sapiens GN=LSM7 PE=1 SV=1 - [LSM7_HUMAN] | 1.070E7 | 8.287E6 | 1.125E7 | 1.339E7 | 7.029 | 6.918 | 7.051 | 7.127 | 6.974 | 7.089 | -0.115 |
| O14654 | Insulin receptor substrate 4 OS=Homo sapiens GN=IRS4 PE=1 SV=1 - [IRS4_HUMAN] | | 4.507E6 | | 5.887E6 | | 6.654 | | 6.770 | 6.654 | 6.770 | -0.116 |
| Q15717 | ELAV-like protein 1 OS=Homo sapiens GN=ELAVL1 PE=1 SV=2 - [ELAV1_HUMAN] | 1.601E7 | 9.951E6 | 1.515E7 | 1.817E7 | 7.204 | 6.998 | 7.180 | 7.259 | 7.101 | 7.220 | -0.119 |
| P25325 | 3-mercaptopyruvate sulfurtransferase OS=Homo sapiens GN=MPST PE=1 SV=3 - [THTM_HUMAN] | 1.665E7 | 9.557E6 | | 1.660E7 | 7.221 | 6.980 | | 7.220 | 7.101 | 7.220 | -0.119 |
| Q9NS69 | Mitochondrial import receptor subunit TOM22 homolog OS=Homo sapiens GN=TOMM22 PE=1 SV=3 - [TOM2 | 2.460E7 | 2.154E7 | 3.235E7 | 2.861E7 | 7.391 | 7.333 | 7.510 | 7.457 | 7.362 | 7.483 | -0.121 |
| B4E2G8 | Uncharacterized protein OS=Homo sapiens GN=CTNNA1 PE=2 SV=1 - [B4E2G8_HUMAN] | 2.470E6 | 5.142E6 | 4.528E6 | 4.911E6 | 6.393 | 6.711 | 6.656 | 6.691 | 6.552 | 6.674 | -0.122 |
| P15121 | Aldose reductase OS=Homo sapiens GN=AKR1B1 PE=1 SV=3 - [ALDR_HUMAN] | 5.001E7 | 4.423E7 | 6.126E7 | 6.438E7 | 7.699 | 7.646 | 7.787 | 7.809 | 7.672 | 7.798 | -0.126 |
| Q9P2E9-2 | Isoform 1 of Ribosome-binding protein 1 OS=Homo sapiens GN=RRBP1 - [RRBP1_HUMAN] | | 2.297E6 | 3.021E6 | 3.116E6 | | 6.361 | 6.480 | 6.494 | 6.361 | 6.487 | -0.126 |
| G3V1R4 | NOL1/NOP2/Sun domain family, member 2, isoform CRA_b OS=Homo sapiens GN=NSUN2 PE=4 SV=1 - [G3V1R4_HUMAN] | | 9.121E6 | 1.578E7 | 9.404E6 | | 6.960 | 7.198 | 6.973 | 6.960 | 7.086 | -0.126 |
| P43243 | Matrin-3 OS=Homo sapiens GN=MATR3 PE=1 SV=2 - [MATR3_HUMAN] | 3.557E7 | 2.804E7 | 4.070E7 | 4.497E7 | 7.551 | 7.448 | 7.610 | 7.653 | 7.499 | 7.631 | -0.132 |

| | | | | | | | | | | | | |
|---|---|---|---|---|---|---|---|---|---|---|---|---|
| Q9NR30-2 | Isoform 2 of Nucleolar RNA helicase 2 OS=Homo sapiens GN=DDX21 - [DDX21_HUMAN] | 1.567E7 | 1.381E7 | 2.060E7 | 1.953E7 | 7.195 | 7.140 | 7.314 | 7.291 | 7.168 | 7.302 | -0.135 |
| P28072 | Proteasome subunit beta type-6 OS=Homo sapiens GN=PSMB6 PE=1 SV=4 - [PSB6_HUMAN] | 1.914E7 | 1.985E7 | 2.421E7 | 2.918E7 | 7.282 | 7.298 | 7.384 | 7.465 | 7.290 | 7.425 | -0.135 |
| P31942-3 | Isoform 3 of Heterogeneous nuclear ribonucleoprotein H3 OS=Homo sapiens GN=HNRNPH3 - [HNRH3_HUM/ | 2.207E7 | 1.649E7 | 2.558E7 | 2.656E7 | 7.344 | 7.217 | 7.408 | 7.424 | 7.281 | 7.416 | -0.136 |
| P22061 | Protein-L-isoaspartate(D-aspartate) O-methyltransferase OS=Homo sapiens GN=PCMT1 PE=1 SV=4 - [PIMT_ | 5.063E7 | 8.588E7 | 1.004E8 | 8.163E7 | 7.704 | 7.934 | 8.002 | 7.912 | 7.819 | 7.957 | -0.138 |
| E9PFD4 | Uncharacterized protein OS=Homo sapiens GN=BZW2 PE=4 SV=1 - [E9PFD4_HUMAN] | 2.301E6 | 3.427E6 | 3.873E6 | 3.860E6 | 6.362 | 6.535 | 6.588 | 6.587 | 6.448 | 6.587 | -0.139 |
| P38159 | Heterogeneous nuclear ribonucleoprotein G OS=Homo sapiens GN=RBMX PE=1 SV=3 - [HNRPG_HUMAN] | 3.030E7 | 2.019E7 | 3.432E7 | 3.382E7 | 7.481 | 7.305 | 7.536 | 7.529 | 7.393 | 7.532 | -0.139 |
| A8CZ64 | Extracellular signal-regulated kinase-2 splice variant OS=Homo sapiens GN=MAPK1 PE=2 SV=1 - [A8CZ64_HUMAN] | | 1.320E7 | 1.449E7 | 2.301E7 | | 7.120 | 7.161 | 7.362 | 7.120 | 7.262 | -0.141 |
| E9PFP2 | Uncharacterized protein OS=Homo sapiens GN=HTATSF1 PE=4 SV=2 - [E9PFP2_HUMAN] | 1.293E7 | 6.956E6 | 1.261E7 | 1.370E7 | 7.112 | 6.842 | 7.101 | 7.137 | 6.977 | 7.119 | -0.142 |
| Q13442 | 28 kDa heat- and acid-stable phosphoprotein OS=Homo sapiens GN=PDAP1 PE=1 SV=1 - [HAP28_HUMAN] | | 5.828E6 | 8.355E6 | 7.852E6 | | 6.766 | 6.922 | 6.895 | 6.766 | 6.908 | -0.143 |
| P23919 | Thymidylate kinase OS=Homo sapiens GN=DTYMK PE=1 SV=4 - [KTHY_HUMAN] | | 4.015E6 | 5.260E6 | 5.987E6 | | 6.604 | 6.721 | 6.777 | 6.604 | 6.749 | -0.145 |
| P40429 | 60S ribosomal protein L13a OS=Homo sapiens GN=RPL13A PE=1 SV=2 - [RL13A_HUMAN] | 7.175E7 | 4.387E7 | 8.294E7 | 7.427E7 | 7.856 | 7.642 | 7.919 | 7.871 | 7.749 | 7.895 | -0.146 |
| E5RGS4 | Uncharacterized protein OS=Homo sapiens GN=PFDN1 PE=4 SV=1 - [E5RGS4_HUMAN] | | 6.487E6 | 8.830E6 | 9.478E6 | | 6.812 | 6.946 | 6.977 | 6.812 | 6.961 | -0.149 |
| P09651-3 | Isoform 2 of Heterogeneous nuclear ribonucleoprotein A1 OS=Homo sapiens GN=HNRNPA1 - [ROA1_HUMAN | 1.571E8 | 1.345E8 | 2.095E8 | 2.041E8 | 8.196 | 8.129 | 8.321 | 8.310 | 8.162 | 8.316 | -0.153 |
| Q9H773 | dCTP pyrophosphatase 1 OS=Homo sapiens GN=DCTPP1 PE=1 SV=1 - [DCTP1_HUMAN] | 1.099E7 | 1.757E7 | 1.802E7 | 2.181E7 | 7.041 | 7.245 | 7.256 | 7.339 | 7.143 | 7.297 | -0.154 |
| Q9BRJ2 | 39S ribosomal protein L45, mitochondrial OS=Homo sapiens GN=MRPL45 PE=1 SV=2 - [RM45_HUMAN] | | 7.732E6 | 9.578E6 | 1.301E7 | | 6.888 | 6.981 | 7.114 | 6.888 | 7.048 | -0.160 |
| O43390 | Heterogeneous nuclear ribonucleoprotein R OS=Homo sapiens GN=HNRNPR PE=1 SV=1 - [HNRPR_HUMAN] | 3.683E7 | 2.521E7 | 4.391E7 | 4.421E7 | 7.566 | 7.402 | 7.643 | 7.646 | 7.484 | 7.644 | -0.160 |
| G3V4W0 | Uncharacterized protein OS=Homo sapiens GN=HNRNPC PE=4 SV=1 - [G3V4W0_HUMAN] | 1.518E8 | 1.098E8 | 1.985E8 | 1.788E8 | 8.181 | 8.041 | 8.298 | 8.252 | 8.111 | 8.275 | -0.164 |
| Q15785 | Mitochondrial import receptor subunit TOM34 OS=Homo sapiens GN=TOMM34 PE=1 SV=2 - [TOM34_HUMAI | 1.186E7 | 1.498E6 | 6.409E6 | 6.124E6 | 7.074 | 6.175 | 6.807 | 6.787 | 6.625 | 6.797 | -0.172 |
| O94776 | Metastasis-associated protein MTA2 OS=Homo sapiens GN=MTA2 PE=1 SV=1 - [MTA2_HUMAN] | 4.493E6 | 3.313E6 | | 5.759E6 | 6.653 | 6.520 | | 6.760 | 6.586 | 6.760 | -0.174 |
| P62273 | 40S ribosomal protein S29 OS=Homo sapiens GN=RPS29 PE=1 SV=2 - [RS29_HUMAN] | 3.572E7 | 2.403E7 | 5.183E7 | 3.712E7 | 7.553 | 7.381 | 7.715 | 7.570 | 7.467 | 7.642 | -0.175 |
| Q9Y5S9-2 | Isoform 2 of RNA-binding protein 8A OS=Homo sapiens GN=RBM8A - [RBM8A_HUMAN] | 2.822E7 | 1.964E7 | 3.052E7 | 4.431E7 | 7.451 | 7.293 | 7.485 | 7.647 | 7.372 | 7.566 | -0.194 |
| Q00796 | Sorbitol dehydrogenase OS=Homo sapiens GN=SORD PE=1 SV=4 - [DHSO_HUMAN] | 8.868E6 | 4.228E6 | 1.083E7 | 8.558E6 | 6.948 | 6.626 | 7.034 | 6.932 | 6.787 | 6.983 | -0.196 |
| Q14566 | DNA replication licensing factor MCM6 OS=Homo sapiens GN=MCM6 PE=1 SV=1 - [MCM6_HUMAN] | 1.010E7 | 7.285E6 | 1.193E7 | 1.537E7 | 7.004 | 6.862 | 7.077 | 7.187 | 6.933 | 7.132 | -0.198 |
| Q9Y2S6 | Coiled-coil domain-containing protein 72 OS=Homo sapiens GN=CCDC72 PE=1 SV=1 - [CCD72_HUMAN] | 1.654E6 | | 2.775E6 | 2.467E6 | 6.219 | | 6.443 | 6.392 | 6.219 | 6.418 | -0.199 |
| Q8WZ42-6 | Isoform 6 of Titin OS=Homo sapiens GN=TTN - [TITIN_HUMAN] | | 1.730E7 | | 2.758E7 | | 7.238 | | 7.441 | 7.238 | 7.441 | -0.203 |
| P49006 | MARCKS-related protein OS=Homo sapiens GN=MARCKSL1 PE=1 SV=2 - [MRP_HUMAN] | 6.066E6 | 5.676E6 | 7.730E6 | 1.180E7 | 6.783 | 6.754 | 6.888 | 7.072 | 6.768 | 6.980 | -0.211 |
| C9IZ01 | Uncharacterized protein OS=Homo sapiens GN=GFM1 PE=4 SV=1 - [C9IZ01_HUMAN] | | 1.376E7 | | 2.259E7 | | 7.139 | | 7.354 | 7.139 | 7.354 | -0.215 |
| Q9Y237 | Peptidyl-prolyl cis-trans isomerase NIMA-interacting 4 OS=Homo sapiens GN=PIN4 PE=1 SV=1 - [PIN4_HUM | 1.148E7 | 6.748E7 | 4.622E7 | 4.667E7 | 7.060 | 7.829 | 7.665 | 7.669 | 7.445 | 7.667 | -0.222 |
| P22626 | Heterogeneous nuclear ribonucleoproteins A2/B1 OS=Homo sapiens GN=HNRNPA2B1 PE=1 SV=2 - [ROA2_H | 2.561E8 | 2.096E8 | 3.858E8 | 3.877E8 | 8.408 | 8.321 | 8.586 | 8.589 | 8.365 | 8.587 | -0.223 |
| P62841 | 40S ribosomal protein S15 OS=Homo sapiens GN=RPS15 PE=1 SV=2 - [RS15_HUMAN] | 3.033E7 | 5.453E7 | 6.724E7 | 7.138E7 | 7.482 | 7.737 | 7.828 | 7.854 | 7.609 | 7.841 | -0.231 |
| O96019 | Actin-like protein 6A OS=Homo sapiens GN=ACTL6A PE=1 SV=1 - [ACL6A_HUMAN] | 3.652E6 | 9.089E6 | 9.936E6 | | 6.563 | 6.959 | 6.997 | | 6.761 | 6.997 | -0.237 |
| Q7Z4H3-2 | Isoform 2 of HD domain-containing protein 2 OS=Homo sapiens GN=HDDC2 - [HDDC2_HUMAN] | 5.895E6 | 6.685E6 | 9.874E6 | 1.421E7 | 6.770 | 6.825 | 6.994 | 7.152 | 6.798 | 7.073 | -0.276 |
| D6RAN8 | Mitochondrial ribosomal protein L27, isoform CRA_c OS=Homo sapiens GN=MRPL27 PE=4 SV=1 - [D6RAN8_ | 1.802E6 | 1.821E6 | | 3.424E6 | 6.256 | 6.260 | | 6.535 | 6.258 | 6.535 | -0.277 |
| E9PF10 | Uncharacterized protein OS=Homo sapiens GN=NUP155 PE=4 SV=1 - [E9PF10_HUMAN] | | 3.516E6 | 6.493E6 | 6.880E6 | | 6.546 | 6.812 | 6.838 | 6.546 | 6.825 | -0.279 |
| G3V5D9 | Uncharacterized protein OS=Homo sapiens GN=APEX1 PE=4 SV=1 - [G3V5D9_HUMAN] | 6.820E6 | 6.368E6 | 1.503E7 | 1.051E7 | 6.834 | 6.804 | 7.177 | 7.022 | 6.819 | 7.099 | -0.280 |
| P63167 | Dynein light chain 1, cytoplasmic OS=Homo sapiens GN=DYNLL1 PE=1 SV=1 - [DYL1_HUMAN] | 3.916E6 | 3.294E6 | 8.145E6 | 6.152E6 | 6.593 | 6.518 | 6.911 | 6.789 | 6.555 | 6.850 | -0.295 |
| Q96H79 | Zinc finger CCCH-type antiviral protein 1-like OS=Homo sapiens GN=ZC3HAV1L PE=1 SV=2 - [ZCCHL_HUMAI | 1.350E6 | | | 2.739E6 | 6.130 | | | 6.438 | 6.130 | 6.438 | -0.307 |
| P62877 | E3 ubiquitin-protein ligase RBX1 OS=Homo sapiens GN=RBX1 PE=1 SV=1 - [RBX1_HUMAN] | | 4.975E6 | 1.031E7 | 1.062E7 | | 6.697 | 7.013 | 7.026 | 6.697 | 7.020 | -0.323 |
| Q13409-6 | Isoform 2F of Cytoplasmic dynein 1 intermediate chain 2 OS=Homo sapiens GN=DYNC1I2 - [DC1I2_HUMAN] | 8.730E6 | 1.839E6 | 8.457E6 | | 6.941 | 6.265 | 6.927 | | 6.603 | 6.927 | -0.324 |
| P47914 | 60S ribosomal protein L29 OS=Homo sapiens GN=RPL29 PE=1 SV=2 - [RL29_HUMAN] | 6.107E7 | 3.912E7 | 1.319E8 | 9.069E7 | 7.786 | 7.592 | 8.120 | 7.958 | 7.689 | 8.039 | -0.350 |
| Q7L2E3-3 | Isoform 3 of Putative ATP-dependent RNA helicase DHX30 OS=Homo sapiens GN=DHX30 - [DHX30_HUMAN] | 1.215E6 | 1.163E6 | | 2.967E6 | 6.085 | 6.066 | | 6.472 | 6.075 | 6.472 | -0.397 |
| Q02978 | Mitochondrial 2-oxoglutarate/malate carrier protein OS=Homo sapiens GN=SLC25A11 PE=1 SV=3 - [M2OM_I | 1.256E7 | 9.137E6 | 2.835E7 | 2.842E7 | 7.099 | 6.961 | 7.453 | 7.454 | 7.030 | 7.453 | -0.423 |
| B5MCW9 | Uncharacterized protein OS=Homo sapiens GN=EML4 PE=4 SV=2 - [B5MCW9_HUMAN] | 1.082E6 | 6.909E6 | | 7.345E6 | 6.034 | 6.839 | | 6.866 | 6.437 | 6.866 | -0.429 |
| D6R9P3 | Uncharacterized protein OS=Homo sapiens GN=HNRNPAB PE=4 SV=1 - [D6R9P3_HUMAN] | 7.647E7 | 4.505E7 | 4.347E7 | 7.356E8 | 7.884 | 7.654 | 7.638 | 8.867 | 7.769 | 8.252 | -0.484 |
| A6NN60 | Uncharacterized protein OS=Homo sapiens GN=CRYZ PE=4 SV=1 - [A6NN60_HUMAN] | 2.027E6 | 8.222E6 | 1.049E7 | 1.569E7 | 6.307 | 6.915 | 7.021 | 7.196 | 6.611 | 7.108 | -0.497 |
| P30049 | ATP synthase subunit delta, mitochondrial OS=Homo sapiens GN=ATP5D PE=1 SV=2 - [ATPD_HUMAN] | 6.386E6 | 5.704E6 | 1.840E7 | 1.984E7 | 6.805 | 6.756 | 7.265 | 7.297 | 6.781 | 7.281 | -0.500 |
| B4DIC4 | Uncharacterized protein OS=Homo sapiens GN=LASP1 PE=2 SV=1 - [B4DIC4_HUMAN] | 9.086E5 | | 4.414E6 | 2.059E6 | 5.958 | | 6.645 | 6.314 | 5.958 | 6.479 | -0.521 |
| Q9HCC0-2 | Isoform 2 of Methylcrotonoyl-CoA carboxylase beta chain, mitochondrial OS=Homo sapiens GN=MCCC2 - [MCCB_HUMAN] | | 2.115E6 | | 7.842E6 | | 6.325 | | 6.894 | 6.325 | 6.894 | -0.569 |
| C8KIM0 | Glutathion reductase delta8+9 alternative splicing variant OS=Homo sapiens GN=GSR PE=2 SV=1 - [C8KIM0 | 1.194E7 | 1.079E6 | 1.335E7 | | 7.077 | 6.033 | 7.126 | | 6.555 | 7.126 | -0.570 |
| Q08752 | Peptidyl-prolyl cis-trans isomerase D OS=Homo sapiens GN=PPID PE=1 SV=3 - [PPID_HUMAN] | 1.464E6 | | | 5.538E6 | 6.165 | | | 6.743 | 6.165 | 6.743 | -0.578 |
| P61081 | NEDD8-conjugating enzyme Ubc12 OS=Homo sapiens GN=UBE2M PE=1 SV=1 - [UBC12_HUMAN] | 2.871E6 | 2.349E6 | 1.148E7 | 8.771E6 | 6.458 | 6.371 | 7.060 | 6.943 | 6.414 | 7.002 | -0.587 |
| P23368 | NAD-dependent malic enzyme, mitochondrial OS=Homo sapiens GN=ME2 PE=1 SV=1 - [MAOM_HUMAN] | 2.140E6 | | 8.794E6 | | 6.330 | | 6.944 | | 6.330 | 6.944 | -0.614 |
| F8WBS5 | Uncharacterized protein OS=Homo sapiens GN=RPL35A PE=4 SV=1 - [F8WBS5_HUMAN] | 2.422E6 | | 1.028E7 | 1.056E7 | 6.384 | | 7.012 | 7.024 | 6.384 | 7.018 | -0.634 |
| P42765 | 3-ketoacyl-CoA thiolase, mitochondrial OS=Homo sapiens GN=ACAA2 PE=1 SV=2 - [THIM_HUMAN] | 8.775E5 | 8.031E6 | 1.446E7 | 1.017E7 | 5.943 | 6.905 | 7.160 | 7.008 | 6.424 | 7.084 | -0.660 |
| Q9NPD3 | Exosome complex component RRP41 OS=Homo sapiens GN=EXOSC4 PE=1 SV=3 - [EXOS4_HUMAN] | 9.767E5 | | 8.951E6 | 5.597E6 | 5.990 | | 6.952 | 6.748 | 5.990 | 6.850 | -0.860 |
| Q9BPU6 | Dihydropyrimidinase-related protein 5 OS=Homo sapiens GN=DPYSL5 PE=1 SV=1 - [DPYL5_HUMAN] | | | | 9.063E5 | | | | 5.957 | 3 | 5.957 | -2.957 |
| F5GX50 | Uncharacterized protein OS=Homo sapiens GN=DIABLO PE=4 SV=1 - [F5GX50_HUMAN] | | | | 1.437E6 | | | | 6.158 | 3 | 6.158 | -3.158 |
| Q96EY7 | Pentatricopeptide repeat-containing protein 3, mitochondrial OS=Homo sapiens GN=PTCD3 PE=1 SV=3 - [PTCD3_HUMAN] | | | 1.106E6 | 4.073E6 | | | 6.044 | 6.610 | 3 | 6.327 | -3.327 |
| P49756 | RNA-binding protein 25 OS=Homo sapiens GN=RBM25 PE=1 SV=3 - [RBM25_HUMAN] | | | | 2.279E6 | | | | 6.358 | 3 | 6.358 | -3.358 |
| B4DSY9 | Uncharacterized protein OS=Homo sapiens GN=PRPF3 PE=2 SV=1 - [B4DSY9_HUMAN] | | | 2.399E6 | | | | 6.380 | | 3 | 6.380 | -3.380 |
| P55265-5 | Isoform 5 of Double-stranded RNA-specific adenosine deaminase OS=Homo sapiens GN=ADAR - [DSRAD_HUMAN] | | | 9.380E6 | 7.252E5 | | | 6.972 | 5.860 | 3 | 6.416 | -3.416 |
| Q96EL3 | 39S ribosomal protein L53, mitochondrial OS=Homo sapiens GN=MRPL53 PE=1 SV=1 - [RM53_HUMAN] | | | 3.231E6 | 2.945E6 | | | 6.509 | 6.469 | 3 | 6.489 | -3.489 |
| F5GXH1 | Uncharacterized protein OS=Homo sapiens GN=COPG PE=4 SV=1 - [F5GXH1_HUMAN] | | | | 3.195E6 | | | | 6.505 | 3 | 6.505 | -3.505 |
| E9PBT8 | Uncharacterized protein OS=Homo sapiens GN=SUMF2 PE=4 SV=1 - [E9PBT8_HUMAN] | | | 3.287E6 | 4.666E6 | | | 6.517 | 6.669 | 3 | 6.593 | -3.593 |
| O75348 | V-type proton ATPase subunit G 1 OS=Homo sapiens GN=ATP6V1G1 PE=1 SV=3 - [VATG1_HUMAN] | | | 6.351E6 | 2.491E6 | | | 6.803 | 6.396 | 3 | 6.600 | -3.600 |
| P55957 | BH3-interacting domain death agonist OS=Homo sapiens GN=BID PE=1 SV=1 - [BID_HUMAN] | | | 8.930E6 | 2.154E6 | | | 6.951 | 6.333 | 3 | 6.642 | -3.642 |
| Q15006 | Tetratricopeptide repeat protein 35 OS=Homo sapiens GN=TTC35 PE=1 SV=1 - [TTC35_HUMAN] | | | 6.192E6 | 3.854E6 | | | 6.792 | 6.586 | 3 | 6.689 | -3.689 |
| E5RH50 | Uncharacterized protein OS=Homo sapiens GN=LARP1 PE=4 SV=1 - [E5RH50_HUMAN] | | | | 4.903E6 | | | | 6.690 | 3 | 6.690 | -3.690 |
| A6NJA2 | Ubiquitin carboxyl-terminal hydrolase OS=Homo sapiens GN=USP14 PE=3 SV=1 - [A6NJA2_HUMAN] | | | 3.656E6 | 7.134E6 | | | 6.563 | 6.853 | 3 | 6.708 | -3.708 |
| E5RFP0 | Uncharacterized protein OS=Homo sapiens GN=NUDCD2 PE=4 SV=1 - [E5RFP0_HUMAN] | | | 8.691E6 | 5.752E6 | | | 6.939 | 6.760 | 3 | 6.849 | -3.849 |
| A6NLC4 | Uncharacterized protein OS=Homo sapiens GN=ATXN10 PE=2 SV=4 - [A6NLC4_HUMAN] | | | 9.533E6 | 5.399E6 | | | 6.979 | 6.732 | 3 | 6.856 | -3.856 |
| Q9Y4W6 | AFG3-like protein 2 OS=Homo sapiens GN=AFG3L2 PE=1 SV=2 - [AFG32_HUMAN] | | | | 7.291E6 | | | | 6.863 | 3 | 6.863 | -3.863 |
| B7Z6P0 | Nucleotide binding protein 2 (MinD homolog, E. coli), isoform CRA_d OS=Homo sapiens GN=NUBP2 PE=2 SV=1 - [B7Z6P0_HUMAN] | | | 8.827E6 | 7.129E6 | | | 6.946 | 6.853 | 3 | 6.899 | -3.899 |
| Q99961 | Endophilin-A2 OS=Homo sapiens GN=SH3GL1 PE=1 SV=1 - [SH3G1_HUMAN] | | | 6.103E6 | 1.038E7 | | | 6.786 | 7.016 | 3 | 6.901 | -3.901 |
| P46976-3 | Isoform GN-1S of Glycogenin-1 OS=Homo sapiens GN=GYG1 - [GLYG_HUMAN] | | | 6.495E6 | 1.005E7 | | | 6.813 | 7.002 | 3 | 6.907 | -3.907 |
| Q9UII2 | ATPase inhibitor, mitochondrial OS=Homo sapiens GN=ATPIF1 PE=1 SV=1 - [ATIF1_HUMAN] | | | 8.615E6 | | | | 6.935 | | 3 | 6.935 | -3.935 |
| Q9H6Z4-3 | Isoform 3 of Ran-binding protein 3 OS=Homo sapiens GN=RANBP3 - [RANB3_HUMAN] | | | 7.852E6 | 9.550E6 | | | 6.895 | 6.980 | 3 | 6.937 | -3.937 |
| F5GWQ7 | Uncharacterized protein OS=Homo sapiens GN=TAF15 PE=4 SV=1 - [F5GWQ7_HUMAN] | | | | 8.944E6 | | | | 6.952 | 3 | 6.952 | -3.952 |
| E7EV06 | Uncharacterized protein OS=Homo sapiens GN=SLC9A3R1 PE=4 SV=1 - [E7EV06_HUMAN] | | | 9.828E6 | | | | 6.992 | | 3 | 6.992 | -3.992 |
| Q1KMD3 | Heterogeneous nuclear ribonucleoprotein U-like protein 2 OS=Homo sapiens GN=HNRNPUL2 PE=1 SV=1 - [HNRL2_HUMAN] | | | 1.231E7 | 1.159E7 | | | 7.090 | 7.064 | 3 | 7.077 | -4.077 |
| B5ME91 | Uncharacterized protein OS=Homo sapiens GN=PDCD4 PE=4 SV=2 - [B5ME91_HUMAN] | | | 8.914E6 | 1.694E7 | | | 6.950 | 7.229 | 3 | 7.089 | -4.089 |

Extended Table 2

### Histones, histone modifying proteins

| Accession | Description | Identified Previously | Ratio H/C |
|---|---|---|---|
| P62805 | Histone H4 OS=Homo sapiens GN=HIST1H4A PE=1 SV=2 - [H4_HUMAN] | | 0.370 |
| P04908 | Histone H2A type 1-B/E OS=Homo sapiens GN=HIST1H2AB PE=1 SV=2 - [H2A1B_HUMAN] | | 0.257 |
| Q16777 | Histone H2A type 2-C OS=Homo sapiens GN=HIST2H2AC PE=1 SV=4 - [H2A2C_HUMAN] | | 0.194 |
| O60814 | Histone H2B type 1-K OS=Homo sapiens GN=HIST1H2BK PE=1 SV=3 - [H2B1K_HUMAN] | | 0.174 |
| O14929-2 | Isoform B of Histone acetyltransferase type B catalytic subunit OS=Homo sapiens GN=HAT1 - [HAT1_HUMAN] | | 0.168 |
| Q09028-3 | Isoform 3 of Histone-binding protein RBBP4 OS=Homo sapiens GN=RBBP4 - [RBBP4_HUMAN] | | 0.143 |
| Q86X55-2 | Isoform 2 of Histone-arginine methyltransferase CARM1 OS=Homo sapiens GN=CARM1 - [CARM1_HUMAN] | | 0.120 |
| Q92522 | Histone H1x OS=Homo sapiens GN=H1FX PE=1 SV=1 - [H1X_HUMAN] | | 0.117 |
| P16403 | Histone H1.2 OS=Homo sapiens GN=HIST1H1C PE=1 SV=2 - [H12_HUMAN] | | 0.098 |
| P10412 | Histone H1.4 OS=Homo sapiens GN=HIST1H1E PE=1 SV=2 - [H14_HUMAN] | | 0.082 |
| P84243 | Histone H3.3 OS=Homo sapiens GN=H3F3A PE=1 SV=2 - [H33_HUMAN] | | -0.027 |

### ADP and ADP-ribose metabolism

| Accession | Description | | Ratio H/C |
|---|---|---|---|
| O75874 | Isocitrate dehydrogenase [NADP] cytoplasmic OS=Homo sapiens GN=IDH1 PE=1 SV=2 - [IDHC_HUMAN] | | 0.333 |
| P53597 | Succinyl-CoA ligase [ADP/GDP-forming] subunit alpha, mitochondrial OS=Homo sapiens GN=SUCLG1 PE=1 SV=4 - [SUCA_HUMAN] | | 0.315 |
| P30043 | Flavin reductase (NADPH) OS=Homo sapiens GN=BLVRB PE=1 SV=3 - [BLVRB_HUMAN] | | 0.313 |
| P09874 | Poly [ADP-ribose] polymerase 1 OS=Homo sapiens GN=PARP1 PE=1 SV=4 - [PARP1_HUMAN] | | 0.280 |
| P61204 | ADP-ribosylation factor 3 OS=Homo sapiens GN=ARF3 PE=1 SV=2 - [ARF3_HUMAN] | | 0.220 |
| P16152 | Carbonyl reductase [NADPH] 1 OS=Homo sapiens GN=CBR1 PE=1 SV=3 - [CBR1_HUMAN] | | 0.197 |
| P05141 | ADP/ATP translocase 2 OS=Homo sapiens GN=SLC25A5 PE=1 SV=7 - [ADT2_HUMAN] | | 0.156 |
| P12236 | ADP/ATP translocase 3 OS=Homo sapiens GN=SLC25A6 PE=1 SV=4 - [ADT3_HUMAN] | | 0.126 |
| Q9UJY4 | ADP-ribosylation factor-binding protein GGA2 OS=Homo sapiens GN=GGA2 PE=1 SV=3 - [GGA2_HUMAN] | | 0.073 |
| B4DFL2 | Isocitrate dehydrogenase [NADP] OS=Homo sapiens GN=IDH2 PE=2 SV=1 - [B4DFL2_HUMAN] | | 0.012 |
| P11908 | Ribose-phosphate pyrophosphokinase 2 OS=Homo sapiens GN=PRPS2 PE=1 SV=2 - [PRPS2_HUMAN] | | 0.157 |
| P60891 | Ribose-phosphate pyrophosphokinase 1 OS=Homo sapiens GN=PRPS1 PE=1 SV=2 - [PRPS1_HUMAN] | | 0.091 |
| Q9UKK9 | ADP-sugar pyrophosphatase OS=Homo sapiens GN=NUDT5 PE=1 SV=1 - [NUDT5_HUMAN] | | 0.366 |

### ATP synthesis, metabolism

| Accession | Description | | Ratio H/C |
|---|---|---|---|
| P56385 | ATP synthase subunit e, mitochondrial OS=Homo sapiens GN=ATP5I PE=1 SV=2 - [ATP5I_HUMAN] | | 4.326 |
| A8MUN4 | Uncharacterized protein OS=Homo sapiens GN=ATP6V1E1 PE=2 SV=1 - [A8MUN4_HUMAN] | | 0.470 |
| P21281 | V-type proton ATPase subunit B, brain isoform OS=Homo sapiens GN=ATP6V1B2 PE=1 SV=3 - [VATB2_HUMAN] | | 0.386 |
| Q9NVI7-2 | Isoform 2 of ATPase family AAA domain-containing protein 3A OS=Homo sapiens GN=ATAD3A - [ATD3A_HUMAN] | | 0.283 |
| P55072 | Transitional endoplasmic reticulum ATPase OS=Homo sapiens GN=VCP PE=1 SV=4 - [TERA_HUMAN] | | 0.218 |
| P55036 | 26S proteasome non-ATPase regulatory subunit 4 OS=Homo sapiens GN=PSMD4 PE=1 SV=1 - [PSMD4_HUMAN] | | 0.214 |
| O95433 | Activator of 90 kDa heat shock protein ATPase homolog 1 OS=Homo sapiens GN=AHSA1 PE=1 SV=1 - [AHSA1_HUMAN] | | 0.203 |
| B1AJY5 | Proteasome (Prosome, macropain) 26S subunit, non-ATPase, 10 OS=Homo sapiens GN=PSMD10 PE=4 SV=1 - [B1AJY5_HUMAN] | | 0.189 |
| O00487 | 26S proteasome non-ATPase regulatory subunit 14 OS=Homo sapiens GN=PSMD14 PE=1 SV=1 - [PSDE_HUMAN] | | 0.188 |
| B7Z3U6 | ATPase, Na+/K+ transporting, alpha 1 polypeptide, isoform CRA_a OS=Homo sapiens GN=ATP1A1 PE=2 SV=1 - [B7Z3U6_HUMAN] | | 0.179 |
| P36542 | ATP synthase subunit gamma, mitochondrial OS=Homo sapiens GN=ATP5C1 PE=1 SV=1 - [ATPG_HUMAN] | | 0.172 |
| P06576 | ATP synthase subunit beta, mitochondrial OS=Homo sapiens GN=ATP5B PE=1 SV=3 - [ATPB_HUMAN] | | 0.171 |
| P25705 | ATP synthase subunit alpha, mitochondrial OS=Homo sapiens GN=ATP5A1 PE=1 SV=1 - [ATPA_HUMAN] | | 0.170 |
| Q15008 | 26S proteasome non-ATPase regulatory subunit 6 OS=Homo sapiens GN=PSMD6 PE=1 SV=1 - [PSMD6_HUMAN] | | 0.170 |
| C9J8H9 | Uncharacterized protein OS=Homo sapiens GN=ATP5J2 PE=4 SV=1 - [C9J8H9_HUMAN] | | 0.162 |
| Q13200 | 26S proteasome non-ATPase regulatory subunit 2 OS=Homo sapiens GN=PSMD2 PE=1 SV=3 - [PSMD2_HUMAN] | | 0.161 |
| O75947-2 | Isoform 2 of ATP synthase subunit d, mitochondrial OS=Homo sapiens GN=ATP5H - [ATP5H_HUMAN] | | 0.158 |
| P05141 | ADP/ATP translocase 2 OS=Homo sapiens GN=SLC25A5 PE=1 SV=7 - [ADT2_HUMAN] | | 0.156 |
| P26196 | Probable ATP-dependent RNA helicase DDX6 OS=Homo sapiens GN=DDX6 PE=1 SV=2 - [DDX6_HUMAN] | | 0.152 |
| P24539 | ATP synthase subunit b, mitochondrial OS=Homo sapiens GN=ATP5F1 PE=1 SV=2 - [AT5F1_HUMAN] | | 0.149 |
| B7Z1R5 | Uncharacterized protein OS=Homo sapiens GN=ATP6V1A PE=2 SV=1 - [B7Z1R5_HUMAN] | | 0.145 |
| O00232 | 26S proteasome non-ATPase regulatory subunit 12 OS=Homo sapiens GN=PSMD12 PE=1 SV=3 - [PSD12_HUMAN] | | 0.140 |
| P48047 | ATP synthase subunit O, mitochondrial OS=Homo sapiens GN=ATP5O PE=1 SV=1 - [ATPO_HUMAN] | | 0.139 |
| P12236 | ADP/ATP translocase 3 OS=Homo sapiens GN=SLC25A6 PE=1 SV=4 - [ADT3_HUMAN] | | 0.126 |
| P51665 | 26S proteasome non-ATPase regulatory subunit 7 OS=Homo sapiens GN=PSMD7 PE=1 SV=2 - [PSD7_HUMAN] | | 0.111 |
| C9JA36 | Uncharacterized protein OS=Homo sapiens GN=ATP1B3 PE=3 SV=1 - [C9JA36_HUMAN] | | 0.110 |
| O00231 | 26S proteasome non-ATPase regulatory subunit 11 OS=Homo sapiens GN=PSMD11 PE=1 SV=3 - [PSD11_HUMAN] | | 0.110 |
| Q9NTK5 | Obg-like ATPase 1 OS=Homo sapiens GN=OLA1 PE=1 SV=2 - [OLA1_HUMAN] | | 0.103 |
| Q92499 | ATP-dependent RNA helicase DDX1 OS=Homo sapiens GN=DDX1 PE=1 SV=2 - [DDX1_HUMAN] | | 0.103 |
| P17844 | Probable ATP-dependent RNA helicase DDX5 OS=Homo sapiens GN=DDX5 PE=1 SV=1 - [DDX5_HUMAN] | | 0.087 |
| E9PN17 | Uncharacterized protein OS=Homo sapiens GN=ATP5L PE=4 SV=1 - [E9PN17_HUMAN] | | 0.082 |
| B4DJ66 | Proteasome (Prosome, macropain) 26S subunit, non-ATPase, 13, isoform CRA_d OS=Homo sapiens GN=PSMD13 PE=2 SV=1 - [B4DJ66_HUMAN] | | 0.060 |
| P53396 | ATP-citrate synthase OS=Homo sapiens GN=ACLY PE=1 SV=3 - [ACLY_HUMAN] | | 0.057 |
| Q92841-4 | Isoform 4 of Probable ATP-dependent RNA helicase DDX17 OS=Homo sapiens GN=DDX17 - [DDX17_HUMAN] | | 0.056 |
| Q7L014 | Probable ATP-dependent RNA helicase DDX46 OS=Homo sapiens GN=DDX46 PE=1 SV=2 - [DDX46_HUMAN] | | 0.040 |
| Q9NUU7 | ATP-dependent RNA helicase DDX19A OS=Homo sapiens GN=DDX19A PE=1 SV=1 - [DD19A_HUMAN] | | 0.019 |
| O00571 | ATP-dependent RNA helicase DDX3X OS=Homo sapiens GN=DDX3X PE=1 SV=3 - [DDX3X_HUMAN] | | 0.008 |
| Q08211 | ATP-dependent RNA helicase A OS=Homo sapiens GN=DHX9 PE=1 SV=4 - [DHX9_HUMAN] | | -0.022 |
| Q99460-2 | Isoform 2 of 26S proteasome non-ATPase regulatory subunit 1 OS=Homo sapiens GN=PSMD1 - [PSMD1_HUMAN] | | -0.023 |
| P61221 | ATP-binding cassette sub-family E member 1 OS=Homo sapiens GN=ABCE1 PE=1 SV=1 - [ABCE1_HUMAN] | | -0.055 |
| P18859 | ATP synthase-coupling factor 6, mitochondrial OS=Homo sapiens GN=ATP5J PE=1 SV=1 - [ATP5J_HUMAN] | | -0.059 |
| Q7L2E3-3 | Isoform 3 of Putative ATP-dependent RNA helicase DHX30 OS=Homo sapiens GN=DHX30 - [DHX30_HUMAN] | | -0.397 |
| P30049 | ATP synthase subunit delta, mitochondrial OS=Homo sapiens GN=ATP5D PE=1 SV=2 - [ATPD_HUMAN] | | -0.500 |
| O75348 | V-type proton ATPase subunit G 1 OS=Homo sapiens GN=ATP6V1G1 PE=1 SV=3 - [VATG1_HUMAN] | | -3.600 |
| Q9UII2 | ATPase inhibitor, mitochondrial OS=Homo sapiens GN=ATPIF1 PE=1 SV=1 - [ATIF1_HUMAN] | | -3.935 |

NAD metabolism

| Accession | Description | Ratio H/C |
|---|---|---|
| O95182 | NADH dehydrogenase [ubiquinone] 1 alpha subcomplex subunit 7 OS=Homo sapiens GN=NDUFA7 PE=1 SV=3 - [NDUA7_HUMAN] | 3.672 |
| Q9P0J0 | NADH dehydrogenase [ubiquinone] 1 alpha subcomplex subunit 13 OS=Homo sapiens GN=NDUFA13 PE=1 SV=3 - [NDUAD_HUMAN] | 0.784 |
| O75874 | Isocitrate dehydrogenase [NADP] cytoplasmic OS=Homo sapiens GN=IDH1 PE=1 SV=2 - [IDHC_HUMAN] | 0.333 |
| P30043 | Flavin reductase (NADPH) OS=Homo sapiens GN=BLVRB PE=1 SV=3 - [BLVRB_HUMAN] | 0.313 |
| P16152 | Carbonyl reductase [NADPH] 1 OS=Homo sapiens GN=CBR1 PE=1 SV=3 - [CBR1_HUMAN] | 0.197 |
| Q9BU61 | NADH dehydrogenase [ubiquinone] 1 alpha subcomplex assembly factor 3 OS=Homo sapiens GN=NDUFAF3 PE=1 SV=1 - [NDUF3_HUMAN] | 0.142 |
| P50213 | Isocitrate dehydrogenase [NAD] subunit alpha, mitochondrial OS=Homo sapiens GN=IDH3A PE=1 SV=1 - [IDH3A_HUMAN] | 0.098 |
| P00387-2 | Isoform 2 of NADH-cytochrome b5 reductase 3 OS=Homo sapiens GN=CYB5R3 - [NB5R3_HUMAN] | 0.078 |
| B4DFL2 | Isocitrate dehydrogenase [NADP] OS=Homo sapiens GN=IDH2 PE=2 SV=1 - [B4DFL2_HUMAN] | 0.012 |
| Q96II6 | NADH dehydrogenase (Ubiquinone) 1 beta subcomplex, 10, 22kDa, isoform CRA_a OS=Homo sapiens GN=NDUFB10 PE=2 SV=1 - [Q96II6_HUMAN] | -0.053 |
| P23368 | NAD-dependent malic enzyme, mitochondrial OS=Homo sapiens GN=ME2 PE=1 SV=1 - [MAOM_HUMAN] | -0.614 |

Kinases, Kinase regulatory proteins

| Accession | Description | Ratio H/C |
|---|---|---|
| Q9UNF0-2 | Isoform 2 of Protein kinase C and casein kinase substrate in neurons protein 2 OS=Homo sapiens GN=PACSIN2 - [PACN2_HUMAN] | 0.834 |
| P36507 | Dual specificity mitogen-activated protein kinase kinase 2 OS=Homo sapiens GN=MAP2K2 PE=1 SV=1 - [MP2K2_HUMAN] | 0.346 |
| P22392-2 | Isoform 3 of Nucleoside diphosphate kinase B OS=Homo sapiens GN=NME2 - [NDKB_HUMAN] | 0.297 |
| Q9H1E3 | Nuclear ubiquitous casein and cyclin-dependent kinases substrate OS=Homo sapiens GN=NUCKS1 PE=1 SV=1 - [NUCKS_HUMAN] | 0.285 |
| P00558 | Phosphoglycerate kinase 1 OS=Homo sapiens GN=PGK1 PE=1 SV=3 - [PGK1_HUMAN] | 0.262 |
| P61024 | Cyclin-dependent kinases regulatory subunit 1 OS=Homo sapiens GN=CKS1B PE=1 SV=1 - [CKS1_HUMAN] | 0.238 |
| Q9BZX2 | Uridine-cytidine kinase 2 OS=Homo sapiens GN=UCK2 PE=1 SV=1 - [UCK2_HUMAN] | 0.220 |
| Q16822 | Phosphoenolpyruvate carboxykinase [GTP], mitochondrial OS=Homo sapiens GN=PCK2 PE=1 SV=3 - [PCKGM_HUMAN] | 0.204 |
| P06493 | Cyclin-dependent kinase 1 OS=Homo sapiens GN=CDK1 PE=1 SV=3 - [CDK1_HUMAN] | 0.175 |
| P00568 | Adenylate kinase isoenzyme 1 OS=Homo sapiens GN=AK1 PE=1 SV=3 - [KAD1_HUMAN] | 0.174 |
| P11908 | Ribose-phosphate pyrophosphokinase 2 OS=Homo sapiens GN=PRPS2 PE=1 SV=2 - [PRPS2_HUMAN] | 0.157 |
| Q9H0C8 | Integrin-linked kinase-associated serine/threonine phosphatase 2C OS=Homo sapiens GN=ILKAP PE=1 SV=1 - [ILKAP_HUMAN] | 0.144 |
| P42771 | Cyclin-dependent kinase inhibitor 2A, isoforms 1/2/3 OS=Homo sapiens GN=CDKN2A PE=1 SV=2 - [CD2A1_HUMAN] | 0.137 |
| P12532 | Creatine kinase U-type, mitochondrial OS=Homo sapiens GN=CKMT1A PE=1 SV=1 - [KCRU_HUMAN] | 0.122 |
| Q9Y3F4 | Serine-threonine kinase receptor-associated protein OS=Homo sapiens GN=STRAP PE=1 SV=1 - [STRAP_HUMAN] | 0.117 |
| P78527 | DNA-dependent protein kinase catalytic subunit OS=Homo sapiens GN=PRKDC PE=1 SV=3 - [PRKDC_HUMAN] | 0.111 |
| P60891 | Ribose-phosphate pyrophosphokinase 1 OS=Homo sapiens GN=PRPS1 PE=1 SV=2 - [PRPS1_HUMAN] | 0.091 |
| P12277 | Creatine kinase B-type OS=Homo sapiens GN=CKB PE=1 SV=1 - [KCRB_HUMAN] | 0.087 |
| P14618 | Pyruvate kinase isozymes M1/M2 OS=Homo sapiens GN=PKM2 PE=1 SV=4 - [KPYM_HUMAN] | 0.076 |
| P67870 | Casein kinase II subunit beta OS=Homo sapiens GN=CSNK2B PE=1 SV=1 - [CSK2B_HUMAN] | -0.001 |
| O60271-7 | Isoform 7 of C-Jun-amino-terminal kinase-interacting protein 4 OS=Homo sapiens GN=SPAG9 - [JIP4_HUMAN] | -0.003 |
| A8CZ64 | Extracellular signal-regulated kinase-2 splice variant OS=Homo sapiens GN=MAPK1 PE=2 SV=1 - [A8CZ64_HUMAN] | -0.141 |
| P23919 | Thymidylate kinase OS=Homo sapiens GN=DTYMK PE=1 SV=4 - [KTHY_HUMAN] | -0.145 |

Phosphatases

| Accession | Description | Ratio H/C |
|---|---|---|
| P62140 | Serine/threonine-protein phosphatase PP1-beta catalytic subunit OS=Homo sapiens GN=PPP1CB PE=1 SV=3 - [PP1B_HUMAN] | 0.244 |
| Q9BSD7 | Cancer-related nucleoside-triphosphatase OS=Homo sapiens GN=NTPCR PE=1 SV=1 - [NTPCR_HUMAN] | 0.218 |
| P24666 | Low molecular weight phosphotyrosine protein phosphatase OS=Homo sapiens GN=ACP1 PE=1 SV=3 - [PPAC_HUMAN] | 0.204 |
| F8VYE8 | Serine/threonine-protein phosphatase OS=Homo sapiens GN=PPP1CC PE=3 SV=1 - [F8VYE8_HUMAN] | 0.197 |
| Q9NRX4 | 14 kDa phosphohistidine phosphatase OS=Homo sapiens GN=PHPT1 PE=1 SV=1 - [PHP14_HUMAN] | 0.147 |
| Q9H0C8 | Integrin-linked kinase-associated serine/threonine phosphatase 2C OS=Homo sapiens GN=ILKAP PE=1 SV=1 - [ILKAP_HUMAN] | 0.144 |
| Q15181 | Inorganic pyrophosphatase OS=Homo sapiens GN=PPA1 PE=1 SV=2 - [IPYR_HUMAN] | 0.143 |
| Q9H2U2-3 | Isoform 3 of Inorganic pyrophosphatase 2, mitochondrial OS=Homo sapiens GN=PPA2 - [IPYR2_HUMAN] | 0.140 |
| P62714 | Serine/threonine-protein phosphatase 2A catalytic subunit beta isoform OS=Homo sapiens GN=PPP2CB PE=1 SV=1 - [PP2AB_HUMAN] | 0.138 |
| Q96C90 | Protein phosphatase 1 regulatory subunit 14B OS=Homo sapiens GN=PPP1R14B PE=1 SV=3 - [PP14B_HUMAN] | 0.127 |
| P53041 | Serine/threonine-protein phosphatase 5 OS=Homo sapiens GN=PPP5C PE=1 SV=1 - [PPP5_HUMAN] | 0.068 |
| E9PMD7 | Serine/threonine-protein phosphatase OS=Homo sapiens GN=PPP1CA PE=3 SV=1 - [E9PMD7_HUMAN] | 0.058 |
| O15355 | Protein phosphatase 1G OS=Homo sapiens GN=PPM1G PE=1 SV=1 - [PPM1G_HUMAN] | 0.052 |
| Q96HS1-2 | Isoform 2 of Serine/threonine-protein phosphatase PGAM5, mitochondrial OS=Homo sapiens GN=PGAM5 - [PGAM5_HUMAN] | -0.022 |
| Q9H773 | dCTP pyrophosphatase 1 OS=Homo sapiens GN=DCTPP1 PE=1 SV=1 - [DCTP1_HUMAN] | -0.154 |

Extended Table 3: PAR-Mass Spec GO Biological Process enrichment Control (T0) Proteins

| Control Proteins GO Biological Process Enrichment | # Genes in Gene Set (K) | # Genes in Overlap (k) | k/K | p-value | FDR q-value |
|---|---|---|---|---|---|
| ORGANELLE_ORGANIZATION_AND_BIOGENESIS | 473 | 8 | 0.0169 | 3.12E-06 | 2.58E-03 |
| ONE_CARBON_COMPOUND_METABOLIC_PROCESS | 26 | 3 | 0.1154 | 1.59E-05 | 6.58E-03 |
| MITOCHONDRION_ORGANIZATION_AND_BIOGENESIS | 48 | 3 | 0.0625 | 1.03E-04 | 2.83E-02 |
| PROTEIN_TARGETING_TO_MITOCHONDRION | 11 | 2 | 0.1818 | 1.88E-04 | 3.88E-02 |

Extended Table 4: PAR-Mass Spec GO Biological Process enrichment Hormone (T30) Proteins

| Control Proteins GO Biological Process Enrichment | # Genes in Gene Set | # Genes in Overlap | k/K | p-value | FDR q-value |
|---|---|---|---|---|---|
| ESTABLISHMENT_OF_LOCALIZATION | 870 | 50 | 0.0575 | 0.00E+00 | 0.00E+00 |
| TRANSPORT | 795 | 46 | 0.0579 | 0.00E+00 | 0.00E+00 |
| BIOPOLYMER_METABOLIC_PROCESS | 1684 | 104 | 0.0618 | 0.00E+00 | 0.00E+00 |
| CELLULAR_PROTEIN_METABOLIC_PROCESS | 1117 | 79 | 0.0707 | 0.00E+00 | 0.00E+00 |
| CELLULAR_MACROMOLECULE_METABOLIC_PROCESS | 1131 | 80 | 0.0707 | 0.00E+00 | 0.00E+00 |
| RNA_METABOLIC_PROCESS | 841 | 60 | 0.0713 | 0.00E+00 | 0.00E+00 |
| PROTEIN_METABOLIC_PROCESS | 1231 | 90 | 0.0731 | 0.00E+00 | 0.00E+00 |
| NUCLEOBASENUCLEOSIDENUCLEOTIDE_AND_NUCLEIC_ACID_METABOLIC_PROCESS | 1244 | 102 | 0.082 | 0.00E+00 | 0.00E+00 |
| ORGANELLE_ORGANIZATION_AND_BIOGENESIS | 473 | 40 | 0.0846 | 0.00E+00 | 0.00E+00 |
| MACROMOLECULE_BIOSYNTHETIC_PROCESS | 321 | 32 | 0.0997 | 0.00E+00 | 0.00E+00 |
| BIOSYNTHETIC_PROCESS | 470 | 49 | 0.1043 | 0.00E+00 | 0.00E+00 |
| CELLULAR_COMPONENT_ASSEMBLY | 298 | 33 | 0.1107 | 0.00E+00 | 0.00E+00 |
| MACROMOLECULAR_COMPLEX_ASSEMBLY | 280 | 33 | 0.1179 | 0.00E+00 | 0.00E+00 |
| CATABOLIC_PROCESS | 225 | 28 | 0.1244 | 0.00E+00 | 0.00E+00 |
| CELLULAR_CATABOLIC_PROCESS | 212 | 28 | 0.1321 | 0.00E+00 | 0.00E+00 |
| CELLULAR_BIOSYNTHETIC_PROCESS | 321 | 45 | 0.1402 | 0.00E+00 | 0.00E+00 |
| RNA_PROCESSING | 173 | 28 | 0.1618 | 0.00E+00 | 0.00E+00 |
| TRANSLATION | 180 | 31 | 0.1722 | 0.00E+00 | 0.00E+00 |
| DNA_METABOLIC_PROCESS | 257 | 27 | 0.1051 | 1.11E-16 | 4.82E-15 |
| APOPTOSIS_GO | 431 | 32 | 0.0742 | 2.33E-15 | 9.58E-14 |
| PROGRAMMED_CELL_DEATH | 432 | 32 | 0.0741 | 2.55E-15 | 9.58E-14 |
| RNA_SPLICING | 91 | 17 | 0.1868 | 2.55E-15 | 9.58E-14 |
| ESTABLISHMENT_OF_CELLULAR_LOCALIZATION | 353 | 29 | 0.0822 | 3.66E-15 | 1.31E-13 |
| INTRACELLULAR_TRANSPORT | 280 | 26 | 0.0929 | 5.44E-15 | 1.87E-13 |
| MRNA_METABOLIC_PROCESS | 84 | 16 | 0.1905 | 1.21E-14 | 3.99E-13 |
| CELLULAR_LOCALIZATION | 371 | 29 | 0.0782 | 1.32E-14 | 4.19E-13 |
| PROTEIN_FOLDING | 58 | 14 | 0.2414 | 1.63E-14 | 4.99E-13 |
| MRNA_PROCESSING_GO_0006397 | 73 | 15 | 0.2055 | 2.49E-14 | 7.33E-13 |
| REGULATION_OF_CELLULAR_METABOLIC_PROCESS | 787 | 40 | 0.0508 | 2.35E-13 | 6.68E-12 |
| CELL_DEVELOPMENT | 577 | 34 | 0.0589 | 2.59E-13 | 7.13E-12 |
| REGULATION_OF_METABOLIC_PROCESS | 799 | 40 | 0.0501 | 3.79E-13 | 1.01E-11 |
| CARBOXYLIC_ACID_METABOLIC_PROCESS | 178 | 19 | 0.1067 | 2.14E-12 | 5.53E-11 |
| ORGANIC_ACID_METABOLIC_PROCESS | 180 | 19 | 0.1056 | 2.62E-12 | 6.55E-11 |
| REGULATION_OF_APOPTOSIS | 341 | 25 | 0.0733 | 3.66E-12 | 8.87E-11 |
| REGULATION_OF_PROGRAMMED_CELL_DEATH | 342 | 25 | 0.0731 | 3.90E-12 | 9.19E-11 |
| TRANSCRIPTION | 753 | 37 | 0.0491 | 5.09E-12 | 1.17E-10 |
| REGULATION_OF_DEVELOPMENTAL_PROCESS | 440 | 28 | 0.0636 | 5.59E-12 | 1.25E-10 |
| RESPONSE_TO_STRESS | 508 | 30 | 0.0591 | 6.34E-12 | 1.38E-10 |
| NEGATIVE_REGULATION_OF_CELLULAR_PROCESS | 646 | 32 | 0.0495 | 1.17E-10 | 2.46E-09 |
| NEGATIVE_REGULATION_OF_APOPTOSIS | 150 | 16 | 0.1067 | 1.19E-10 | 2.46E-09 |
| NEGATIVE_REGULATION_OF_PROGRAMMED_CELL_DEATH | 151 | 16 | 0.106 | 1.32E-10 | 2.65E-09 |
| MACROMOLECULE_CATABOLIC_PROCESS | 137 | 15 | 0.1095 | 3.13E-10 | 6.14E-09 |
| NEGATIVE_REGULATION_OF_BIOLOGICAL_PROCESS | 677 | 32 | 0.0473 | 3.76E-10 | 7.22E-09 |
| ESTABLISHMENT_OF_PROTEIN_LOCALIZATION | 190 | 17 | 0.0895 | 5.03E-10 | 9.42E-09 |
| DNA_REPLICATION | 102 | 13 | 0.1275 | 7.15E-10 | 1.31E-08 |
| GENERATION_OF_PRECURSOR_METABOLITES_AND_ENERGY | 123 | 14 | 0.1138 | 7.28E-10 | 1.31E-08 |
| NEGATIVE_REGULATION_OF_DEVELOPMENTAL_PROCESS | 197 | 17 | 0.0863 | 8.80E-10 | 1.54E-08 |
| PROTEIN_RNA_COMPLEX_ASSEMBLY | 67 | 11 | 0.1642 | 9.22E-10 | 1.59E-08 |
| RIBONUCLEOPROTEIN_COMPLEX_BIOGENESIS_AND_ASSEMBLY | 86 | 12 | 0.1395 | 1.11E-09 | 1.88E-08 |
| NITROGEN_COMPOUND_METABOLIC_PROCESS | 155 | 15 | 0.0968 | 1.78E-09 | 2.93E-08 |
| MACROMOLECULE_LOCALIZATION | 235 | 18 | 0.0766 | 1.92E-09 | 3.11E-08 |
| PROTEIN_TRANSPORT | 157 | 15 | 0.0955 | 2.12E-09 | 3.37E-08 |
| CELLULAR_RESPIRATION | 19 | 7 | 0.3684 | 2.39E-09 | 3.65E-08 |
| TRNA_METABOLIC_PROCESS | 19 | 7 | 0.3684 | 2.39E-09 | 3.65E-08 |
| PROTEIN_LOCALIZATION | 214 | 17 | 0.0794 | 3.12E-09 | 4.68E-08 |
| PROTEIN_COMPLEX_ASSEMBLY | 167 | 15 | 0.0898 | 4.97E-09 | 7.32E-08 |
| REGULATION_OF_GENE_EXPRESSION | 673 | 30 | 0.0446 | 5.14E-09 | 7.43E-08 |
| TRANSCRIPTION_DNA_DEPENDENT | 636 | 29 | 0.0456 | 5.63E-09 | 8.00E-08 |
| RNA_BIOSYNTHETIC_PROCESS | 638 | 29 | 0.0455 | 6.03E-09 | 8.43E-08 |
| CELLULAR_COMPONENT_DISASSEMBLY | 33 | 8 | 0.2424 | 7.18E-09 | 9.87E-08 |
| AMINO_ACID_AND_DERIVATIVE_METABOLIC_PROCESS | 101 | 12 | 0.1188 | 7.30E-09 | 9.87E-08 |
| REGULATION_OF_CELLULAR_COMPONENT_ORGANIZATION_AND_BIOGENESIS | 125 | 13 | 0.104 | 8.95E-09 | 1.19E-07 |
| CELLULAR_LIPID_CATABOLIC_PROCESS | 35 | 8 | 0.2286 | 1.19E-08 | 1.56E-07 |
| SIGNAL_TRANSDUCTION | 1634 | 50 | 0.0306 | 1.71E-08 | 2.21E-07 |
| ALCOHOL_METABOLIC_PROCESS | 88 | 11 | 0.125 | 1.80E-08 | 2.29E-07 |
| ENERGY_DERIVATION_BY_OXIDATION_OF_ORGANIC_COMPOUNDS | 37 | 8 | 0.2162 | 1.91E-08 | 2.39E-07 |
| LIPID_CATABOLIC_PROCESS | 38 | 8 | 0.2105 | 2.39E-08 | 2.94E-07 |
| ANTI_APOPTOSIS | 118 | 12 | 0.1017 | 4.29E-08 | 5.21E-07 |
| REGULATION_OF_NUCLEOBASENUCLEOSIDENUCLEOTIDE_AND_NUCLEIC_ACID_METABC | 618 | 27 | 0.0437 | 4.53E-08 | 5.41E-07 |
| INTRACELLULAR_PROTEIN_TRANSPORT | 145 | 13 | 0.0897 | 5.36E-08 | 6.31E-07 |
| ESTABLISHMENT_AND_OR_MAINTENANCE_OF_CHROMATIN_ARCHITECTURE | 77 | 10 | 0.1299 | 5.54E-08 | 6.44E-07 |
| AMINO_ACID_METABOLIC_PROCESS | 78 | 10 | 0.1282 | 6.29E-08 | 7.20E-07 |
| APOPTOTIC_PROGRAM | 60 | 9 | 0.15 | 7.16E-08 | 8.09E-07 |
| CHROMOSOME_ORGANIZATION_AND_BIOGENESIS | 124 | 12 | 0.0968 | 7.47E-08 | 8.33E-07 |
| DNA_REPAIR | 125 | 12 | 0.096 | 8.17E-08 | 8.99E-07 |
| CELLULAR_CARBOHYDRATE_METABOLIC_PROCESS | 126 | 12 | 0.0952 | 8.93E-08 | 9.69E-07 |
| CYTOSKELETON_ORGANIZATION_AND_BIOGENESIS | 208 | 15 | 0.0721 | 9.45E-08 | 1.01E-06 |
| CARBOHYDRATE_METABOLIC_PROCESS | 180 | 14 | 0.0778 | 9.92E-08 | 1.05E-06 |
| APOPTOTIC_NUCLEAR_CHANGES | 19 | 6 | 0.3158 | 1.01E-07 | 1.05E-06 |
| RESPONSE_TO_DNA_DAMAGE_STIMULUS | 162 | 13 | 0.0802 | 1.97E-07 | 2.03E-06 |
| POSITIVE_REGULATION_OF_BIOLOGICAL_PROCESS | 709 | 28 | 0.0395 | 2.04E-07 | 2.08E-06 |
| CELL_PROLIFERATION_GO_0008283 | 513 | 23 | 0.0448 | 2.86E-07 | 2.88E-06 |
| AMINE_METABOLIC_PROCESS | 141 | 12 | 0.0851 | 3.07E-07 | 3.05E-06 |
| NUCLEOBASENUCLEOSIDE_AND_NUCLEOTIDE_METABOLIC_PROCESS | 52 | 8 | 0.1538 | 3.14E-07 | 3.09E-06 |
| BIOPOLYMER_CATABOLIC_PROCESS | 117 | 11 | 0.094 | 3.50E-07 | 3.40E-06 |
| DNA_CATABOLIC_PROCESS | 23 | 6 | 0.2609 | 3.59E-07 | 3.44E-06 |
| DNA_FRAGMENTATION_DURING_APOPTOSIS | 13 | 5 | 0.3846 | 3.99E-07 | 3.78E-06 |
| COFACTOR_METABOLIC_PROCESS | 54 | 8 | 0.1481 | 4.25E-07 | 3.98E-06 |
| REGULATION_OF_TRANSCRIPTION | 566 | 24 | 0.0424 | 4.32E-07 | 4.00E-06 |
| NITROGEN_COMPOUND_BIOSYNTHETIC_PROCESS | 26 | 6 | 0.2308 | 7.92E-07 | 7.26E-06 |
| RESPONSE_TO_CHEMICAL_STIMULUS | 314 | 17 | 0.0541 | 8.06E-07 | 7.31E-06 |
| AEROBIC_RESPIRATION | 15 | 5 | 0.3333 | 9.11E-07 | 8.17E-06 |
| NUCLEOTIDE_METABOLIC_PROCESS | 42 | 7 | 0.1667 | 9.91E-07 | 8.79E-06 |
| GLUCOSE_METABOLIC_PROCESS | 28 | 6 | 0.2143 | 1.27E-06 | 1.11E-05 |
| CELLULAR_LIPID_METABOLIC_PROCESS | 255 | 15 | 0.0588 | 1.28E-06 | 1.11E-05 |
| LIPID_METABOLIC_PROCESS | 325 | 17 | 0.0523 | 1.29E-06 | 1.11E-05 |
| RESPONSE_TO_OXIDATIVE_STRESS | 46 | 7 | 0.1522 | 1.88E-06 | 1.60E-05 |
| NUCLEAR_ORGANIZATION_AND_BIOGENESIS | 30 | 6 | 0.2 | 1.96E-06 | 1.65E-05 |
| RESPONSE_TO_ENDOGENOUS_STIMULUS | 200 | 13 | 0.065 | 2.16E-06 | 1.80E-05 |
| POSITIVE_REGULATION_OF_CELLULAR_PROCESS | 668 | 25 | 0.0374 | 2.37E-06 | 1.96E-05 |

Extended Table 5

| Gene Set Name | # Genes in Gene Set | # Genes in Overla | p-value | FDR q-value |
|---|---|---|---|---|
| TRANSCRIPTION | 753 | 62 | 0.00E+00 | 0.00E+00 |
| SYSTEM_DEVELOPMENT | 861 | 67 | 0.00E+00 | 0.00E+00 |
| SIGNAL_TRANSDUCTION | 1634 | 139 | 0.00E+00 | 0.00E+00 |
| PROTEIN_METABOLIC_PROCESS | 1231 | 82 | 0.00E+00 | 0.00E+00 |
| NUCLEOBASENUCLEOSIDENUCLEOTIDE_AND_NUCLEIC_ACID_METABOI | 1244 | 82 | 0.00E+00 | 0.00E+00 |
| MULTICELLULAR_ORGANISMAL_DEVELOPMENT | 1049 | 87 | 0.00E+00 | 0.00E+00 |
| CELLULAR_PROTEIN_METABOLIC_PROCESS | 1117 | 74 | 0.00E+00 | 0.00E+00 |
| CELLULAR_MACROMOLECULE_METABOLIC_PROCESS | 1131 | 76 | 0.00E+00 | 0.00E+00 |
| CELL_SURFACE_RECEPTOR_LINKED_SIGNAL_TRANSDUCTION_GO_000 | 641 | 54 | 0.00E+00 | 0.00E+00 |
| CELL_PROLIFERATION_GO_0008283 | 513 | 48 | 0.00E+00 | 0.00E+00 |
| BIOPOLYMER_METABOLIC_PROCESS | 1684 | 108 | 0.00E+00 | 0.00E+00 |
| ANATOMICAL_STRUCTURE_DEVELOPMENT | 1013 | 77 | 0.00E+00 | 0.00E+00 |
| TRANSCRIPTION_DNA_DEPENDENT | 636 | 52 | 1.11E-16 | 1.57E-14 |
| REGULATION_OF_GENE_EXPRESSION | 673 | 54 | 1.11E-16 | 1.57E-14 |
| RNA_BIOSYNTHETIC_PROCESS | 638 | 52 | 2.22E-16 | 2.97E-14 |
| REGULATION_OF_CELLULAR_METABOLIC_PROCESS | 787 | 58 | 3.33E-16 | 4.23E-14 |
| RNA_METABOLIC_PROCESS | 841 | 60 | 4.44E-16 | 5.37E-14 |
| REGULATION_OF_METABOLIC_PROCESS | 799 | 58 | 6.66E-16 | 7.69E-14 |
| INTRACELLULAR_SIGNALING_CASCADE | 667 | 52 | 1.11E-15 | 1.23E-13 |
| REGULATION_OF_TRANSCRIPTION | 566 | 46 | 1.15E-14 | 1.22E-12 |
| ORGAN_DEVELOPMENT | 571 | 45 | 6.73E-14 | 6.47E-12 |
| REGULATION_OF_NUCLEOBASENUCLEOSIDENUCLEOTIDE_AND_NUCLE | 618 | 47 | 6.87E-14 | 6.47E-12 |
| BIOPOLYMER_MODIFICATION | 650 | 48 | 1.10E-13 | 1.00E-11 |
| NEGATIVE_REGULATION_OF_BIOLOGICAL_PROCESS | 677 | 49 | 1.29E-13 | 1.13E-11 |
| ENZYME_LINKED_RECEPTOR_PROTEIN_SIGNALING_PATHWAY | 140 | 22 | 2.76E-13 | 2.33E-11 |
| ENZYME_REGULATOR_ACTIVITY | 323 | 32 | 6.95E-13 | 5.70E-11 |
| NEGATIVE_REGULATION_OF_CELLULAR_PROCESS | 646 | 46 | 1.29E-12 | 1.02E-10 |
| PROTEIN_MODIFICATION_PROCESS | 631 | 45 | 2.15E-12 | 1.66E-10 |
| RECEPTOR_ACTIVITY | 583 | 43 | 2.22E-12 | 1.66E-10 |
| CELL_CELL_SIGNALING | 404 | 35 | 2.89E-12 | 2.10E-10 |
| TRANSCRIPTION_FROM_RNA_POLYMERASE_II_PROMOTER | 457 | 37 | 5.17E-12 | 3.55E-10 |
| DNA_BINDING | 602 | 42 | 2.34E-11 | 1.57E-09 |
| REGULATION_OF_TRANSCRIPTIONDNA_DEPENDENT | 461 | 36 | 2.79E-11 | 1.77E-09 |
| PID_PDGFRBPATHWAY | 129 | 19 | 3.69E-11 | 2.29E-09 |
| POSITIVE_REGULATION_OF_CELLULAR_PROCESS | 668 | 44 | 5.08E-11 | 2.94E-09 |
| REGULATION_OF_RNA_METABOLIC_PROCESS | 471 | 36 | 5.10E-11 | 2.94E-09 |
| POST_TRANSLATIONAL_PROTEIN_MODIFICATION | 476 | 36 | 6.85E-11 | 3.87E-09 |
| POSITIVE_REGULATION_OF_BIOLOGICAL_PROCESS | 709 | 45 | 1.02E-10 | 5.66E-09 |
| GTPASE_REGULATOR_ACTIVITY | 125 | 18 | 1.77E-10 | 9.54E-09 |
| IMMUNE_RESPONSE | 235 | 24 | 2.76E-10 | 1.46E-08 |
| NERVOUS_SYSTEM_DEVELOPMENT | 385 | 31 | 3.12E-10 | 1.62E-08 |
| TRANSCRIPTION_FACTOR_ACTIVITY | 354 | 29 | 7.77E-10 | 3.95E-08 |
| REGULATION_OF_CELL_PROLIFERATION | 308 | 26 | 3.09E-09 | 1.54E-07 |
| IMMUNE_SYSTEM_PROCESS | 332 | 27 | 3.47E-09 | 1.69E-07 |
| RESPONSE_TO_VIRUS | 50 | 11 | 6.30E-09 | 3.02E-07 |
| ANATOMICAL_STRUCTURE_MORPHOGENESIS | 376 | 28 | 1.23E-08 | 5.69E-07 |
| RECEPTOR_BINDING | 377 | 28 | 1.30E-08 | 5.81E-07 |
| REGULATION_OF_SIGNAL_TRANSDUCTION | 222 | 21 | 1.39E-08 | 6.08E-07 |
| MULTI_ORGANISM_PROCESS | 165 | 18 | 1.64E-08 | 7.06E-07 |
| RESPONSE_TO_OTHER_ORGANISM | 83 | 13 | 2.11E-08 | 8.81E-07 |
| NUCLEOSIDE_TRIPHOSPHATASE_ACTIVITY | 212 | 20 | 3.19E-08 | 1.29E-06 |
| GUANYL_NUCLEOTIDE_EXCHANGE_FACTOR_ACTIVITY | 46 | 10 | 3.50E-08 | 1.37E-06 |
| PID_P53DOWNSTREAMPATHWAY | 137 | 16 | 3.87E-08 | 1.49E-06 |
| RESPONSE_TO_BIOTIC_STIMULUS | 120 | 15 | 4.10E-08 | 1.55E-06 |
| TISSUE_DEVELOPMENT | 138 | 16 | 4.29E-08 | 1.60E-06 |
| TRANSFERASE_ACTIVITY_TRANSFERRING_PHOSPHORUS_CONTAINING | 424 | 29 | 4.38E-08 | 1.61E-06 |
| TRANSFORMING_GROWTH_FACTOR_BETA_RECEPTOR_SIGNALING_PAT | 36 | 9 | 4.60E-08 | 1.67E-06 |
| PID_ERBB1_DOWNSTREAM_PATHWAY | 105 | 14 | 5.05E-08 | 1.81E-06 |
| PYROPHOSPHATASE_ACTIVITY | 226 | 20 | 9.23E-08 | 3.21E-06 |
| PROTEIN_KINASE_CASCADE | 293 | 23 | 9.39E-08 | 3.23E-06 |
| HYDROLASE_ACTIVITY_ACTING_ON_ACID_ANHYDRIDES | 228 | 20 | 1.07E-07 | 3.56E-06 |
| SUBSTRATE_SPECIFIC_TRANSPORTER_ACTIVITY | 392 | 27 | 1.10E-07 | 3.63E-06 |
| CELL_DEVELOPMENT | 577 | 34 | 1.16E-07 | 3.79E-06 |
| RESPONSE_TO_STRESS | 508 | 31 | 1.95E-07 | 6.20E-06 |
| ESTABLISHMENT_OF_LOCALIZATION | 870 | 43 | 3.57E-07 | 1.11E-05 |
| DEFENSE_RESPONSE | 270 | 21 | 3.91E-07 | 1.19E-05 |
| TRANSMEMBRANE_RECEPTOR_ACTIVITY | 418 | 27 | 3.94E-07 | 1.19E-05 |
| PID_EPOPATHWAY | 34 | 8 | 4.34E-07 | 1.29E-05 |
| KINASE_ACTIVITY | 369 | 25 | 4.37E-07 | 1.29E-05 |
| TRANSMEMBRANE_RECEPTOR_PROTEIN_SERINE_THREONINE_KINASE | 47 | 9 | 5.41E-07 | 1.56E-05 |
| PID_IL6_7PATHWAY | 47 | 9 | 5.41E-07 | 1.56E-05 |
| REGULATION_OF_MOLECULAR_FUNCTION | 324 | 23 | 5.59E-07 | 1.60E-05 |
| TRANSPORT | 795 | 40 | 5.83E-07 | 1.64E-05 |
| MOLECULAR_ADAPTOR_ACTIVITY | 49 | 9 | 7.86E-07 | 2.17E-05 |
| PID_GMCSF_PATHWAY | 37 | 8 | 8.72E-07 | 2.36E-05 |
| ACTIN_FILAMENT_BASED_PROCESS | 115 | 13 | 1.05E-06 | 2.78E-05 |
| REGULATION_OF_DEVELOPMENTAL_PROCESS | 440 | 27 | 1.06E-06 | 2.78E-05 |
| RESPONSE_TO_CHEMICAL_STIMULUS | 314 | 22 | 1.22E-06 | 3.16E-05 |
| PID_SMAD2_3NUCLEARPATHWAY | 82 | 11 | 1.28E-06 | 3.28E-05 |
| NEGATIVE_REGULATION_OF_CELL_PROLIFERATION | 156 | 15 | 1.29E-06 | 3.28E-05 |

Extended Table 6

| Gene Set Name | # Genes in ( | # Genes in ( | p-value | FDR q-value |
|---|---|---|---|---|
| ANATOMICAL_STRUCTURE_DEVELOPMENT | 1013 | 59 | 0.00E+00 | 0.00E+00 |
| INTRACELLULAR_SIGNALING_CASCADE | 667 | 47 | 0.00E+00 | 0.00E+00 |
| NEGATIVE_REGULATION_OF_BIOLOGICAL_PROCESS | 677 | 47 | 0.00E+00 | 0.00E+00 |
| NEGATIVE_REGULATION_OF_CELLULAR_PROCESS | 646 | 46 | 0.00E+00 | 0.00E+00 |
| SIGNAL_TRANSDUCTION | 1634 | 89 | 0.00E+00 | 0.00E+00 |
| MULTICELLULAR_ORGANISMAL_DEVELOPMENT | 1049 | 57 | 1.11E-16 | 4.03E-14 |
| SYSTEM_DEVELOPMENT | 861 | 50 | 1.11E-16 | 4.03E-14 |
| CELL_DEVELOPMENT | 577 | 40 | 4.44E-16 | 1.41E-13 |
| BIOPOLYMER_METABOLIC_PROCESS | 1684 | 68 | 3.23E-14 | 7.85E-12 |
| PROTEIN_KINASE_CASCADE | 293 | 27 | 3.40E-14 | 7.85E-12 |
| ORGAN_DEVELOPMENT | 571 | 36 | 2.29E-13 | 4.85E-11 |
| REGULATION_OF_DEVELOPMENTAL_PROCESS | 440 | 31 | 6.13E-13 | 1.20E-10 |
| ENZYME_REGULATOR_ACTIVITY | 323 | 26 | 2.30E-12 | 4.17E-10 |
| TRANSCRIPTION | 753 | 40 | 2.46E-12 | 4.17E-10 |
| KINASE_ACTIVITY | 369 | 27 | 8.04E-12 | 1.28E-09 |
| REGULATION_OF_TRANSCRIPTION | 566 | 33 | 1.87E-11 | 2.79E-09 |
| PHOSPHOTRANSFERASE_ACTIVITY_ALCOHOL_GROUP_AS_ACCEPTOR | 334 | 25 | 2.94E-11 | 4.15E-09 |
| TRANSFERASE_ACTIVITY_TRANSFERRING_PHOSPHORUS_CONTAINING_GROUPS | 424 | 28 | 3.70E-11 | 4.95E-09 |
| REGULATION_OF_NUCLEOBASENUCLEOSIDENUCLEOTIDE_AND_NUCLEIC_ACID_METAI | 618 | 34 | 4.39E-11 | 5.57E-09 |
| PROGRAMMED_CELL_DEATH | 432 | 28 | 5.70E-11 | 6.90E-09 |
| PROTEIN_METABOLIC_PROCESS | 1231 | 50 | 7.88E-11 | 9.10E-09 |
| POSITIVE_REGULATION_OF_CELLULAR_PROCESS | 668 | 35 | 8.54E-11 | 9.44E-09 |
| REGULATION_OF_MOLECULAR_FUNCTION | 324 | 24 | 9.05E-11 | 9.54E-09 |
| TRANSCRIPTION_DNA_DEPENDENT | 636 | 34 | 9.39E-11 | 9.54E-09 |
| RNA_BIOSYNTHETIC_PROCESS | 638 | 34 | 1.02E-10 | 9.60E-09 |
| POST_TRANSLATIONAL_PROTEIN_MODIFICATION | 476 | 29 | 1.12E-10 | 1.02E-08 |
| REGULATION_OF_CATALYTIC_ACTIVITY | 276 | 22 | 1.36E-10 | 1.17E-08 |
| REGULATION_OF_CELLULAR_METABOLIC_PROCESS | 787 | 38 | 1.38E-10 | 1.17E-08 |
| REGULATION_OF_METABOLIC_PROCESS | 799 | 38 | 2.12E-10 | 1.74E-08 |
| APOPTOSIS_GO | 431 | 27 | 2.67E-10 | 2.10E-08 |
| REGULATION_OF_PROGRAMMED_CELL_DEATH | 342 | 24 | 2.73E-10 | 2.10E-08 |
| NUCLEOBASENUCLEOSIDENUCLEOTIDE_AND_NUCLEIC_ACID_METABOLIC_PROCESS | 1244 | 49 | 3.44E-10 | 2.57E-08 |
| ANATOMICAL_STRUCTURE_MORPHOGENESIS | 376 | 25 | 3.59E-10 | 2.61E-08 |
| DNA_BINDING | 602 | 32 | 3.87E-10 | 2.73E-08 |
| POSITIVE_REGULATION_OF_BIOLOGICAL_PROCESS | 709 | 35 | 4.18E-10 | 2.87E-08 |
| ESTABLISHMENT_OF_LOCALIZATION | 870 | 39 | 6.63E-10 | 4.34E-08 |
| TRANSPORT | 795 | 37 | 6.65E-10 | 4.34E-08 |
| CELLULAR_PROTEIN_METABOLIC_PROCESS | 1117 | 45 | 9.26E-10 | 5.88E-08 |
| PROTEIN_AMINO_ACID_PHOSPHORYLATION | 279 | 21 | 1.01E-09 | 6.12E-08 |
| REGULATION_OF_TRANSCRIPTIONDNA_DEPENDENT | 461 | 27 | 1.16E-09 | 6.87E-08 |
| CELLULAR_MACROMOLECULE_METABOLIC_PROCESS | 1131 | 45 | 1.36E-09 | 7.79E-08 |
| REGULATION_OF_APOPTOSIS | 341 | 23 | 1.38E-09 | 7.79E-08 |
| PHOSPHORYLATION | 313 | 22 | 1.46E-09 | 8.03E-08 |
| PROTEIN_KINASE_ACTIVITY | 285 | 21 | 1.48E-09 | 8.03E-08 |
| REGULATION_OF_GENE_EXPRESSION | 673 | 33 | 1.57E-09 | 8.29E-08 |
| REGULATION_OF_RNA_METABOLIC_PROCESS | 471 | 27 | 1.84E-09 | 9.37E-08 |
| PROTEIN_MODIFICATION_PROCESS | 631 | 31 | 4.70E-09 | 2.34E-07 |
| REGULATION_OF_SIGNAL_TRANSDUCTION | 222 | 18 | 4.92E-09 | 2.40E-07 |
| NEGATIVE_REGULATION_OF_DEVELOPMENTAL_PROCESS | 197 | 17 | 5.14E-09 | 2.47E-07 |
| TRANSCRIPTION_REPRESSOR_ACTIVITY | 152 | 15 | 6.63E-09 | 3.06E-07 |
| RESPONSE_TO_EXTERNAL_STIMULUS | 312 | 21 | 7.40E-09 | 3.36E-07 |
| REGULATION_OF_PROTEIN_KINASE_ACTIVITY | 155 | 15 | 8.67E-09 | 3.86E-07 |
| BIOPOLYMER_MODIFICATION | 650 | 31 | 9.34E-09 | 4.09E-07 |
| RNA_METABOLIC_PROCESS | 841 | 36 | 9.97E-09 | 4.29E-07 |
| REGULATION_OF_KINASE_ACTIVITY | 157 | 15 | 1.03E-08 | 4.37E-07 |
| CYTOSKELETAL_PROTEIN_BINDING | 159 | 15 | 1.23E-08 | 4.95E-07 |
| REGULATION_OF_TRANSFERASE_ACTIVITY | 161 | 15 | 1.45E-08 | 5.77E-07 |
| REGULATION_OF_BIOLOGICAL_QUALITY | 419 | 24 | 1.48E-08 | 5.80E-07 |
| NEGATIVE_REGULATION_OF_TRANSCRIPTION | 188 | 16 | 1.76E-08 | 6.68E-07 |
| POSITIVE_REGULATION_OF_CATALYTIC_ACTIVITY | 165 | 15 | 2.03E-08 | 7.57E-07 |
| GTPASE_REGULATOR_ACTIVITY | 125 | 13 | 3.59E-08 | 1.30E-06 |
| RESPONSE_TO_STRESS | 508 | 26 | 3.66E-08 | 1.31E-06 |
| MAPKKK_CASCADE_GO_0000165 | 104 | 12 | 3.74E-08 | 1.32E-06 |
| RECEPTOR_ACTIVITY | 583 | 28 | 4.20E-08 | 1.46E-06 |
| REGULATION_OF_MAP_KINASE_ACTIVITY | 67 | 10 | 4.26E-08 | 1.46E-06 |
| CALMODULIN_BINDING | 25 | 7 | 4.79E-08 | 1.62E-06 |
| TRANSCRIPTION_FACTOR_ACTIVITY | 354 | 21 | 6.51E-08 | 2.15E-06 |
| CELL_SURFACE_RECEPTOR_LINKED_SIGNAL_TRANSDUCTION_GO_0007166 | 641 | 29 | 8.62E-08 | 2.81E-06 |
| NEGATIVE_REGULATION_OF_NUCLEOBASENUCLEOSIDENUCLEOTIDE_AND_NUCLEIC_/ | 211 | 16 | 8.87E-08 | 2.85E-06 |

| | | | | |
|---|---|---|---|---|
| SULFOTRANSFERASE_ACTIVITY | 28 | 7 | 1.14E-07 | 3.61E-06 |
| ACTIN_BINDING | 76 | 10 | 1.46E-07 | 4.42E-06 |
| CELL_PROLIFERATION_GO_0008283 | 513 | 25 | 1.69E-07 | 4.99E-06 |
| TRANSMEMBRANE_TRANSPORTER_ACTIVITY | 375 | 21 | 1.71E-07 | 4.99E-06 |
| RESPONSE_TO_CHEMICAL_STIMULUS | 314 | 19 | 2.02E-07 | 5.83E-06 |
| NERVOUS_SYSTEM_DEVELOPMENT | 385 | 21 | 2.64E-07 | 7.53E-06 |
| TRANSFERASE_ACTIVITY_TRANSFERRING_SULFUR_CONTAINING_GROUPS | 32 | 7 | 3.07E-07 | 8.57E-06 |
| TRANSMEMBRANE_RECEPTOR_PROTEIN_TYROSINE_KINASE_SIGNALING_PATHWAY | 83 | 10 | 3.40E-07 | 9.29E-06 |
| PID_HIF1_TFPATHWAY | 66 | 9 | 4.51E-07 | 1.21E-05 |
| POSITIVE_REGULATION_OF_TRANSFERASE_ACTIVITY | 86 | 10 | 4.76E-07 | 1.26E-05 |
| POSITIVE_REGULATION_OF_DEVELOPMENTAL_PROCESS | 218 | 15 | 7.73E-07 | 1.99E-05 |
| PID_P53DOWNSTREAMPATHWAY | 137 | 12 | 7.83E-07 | 1.99E-05 |

Extended Table 7

| Database_Pathway Annotation | # Genes in Gene Set | # Genes in O | p-value | FDR q-value |
|---|---|---|---|---|
| ***Nudix Dependent Genes*** | | | | |
| REACTOME_IMMUNE_SYSTEM | 933 | 92 | 0.00E+00 | 0.00E+00 |
| REACTOME_CYTOKINE_SIGNALING_IN_IMMUNE_SYSTEM | 270 | 49 | 0.00E+00 | 0.00E+00 |
| REACTOME_INTERFERON_SIGNALING | 159 | 38 | 0.00E+00 | 0.00E+00 |
| REACTOME_INTERFERON_ALPHA_BETA_SIGNALING | 64 | 24 | 0.00E+00 | 0.00E+00 |
| KEGG_PATHWAYS_IN_CANCER | 328 | 34 | 3.57E-14 | 3.63E-12 |
| REACTOME_INTERFERON_GAMMA_SIGNALING | 63 | 15 | 3.24E-12 | 2.28E-10 |
| KEGG_CYTOKINE_CYTOKINE_RECEPTOR_INTERACTION | 267 | 27 | 2.69E-11 | 1.75E-09 |
| REACTOME_ADAPTIVE_IMMUNE_SYSTEM | 539 | 39 | 4.15E-11 | 2.51E-09 |
| REACTOME_DEVELOPMENTAL_BIOLOGY | 396 | 29 | 9.85E-09 | 4.63E-07 |
| REACTOME_ANTIVIRAL_MECHANISM_BY_IFN_STIMULATED_GENES | 66 | 12 | 1.29E-08 | 5.81E-07 |
| KEGG_MAPK_SIGNALING_PATHWAY | 267 | 23 | 1.70E-08 | 7.20E-07 |
| REACTOME_AXON_GUIDANCE | 251 | 22 | 2.53E-08 | 1.04E-06 |
| KEGG_TOLL_LIKE_RECEPTOR_SIGNALING_PATHWAY | 102 | 14 | 3.47E-08 | 1.37E-06 |
| REACTOME_TRANSMEMBRANE_TRANSPORT_OF_SMALL_MOLECULES | 413 | 28 | 8.95E-08 | 3.16E-06 |
| REACTOME_HEMOSTASIS | 466 | 30 | 9.75E-08 | 3.30E-06 |
| REACTOME_INNATE_IMMUNE_SYSTEM | 279 | 22 | 1.64E-07 | 5.28E-06 |
| REACTOME_SIGNALING_BY_FGFR_MUTANTS | 44 | 9 | 2.98E-07 | 9.35E-06 |
| KEGG_CHEMOKINE_SIGNALING_PATHWAY | 190 | 17 | 7.16E-07 | 2.00E-05 |
| KEGG_FC_EPSILON_RI_SIGNALING_PATHWAY | 79 | 11 | 8.73E-07 | 2.36E-05 |
| KEGG_REGULATION_OF_ACTIN_CYTOSKELETON | 216 | 18 | 9.65E-07 | 2.58E-05 |
| ***Nudix Independent Genes*** | | | | |
| KEGG_PATHWAYS_IN_CANCER | 328 | 30 | 1.55E-15 | 4.39E-13 |
| REACTOME_SIGNALING_BY_RHO_GTPASES | 113 | 15 | 1.01E-10 | 9.60E-09 |
| REACTOME_SIGNALING_BY_GPCR | 920 | 40 | 9.59E-10 | 5.94E-08 |
| REACTOME_PLATELET_ACTIVATION_SIGNALING_AND_AGGREGATION | 208 | 18 | 1.75E-09 | 9.06E-08 |
| KEGG_PROSTATE_CANCER | 89 | 12 | 6.25E-09 | 2.94E-07 |
| REACTOME_TRANSMEMBRANE_TRANSPORT_OF_SMALL_MOLECULES | 413 | 24 | 1.13E-08 | 4.69E-07 |
| KEGG_ENDOCYTOSIS | 183 | 16 | 1.20E-08 | 4.92E-07 |
| KEGG_MAPK_SIGNALING_PATHWAY | 267 | 19 | 1.57E-08 | 6.03E-07 |
| REACTOME_HEMOSTASIS | 466 | 25 | 2.70E-08 | 9.94E-07 |
| KEGG_AXON_GUIDANCE | 129 | 13 | 5.23E-08 | 1.75E-06 |
| KEGG_REGULATION_OF_ACTIN_CYTOSKELETON | 216 | 16 | 1.22E-07 | 3.84E-06 |
| REACTOME_SIGNALLING_BY_NGF | 217 | 16 | 1.30E-07 | 4.04E-06 |
| KEGG_ALDOSTERONE_REGULATED_SODIUM_REABSORPTION | 42 | 8 | 1.36E-07 | 4.17E-06 |
| REACTOME_NRAGE_SIGNALS_DEATH_THROUGH_JNK | 43 | 8 | 1.65E-07 | 4.94E-06 |
| KEGG_FOCAL_ADHESION | 201 | 15 | 2.74E-07 | 7.75E-06 |
| KEGG_WNT_SIGNALING_PATHWAY | 151 | 13 | 3.34E-07 | 9.21E-06 |
| KEGG_CYTOKINE_CYTOKINE_RECEPTOR_INTERACTION | 267 | 17 | 4.32E-07 | 1.17E-05 |
| KEGG_ADIPOCYTOKINE_SIGNALING_PATHWAY | 67 | 9 | 5.14E-07 | 1.35E-05 |
| KEGG_PANCREATIC_CANCER | 70 | 9 | 7.53E-07 | 1.95E-05 |
